# Same strain, two genomes: Creating a consensus circular genome of *Myxococcus xanthus* DZ2 revealed the diversity of a large polyploid prophage region within the close-related myxobacterial strains

**DOI:** 10.1101/2024.12.30.630742

**Authors:** Utkarsha Mahanta, Gaurav Sharma

**Affiliations:** Department of Biotechnology, Indian Institute of Technology Hyderabad, Sangareddy, Telangana, India

**Keywords:** Evolution, genome variation, homology, genome circularization, genome rearrangement, genome sequencing, kin discrimination, microbial diversity, Comparative genomics

## Abstract

One of the well-known myxobacterial model organism, i.e., *Myxococcus xanthus* DZ2 strain has three reported assemblies of which two complete assemblies (MxDZ2_Tam and MxDZ2_Nan) have been reported in the last three years for the same strain procured from same culture collection stock. Raw assemblies of this same strain were reported circular and differed by 6.4 kb, raising questions about their accuracy. Therefore, our computational analysis, removing duplicate ends, aligning genomes to the origin of replication, and circularization, revealed a minimal size difference of only 32 bp, with MxDZ2_Tam being slightly larger. Further comparative analysis identified 40 sequence variations: 38 indels and 2 substitutions, with 24 variations concerning 18 coding genes. Eight of these 24 variations triggered frameshift mutations, seven of them encountered a premature stop codon followed by a second ORF and the remaining three had a premature stop codon followed by a non-coding region. 78% of these variations were detected in the regions which had repeated bases, especially guanine and cytosine, which is one of the well-known sequencing limitations. Although PacBio HiFi technology, used for the complete assemblies, boasts a low error rate, it remains higher than the 454-platform used for the earlier MxDZ2_Kirby assembly. Using MxDZ2_Kirby as a reference, we determined that most variations were specific to MxDZ2_Nan, while MxDZ2_Tam showed greater consistency with MxDZ2_Kirby. Based on these findings, we constructed a consensus ‘truly circular’ genome for the same *M. xanthus* DZ2 strain. Additionally, we monitored divergence between *M. xanthus* DZ2 and its close relative *M. xanthus* DK1622 in the number of Mx-alpha regions, which are linked to interspecies and intraspecies antagonism via the toxin gene sitA. Further comparative genome analysis across 61 myxobacteria revealed the presence of Mx-alpha regions in five other organisms, though only *M. xanthus* DZ2 and DZF1 contained all three Mx-alpha regions. Overall, this study underscores the need for meticulous validation of sequencing-based genome assemblies and their variations along with highlighting the genomic diversity within *M. xanthus* strains based on a large polyploid prophage region, known as Mx-alpha.

**Importance:** Sequencing an organism is an efficient way to investigate its physiology and function. Through this study, we focused on resolving differences between two genome assemblies of the same strain, i.e., *M. xanthus* DZ2, procured from the same culture collection and assembled via two different organisations. We identified 40 differences between these two, which led to inconsistencies in how some proteins were predicted and annotated. By comparing these assemblies with a third draft genome, we created a more accurate and reliable consensus ‘truly circular’ genome for this strain. We also investigated a particular region of the genome, known as Mx Alpha, that contains genes for toxin-antitoxin systems, which play a role in helping the bacteria recognize and interact with related strains. These findings provide new insights into the diversity of this region in *Myxococcus* and its phylum Myxococcota relatives, shedding light on how these bacteria interact within their communities.

## Introduction

Organisms keep evolving incessantly in response to ever-changing environmental conditions through a compendium of genomic mutations. For each such mutation to persist in the genotype across generations, an equilibrium is required between the robustness and evolvability of that mutation (1). Mutations, whether synonymous or non-synonymous, can generate diverse variations. Long-term evolution experiment (LTEE) with *Escherichia coli* has already revealed several genome rearrangements and variations/SNPs over time across generations, with the study now surpassing 80,000 generations (2). In the similar line, sequencing two or more strains having the same origin but separated by different generations at different places may reveal how significantly each of those strains has evolved and what type of variations each strain type has gained to stay fit or naturally selected (3). However, in this current sequencing and high-throughput era, researchers must be aware of several such differences majorly derived from technological limitations, instead of the real mutations.

Most of the next-generation (including Illumina and 454) and third generation (PacBio High-Fidelity (HiFi) and Nanopore) sequencing platforms offer a high read accuracy or fidelity while sequencing any genome. Even after using stringent high-quality sequencing reads and similar assembly processes, the final genome assembly of any organism might have a few or more discrepancies. Such discrepancies or variations within even a complete genome of an organism sequenced by one research team cannot be identified unless another team decides to sequence the exact same organism procured from the same source as the first research team.

Myxobacteria are a group of ubiquitously present gram-negative predatory bacteria which were previously classified under the class of Deltaproteobacteria based on 16s rRNA and morphology. However, with the recent advancements in taxonomy and classification, these bacteria form a new separate phylum under the name of Myxococcota with a class named Myxococcia and a subclass called Polyangia (4, 5). *Myxococcus xanthus* is the model organism of this group of bacteria which belongs to the class of Myxococcia, order Myxococcales, suborder Cystobacterineae and family Myxococcaceae. This organism (Beebe isolate) was first classified and reported in the year 1941 by J. M. Beebe (6), and was relocated to UC Berkeley to be further maintained in Roger Stanier’s strain collection as strain FB [**Figure-1**] (7). FB strain was further derived into the well-known model organism of the group, DK 1622 after going through several in-between mutants such as DK101 (further derived into another well-known strain, DZF1), DK320, DK1217, and then DK 1622 (7). Along with FB strain, another strain, DZ2 was stored independently by David Zusman (abbreviated as DZ) in his lab, away from the Stanier collection. It is not yet clear which strain is the direct ancestor of DZ2 and FB. One major similarity between DZ2 and DK 1622 is their A^+^S^+^ nature suggesting that they display both A and S motility, whereas other in-between mutants have a range of A^+/-^S^+/-^ activities. Overall, from a long time, *M. xanthus* DZ2 is being used by several myxobacterial biologists as a model organism to study diverse myxobacterial physiology.

Considering its importance in understanding myxobacterial physiology, it was first sequenced in 2013 by John Kirby’s group at the University of Iowa, USA, as a draft assembly (GCF_000278585.1) consisting of 87 contigs (8) using 454 sequencing platform followed by assembly using Newbler v. 11/12/2012. Eight years later in 2021, the first complete genome (GCF_018517205.1) was reported (9) by Tam Mignot’s lab at CNRS, France and in the subsequent year another assembly (GCF_020827275.1) of its complete genome was published (10) by Beiyan Nan’s lab at Texas A&M. Their strain description reveals that both Tam’s and Nan’s lab isolated the DNA for genome sequencing from DZ2 cells, derived from the frozen stock from the Roger Stanier collection at UC Berkeley, suggesting both must be similar. Overall, all these three strains have the same origin from David Zusman’s lab at UC Berkeley, but they have been sub-cultured, sequenced and assembled at three different places (Iowa, USA; Marseille, France; Texas, USA) at three different times (2013; 2021; 2022). Two out of these three assemblies, i.e., GCF_018517205.1 and GCF_020827275.1, represent complete genome assemblies of the same strain, but their genome sizes are reported to differ by 6.4 kb and several other genome statistics also do not resemble each other [**Figure-1**].

Moreover, it has been reported earlier that DZ2 and DK 1622 are different from each other in the context of a Mx alpha region (∼200-kb prophage-like element), which got deleted in DK 1622 leading to the latter having only the Mx alpha 3 region whereas the former has all three of them (7). Some of the genes in Mx alpha unit could be found in all the repeats whereas some are unique to those units. One such unique gene operon is SitBAI which is involved in antagonism where the toxin protein SitA is transferred through outer membrane exchange and the species or strains lacking this SitI protein, which provides immunity, is killed (7, 11, 12). In this case, DZ2 can readily kill DK 1622 because the latter lacks *sitI*1 and *sitI*2 genes and hence has no immunity against *sitA*1 and *sitA*2 toxins. However, DZ2 can survive the attack of SitA3 from DK 1622 as it contains its immunity gene *sitI*3.

Therefore, this study aims to find out how and why these two complete genome assemblies GCF_018517205.1 and GCF_020827275.1 are different from each other, what kind of variations are present in each of them, how those discrepancies can be of concern for experimental and computational researchers and can we provide a consensus genome assembly of this well-known myxobacterial model organism, *Myxococcus xanthus* DZ2. We further investigated the Mx alpha region distribution and classification across some selected members of the myxobacteria, to get an idea of how prevalent the occurrence and enumeration of this prophage region is within the phylum Myxococcota.

## Methodology

### Data source

Three assemblies of the same *Myxococcus xanthus* DZ2 strain are currently available in open access, of which the first one is a draft, <u>GCF_000278585.1</u> (MxDZ2_Kirby) published in 2013 using 454 sequencing technology (8) and the other two are complete: <u>GCF_018517205.1</u> (MxDZ2_Tam) published in 2021 using PacBio high-fidelity (HiFi) technology (9) and <u>GCF_020827275.1</u> (MxDZ2_Nan), published in 2022 using DNBSEQ and PacBio HiFi sequencing technologies (10). The genomic and proteomic content of these assemblies were downloaded from the NCBI RefSeq Database (https://www.ncbi.nlm.nih.gov/refseq/) for analysis. In this manuscript, the tags MxDZ2_Kirby, MxDZ2_Tam and MxDZ2_Nan will be used to refer the assemblies GCF_000278585.1, GCF_018517205.1 and GCF_020827275.1, respectively.

For gene/CDS names, we have used locus_tag information in a bit modified manner to make it convenient for readers. In the standard manner, locus tags start with an organism tag prefix followed by RS numbers. In this research, it is MXDZ, JTM82, and K1515 for MxDZ2_Kirby, MxDZ2_Tam and MxDZ2_Nan respectively followed by RS numbers; however, we have used Kirby, Tam, and Nan as respective locus tag prefix followed by their RS numbers.

### Alignment

Various alignment tools like Mauve v2.4.0 (13), Pygenomeviz 0.4.4 (https://pypi.org/project/pygenomeviz/), MUMer 4.0.0rc1 (14) and MAFFT v7.526 (15) with their default parameters were used to align the assembled genomes against each other. Mauve progressively aligns multiple genomes to detect evolutionary events such as rearrangement and inversion whereas MUMer with the help of Maximal Unique Matches (MUMs) provides variations information along with the alignment. Pygenomeviz was used along Mauve for visualization of the genomic content of the three assemblies with respect to each other [**Supplementary Figure-1**] and multiple sequence alignment of the nucleotide and protein sequences was done via MAFFT using the auto parameter [**Supplementary File-1**].

### Homology detection

To detect the homologous and repeated proteins within the three assemblies, the amino acid and nucleotide sequence of the genes were subjected to blast v2.13.0 (16) via blastp and blastn respectively with an e-value of 10^-5^.

### Circularization

On detection of repeated proteins at the beginning and end of the genomes, the redundant proteins were removed keeping only the protein with longer length for further analysis [**Figure-2**] and subsequently genome statistics were calculated for all three assemblies [**Figure-1**]. Based on dnaA-dnaN-gyrB conserved region along with GC skew and comparative genomics, the origin of replication of both genomes was found using Ori-Finder 2022 (17).

**Figure-1:**
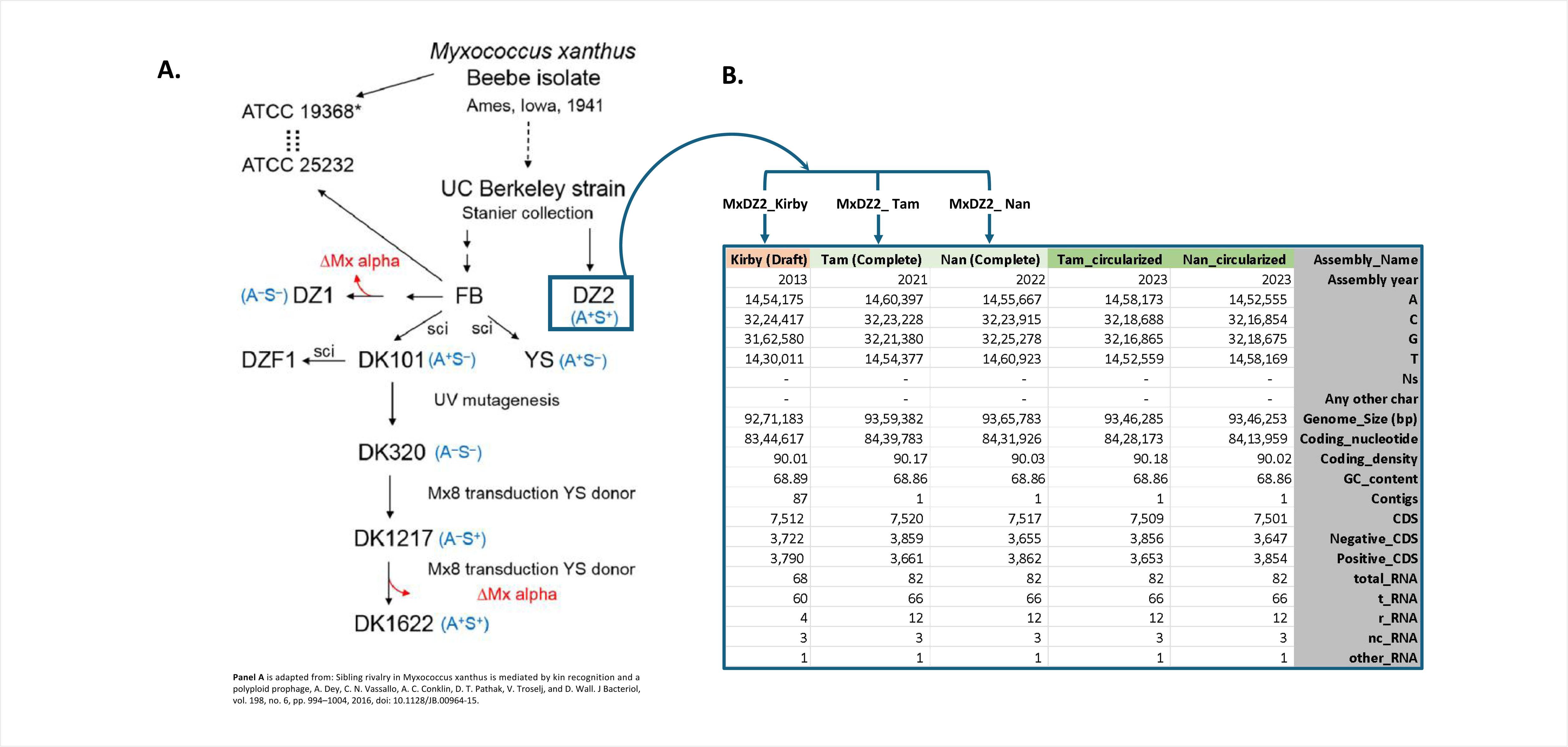
**(a)** Pictorial description of the ancestry of various strains of *M. xanthus* adapted from Dey A et al (7), showcasing the putative evolution of lab cultures belonging to *M. xanthus* species. **(b)** Genome statistics of all three *M. xanthus* DZ2 assemblies before and after circularization.

### Realignment

Neither MxDZ2_Tam nor MxDZ2_Nan genomic sequence starts from the *oriC*, so both were rearranged in a manner that the segment of the genome preceding the *oriC* was truncated and attached to the end of the sequence for both the genomes. Further genomic alignment between MxDZ2_Tam and MxDZ2_Nan reveals that these two genomes are reverse complement of each other [**Figure-2, Supplementary Figure-1**].

**Figure-2:**
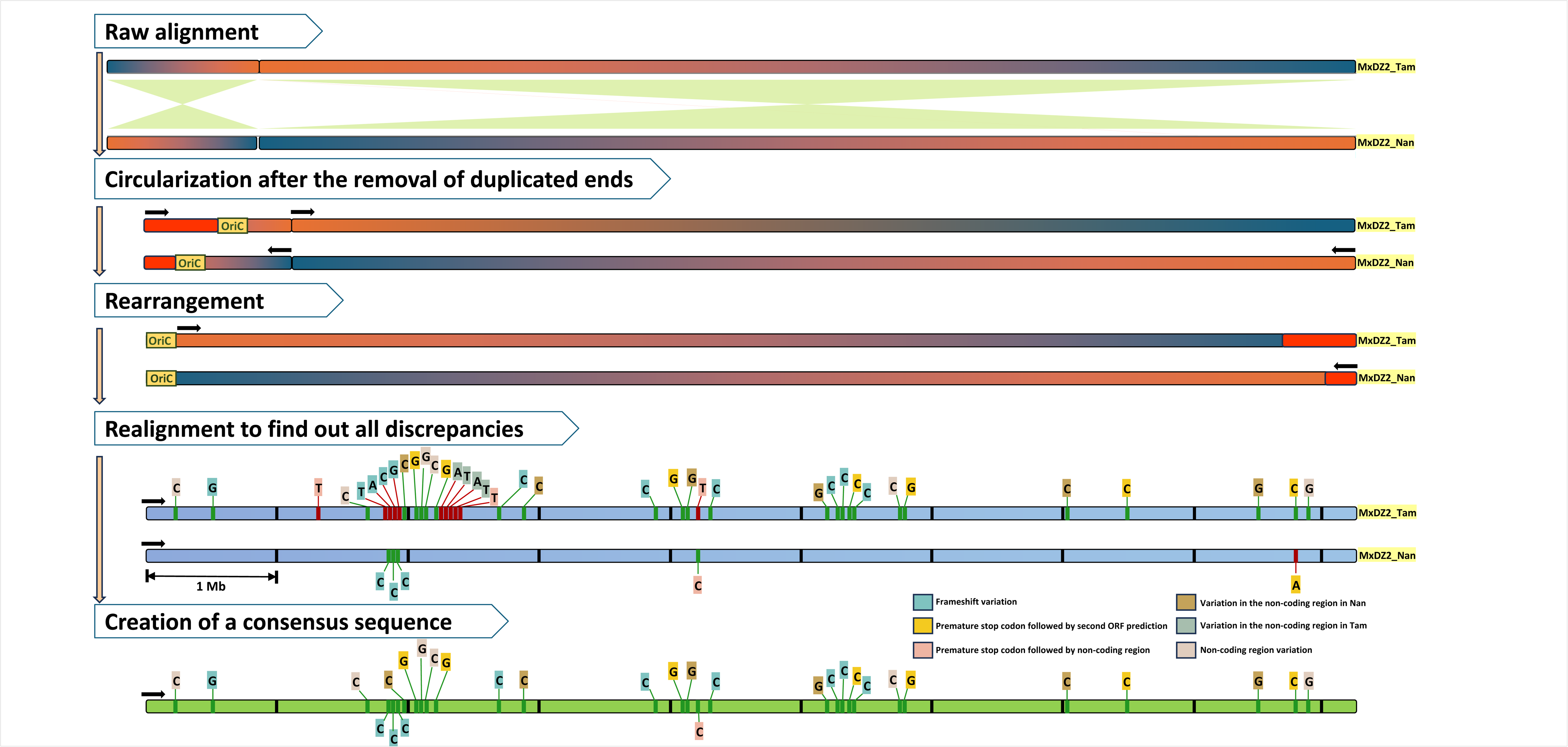
A pictorial methodology showing the comparison of MxDZ2_Nan and MxDZ2_Tam assemblies to reveal their raw and refined alignments along with the variations detected, which have been further listed in Supplementary Table-1. This whole process was employed to create a consensus sequence of MxDZ2.

### Mutational analysis

Nucleotide discrepancies were identified between the two genomes (MxDZ2_Tam and MxDZ2_Nan) using NUCmer (NUCleotide MUMmer) 4.0.0rc1 (14) with the default parameters and subsequently the genes in which these mutations are observed were aligned to each other along with their respective homologous genes in MxDZ2_Kirby using MAFFT [**Supplementary Table-1, Supplementary File-1**].

### Consensus sequence

MxDZ2_Kirby genome has been assembled using 454 reads which have more coverage and therefore, high accuracy. On occasions, where we identified nucleotide discrepancies, we used MxDZ2_Kirby assembly as a reference to find the correct nucleotide. Such correct nucleotides at all identified discrepancy sites were used to build a consensus sequence for *M. xanthus* DZ2, which shall from now on be referred to as MxDZ2_Consensus [**Figure-2**].

### Comparison of MxDZ2_Consensus with other strains of *M. xanthus*

MxDZ2_Consensus was compared to *M. xanthus* DK 1622, which is a complete genome published in the year 2006 (18) and with *M. xanthus* DZF1, which is a draft genome reported in 2013 (19). In *M. xanthus* DK 1622 a genomic region of ∼200kb deletion was detected which corresponds to the two missing Mx alpha units as previously reported (7).

### Detection of Mx alpha units across myxobacteria

Mx alpha units orthologs were searched across *M. xanthus* strains DZ2, DK 1622 and DZF1 along with a selected dataset of 63 myxobacteria encompassing at least one member from all known families using Proteinortho6 (20) with the default parameters. A total of 235 single copy orthologs were detected in this dataset along with *Bdellovibrio bacteriovorus* as an outgroup, which were subjected to multiple sequence alignment using MUSCLE5 (21) and subsequently concatenated phylogeny was built using IQ-TREE version 1.6.12 (22) with LG+F+I+G4 model. Another four core Mx alpha gene (single gene and concatenated) phylogenies consisting of orthologs of MXAN_RS08880, MXAN_RS08910, MXAN_RS09000 and MXAN_RS09080 were built using the same method described previously with LG+F+G4 model for the concatenated tree. The visualization of these phylogenies was done using iTOL v6 (23).

### Presence of polymorphic toxin families within the dataset

Each Mx alpha region is associated with a toxin protein SitA, or swarm inhibition toxin A which is responsible for kin discrimination. SitA1, SitA2 and SitA3 proteins are flanked by an accessory protein SitB and have a downstream SitI immunity gene (12). SitA is a signal peptide with N-terminal lipoprotein and has a C-terminal nuclease domain which was detected using SignalP6 (24) and HHpred 3.3.0 (25) search against PDB (26), Pfam (27) and NCBI CDD (28) databases respectively using default parameters. Also as reported by Vassallo C et al (11), myxobacterial genome encodes for six such polymorphic toxin families, with SitA1 and SitA2 belonging to one family because of high sequence identity and the remaining SitA3 to SitA7 belonging to individual toxin families. The presence of these six families of toxin proteins in our dataset was detected by homology search against the database consisting of these SitA proteins as reported previously (11).

### Genomic island and prophage region detection

All *M. xanthus* strains considered in this study along with those complete assembly myxobacterial genomes with considerable counts of Mx alpha homologs were subjected to genomic island detection using IslandViewer 4 (29) and subsequent prophage region were detected using PhiSpy version 4.2.21 (30).

## Results

### Both complete assemblies are significantly different in their genomic and proteomic content

The first draft genome assembly of *Myxococcus xanthus* DZ2, MxDZ2_Kirby, had 87 contigs with a total of 9,271,183 base pairs of which 8,344,617 (90.01%) encoded for 7,512 proteins and 68 RNAs. Eight years later the first complete genome assembly MxDZ2_Tam was published, which possesses 7,520 genes along with 82 RNAs encoded by 8,439,783 (90.17%) nucleotides out of a total of 9,359,382 base pairs. Another complete genome assembly MxDZ2_Nan was published in the subsequent year and had 8,431,926 (90.03%) out of 9,365,783 base pairs encoding for 7,517 proteins and 82 RNAs [**Figure-1**]. MxDZ2_Kirby is ∼90kb smaller in genome size as compared to both complete assemblies. Even both the complete assemblies differ by 6,401 base pairs with MxDZ2_Nan being the larger one. Mauve alignment revealed that both complete assemblies have large-scale genomic rearrangements as compared to each other showcasing inversions [**Figure-2**]. Moreover, both genomes also show rearrangements as compared to *M. xanthus* DK 1622, the genome of their nearest relative [**Supplementary Figure-1, 2**]. Alignment of both MxDZ2 assemblies with MxDK1622 revealed that the MxDZ2_Tam is more syntenic and ordered as compared to MxDZ2_Nan assembly [**Supplementary Figure-1**].

### Duplicated proteins at N and C terminals suggest that both assemblies are not circular as reported earlier

Most myxobacteria have been reported to have a single circular chromosome, especially the close relative of MxDZ2, i.e., *M. xanthus* DK 1622 (18). Our analysis disclosed that although MxDZ2_Tam and MxDZ2_Nan are complete assemblies, they do not represent a circular genome. Homology analysis detected several duplicated proteins present in an order at the beginning and end of the genomic sequences within each complete assembly. From MxDZ2_Tam blastp analysis, it was found that proteins 1-10 (RS37890 - RS00050) were homologous to proteins 7510-7519 (RS37835 - RS37880), respectively. However, protein sequence similarity could not be detected between protein 11 (RS00050) and protein 7520 (RS37885) but their nucleotide sequence similarity was unrevealed by blastn analysis. Similarly, in the case of MxDZ2_Nan assembly, blastp analysis detected proteins 1 to 14 (RS00005 - RS00070) had sequence similarity with proteins 7502 to 7515 (RS37900 - RS37965) respectively and protein 15 (RS00075) exhibiting sequence similarity with protein 7516 and 7517 (RS37970 and RS37975). These redundant proteins were removed while keeping the longer protein sequences followed by truncation for both MxDZ2_Tam and MxDZ2_Nan assemblies resulting in 7,509 and 7,501 encoded proteins and their genome sizes decreased by 13,097 (0.13%) and 19,530 (0.21%) base pairs, respectively [**Figure-1**]. In the raw assemblies, MxDZ2_Nan was 6.4kb larger than MxDZ2_Tam; however, after circularization, this study revealed that MxDZ2_Tam is only 32 bp larger than MxDZ2_Nan, showcasing the extent of more duplicated ends in MxDZ2_Nan, that has been truncated now.

### Whole genome alignment revealed the requirement of reverse complementation of one of the complete assemblies

Origin of replication (*oriC*) sites were detected at 1,023,040 - 1,023,346 bp and 141,035 - 141,341 bp for MxDZ2_Tam and MxDZ2_Nan, respectively. In order to start the genome sequence from *oriC*, 1-1,023,039 nt and 1-141,034 nt were truncated from the beginning of the genome sequence of MxDZ2_Tam and MxDZ2_Nan respectively and added to the end of the sequence. The rearranged genomic sequences of MxDZ2_Tam and MxDZ2_Nan were aligned to each other revealing that they were reverse complement to each other [**Figure-2, Supplementary Figure-1**].

### Forty variations make rearranged circularized complete assemblies different by 32 bps

Based on the realignment of both circularized complete assemblies, 40 variations were detected of which 38 were indels and two were substitutions. Within 35 out of 38 indels, MxDZ2_Nan assembly had a missing base whereas in the remaining three indels, MxDZ2_Nan had an additional cytosine base as compared to MxDZ2_Tam. The only two identified substitutions were transversions (C-A/T-G) [**Supplementary Table-1**].

Of the forty identified variations, twenty-four were present in the coding region in both assemblies, with two being substitutions and twenty-two being indels [**Supplementary Table-1**]. Out of these twenty-two indels, in seven cases, one indel per ORF led to a <u>frameshift mutation</u> resulting in different amino acids across the protein length, however, the reported proteins are of equal lengths with protein in one of the assemblies having multiple stop codons, which is because of wrong translation. Moreover, in one case, we found seven indels in one single ORF, causing several frameshift mutations across the protein length, but the gene calling, and annotation have predicted almost the same length for both assemblies with one having multiple stop codons in between. These results have been described in detail in **Supplementary File-1** depicting that protein alignments in all frameshift cases showcase little to no similarity even though they are of equal length because of the shift in the reading frames induced by the indels. In one example, Nan_RS36290 and Tam_RS06590 are 926 and 927 base pairs with eight and seven guanine base repeats, respectively. The predicted protein length (308 amino acids) is similar but multiple stop codons are present in MxDZ2_Nan protein due to change in its reading frame. MxDZ2_Kirby and MxDZ2_Tam genes are completely identical.

Besides these frameshift mutations, seven cases were identified in which a mutation has led to <u>frameshift mutation in either of the complete assemblies resulting in the reading frame</u> <u>reaching a stop codon</u> early and splitting up of a single protein into two proteins, thus predicting short proteins in one of the assemblies [**Supplementary File-1]**. In one of the cases, Nan_RS29755 and Tam_RS13120 share a point mutation in which Tam_RS13120 has four cytosine (complement) base repeats which is three in Nan_RS29755 leading to a frameshift mutation resulting in early reading of stop codon in Nan_RS29755. Tam_RS13120 is a protein of length 219 which has shared homology with two proteins in MxDZ2_Nan assembly: Nan_RS29755 of length 151 and Nan_RS29760 of length 68. MSA with MxDZ2_Kirby revealed that Kirby_RS0202735 also has four guanine repeats, similar to MxDZ2_Tam assembly.

In the above-mentioned fifteen cases, it was vividly observed that 14 of them have occurred where a particular base especially cytosine or guanine was present multiple times at that position of the genome. Also, alignment with homologs in MxDZ2_Kirby displayed that in 14 out of the 15 cases, MxDZ2_Kirby was exactly like MxDZ2_Tam.

Also, there are three instances in which a mutation has led to <u>frameshift mutations causing</u> <u>the reading frame to reach a stop codon early resulting in a shorter protein</u> and the remaining nucleotide becoming a part of the non-coding region because of the lack of a start codon in the rest of the part of the protein. Consequently, there is a huge difference in the length of the proteins encoded by the two assemblies. For example, in first case, Nan_RS32920 and Tam_RS09970 share a point mutation in which Tam_RS09970 has a thymine base which is absent in Nan_RS32920 leading to a frameshift mutation and early stop codon reading in Tam_RS09970 making it shorter by 314 amino acids. Nan_RS32920 is of 2255 bases encoding 751 amino acids and Tam_RS09970 is 1314 bases in length translating into 437 amino acids where the remaining 942 bases become a part of the non-coding region. Kirby_RS0217305 does not have adenine (complement), like MxDZ2_Nan assembly. Unlike other cases in which ∼93% (14 out of 15) of them MxDZ2_Kirby was like MxDZ2_Tam, here in the above two out of three cases MxDZ2_Kirby was similar to MxDZ2_Nan. Moreover, we also encountered a special case in which MxDZ2_Kirby was similar to neither MxDZ2_Tam nor MxDZ2_Nan.

Additionally, there were 16 variations detected in the non-coding region of the genome(s), of which six were in MxDZ2_Nan non-coding region, four in MxDZ2_Tam non-coding region and six in non-coding region of both the genomes. In one such example, Tam_RS12675 shared an indel with the non-coding region preceding Nan_RS30200 in which Tam_RS12675 had ten guanine (complement) base repeats which was nine in case of Nan_RS30200. Comparing with MxDZ2_Kirby, it was found that MxDZ2_Kirby also had nine repeats in the non-coding region, similar to MxDZ2_Nan assembly.

### MxDZ2_Kirby was used as a reference to build a consensus sequence for *M. xanthus* DZ2

MxDZ2_Kirby can be considered as a better assembly amidst these three because of utilizing short 454 reads which have more coverage and fewer associated errors (like Illumina) (31) as compared to Nanopore or PacBio. Therefore, to determine which of the above indels or substitutions must be correct in the *M. xanthus* consensus DZ2 genome, we took help from the genomic sequence of MxDZ2_Kirby and assigned that respective base present in MxDZ2_Kirby to the consensus sequence [**Figure-2**]. Hereby we report a curated consensus sequence of *M. xanthus* DZ2 with 9,346,278 base pairs accounting for 68% GC content (1,458,168 adenine, 1,452,555 thymine, 3,216,865 guanine and 3,218,690 cytosine).

### *M. xanthus* DZ2 encodes for three Mx alpha units in tandem within its genome

Mx alpha is a polyploid prophage region involved in kin discrimination via the presence of a toxin-antitoxin cassette. The toxin is delivered through outer membrane exchange and the organisms lacking the antitoxin gene are killed (7). Two of three *M. xanthus* strains (DZ2 and DZF1) considered in this study have three Mx alpha units whereas *M. xanthus* DK 1622 has a deletion of ∼200kb resulting in the presence of only one of these units. As reported by Dey A et al (7), Mx alpha unit in *M. xanthus* DK 1622 spans from MXAN_1800 to MXAN_1901 whose corresponding updated locus tags are from MXAN_RS08710 to MXAN_RS36675. *M. xanthus* DZF1 encompasses three Mx alpha units arranged within 16 contigs [**Figure-5**]. *M. xanthus* DZ2 has three Mx alpha units encoded within the following locus tags: for MxDZ2_Tam - RS13240 to RS13765, RS13845 to RS14290 and RS14310 to RS14820; for MxDZ2_Nan - RS29640 to RS29115, RS29035 to RS28590 and RS28570 to RS28055 [**Figure-4, 5**].

### Homologs of Mx alpha were detected in our selected dataset of myxobacteria

Utilizing orthology analysis, we detected Mx alpha genes encoded within other myxobacteria ranging from 1 to 136 homologs per copy within our dataset, which increases to a maximum of 191 proteins in case of extrapolated counts. [**Figure-3**]. On average, the class Myxococcia has higher counts of these homologs compared to Polyangia. Surprisingly, a few order Haliangiales members, especially from family Kofleriaceae including *Haliangium ochraceum* depict the presence of a large number of Mx alpha genes in their genomes. Within Myxococcia, the families Anaeromyxobacteraceae and Vulgatibacteraceae have the least Mx alpha genes with Anaeromyxobacteraceae having just 2 or 3 genes and Vulgatibacteraceae having just 1 gene. Similarly, the presence of several families of SitA proteins was found to be more prominent in Myxococcia than in Polyangia [**Figure-3**].

**Figure-3:**
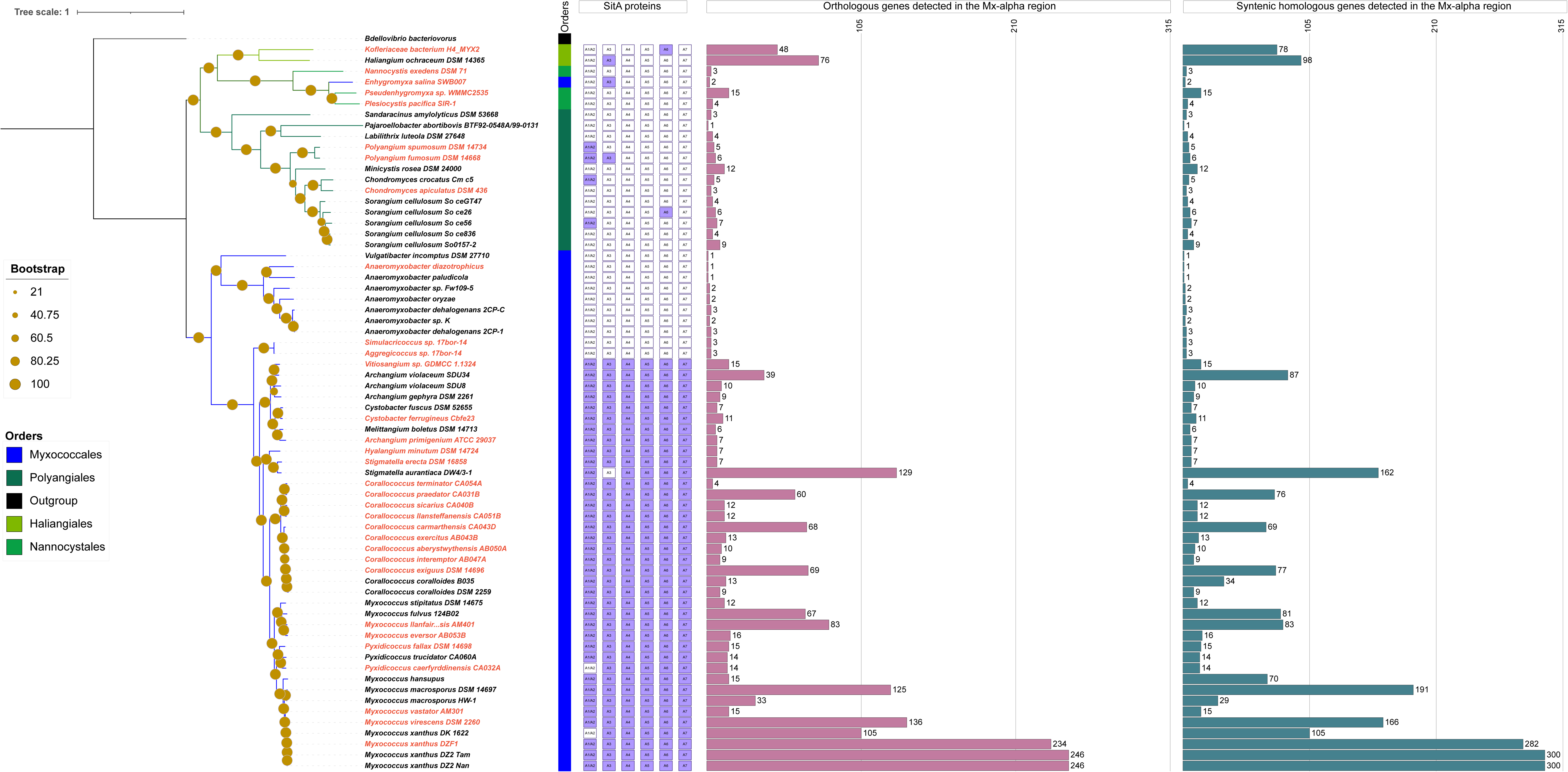
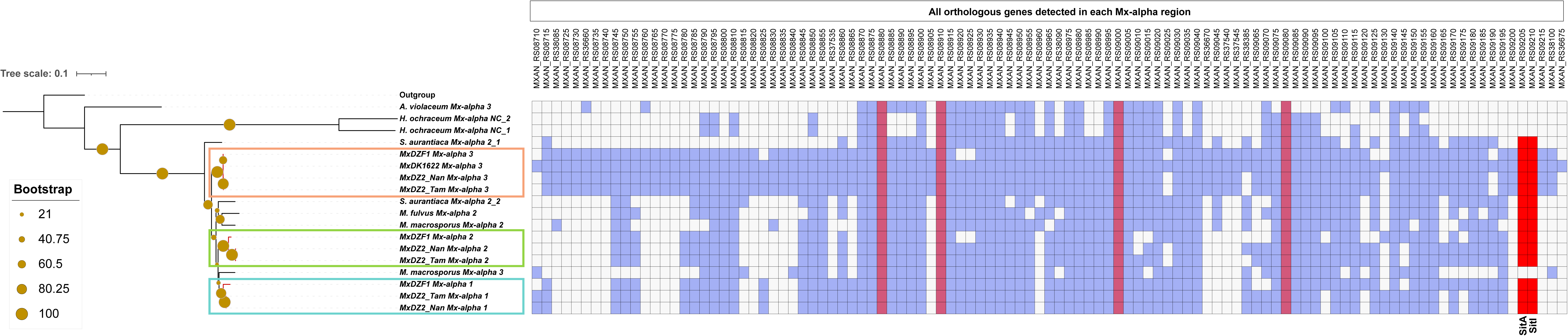
Distribution of SitA proteins and all encoded proteins within the Mx Alpha regions across myxobacterial phylogeny: The concatenated alignments of 235 single copy orthologous genes were used to build a maximum likelihood phylogeny of selected members of the phylum Myxococcota along with *B. bacteriovorus* as an outgroup. The terminal taxa labels in black and red colour represent complete and draft assemblies respectively, whereas the coloured strip indicates the class to which these species belong. The second panel of boxes represent the six families of SitA proteins where a purple colour box indicates the presence of proteins belonging to that family whereas a white box indicates its absence. Two bar plots provide the number of homologs within all detected Mx alpha regions; the first bar plot is the exact number of orthologous genes detected with the above-mentioned parameters in the analysis, and the second one is an extended count of the same where the number of genes was calculated based on the first and the last homolog that was detected. It must be noted that the majority of the genes in these Mx alpha regions are present in synteny but with lesser sequence identity, because of which, there is a difference in the two bar plot numbers. On average, each Mx alpha unit has 80-100 genes.

### Prophage region lies within Mx alpha unit which in turn lies within genomic island(s)

Genomic islands are non-native cluster of genes in the bacterial genome which are acquired through horizontal gene transfer events whereas prophage regions are subset of genomic islands harbouring phage-like genes. Analysis of the complete genomes in our dataset revealed a significant number of Mx alpha homologs [**Figure-5**], identified in five organisms: *Myxococcus macrosporus* DSM 14697, *Myxococcus fulvus* 124B02, *Stigmatella aurantiaca* DW4/3-1, *Archangium violaceum* SDU34, and *Haliangium ochraceum* DSM 14365, along with *M. xanthus* spp. Notably, Mx alpha regions were consistently located within one genomic island or spanned two genomic islands, further encompassed prophage regions [**Figure-5**]. The number of Mx alpha units varied from zero to three across our myxobacterial dataset with only *M. xanthus* DZ2 and *M. xanthus* DZF1 exhibiting the maximum count of three Mx alpha units [**Figure-5**].

### Phylogenetic analysis of conserved Mx alpha genes classified Mx alpha regions

Subsequently, we utilized the phylogenetic analysis considering the core orthologs of Mx alpha genes [**Figure-4**] to classify the Mx alpha regions but most of them could not completely resolve the Mx alpha regions even within *M. xanthus*. Only orthologs of MXAN_RS08880, MXAN_RS08910, MXAN_RS09000 and MXAN_RS09080 were able to resolve Mx alpha units of *M. xanthus* but the single gene, as well as concatenated gene phylogenies, could not resolve the Mx alpha classification for the rest of the organisms [**Figure-4, Supplementary Figure-3**].

**Figure-4:**
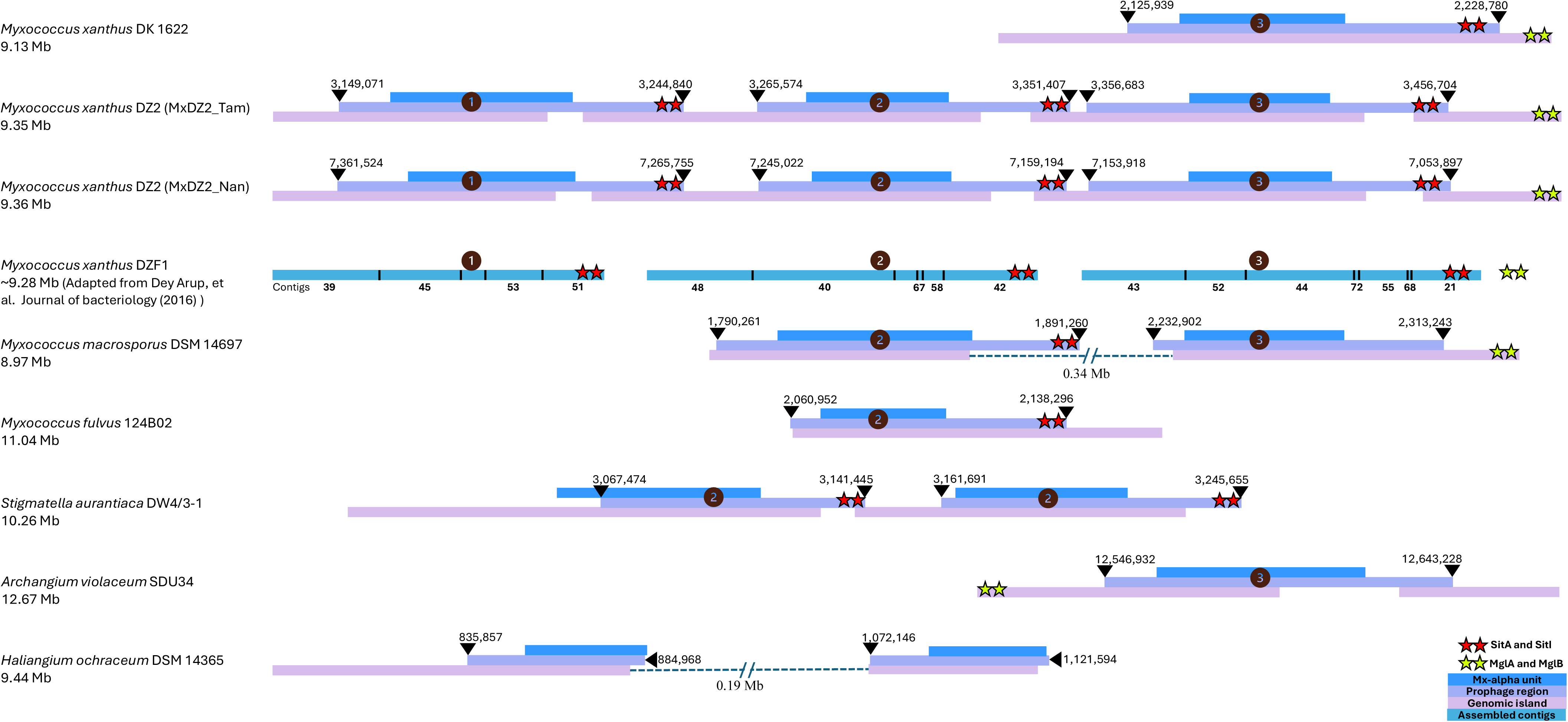
Phylogenetic analysis for Mx alpha type classification: This phylogeny, built using a concatenated alignment of four core Mx alpha genes (depicted in maroon colour), showcases the clustering of respective Mx alpha units in specific clades followed by the depiction of homologs detected in the Mx alpha regions of the respective species. Here, a coloured box indicates the presence of the respective gene above the sequence homology cutoff, and the white box denotes their absence in the organism; however, it must be noted that these absentees have lesser sequence identity with their respective counterparts. SitAI are marked in red coloured boxes. The various Mx alpha units in these studies species have been demarcated within the colour-coded boxes over the phylogeny branches and labels. The Mx alpha unit numbers after the organism name have been abbreviated as follows: (NC: Not characterised), (_1: first to appear based on genomic location), (_2: second to appear based on genomic location).

### SitA1, SitA2 and SitA3 were identified using their distinct domain architecture

As reported for *M. xanthus*, each Mx alpha region contains a toxin gene *sitA*, which is responsible for killing non-kin species/strains not encoding its antitoxin counterpart *sitI*, which is always present downstream of *sitA*. The three SitA proteins associated with the three Mx alpha units in MxDZ2_Tam are: RS38135, RS14285 and RS14805 and in MxDZ2_Nan are: RS29120, RS28595 and RS28070. For all the *M. xanthus* strains considered in this study, the SitA1 protein showed high probability hits with <u>PF14414.9</u> (pfam domain) which is a nuclease belonging to the superfamily of HNH/ENDO VII with conserved WHH. Similarly, SitA2 protein showed hits with <u>3ZBI_I</u> which is a bacterial type IV secretion system and SitA3 showed hits with <u>4G6V_G</u> which is a CdiA-CT/CdiI toxin and immunity complex [**Supplementary Table-2**]. However, N-terminal signal peptide was not detected in SitA1 homologs, i.e., Tam_RS38135 and Nan_RS29120, but lipoprotein signal peptide was detected in the proteins preceding them. In the case of SitA3, Tam_RS14805 did not show the presence of signal peptide but it was present in Nan_RS28070 [**Supplementary Table-2**]. We believe that these two cases have arisen because of gene annotation problems and signal peptide should be a part of SitA proteins.

In MxDZ2_Tam and MxDZ2_Nan, three SitA proteins are associated with each of the three Mx alpha subunits. Consequently, homologs of SitA1, SitA2, and SitA3 were utilized as molecular markers to differentiate between Mx alpha 1, Mx alpha 2, and Mx alpha 3 within the dataset of myxobacteria. For *Stigmatella aurantiaca* DW4/3-1, two SitA proteins associated with the two Mx alpha regions exhibited homology with SitA2 proteins of MxDZ2_Tam and MxDZ2_Nan. These proteins also showed significant alignment with <u>7O3J</u>, a type IV secretion system in bacteria. These observations suggest a possible duplication of Mx alpha 2 unit in this organism [**Figure-5, Supplementary Figure-3**]. However, SitA homologs were not detected within the Mx alpha regions of *Archangium violaceum* SDU34, *Haliangium ochraceum* DSM 14365, and one of the Mx alpha regions in *Myxococcus macrosporus* DSM 14697. As a result, Mx alpha classification based on SitA proteins could not be conducted for these three cases.

**Figure-5:**
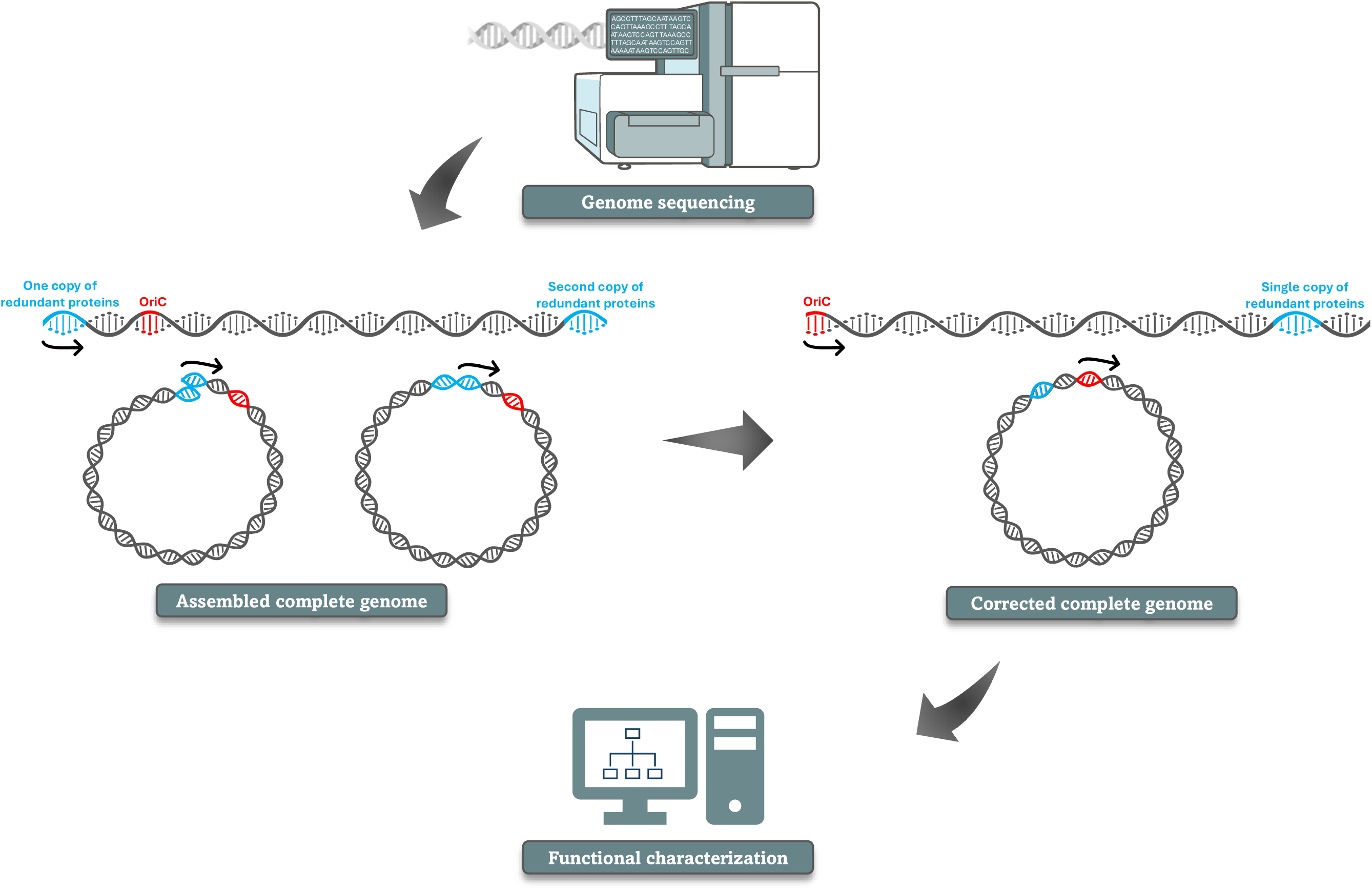
**Diversity, classification and distribution of Mx alpha regions amongst the complete myxobacterial genomes**: This figure depicts the location of detected prophage (Mx alpha) regions, some of their specific genes (SitA and SitI), genomic islands (colour coded as per the right-bottom panel), along with syntenic motility genes, MglA and MglB. SitAI and MglAB are marked in stars of red and neon green colours respectively. The numbers in brown circle indicate the Mx alpha region they belong to as per the proposed classification by Dey A et al (7) and our observations.

SitA also has a downstream immunity protein SitI which renders protection to species from the toxicity of SitA. We looked into the domains associated with SitI proteins and found that SitI1 showed hits with <u>3D5P</u> which belongs to the well-known SUKH-1 superfamily of primary bacterial immunity proteins, SitI2 showed hits with PF19794.2 and PF12713.10 which are mostly domains of unknown function and SitI3 showed hits with PF09816.12 which is known to activate transcription elongation by facilitating the binding of RNA polymerase II [**Supplementary Table-2**]. Based on these directions, we were able to identify all SitA and SitI homologs and used them in the classification of Mx alpha regions.

### MglA and MglB were detected near the end of the genomic island containing Mx alpha 3

In *M. xanthus*, A (Adventurous) and S (Social) motility is known to be regulated by several genes including a *mgl* operon consisting of two genes, *mglA* and *mglB.* Our analysis detected the presence of these two genes in a syntenic manner towards the end of the genomic island containing Mx alpha 3 in *M. xanthus* DK 1622, MxDZ2_Tam and MxDZ2_Nan. Further, we detected the presence of these two genes in contig 21 of *M. xanthus* DZF1, which is the last contig containing Mx alpha 3. Hence the presence of these genes can be considered in the classification of the Mx alpha units within the five organisms. We further detected MglA and MglB at the end of the genomic island of *Archangium violaceum* SDU34 and in the second genomic island of *Myxococcus macrosporus* DSM 14697. Thus, it allowed us to classify the Mx alpha regions in organisms of our dataset except *Haliangium ochraceum* DSM 14365 [**Figure-5**].

## Discussion

Independent sequencing efforts for a singular bacterial strain are uncommon, leading to the assumption that existing genome assembly represents the most accurate version available. However, discrepancies in genomic architecture and sequence among ostensibly identical strains pose significant challenges for experimental and computational analyses. In such instances, elucidating genuine genetic disparities and consolidating them into a consensus assembly is imperative for facilitating research on this model organism.

*Myxococcus xanthus,* the model organism of the group of bacteria known as myxobacteria, has a genome size of around 9Mb encoding for approximately seven thousand genes. Multiple strains of *M. xanthus* with varying assembly quality have been reported in the NCBI database (https://www.ncbi.nlm.nih.gov/genome/browse#!/prokaryotes/) in the last two decades. This study focuses on three assemblies of the *M. xanthus* DZ2 strain submitted by independent research groups. The first, reported in 2013, is a contig-level assembly generated by John Kirby’s group using 454 GS-FLX Titanium technology. In contrast, two subsequent assemblies, submitted in 2022 and 2023, are complete genome assemblies produced by Tam Mignot’s group using PacBio long high-fidelity (HiFi) reads and Beiyan Nan’s lab employing a combination of DNBSEQ and PacBio HiFi sequencing technologies, respectively.

We further systematically examined the genomic variations between these assemblies to provide a comprehensive understanding of the factors contributing to such disparities and to establish a foundation for a consensus assembly. The present study identified 11 and 16 repetitive proteins at both termini of the MxDZ2_Tam and MxDZ2_Nan assemblies, respectively, with genomic segments that are reverse complements of their counterparts in the alternative assembly. Following circularization and rearrangement of their respective assembly sections, a total of 40 variations of frameshift, missense, and nonsense nature, were detected, with 62.5% (25 mutations) localized within the coding regions. Although both assemblies have been sequenced using PacBio HiFi technology which claims to be more than 99.5% accurate for long-read sequencing (32), there is still a probability that these variations are sequencing errors in one of the assemblies. To distinguish between a real mutation and a sequencing error we request the researchers to take the reported leads and verify them by considering the above-mentioned proteins with 50 base pairs upstream and downstream and subjecting them to PCR-based amplification followed by Sanger sequencing which could throw some light into this matter. Conclusively, the first part of this study generated the circular chromosome versions of MxDZ2_Tam and MxDZ2_Nan assemblies using both end trimming, *oriC* site identification, and circularization methods and further used MxDZ2_Kirby as a reliable reference for resolving indels and substitutions. Consequently, we present a curated consensus sequence of *M. xanthus* DZ2 comprising 9,346,278 base pairs with 68% GC content. To verify if these variations are real mutations or just sequencing errors, validation by PCR-based amplification followed by Sanger sequencing must be performed by experimental biologists.

A closely related strain to *Myxococcus xanthus* DZ2, DK 1622, exhibits differences in the number of Mx alpha regions, with the latter harbouring a deletion of two out of three Mx alpha regions. Each Mx alpha region encodes a specific toxin-antitoxin cassette that mediates antagonistic interactions by delivering toxin proteins (SitA) via outer membrane exchange. The SitA toxins are lethal in the absence of their cognate immunity protein, SitI, in recipient cells, thereby conferring a competitive advantage to the donor strain through effective discrimination. This study investigated the homologs of Mx alpha regions across sixty-one diverse myxobacterial species out of which *Myxococcus macrosporus* DSM 14697, *Myxococcus fulvus* 124B02, *Stigmatella aurantiaca* DW4/3-1, *Archangium violaceum* SDU34, and *Haliangium ochraceum* DSM 14365 were identified having these regions along with *M. xanthus* spp. However, none of these species exhibited three Mx alpha regions akin to *M. xanthus* DZ2 and DZF1. We classified the Mx alpha regions across these species using phylogeny of conserved ortholog proteins, SitA protein homologs and subsequent domain analyses. This approach enabled the classification of *M. macrosporus* Mx alpha 2, *M. fulvus* Mx alpha 2, and a duplication event in *S. aurantiaca* resulting in Mx alpha 2_1 and Mx alpha 2_2 [**Figure-5**].

This classification was also supported by the syntenic presence of a few proteins such as two motility genes, *mglA* and *mglB*, were identified at the terminus of genomic islands encoding Mx alpha 3 in *M. xanthus* DK 1622, DZ2, and DZF1. Comparative genomic analyses revealed the presence of these motility genes in the genomic islands of *M. macrosporus* Mx alpha 3 and *A. violaceum* Mx alpha 3, too. However, the Mx alpha regions NC_1 and NC_2 in *H. ochraceum* could not be classified based on homology with SitA or the presence of Mgl proteins. To address this classification ambiguity, further experimental studies are recommended, specifically, competitive assays involving *H. ochraceum* and species harbouring diverse combinations of Mx alpha regions could provide insights into the functional and evolutionary significance of its unclassified Mx alpha units.

In conclusion, the authors would like to argue that the computational data must be cross-checked for such redundancy before submitting online. Notably, these two complete assemblies were not quality-checked before submitting to the NCBI and publishing as an article as they have redundant sequences in the end and do not qualify as a circular genome. Both genomes have been submitted as a circular assembly as depicted in their Nucleotide pages (<u>MxDZ2_Tam</u> and <u>MxDZ2_Nan</u>) and publications (9, 10). These assemblies differ from each other by 6.4 kb despite originating from the same strain obtained from a shared culture collection. These discrepancies challenge the assumption of equivalence and underscore the need to investigate the root cause of these differences. The final circular versions of MxDZ2_Tam and MxDZ2_Nan assemblies have 13,097 (0.13%) and 19,530 (0.21%) base pairs less as compared to their submitted version. This further highlights the requirement of tools which can easily detect such issues in genome assemblies and further recommends the experimental and computational biologists to be more aware of quality checks before submitting and publishing data in open access.

## Supplementary Data Legends

**Supplementary Figure-1:** Synteny comparison of three *M. xanthus* DZ2 assemblies (two complete and one draft) along with *M. xanthus* DK 1622 complete genome using MAUVE tool.

**Supplementary Figure-2:** Dot plots exhibiting the alignments of raw, circularized and rearranged alignments of MxDZ2_Nan, MxDZ2_Tam, consensus sequence of MxDZ2, and *M. xanthus* DK 1622.

**Supplementary Figure-3:** Several single gene phylogenies were built to classify the Mx alpha region of the various organisms.

**Supplementary Table-1:** Nucleotide location and variation information for all 40 detected variations within MxDZ2_Nan and MxDZ2_Tam assemblies. It also shows the comparison with MxDZ2_Kirby as in all the cases, one of them shows similarity with MxDZ2_Kirby assembly. To make a consensus, we have used this information in consideration.

**Supplementary Table-2**: Validation of SitA (SitA1, SitA2 and SitA3) and SitI (SitI1, SitI2 and SitI3) proteins using the best domain identified by HHpred and signal peptide encoded.

**Supplementary File-1:** MAFFT-based multiple sequence alignments of MxDZ2_Tam, MxDZ2_Nan, and MxDZ2_Kirby genomes have been reported here for all forty mutational hotspot regions in both nucleotide and protein sequences.

## Ethics approval and consent to participate

Not applicable

## Availability of data and materials

Authors have used open-source tools in this analysis. All tool versions have been provided in the methodology. The final consensus *M. xanthus* DZ2 genome sequence and its annotations have been deposited to the open-access portal, NCBI, which will be available soon.

## Competing interests

The authors declare no conflict of interest to disclose.

## Funding

UM is supported by DST-INSPIRE fellowship by Department of Science and Technology (Government of India). GS acknowledges the Department of Science and Technology (DST)-INSPIRE Faculty Award by the Government of India and seed grant from the IIT Hyderabad for supporting his research.

## Authors’ contributions

GS generated the idea and supervised the project. UM performed the computational analysis and wrote the first draft of the manuscript. UM and GS edited and finalized the manuscript.

## Notes

### Competing Interest Statement

The authors have declared no competing interest.

