## Supplementary file 1 for "Same strain, two genomes: Creating a consensus circular genome of *Myxococcus xanthus* DZ2 revealed the diversity of a large polyploid prophage region within the close-related myxobacterial strains"

**Frameshift mutations:**

In seven cases, one indel per ORF led to a frameshift mutation resulting in different amino acids across the protein length, however, the reported proteins are of equal lengths with protein in one of the assemblies having multiple stop codons, which is wrong translation. Moreover, in one case, we found seven indels in one single ORF, causing several frameshift mutations across the protein length, but the gene calling, and annotation have predicted almost the same length for both assemblies with one having multiple stop codons in between. These results have been explained below and further in **Supplementary File 2**:

- **Frameshift mutation 1:** Nan_RS36290 and Tam_RS06590 are 926 and 927 base pairs with eight and seven guanine base repeats, respectively. The protein length (308 amino acids) is similar but multiple stop codons are present in MxDZ2_Nan protein due to change in its reading frame. MxDZ2_Kirby and MxDZ2_Tam genes are identical.
- **Frameshift mutations 2:** We observed seven indels between Nan_RS30255 and Tam_RS12620, of which 3 guanine (complement) bases were inserted in MxDZ2_Nan gene (also present in its homolog, Kirby_RS0203280) and the remaining 4 bases (adenine, thymine, guanine, cytosine; all complement) were inserted in MxDZ2_Tam gene (absent in MxDZ2_Kirby gene), resulting in MxDZ2_Nan gene having 2079 bases encoding for 692 amino acids and MxDZ2_Tam gene having 2080 bases encoding for 693 amino acids. Contrary to the above frameshift mutation cases, in this case, MxDZ2_Kirby and MxDZ2_Nan nucleotide and protein sequences are completely identical to each other.
- **Frameshift mutation 3:** Nan_RS26665 and Tam_RS16225 are 1340 and 1341 base pairs respectively having six guanine (complement) base repeats in the latter whereas only five of them in the former. Although both encode 446 amino acids long proteins, there are multiple stop codons observed in MxDZ2_Nan protein due to shift in the reading frame. Its homologous MxDZ2_Kirby gene is similar to MxDZ2_Tam assembly.
- **Frameshift mutation 4:** Both genes, i.e., Nan_RS21765 and Tam_RS21100 are 1790 and 1791 base pairs in length respectively having six and five cytosine base repeats, respectively. Both encode 596 amino acids long proteins; however, multiple stop codons can be seen in the MxDZ2_Nan encoded protein due to shift in the reading frame. Its homologous MxDZ2_Kirby gene also has six guanine base repeats, again showcasing similarity with MxDZ2_Tam assembly.
- **Frameshift mutation 5:** Nan_RS19765 and Tam_RS23100 are 362 and 363 base pairs long respectively with five and four cytosine base repeats. Alignment with MxDZ2_Kirby revealed that Kirby_RS0218355 also has five repeats as seen in MxDZ2_Tam gene. Because of this frameshift mutation, several stop codons were observed in MxDZ2_Nan, though both Nan_RS19765 and Tam_RS23100 reported to encode proteins of length 120 amino acids.
- **Frameshift mutation 6:** Nan_RS16675 and Tam_RS26190 are 1379 and 1380 base pairs respectively with seven and six guanine (complement) repeats reported, respectively. Both encode a 459 amino acids long protein with multiple stop codons observed in like MxDZ2_Nan gene due to shift in the reading frame. Comparing it with Kirby_RS0232470 showcased seven cytosine repeats, as reported in MxDZ2_Tam counterpart.
- **Frameshift mutation 7:** Nan_RS16615 and Tam_RS26250 are 779 and 780 base pairs respectively with the latter having four guanine (complement) base repeats compared to three in the former. Even though both are described to encode proteins of length 259 but there are several stop codons observed in MxDZ2_Nan gene due to frameshift mutation. Alignment with Kirby_RS0232530 revealed that MxDZ2_Kirby gene also have four cytosine bases like gene in the MxDZ2_Tam assembly.
- **Frameshift mutation 8:** Nan_RS16250 and Tam_RS26610 are reported to be of the length 4265 and 4266 base pairs respectively with MxDZ2_Tam gene having six guanine (complement) base repeats whereas only five of them are reported in the respective MxDZ2_Nan gene. Even though both are described to encode proteins of length 1421, several stop codons were observed in MxDZ2_Nan gene due to frameshift mutation. When these two proteins were aligned along with Kirby_RS0227330, we found six cytosine bases in the latter, like MxDZ2_Tam assembly.

**Premature stop codon followed by second ORF prediction:**

Besides these frameshift mutations, seven cases were identified in which a mutation has led to frameshift mutation in either of the complete assemblies resulting in the reading frame reaching a stop codon early and splitting up of a single protein into two proteins, thus predicting short proteins in one of the assemblies as mentioned below and in **Supplementary File 1**:

- **Premature stop codon followed by second ORF prediction 1:** Nan_RS29755 and Tam_RS13120 share a point mutation in which Tam_RS13120 has four cytosine (complement) base repeats which is three in Nan_RS29755 leading to a frameshift mutation and resulting in early reading of stop codon in Nan_RS29755. Tam_RS13120 is a protein length 219 which has shared homology with two proteins in MxDZ2_Nan assembly: Nan_RS29755 of length 151 and Nan_RS29760 of length 68. MSA with MxDZ2_Kirby revealed that Kirby_RS0202735 also has four guanine repeats, similar to MxDZ2_Tam assembly.
- **Premature stop codon followed by second ORF prediction 2:** Nan_RS29095 and Tam_RS13780 share a point mutation in which Tam_RS13780 has five guanine (complement) base repeats which is four in Nan_RS29095 leading to a frameshift mutation and resulting in early reading of stop codon in Nan_RS29095. Tam_RS13780 encodes for a protein of length 472 which has shared homology with two proteins in MxDZ2_Nan assembly: Nan_RS29095 of length 245 and Nan_RS29100 of length 248. MSA with MxDZ2_Kirby revealed that Kirby_RS0235795 also has five cytosine repeats, similar to MxDZ2_Tam assembly.
- **Premature stop codon followed by second ORF prediction 3:** Nan_RS20945 and Tam_RS21920 share a point mutation in which Tam_RS21920 has six guanine base repeats which is five in Nan_RS20945 leading to a frameshift mutation and resulting in next codon to be a stop codon (TGA) in Nan_RS20945. Tam_RS21920 encodes for a protein of length 579 which has shared homology with two proteins in MxDZ2_Nan assembly: Nan_RS20940 of length 67 and Nan_RS20945 of length 512. MSA with MxDZ2_Kirby revealed that Kirby_RS0229920 also has six cytosine (complement) repeats, similar to MxDZ2_Tam assembly.
- **Premature stop codon followed by second ORF prediction 4:** Tam_RS26590 and Nan_RS16270 share a point mutation with four and three guanine (complement) base repeats, respectively, further leading to a frameshift mutation and an early reading of stop codon for MxDZ2_Nan gene. Tam_RS26590 encodes for a protein of length 1435 which share homology with two proteins in MxDZ2_Nan assembly: Nan_RS16270 (630 amino acids) and Nan_RS16275 (806 amino acids). MxDZ2_Kirby gene (Kirby_RS0227310) also has four cytosine base repeats, similar to MxDZ2_Tam assembly.
- **Premature stop codon followed by second ORF prediction 5:** Nan_RS14965 and Tam_RS27885 share a point mutation in which MxDZ2_Tam has four cytosine (complement) base repeats but three in MxDZ2_Nan leading to a frameshift mutation and further resulting in MxDZ2_Nan gene with an early reading of stop codon. MxDZ2_Tam gene encodes 775 amino acid protein which share homology with two proteins in MxDZ2_Nan assembly: Nan_RS14965 of length 288 amino acids and Nan_RS14970 of length 494 amino acids. MSA with MxDZ2_Kirby revealed that Kirby_RS0206915 also has four guanine repeats, like MxDZ2_Tam assembly.
- **Premature stop codon followed by second ORF prediction 6:** Nan_RS08040 and Tam_RS34805 share a point mutation in which the latter has five cytosine base repeats whereas MxDZ2_Nan gene has four which creates a frameshift mutation resulting in an early reading of stop codon in Nan_RS08040. Nan_RS08040 encodes a protein of length 345; however, it shares partial-length homology with two proteins in MxDZ2_Tam assembly: Tam_RS34805 which is 296 amino acids protein and Tam_RS34810 which is 46 amino acids protein. MSA with MxDZ2_Kirby homolog revealed that Kirby_RS0231740 has five guanine (complement) repeats, as in MxDZ2_Tam assembly.
- **Premature stop codon followed by second ORF prediction 7:** Nan_RS02815 and Tam_RS02175 share an indel in which cytosine base in MxDZ2_Tam has been replaced by adenine base in MxDZ2_Nan leading to nonsense mutation and splitting up of MxDZ2_Tam gene (994 amino acids) into two proteins in MxDZ2_Nan assembly: Nan_RS02810 (length 324 amino acids) and Nan_RS02815 (length 662 amino acids). MSA with their Kirby_RS0212590 homolog revealed the presence of a guanine (complement) base like MxDZ2_Tam assembly.

In the above-discussed 15 cases of mutation, it was vividly observed that 14 of them have occurred where a particular base especially cytosine or guanine was present multiple times at that position of the genome. Also, alignment with homologs in MxDZ2_Kirby displayed that in 14 out of the 15 cases, MxDZ2_Kirby was exactly like MxDZ2_Tam.

Apart from the above observations, there are three instances in which a mutation has led to frameshift mutations causing the reading frame to reach a stop codon early resulting in a shorter protein and the remaining nucleotide becoming a part of the non-coding region because of the lack of a start codon in the rest of the part of the protein. Consequently, there is a huge difference in the length of the proteins encoded by the two assemblies:

- **Premature stop codon followed by non-coding region 1:** Nan_RS32920 and Tam_RS09970 share a point mutation in which Tam_RS09970 has a thymine base which is absent in Nan_RS32920 leading to a frameshift mutation and early stop codon reading in Tam_RS09970 making it shorter by 314 amino acids. Nan_RS32920 is of 2255 bases encoding 751 amino acids and Tam_RS09970 is 1314 bases in length translating into 437 amino acids where the remaining 942 bases become a part of the non-coding region. Kirby_RS0217305 does not have adenine (complement), similar to MxDZ2_Nan assembly.
- **Premature stop codon followed by non-coding region 2:** Nan_RS28750 and Tam_RS38160 share a point mutation in which Tam_RS38160 has an additional adenine (complement) base which is missing in Nan_RS28750 leading to a frameshift mutation and resulting in early reading of stop codon in Tam_RS38160 which makes it shorter by 61 amino acids. Nan_RS28750 is of 543 nucleotides encoding 180 amino acids and Tam_RS38160 is 360 nucleotides long translating into 119 amino acids where the remaining 183 bases become a part of the non-coding region. Although Kirby_RS39310 does not have thymine similar to MxDZ2_Nan assembly but the gene encoded by Kirby_RS39310 is 51 nucleotides and 16 amino acids shorter than Nan_RS28750. Unlike the rest of the cases, here MxDZ2_Kirby is not completely similar to any of the two assemblies.
- **Premature stop codon followed by non-coding region 3:** Nan_RS20225 and Tam_RS22640 have a point mutation in which a cytosine (complement) base in Nan_RS20225 is replaced by adenine (complement) in Tam_RS22640. The succeeding bases to this mutation are adenine and guanine which has resulted in nonsense mutation (TAG) in Tam_RS22640 causing the translation to stop, making the protein shorted by 14 amino acids. Nan_RS20225 is of 654 bases encoding 217 amino acids and Tam_RS22640 is 612 bases in length translating into 203 amino acids where the remaining 42 bases become a part of the non-coding region. Kirby_RS0218815 has a guanine base similar to Nan assembly.

Unlike the previous cases in which ~93% (14 out of 15) of them MxDZ2_Kirby was like MxDZ2_Tam, here in the above two out of three cases MxDZ2_Kirby was similar to MxDZ2_Nan. Moreover, we also encountered a special case in which MxDZ2_Kirby was similar to neither MxDZ2_Tam nor MxDZ2_Nan.

Additionally, there were 16 variations detected in the non-coding region of the genome(s), of which 6 were in MxDZ2_Nan non-coding region, 4 in MxDZ2_Tam non-coding region and 6 in non-coding region of both the genomes which are further illustrated below:

- **Non-coding region mutation in MxDZ2_Nan 1:** Tam_RS12675 shared an indel with the non-coding region prior to Nan_RS30200 in which Tam_RS12675 had ten guanine (complement) base repeats which was nine in case of Nan_RS30200. Comparing with MxDZ2_Kirby, it was found that MxDZ2_Kirby also had nine repeats in the non-coding region which is similar to MxDZ2_Nan assembly.
- **Non-coding region mutation in MxDZ2_Nan 2:** An indel is shared between Tam_RS17205 and the non-coding region prior to Nan_RS25680 in which the former has five cytosine base repeats and only four were found in the latter. Comparing with MxDZ2_Kirby, Kirby_RS38320 also had five cytosine bases in the coding region similar to MxDZ2_Tam.
- **Non-coding region mutation in MxDZ2_Nan 3:** An indel was observed between Tam_RS22030 and the non-coding region preceding Nan_RS20830 in which the former had seven guanine base repeats while the latter had only six of them. Comparing with MxDZ2_Kirby, it was found that Kirby_RS0230030 also had seven guanine bases in the coding region similar to MxDZ2_Tam.
- **Non-coding region mutation in MxDZ2_Nan 4:** Another indel was observed between Tam_RS25820 and the non-coding region preceding Nan_RS17045 in which the former has five cytosine (complement) base repeats and the latter reported only four of them. Comparing with MxDZ2_Kirby, it was found that Kirby_RS0232095 also had five of those repeats in the coding region similar to MxDZ2_Tam.
- **Non-coding region mutation in MxDZ2_Nan 5:** Tam_RS32670 and the non-coding region succeeding Nan_RS10175 share an indel in which the former has eight guanine (complement) base repeats whereas the latter reported only seven of them. Upon comparison with MxDZ2_Kirby, it was found that Kirby_RS0206555 also had eight of those repeats in the coding region similar to MxDZ2_Tam.
- **Non-coding region mutation in MxDZ2_Nan 6:** Tam_RS01140 and the non-coding region preceding Nan_RS03855 share an indel in which the former has seven cytosine (complement) base repeats whereas the latter reported only six of them. Upon comparison with MxDZ2_Kirby, it was found that Kirby_RS0209420 also had seven of those repeats in the coding region similar to MxDZ2_Tam.
- **Non-coding region mutation in MxDZ2_Tam 1:** The non-coding region succeeding Tam_RS38160 and coding region of Nan_RS28750 shared four indels in which the former has two additional adenine and two additional thymine bases. Upon comparing with MxDZ2_Kirby homologs, it was found that did not have one of the additional adenine and thymine but the remaining two bases could not be detected because the sequence hit the contig boundary, making MxDZ2_Kirby partially similar to MxDZ2_Nan.
- **Non-coding region mutation 1-5:** There were six variations detected in the five non-coding regions of both the assemblies with Tam’s non-coding regions having an additional cytosine or guanine base. When compared to its counterpart in MxDZ2_Kirby, it was found that MxDZ2_Kirby’s non-coding regions were similar to MxDZ2_Tam.

**CLUSTAL format alignment of variations by MAFFT L-INS-i (v7.526)**

**Frameshift mutation 1**

| **Tam_location** | **Tam_nucleotide** | **Kirby_nucleotide** | **Nan_nucleotide** | **Nan_location** |
| --- | --- | --- | --- | --- |
| **508598** | **G** | **G** | **.** | **508596** |

```
multiple sequence alignment (nucleotide)

MxDZ2_Kirby_RS0221645 atggacctgctcagggttgagcccggcggcttcgctcgctacgaaggggcgatgccgtcc
MxDZ2_Tam_RS06590     atggacctgctcagggttgagcccggcggcttcgctcgctacgaaggggcgatgccgtcc
MxDZ2_Nan_RS36290     atggacctgctcagggttgagcccggcggcttcgctcgctacgaaggggcgatgccgtcc
                      ************************************************************

MxDZ2_Kirby_RS0221645 gagccacaaccgccgtcctgggggcctgattcgtcgtcgcgcctggacttcgaaaaaggg
MxDZ2_Tam_RS06590     gagccacaaccgccgtcctgggggcctgattcgtcgtcgcgcctggacttcgaaaaaggg
MxDZ2_Nan_RS36290     gagccacaaccgccgtcctgggggcctgattcgtcgtcgcgcctggacttcgaaaaaggg
                      ************************************************************

MxDZ2_Kirby_RS0221645 cggctgccgatatgggcggggcgccttcaggtccaccctcggggcgggcgcgccaccgcc
MxDZ2_Tam_RS06590     cggctgccgatatgggcggggcgccttcaggtccaccctcggggcgggcgcgccaccgcc
MxDZ2_Nan_RS36290     cggctgccgatatgggcggggcgccttcaggtccaccctcggggcgggcgcgccaccgcc
                      ************************************************************

MxDZ2_Kirby_RS0221645 gtgcacgcctacgccgtggtgatgctggtgacgcgaggcgagtcgaagatgcggcacgcg
MxDZ2_Tam_RS06590     gtgcacgcctacgccgtggtgatgctggtgacgcgaggcgagtcgaagatgcggcacgcg
MxDZ2_Nan_RS36290     gtgcacgcctacgccgtggtgatgctggtgacgcgaggcgagtcgaagatgcggcacgcg
                      ************************************************************

MxDZ2_Kirby_RS0221645 ggggacctggtggtccgggcgggagacgtccacctgattccgcccggagatgcacacggc
MxDZ2_Tam_RS06590     ggggacctggtggtccgggcgggagacgtccacctgattccgcccggagatgcacacggc
MxDZ2_Nan_RS36290     ggggacctggtggtccgggcgggagacgtccacctgattccgcccggagatgcacacggc
                      ************************************************************

MxDZ2_Kirby_RS0221645 gccggcgcttcgaacgcggaagggtggggcgtggccttccatccggatgccttgggcgag
MxDZ2_Tam_RS06590     gccggcgcttcgaacgcggaagggtggggcgtggccttccatccggatgccttgggcgag
MxDZ2_Nan_RS36290     gccggcgcttcgaacgcggaagggtggggcgtggccttccatccggatgccttgggcgag
                      ************************************************************


MxDZ2_Kirby_RS0221645 gacggtggggccgcgaaccgcctggggccgttgctgcgggtgcgcaagggctgtcacccc
MxDZ2_Tam_RS06590     gacggtggggccgcgaaccgcctggggccgttgctgcgggtgcgcaagggctgtcacccc
MxDZ2_Nan_RS36290     gacggtggggccgcgaaccgcctggggccgttgctgcgggtgcgcaagggctgtcacccc
                      ************************************************************

MxDZ2_Kirby_RS0221645 gtgctgcggcccacgctgccgcagcgccggaggctggcgcggtggatgcgcctgctgtcg
MxDZ2_Tam_RS06590     gtgctgcggcccacgctgccgcagcgccggaggctggcgcggtggatgcgcctgctgtcg
MxDZ2_Nan_RS36290     gtgctgcggcccacgctgccgcagcgccggaggctggcgcggtggatgcgcctgctgtcg
                      ************************************************************

MxDZ2_Kirby_RS0221645 gaggaggtggcccggaacgagccGgggggggaggaagccgcgcgctcgctgctgcggctg
MxDZ2_Tam_RS06590     gaggaggtggcccggaacgagccGgggggggaggaagccgcgcgctcgctgctgcggctg
MxDZ2_Nan_RS36290     gaggaggtggcccggaacgagcc-gggggggaggaagccgcgcgctcgctgctgcggctg
                      *********************** ************************************

MxDZ2_Kirby_RS0221645 gtcctcatcgaactggagcgcatgacggcccaaacggggacgagcgagccgccgtccctg
MxDZ2_Tam_RS06590     gtcctcatcgaactggagcgcatgacggcccaaacggggacgagcgagccgccgtccctg
MxDZ2_Nan_RS36290     gtcctcatcgaactggagcgcatgacggcccaaacggggacgagcgagccgccgtccctg
                      ************************************************************

MxDZ2_Kirby_RS0221645 gggctgagccgcaaggcgctcacgtacatcgagacgcactgtctggagccgctgtcgctc
MxDZ2_Tam_RS06590     gggctgagccgcaaggcgctcacgtacatcgagacgcactgtctggagccgctgtcgctc
MxDZ2_Nan_RS36290     gggctgagccgcaaggcgctcacgtacatcgagacgcactgtctggagccgctgtcgctc
                      ************************************************************

MxDZ2_Kirby_RS0221645 gcgcaggtggcgaaggcgctggggcgctcctccgcgcacgtcgcgggcgtggtgcggcag
MxDZ2_Tam_RS06590     gcgcaggtggcgaaggcgctggggcgctcctccgcgcacgtcgcgggcgtggtgcggcag
MxDZ2_Nan_RS36290     gcgcaggtggcgaaggcgctggggcgctcctccgcgcacgtcgcgggcgtggtgcggcag
                      ************************************************************

MxDZ2_Kirby_RS0221645 gagacgggccgcacggtgggcgagtggattctggagtgccggatggcggaggcacgccgc
MxDZ2_Tam_RS06590     gagacgggccgcacggtgggcgagtggattctggagtgccggatggcggaggcacgccgc
MxDZ2_Nan_RS36290     gagacgggccgcacggtgggcgagtggattctggagtgccggatggcggaggcacgccgc
                      ************************************************************

MxDZ2_Kirby_RS0221645 cggctgcgtggcacggacgagcgcgtggacatcatcgcggagcgcgtgggctacgcggac
MxDZ2_Tam_RS06590     cggctgcgtggcacggacgagcgcgtggacatcatcgcggagcgcgtgggctacgcggac
MxDZ2_Nan_RS36290     cggctgcgtggcacggacgagcgcgtggacatcatcgcggagcgcgtgggctacgcggac
                      ************************************************************


MxDZ2_Kirby_RS0221645 gtgacgcacttcatccgtcagttccgccgcgtccacggggtgactccggccgcgtggcgg
MxDZ2_Tam_RS06590     gtgacgcacttcatccgtcagttccgccgcgtccacggggtgactccggccgcgtggcgg
MxDZ2_Nan_RS36290     gtgacgcacttcatccgtcagttccgccgcgtccacggggtgactccggccgcgtggcgg
                      ************************************************************

MxDZ2_Kirby_RS0221645 cgcaaggcgacggctggggctccatga
MxDZ2_Tam_RS06590     cgcaaggcgacggctggggctccatga
MxDZ2_Nan_RS36290     cgcaaggcgacggctggggctccatga
                      ***************************
-----------------------------------------------------------------------------------
multiple sequence alignment (protein)

MxDZ2_Kirby_RS0221645 MDLLRVEPGGFARYEGAMPSEPQPPSWGPDSSSRLDFEKGRLPIWAGRLQVHPRGGRATA
MxDZ2_Tam_RS06590     MDLLRVEPGGFARYEGAMPSEPQPPSWGPDSSSRLDFEKGRLPIWAGRLQVHPRGGRATA
MxDZ2_Nan_RS36290     MDLLRVEPGGFARYEGAMPSEPQPPSWGPDSSSRLDFEKGRLPIWAGRLQVHPRGGRATA
                      ************************************************************

MxDZ2_Kirby_RS0221645 VHAYAVVMLVTRGESKMRHAGDLVVRAGDVHLIPPGDAHGAGASNAEGWGVAFHPDALGE
MxDZ2_Tam_RS06590     VHAYAVVMLVTRGESKMRHAGDLVVRAGDVHLIPPGDAHGAGASNAEGWGVAFHPDALGE
MxDZ2_Nan_RS36290     VHAYAVVMLVTRGESKMRHAGDLVVRAGDVHLIPPGDAHGAGASNAEGWGVAFHPDALGE
                      ************************************************************

MxDZ2_Kirby_RS0221645 DGGAANRLGPLLRVRKGCHPVLRPTLPQRRRLARWMRLLSEEVARNEPGGEEAARSLLRL
MxDZ2_Tam_RS06590     DGGAANRLGPLLRVRKGCHPVLRPTLPQRRRLARWMRLLSEEVARNEPGGEEAARSLLRL
MxDZ2_Nan_RS36290     DGGAANRLGPLLRVRKGCHPVLRPTLPQRRRLARWMRLLSEEVARNEPGGRKPRARC---
                      **************************************************.:.

MxDZ2_Kirby_RS0221645 VLIELERMTAQTGTSEPPSLGLSRKALTYIETHCLEPLSLAQVAKALGRSSAHVAGVVRQ
MxDZ2_Tam_RS06590     VLIELERMTAQTGTSEPPSLGLSRKALTYIETHCLEPLSLAQVAKALGRSSAHVAGVVRQ
MxDZ2_Nan_RS36290     ---------------------------------CGWSSSNWSARPKRGRASRRPWGAARR
                                                       *  . *  ..    **:* :  *..*:

MxDZ2_Kirby_RS0221645 ETGRTVGEWILECRMAEARRRLRGTDERVDIIAERVGYADVTHFIRQFRRVHGVTPAAWR
MxDZ2_Tam_RS06590     ETGRTVGEWILECRMAEARRRLRGTDERVDIIAERVGYADVTHFIRQFRRVHGVTPAAWR
MxDZ2_Nan_RS36290     SRTSRRTVWS-RCRSRRWRRRWGAPPRTSRAWCGRR---------RAARWASGFWSAGWR
                      .       *  .**  . ***  .. .     . *          *  * . *. .*.**

MxDZ2_Kirby_RS0221645 RKATA------------------------------------------GAP
MxDZ2_Tam_RS06590     RKATA------------------------------------------GAP
MxDZ2_Nan_RS36290     RHAAGCVARTSAWTSSRSAWATRTRTSSVSSAASTGLRPRGGARRRLGLH
                      *:*:.                                          *
```

**Frameshift mutations 2**

| **Tam_location** | **Tam_nucleotide** | **Kirby_nucleotide** | **Nan_nucleotide** | **Nan_location** |
| --- | --- | --- | --- | --- |
| **2998327** | **A(complement)** | **.** | **.** | **1975353** |
| **2998344** | **T(complement)** | **.** | **.** | **1975369** |
| **2998473** | **.** | **G(complement)** | **G(complement)** | **1975499** |
| **2998482** | **.** | **G(complement)** | **G(complement)** | **1975509** |
| **2998559** | **.** | **G(complement)** | **G(complement)** | **1975587** |
| **2998596** | **G(complement)** | **.** | **.** | **1975623** |
| **2998860** | **C(complement)** | **.** | **.** | **1975886** |

```
multiple sequence alignment (nucleotide)

MxDZ2_Kirby_RS0203280 atggagttgtcgagtctcgagtcgagtttctgggcaatcgcaccgggcgtggtcggcgcc
MxDZ2_Nan_RS30255     atggagttgtcgagtctcgagtcgagtttctgggcaatcgcaccgggcgtggtcggcgcc
MxDZ2_Tam_RS12620     atggagttgtcgagtctcgagtcgagtttctgggcaatcgcaccgggcgtggtcggcgcc
                      ************************************************************

MxDZ2_Kirby_RS0203280 gtggggctgctcttcgctgcggtcttctacttccgcgtcaaggcgctcccggagggtgac
MxDZ2_Nan_RS30255     gtggggctgctcttcgctgcggtcttctacttccgcgtcaaggcgctcccggagggtgac
MxDZ2_Tam_RS12620     gtggggctgctcttcgctgcggtcttctacttccgcgtcaaggcgctcccggagggtgac
                      ************************************************************

MxDZ2_Kirby_RS0203280 gcgacgatgaaccgcatcgcgggctacatccgcgacggcgcgatggcgttcctcgtccgc
MxDZ2_Nan_RS30255     gcgacgatgaaccgcatcgcgggctacatccgcgacggcgcgatggcgttcctcgtccgc
MxDZ2_Tam_RS12620     gcgacgatgaaccgcatcgcgggctacatccgcgacggcgcgatggcgttcctcgtccgc
                      ************************************************************

MxDZ2_Kirby_RS0203280 gagtacaaggtgctcgccgcgtactgcgcggtcatcgcggtgctcatcggcctggc-gct
MxDZ2_Nan_RS30255     gagtacaaggtgctcgccgcgtactgcgcggtcatcgcggtgctcatcggcctggc-gct
MxDZ2_Tam_RS12620     gagtacaaggtgctcgccgcgtactgcgcggtcatcgcggtgctcatcggcctggcGgct
                      ******************************************************** ***

MxDZ2_Kirby_RS0203280 ggggccgctcgccagcgggagcttcgtgctcggcgcgttcctctcgctgctggccggcta
MxDZ2_Nan_RS30255     ggggccgctcgccagcgggagcttcgtgctcggcgcgttcctctcgctgctggccggcta
MxDZ2_Tam_RS12620     ggggccgctcgccagcgggagcttcgtgctcggcgcgttcctctcgctgctggccggcta
                      ************************************************************

MxDZ2_Kirby_RS0203280 catcggcatgaaggccgcgacgttcgcgaacgtgcgcaccgcgcaggccgcgcgcaccgg
MxDZ2_Nan_RS30255     catcggcatgaaggccgcgacgttcgcgaacgtgcgcaccgcgcaggccgcgcgcaccgg
MxDZ2_Tam_RS12620     catcggcatgaaggccgcgacgttcgcgaacgtgcgcaccgcgcaggccgcgcgcaccgg
                      ************************************************************

MxDZ2_Kirby_RS0203280 ctccaagccgaacgcgctgctggtcgcgctcgacggtggcgcggtcatgggcctcgccgt
MxDZ2_Nan_RS30255     ctccaagccgaacgcgctgctggtcgcgctcgacggtggcgcggtcatgggcctcgccgt
MxDZ2_Tam_RS12620     ctccaagccgaacgcgctgctggtcgcgctcgacggtggcgcggtcatgggcctcgccgt
                      ************************************************************

MxDZ2_Kirby_RS0203280 cgccggcctgggcctcatcggcatgggcggcgtctactacgccttccaggggcatccgca
MxDZ2_Nan_RS30255     cgccggcctgggcctcatcggcatgggcggcgtctactacgccttccaggggcatccgca
MxDZ2_Tam_RS12620     cgccggcctgggcctcatcggcatgggcggcgtctactacgccttccaggggcatccgca
                      ************************************************************

MxDZ2_Kirby_RS0203280 gctttcccccgtcctccact-ccttcgccgtgggcgccagctccatcgcgctcttcgCcc
MxDZ2_Nan_RS30255     gctttcccccgtcctccact-ccttcgccgtgggcgccagctccatcgcgctcttcgCcc
MxDZ2_Tam_RS12620     gctttcccccgtcctccactCccttcgccgtgggcgccagctccatcgcgctcttcg-cc
                      ******************** ************************************ **

MxDZ2_Kirby_RS0203280 gcgtgggcggcggcatctacaccaaggccgctgacgtcggctccgacatcgccggcaagg
MxDZ2_Nan_RS30255     gcgtgggcggcggcatctacaccaaggccgctgacgtcggctccgacatcgccggcaagg
MxDZ2_Tam_RS12620     gcgtgggcggcggcatctacaccaaggccgctgacgtcggctccgacatcgccggcaagg
                      ************************************************************

MxDZ2_Kirby_RS0203280 tcattgagaacatCcccgaggacgaCccgcgcaacccgggcgtcatcgccgacaacgtgg
MxDZ2_Nan_RS30255     tcattgagaacatCcccgaggacgaCccgcgcaacccgggcgtcatcgccgacaacgtgg
MxDZ2_Tam_RS12620     tcattgagaacat-cccgaggacga-ccgcgcaacccgggcgtcatcgccgacaacgtgg
                      ************* *********** **********************************

MxDZ2_Kirby_RS0203280 gcgacaacgtgggtgacgtggccggcatgggcgccgacatctacgagtccatggtggccg
MxDZ2_Nan_RS30255     gcgacaacgtgggtgacgtggccggcatgggcgccgacatctacgagtccatggtggccg
MxDZ2_Tam_RS12620     gcgacaacgtgggtgacgtggccggcatgggcgccgacatctacgagtccatggtggccg
                      ************************************************************

MxDZ2_Kirby_RS0203280 cgattgtcgccgccatggccatcgcgctgaccgcc-agcgccgcggacctctc-ccgcct
MxDZ2_Nan_RS30255     cgattgtcgccgccatggccatcgcgctgaccgcc-agcgccgcggacctctc-ccgcct
MxDZ2_Tam_RS12620     cgattgtcgccgccatggccatcgcgctgaccgccAagcgccgcggacctctcTccgcct
                      *********************************** ***************** ******

MxDZ2_Kirby_RS0203280 cgtggtggacccctccgcgatcggaagcgcgaaggtggcgggcgtggtgatcccgctggt
MxDZ2_Nan_RS30255     cgtggtggacccctccgcgatcggaagcgcgaaggtggcgggcgtggtgatcccgctggt
MxDZ2_Tam_RS12620     cgtggtggacccctccgcgatcggaagcgcgaaggtggcgggcgtggtgatcccgctggt
                      ************************************************************

MxDZ2_Kirby_RS0203280 gctgtccgccgtgggcctggtggtcagcctgctgagcatcttcatcgcgcgcgcgctcaa
MxDZ2_Nan_RS30255     gctgtccgccgtgggcctggtggtcagcctgctgagcatcttcatcgcgcgcgcgctcaa
MxDZ2_Tam_RS12620     gctgtccgccgtgggcctggtggtcagcctgctgagcatcttcatcgcgcgcgcgctcaa
                      ************************************************************

MxDZ2_Kirby_RS0203280 gcacatgaaccccgcgcaggtgctgcgcagcgccctcatcctgcccccggtcatcctggt
MxDZ2_Nan_RS30255     gcacatgaaccccgcgcaggtgctgcgcagcgccctcatcctgcccccggtcatcctggt
MxDZ2_Tam_RS12620     gcacatgaaccccgcgcaggtgctgcgcagcgccctcatcctgcccccggtcatcctggt
                      ************************************************************

MxDZ2_Kirby_RS0203280 gggtctgtccttcgtgatgatgcaggtgttcggcctgtcccaggccatcacggtggcgct
MxDZ2_Nan_RS30255     gggtctgtccttcgtgatgatgcaggtgttcggcctgtcccaggccatcacggtggcgct
MxDZ2_Tam_RS12620     gggtctgtccttcgtgatgatgcaggtgttcggcctgtcccaggccatcacggtggcgct
                      ************************************************************

MxDZ2_Kirby_RS0203280 ggccgcgggcgccttcggcggcgccatcatcggcctcgtcacggactactacacgtcgtc
MxDZ2_Nan_RS30255     ggccgcgggcgccttcggcggcgccatcatcggcctcgtcacggactactacacgtcgtc
MxDZ2_Tam_RS12620     ggccgcgggcgccttcggcggcgccatcatcggcctcgtcacggactactacacgtcgtc
                      ************************************************************

MxDZ2_Kirby_RS0203280 cacgccggtgcagcgcatcgcggaggcctccgtcacgggtgccggcaccaacctcatccg
MxDZ2_Nan_RS30255     cacgccggtgcagcgcatcgcggaggcctccgtcacgggtgccggcaccaacctcatccg
MxDZ2_Tam_RS12620     cacgccggtgcagcgcatcgcggaggcctccgtcacgggtgccggcaccaacctcatccg
                      ************************************************************

MxDZ2_Kirby_RS0203280 cggcctcgccgtcggcatggagagcgtgggcatcccgatggccaccatcgccgtggtggc
MxDZ2_Nan_RS30255     cggcctcgccgtcggcatggagagcgtgggcatcccgatggccaccatcgccgtggtggc
MxDZ2_Tam_RS12620     cggcctcgccgtcggcatggagagcgtgggcatcccgatggccaccatcgccgtggtggc
                      ************************************************************

MxDZ2_Kirby_RS0203280 ctacatcgccgaccaggcgcttggcctgtacggcatcgcgctggcggccgtgggcatgct
MxDZ2_Nan_RS30255     ctacatcgccgaccaggcgcttggcctgtacggcatcgcgctggcggccgtgggcatgct
MxDZ2_Tam_RS12620     ctacatcgccgaccaggcgcttggcctgtacggcatcgcgctggcggccgtgggcatgct
                      ************************************************************

MxDZ2_Kirby_RS0203280 gggcggcaccgccgtggtgatgaccgtggatgcctacggccccatctccgacaacgccgg
MxDZ2_Nan_RS30255     gggcggcaccgccgtggtgatgaccgtggatgcctacggccccatctccgacaacgccgg
MxDZ2_Tam_RS12620     gggcggcaccgccgtggtgatgaccgtggatgcctacggccccatctccgacaacgccgg
                      ************************************************************


MxDZ2_Kirby_RS0203280 tggcatctccgagatgtccggcctgggccccgaggtccgcgccatcaccgatgagctgga
MxDZ2_Nan_RS30255     tggcatctccgagatgtccggcctgggccccgaggtccgcgccatcaccgatgagctgga
MxDZ2_Tam_RS12620     tggcatctccgagatgtccggcctgggccccgaggtccgcgccatcaccgatgagctgga
                      ************************************************************

MxDZ2_Kirby_RS0203280 cgcggtgggcaacaccacggcggccatcggcaagggcttcgccatcggctcggcgacgct
MxDZ2_Nan_RS30255     cgcggtgggcaacaccacggcggccatcggcaagggcttcgccatcggctcggcgacgct
MxDZ2_Tam_RS12620     cgcggtgggcaacaccacggcggccatcggcaagggcttcgccatcggctcggcgacgct
                      ************************************************************

MxDZ2_Kirby_RS0203280 caccgtcatcgcgctcttctccgccttcaacctcgaggtcaaccacacgcgcatcgccgc
MxDZ2_Nan_RS30255     caccgtcatcgcgctcttctccgccttcaacctcgaggtcaaccacacgcgcatcgccgc
MxDZ2_Tam_RS12620     caccgtcatcgcgctcttctccgccttcaacctcgaggtcaaccacacgcgcatcgccgc
                      ************************************************************

MxDZ2_Kirby_RS0203280 tggcctgccggagatgagcctgcaactgaccaacccgaacgtcatcgtcggcctgctgct
MxDZ2_Nan_RS30255     tggcctgccggagatgagcctgcaactgaccaacccgaacgtcatcgtcggcctgctgct
MxDZ2_Tam_RS12620     tggcctgccggagatgagcctgcaactgaccaacccgaacgtcatcgtcggcctgctgct
                      ************************************************************

MxDZ2_Kirby_RS0203280 gggctccatcctcccgttcctggtgggcgcctccacgatgctcgccgtgggtcgcgcggc
MxDZ2_Nan_RS30255     gggctccatcctcccgttcctggtgggcgcctccacgatgctcgccgtgggtcgcgcggc
MxDZ2_Tam_RS12620     gggctccatcctcccgttcctggtgggcgcctccacgatgctcgccgtgggtcgcgcggc
                      ************************************************************

MxDZ2_Kirby_RS0203280 cggtgccatcgtcgaggagattggccgtcagttccgcgagattccggggctgatggagct
MxDZ2_Nan_RS30255     cggtgccatcgtcgaggagattggccgtcagttccgcgagattccggggctgatggagct
MxDZ2_Tam_RS12620     cggtgccatcgtcgaggagattggccgtcagttccgcgagattccggggctgatggagct
                      ************************************************************

MxDZ2_Kirby_RS0203280 caaggcggacccggaccccaagaagatcgtcgacatcgccaccaagagcgccctgcaaga
MxDZ2_Nan_RS30255     caaggcggacccggaccccaagaagatcgtcgacatcgccaccaagagcgccctgcaaga
MxDZ2_Tam_RS12620     caaggcggacccggaccccaagaagatcgtcgacatcgccaccaagagcgccctgcaaga
                      ************************************************************

MxDZ2_Kirby_RS0203280 gatggtcttcccgggcatcatcgccgtcgccgccccgcccctggtgggctacctgctggg
MxDZ2_Nan_RS30255     gatggtcttcccgggcatcatcgccgtcgccgccccgcccctggtgggctacctgctggg
MxDZ2_Tam_RS12620     gatggtcttcccgggcatcatcgccgtcgccgccccgcccctggtgggctacctgctggg
                      ************************************************************


MxDZ2_Kirby_RS0203280 ccctggtgcgctggcgggtctgctggccggctcgctcgtcgtcggcgccacgatggccct
MxDZ2_Nan_RS30255     ccctggtgcgctggcgggtctgctggccggctcgctcgtcgtcggcgccacgatggccct
MxDZ2_Tam_RS12620     ccctggtgcgctggcgggtctgctggccggctcgctcgtcgtcggcgccacgatggccct
                      ************************************************************

MxDZ2_Kirby_RS0203280 ctacatggccaacgcgggcggcgcgtgggacaacgccaagaagttcatcgagaagggcaa
MxDZ2_Nan_RS30255     ctacatggccaacgcgggcggcgcgtgggacaacgccaagaagttcatcgagaagggcaa
MxDZ2_Tam_RS12620     ctacatggccaacgcgggcggcgcgtgggacaacgccaagaagttcatcgagaagggcaa
                      ************************************************************

MxDZ2_Kirby_RS0203280 gctgccgggccacgcgaagggctccgcggtccacaaggccgccgtcgtcggtgacatggt
MxDZ2_Nan_RS30255     gctgccgggccacgcgaagggctccgcggtccacaaggccgccgtcgtcggtgacatggt
MxDZ2_Tam_RS12620     gctgccgggccacgcgaagggctccgcggtccacaaggccgccgtcgtcggtgacatggt
                      ************************************************************

MxDZ2_Kirby_RS0203280 cggtgacccgttcaaggacacctccggtcctggcgtggccatcctcatcaaggtcatgag
MxDZ2_Nan_RS30255     cggtgacccgttcaaggacacctccggtcctggcgtggccatcctcatcaaggtcatgag
MxDZ2_Tam_RS12620     cggtgacccgttcaaggacacctccggtcctggcgtggccatcctcatcaaggtcatgag
                      ************************************************************

MxDZ2_Kirby_RS0203280 cgtcgtgtccctgctcgtggcctcgctcatcgcgctgcggtag
MxDZ2_Nan_RS30255     cgtcgtgtccctgctcgtggcctcgctcatcgcgctgcggtag
MxDZ2_Tam_RS12620     cgtcgtgtccctgctcgtggcctcgctcatcgcgctgcggtag
                      *******************************************


-----------------------------------------------------------------------------------
multiple sequence alignment (protein)


MxDZ2_Kirby_RS0203280 MELSSLESSFWAIAPGVVGAVGLLFAAVFYFRVKALPEGDATMNRIAGYIRDGAMAFLVR
MxDZ2_Nan_RS30255     MELSSLESSFWAIAPGVVGAVGLLFAAVFYFRVKALPEGDATMNRIAGYIRDGAMAFLVR
MxDZ2_Tam_RS12620     MELSSLESSFWAIAPGVVGAVGLLFAAVFYFRVKALPEGDATMNRIAGYIRDGAMAFLVR
                      ************************************************************

MxDZ2_Kirby_RS0203280 EYKVLAAYCAVIAVLIGLALG------------PLASGSFVLGAFLSLLAGY-IGMKAAT
MxDZ2_Nan_RS30255     EYKVLAAYCAVIAVLIGLALG------------PLASGSFVLGAFLSLLAGY-IGMKAAT
MxDZ2_Tam_RS12620     EYKVLAAYCAVIAVLIGLAAGAARQRELRARRVPLAAGR------LHRHEGRDVRERAHR
                      ******************* *            ***:*       *    *  :  :*


MxDZ2_Kirby_RS0203280 FANVRTAQAARTGSKPNALLVALDGGAVMGLAVAGLGLIGMGGVYYAFQGHPQLSPV---
MxDZ2_Nan_RS30255     FANVRTAQAARTGSKPNALLVALDGGAVMGLAVAGLGLIGMGGVYYAFQGHPQLSPV---
MxDZ2_Tam_RS12620     AGRAHRLQAERAAGRARRWRGHGPRRRRPGPHRHGRRLLRLPGASAAFPRPPLPSPWAPA
                       ...:  ** *:..:..            *    *  *: : *.  **   *  **

MxDZ2_Kirby_RS0203280 -----------LHS------------------FAVGASSIALFA----------------
MxDZ2_Nan_RS30255     -----------LHS------------------FAVGASSIALFA----------------
MxDZ2_Tam_RS12620     PSRSSPRGRRHLHQGRRRLRHRRQGHEHPEDDRATRASSPTTWATTWVTWPAWAPTSTSP
                                 **.                   *. *** : :*

MxDZ2_Kirby_RS0203280 -------------------------------RVGGGIYTKAADVGSDIAGKVIENI----
MxDZ2_Nan_RS30255     -------------------------------RVGGGIYTKAADVGSDIAGKVIENI----
MxDZ2_Tam_RS12620     WWPRLSPPWPSRPPSAADLSPPRGGPLRDRKREGGGRGDPAGAVRRGPGGQPAEHLHRAR
                                                     * ***    *. *  . .*:  *::

MxDZ2_Kirby_RS0203280 -------------PEDDPRNPG--VIADNVGDNVGDVAGMGADIYESMVAAIVAAMAIAL
MxDZ2_Nan_RS30255     -------------PEDDPRNPG--VIADNVGDNVGDVAGMGADIYESMVAAIVAAMAIAL
MxDZ2_Tam_RS12620     AQAHEPRAGAAQRPHPAPGHPGGSVLRDDAG--VRPVPGHHGG-----------------
                                   *.  * :**  *: *:.*  *  *.*  ..

MxDZ2_Kirby_RS0203280 TASAADLSRLVVDPSAIGSAKVAGVVIPLVLSAVGLVVSLLSIFIARALKHMNPAQVLRS
MxDZ2_Nan_RS30255     TASAADLSRLVVDPSAIGSAKVAGVVIPLVLSAVGLVVSLLSIFIARALKHMNPAQVLRS
MxDZ2_Tam_RS12620     -AGRGRLRR----------------------------------------RHHRPRH----
                       *. . * *                                        :* .* :

MxDZ2_Kirby_RS0203280 ALILPPVILVGLSFVMMQVFGLSQAITVALAAGAFGGAIIGLVTDYYTSSTPVQRIAEAS
MxDZ2_Nan_RS30255     ALILPPVILVGLSFVMMQVFGLSQAITVALAAGAFGGAIIGLVTDYYTSSTPVQRIAEAS
MxDZ2_Tam_RS12620     ----------GLLHV------------VHAGAAHRGGLRHG--CRHQPHPRPRRRHGE--
                                ** .*            *  .*.  **   *    : . . * :* .*

MxDZ2_Kirby_RS0203280 VTGAGTNLIRGLAVGMESVGIPMATIAVVAYIADQALGLYGIALAAVGMLGGTAVVMTVD
MxDZ2_Nan_RS30255     VTGAGTNLIRGLAVGMESVGIPMATIAVVAYIADQALGLYGIALAAVGMLGGTAVVMTVD
MxDZ2_Tam_RS12620     ---------RGHPDGHHRRG-----------------GLHRRPGA---------------
                               ** . * .  *                 **:  . *

MxDZ2_Kirby_RS0203280 AYGPISDNAGGI---------SEMSGLGPEVRAITDELDAV-----GNTTAAIGKG----
MxDZ2_Nan_RS30255     AYGPISDNAGGI---------SEMSGLGPEVRAITDELDAV-----GNTTAAIGKG----
MxDZ2_Tam_RS12620     --WPVRHRAGGRGHAGRHRRGDDRGCLRPHLRQRRWHLRDVRPGPRGPRHHRAGRGGQHH
                         *: ..***          .: . * *.:*    .*  *     *      *:*


MxDZ2_Kirby_RS0203280 --------FAIGSATLTVIALF------------------SAFNLEVNHTRIAAGLPEMS
MxDZ2_Nan_RS30255     --------FAIGSATLTVIALF------------------SAFNLEVNHTRIAAGLPEMS
MxDZ2_Tam_RS12620     GGHRQGLRHRLGDAHRHRALLRLQPRGQPHAHRRWPAGDEPATDQPERHRRPAAGL----
                              . :*.*      *                   .* :   .* * ****

MxDZ2_Kirby_RS0203280 LQLTNPNVIVGLLLGSILPFLVGASTMLAVGRAAGAIVEEIGRQFRE-----IP---GLM
MxDZ2_Nan_RS30255     LQLTNPNVIVGLLLGSILPFLVGASTMLAVGRAAGAIVEEIGRQFRE-----IP---GLM
MxDZ2_Tam_RS12620     ----HPPV--------------------PGGRLHDARRGSRGRCHRRGDWPSVPRDSGAD
                          :* *                    . **  .*   . ** .*.     :*   *


MxDZ2_Kirby_RS0203280 ELKADPDPKKIVDIATKSALQEMV----FPGIIAVAAPPLVGYLLGPGALAGLLAGSLVV
MxDZ2_Nan_RS30255     ELKADPDPKKIVDIATKSALQEMV----FPGIIAVAAPPLVGYLLGPGALAGLLAGSLVV
MxDZ2_Tam_RS12620     GAQGGPGPQE----DRRHRHQERPARDGLPGHHRRRRP-------APG---GLPAGPWCA
                        :..*.*::      :   **      :**      *       .**   ** **.  .

MxDZ2_Kirby_RS0203280 GATMALYMANAGGAWDNAKKFIEKGKLPG------------HAKGSAV------------
MxDZ2_Nan_RS30255     GATMALYMANAGGAWDNAKKFIEKGKLPG------------HAKGSAV------------
MxDZ2_Tam_RS12620     G----------GSAGRLARRRRHDGPLHGQRGRRVGQRQEVHREGQAAGPREGLRGPQGR
                      *          *.*   *::  ..* * *            * :*.*.

MxDZ2_Kirby_RS0203280 ---HKAAVVGDM---VGDPF----KDTSGPGVAILIKVMSVVSLLVASLIALR
MxDZ2_Nan_RS30255     ---HKAAVVGDM---VGDPF----KDTSGPGVAILIKVMSVVSLLVASLIALR
MxDZ2_Tam_RS12620     RRRHGRPVQGHLRSWRGHPHQGHERRVPARGLAHRAAV---------------
                         *  .* *.:    *.*.    : ... *:*    *
```

**Frameshift mutation 3**

| **Tam_location** | **Tam_nucleotide** | **Kirby_nucleotide** | **Nan_nucleotide** | **Nan_location** |
| --- | --- | --- | --- | --- |
| **2736166** | **G (complement)** | **G (complement)** | **.** | **2736150** |

```
multiple sequence alignment (nucleotide)

MxDZ2_Kirby_RS0200050 atgcgaaccctatcaatgcccctcatcgtcgccgccctcctatctggctgcagccagacc
MxDZ2_Tam_RS16225     atgcgaaccctatcaatgcccctcatcgtcgccgccctcctatctggctgcagccagacc
MxDZ2_Nan_RS26665     atgcgaaccctatcaatgcccctcatcgtcgccgccctcctatctggctgcagccagacc
                      ************************************************************

MxDZ2_Kirby_RS0200050 ccagaccccacgccaacgccccaagactcgggcgtcccgcagctggactcgggcgtcccg
MxDZ2_Tam_RS16225     ccagaccccacgccaacgccccaagactcgggcgtcccgcagctggactcgggcgtcccg
MxDZ2_Nan_RS26665     ccagaccccacgccaacgccccaagactcgggcgtcccgcagctggactcgggcgtcccg
                      ************************************************************

MxDZ2_Kirby_RS0200050 ccggaagactcgggcgttccgccagatgacgcgggcgtccctccagatgacgcgggcgtt
MxDZ2_Tam_RS16225     ccggaagactcgggcgttccgccagatgacgcgggcgtccctccagatgacgcgggcgtt
MxDZ2_Nan_RS26665     ccggaagactcgggcgttccgccagatgacgcgggcgtccctccagatgacgcgggcgtt
                      ************************************************************

MxDZ2_Kirby_RS0200050 ccgccagaagatccgcttcccaccggCcccccggtcccgcagggcccgcccaacgtgccc
MxDZ2_Tam_RS16225     ccgccagaagatccgcttcccaccggCcccccggtcccgcagggcccgcccaacgtgccc
MxDZ2_Nan_RS26665     ccgccagaagatccgcttcccaccgg-cccccggtcccgcagggcccgcccaacgtgccc
                      ************************** *********************************

MxDZ2_Kirby_RS0200050 gagttcgatccggccttccccggacagacacgcgtcccagccatccagacgcagacgccc
MxDZ2_Tam_RS16225     gagttcgatccggccttccccggacagacacgcgtcccagccatccagacgcagacgccc
MxDZ2_Nan_RS26665     gagttcgatccggccttccccggacagacacgcgtcccagccatccagacgcagacgccc
                      ************************************************************

MxDZ2_Kirby_RS0200050 atcgaggtcaccgagattgcctcgggcttcaggaatccctgggccatcgccttcctgccc
MxDZ2_Tam_RS16225     atcgaggtcaccgagattgcctcgggcttcaggaatccctgggccatcgccttcctgccc
MxDZ2_Nan_RS26665     atcgaggtcaccgagattgcctcgggcttcaggaatccctgggccatcgccttcctgccc
                      ************************************************************

MxDZ2_Kirby_RS0200050 gaccagcgcatgctggtgacagagaagcccaccggctcgctctacatcgtcacgccgcag
MxDZ2_Tam_RS16225     gaccagcgcatgctggtgacagagaagcccaccggctcgctctacatcgtcacgccgcag
MxDZ2_Nan_RS26665     gaccagcgcatgctggtgacagagaagcccaccggctcgctctacatcgtcacgccgcag
                      ************************************************************

MxDZ2_Kirby_RS0200050 ggcgcgaagtctcccgccgtcgtggggctgcccaacgtggatggccgtggacaggggggc
MxDZ2_Tam_RS16225     ggcgcgaagtctcccgccgtcgtggggctgcccaacgtggatggccgtggacaggggggc
MxDZ2_Nan_RS26665     ggcgcgaagtctcccgccgtcgtggggctgcccaacgtggatggccgtggacaggggggc
                      ************************************************************

MxDZ2_Kirby_RS0200050 ctgctcgacgtggaggtcggccccgactacgcgcagagccagctcatctactggacctat
MxDZ2_Tam_RS16225     ctgctcgacgtggaggtcggccccgactacgcgcagagccagctcatctactggacctat
MxDZ2_Nan_RS26665     ctgctcgacgtggaggtcggccccgactacgcgcagagccagctcatctactggacctat
                      ************************************************************

MxDZ2_Kirby_RS0200050 tccgagccacgccagggcggcaacgggctcgcggtggcgcgtgcgaaactcgtggacggt
MxDZ2_Tam_RS16225     tccgagccacgccagggcggcaacgggctcgcggtggcgcgtgcgaaactcgtggacggt
MxDZ2_Nan_RS26665     tccgagccacgccagggcggcaacgggctcgcggtggcgcgtgcgaaactcgtggacggt
                      ************************************************************

MxDZ2_Kirby_RS0200050 gcgcagcctcgcgtggaggacgtccaggtcatcttccgcatgatgcccacgctcgagtcg
MxDZ2_Tam_RS16225     gcgcagcctcgcgtggaggacgtccaggtcatcttccgcatgatgcccacgctcgagtcg
MxDZ2_Nan_RS26665     gcgcagcctcgcgtggaggacgtccaggtcatcttccgcatgatgcccacgctcgagtcg
                      ************************************************************

MxDZ2_Kirby_RS0200050 acgctgcactcggggggacggctggtattcacccccgacggcaagctgttcgtcacgctc
MxDZ2_Tam_RS16225     acgctgcactcggggggacggctggtattcacccccgacggcaagctgttcgtcacgctc
MxDZ2_Nan_RS26665     acgctgcactcggggggacggctggtattcacccccgacggcaagctgttcgtcacgctc
                      ************************************************************

MxDZ2_Kirby_RS0200050 ggggagcgctccatcctcgcgggccgggtgcaggcgcaggaactgaacagccacttcggc
MxDZ2_Tam_RS16225     ggggagcgctccatcctcgcgggccgggtgcaggcgcaggaactgaacagccacttcggc
MxDZ2_Nan_RS26665     ggggagcgctccatcctcgcgggccgggtgcaggcgcaggaactgaacagccacttcggc
                      ************************************************************

MxDZ2_Kirby_RS0200050 aaggtggtccgcatcaatcccgacggctccgtgccccaggacaacccctacgtgaacacc
MxDZ2_Tam_RS16225     aaggtggtccgcatcaatcccgacggctccgtgccccaggacaacccctacgtgaacacc
MxDZ2_Nan_RS26665     aaggtggtccgcatcaatcccgacggctccgtgccccaggacaacccctacgtgaacacc
                      ************************************************************

MxDZ2_Kirby_RS0200050 gagggggcgaagccggaaatctggtcggtgggccaccgcaacgtcctgtcggcggcgctc
MxDZ2_Tam_RS16225     gagggggcgaagccggaaatctggtcggtgggccaccgcaacgtcctgtcggcggcgctc
MxDZ2_Nan_RS26665     gagggggcgaagccggaaatctggtcggtgggccaccgcaacgtcctgtcggcggcgctc
                      ************************************************************

MxDZ2_Kirby_RS0200050 gacagccagaaccggctgtggacggtcgaaatgggaccgcgcgggggtgacgagctcaac
MxDZ2_Tam_RS16225     gacagccagaaccggctgtggacggtcgaaatgggaccgcgcgggggtgacgagctcaac
MxDZ2_Nan_RS26665     gacagccagaaccggctgtggacggtcgaaatgggaccgcgcgggggtgacgagctcaac
                      ************************************************************

MxDZ2_Kirby_RS0200050 cgccccgaggccggcaaggattacggctggcccaccattgggtatggcgaggagtactcc
MxDZ2_Tam_RS16225     cgccccgaggccggcaaggattacggctggcccaccattgggtatggcgaggagtactcc
MxDZ2_Nan_RS26665     cgccccgaggccggcaaggattacggctggcccaccattgggtatggcgaggagtactcc
                      ************************************************************

MxDZ2_Kirby_RS0200050 ggcgcgcccatccacgagagccctcgtggcccaggcatggagcagcccgtgtactactgg
MxDZ2_Tam_RS16225     ggcgcgcccatccacgagagccctcgtggcccaggcatggagcagcccgtgtactactgg
MxDZ2_Nan_RS26665     ggcgcgcccatccacgagagccctcgtggcccaggcatggagcagcccgtgtactactgg
                      ************************************************************

MxDZ2_Kirby_RS0200050 gacccggtcatttctccctcggggatgaccatctactcggggaccctgttccccgagtgg
MxDZ2_Tam_RS16225     gacccggtcatttctccctcggggatgaccatctactcggggaccctgttccccgagtgg
MxDZ2_Nan_RS26665     gacccggtcatttctccctcggggatgaccatctactcggggaccctgttccccgagtgg
                      ************************************************************

MxDZ2_Kirby_RS0200050 cggaacaacatcttcatcggaggcctgtccagccaggcgctggtgcggctcatgatgcgg
MxDZ2_Tam_RS16225     cggaacaacatcttcatcggaggcctgtccagccaggcgctggtgcggctcatgatgcgg
MxDZ2_Nan_RS26665     cggaacaacatcttcatcggaggcctgtccagccaggcgctggtgcggctcatgatgcgg
                      ************************************************************

MxDZ2_Kirby_RS0200050 aatgaccgcgtggtgggcgaggagcatctcctcaaggacctgggcgtgcgcatccgcgag
MxDZ2_Tam_RS16225     aatgaccgcgtggtgggcgaggagcatctcctcaaggacctgggcgtgcgcatccgcgag
MxDZ2_Nan_RS26665     aatgaccgcgtggtgggcgaggagcatctcctcaaggacctgggcgtgcgcatccgcgag
                      ************************************************************

MxDZ2_Kirby_RS0200050 gtggtgcagggccccgacggagccctctacctgctcaccgacgccaccaacggaaagctg
MxDZ2_Tam_RS16225     gtggtgcagggccccgacggagccctctacctgctcaccgacgccaccaacggaaagctg
MxDZ2_Nan_RS26665     gtggtgcagggccccgacggagccctctacctgctcaccgacgccaccaacggaaagctg
                      ************************************************************

MxDZ2_Kirby_RS0200050 ctcaaggtcacgccgcgctga
MxDZ2_Tam_RS16225     ctcaaggtcacgccgcgctga
MxDZ2_Nan_RS26665     ctcaaggtcacgccgcgctga
                      *********************

-----------------------------------------------------------------------------------
multiple sequence alignment (protein)


MxDZ2_Kirby_RS0200050 MRTLSMPLIVAALLSGCSQTPDPTPTPQDSGVPQLDSGVPPEDSGVPPDDAGVPPDDAGV
MxDZ2_Tam_RS16225     MRTLSMPLIVAALLSGCSQTPDPTPTPQDSGVPQLDSGVPPEDSGVPPDDAGVPPDDAGV
MxDZ2_Nan_RS26665     MRTLSMPLIVAALLSGCSQTPDPTPTPQDSGVPQLDSGVPPEDSGVPPDDAGVPPDDAGV
                      ************************************************************

MxDZ2_Kirby_RS0200050 PPEDPLPTGPPVPQGPPNV-------PEFDPAFPGQTRVPAIQTQTPIEVTEIASGFRNP
MxDZ2_Tam_RS16225     PPEDPLPTGPPVPQGPPNV-------PEFDPAFPGQTRVPAIQTQTPIEVTEIASGFRNP
MxDZ2_Nan_RS26665     PPEDPLPTGPRSRRARPTCPSSIRPSPDRHASQPSRRRRPSRSPRLP-----RASGIPGP
                      **********   :. *.        *: ..: *.: * *: ..: *      ***: .*

MxDZ2_Kirby_RS0200050 WAIAFLPDQRMLVTEKPTGSLYIVTPQGAKS---------PAVVGLPNVDGRGQGGLLDV
MxDZ2_Tam_RS16225     WAIAFLPDQRMLVTEKPTGSLYIVTPQGAKS---------PAVVGLPNVDGRGQGGLLDV
MxDZ2_Nan_RS26665     ------------SPSCPTSACWQRSPPARSTSSRRRARSLPPSWGCPTWMAVDRGACSTW
                                   .. **.: :  :* . .:         *.  * *.  . .:*.

MxDZ2_Kirby_RS0200050 EVGPDYAQSQLIYWTYSEPRQGGNGLAVARAKLVDGAQPRVED---VQVIFRMMPTLEST
MxDZ2_Tam_RS16225     EVGPDYAQSQLIYWTYSEPRQGGNGLAVARAKLVDGAQPRVED---VQVIFRMMPTLEST
MxDZ2_Nan_RS26665     RSAPTTRRA-----------SSSTGPIPSHARAATGSRWRVRNSWTVRSLAWRTSRSSSA
                      . .*   ::           ....*   ::*: . *:: **.:   *: :    .  .*:

MxDZ2_Kirby_RS0200050 LHSGGRLVFTPDG---KLFVTLGERSILAGRVQAQELNSHFGKVVRIN---PDGSVPQDN
MxDZ2_Tam_RS16225     LHSGGRLVFTPDG---KLFVTLGERSILAGRVQAQELNSHFGKVVRIN---PDGSVPQDN
MxDZ2_Nan_RS26665     CPRSSRRCTRGDGWYSPPTASCSSRSGSAPSSRAGCRRRNTATSARWSASIPTAPCPRTT
                         ..*     **      .: ..**  *   :*   . : .. .* .   * .. *: .

MxDZ2_Kirby_RS0200050 PYV-NTEGAKPEIWSVGH--RNVLSAALDSQNRLWTVEMGPRGGDELNRPEAGKDYGWPT
MxDZ2_Tam_RS16225     PYV-NTEGAKPEIWSVGH--RNVLSAALDSQNRLWTVEMGPRGGDELNRPEAGKDYGWPT
MxDZ2_Nan_RS26665     PTTPRGRSRKSGRWATATSCRRRSTARTGCGRSKWDRA----GVTSSTAPRPARITAGPP
                      * . . .. *.  *:..   *.  :*  .. .  *       *  . . *...:  . *.

MxDZ2_Kirby_RS0200050 IGYGEEYSGAPIHES------------------PRGPGMEQPVYYWDPVISPSGMTIYSG
MxDZ2_Tam_RS16225     IGYGEEYSGAPIHES------------------PRGPGMEQPVYYWDPVISPSGMTIYSG
MxDZ2_Nan_RS26665     LGMARSTPARPSTRALVAQAWSSPCTTGTRSFLPRGPSTRGPC-------SPSGGTTSSS
                      :* ... .. *  .:                  ****. . *        **** *  *.


MxDZ2_Kirby_RS0200050 TLFPEWR-------NNIFIGGLSSQALVRLMMRNDRVVGEEHLLKDLGVRIREVVQGPDG
MxDZ2_Tam_RS16225     TLFPEWR-------NNIFIGGLSSQALVRLMMRNDRVVGEEHLLKDLGVRIREVVQGPDG
MxDZ2_Nan_RS26665     EACPARRWCGSCGMTAWWARSISSRTWACASARWCRA-----------------------
                         *  *       .  :  .:**:: .    *  *.

MxDZ2_Kirby_RS0200050 ALYLLTDATNGKLLKVTP----------R
MxDZ2_Tam_RS16225     ALYLLTDATNGKLLKVTP----------R
MxDZ2_Nan_RS26665     -------PTEPSTCSPTPPTESCSRSRRA
                             .*: .  . **
```

**Frameshift mutation 4**

| **Tam_location** | **Tam_nucleotide** | **Kirby_nucleotide** | **Nan_nucleotide** | **Nan_location** |
| --- | --- | --- | --- | --- |
| **3936061** | **C** | **C** | **.** | **3936043** |

```
multiple sequence alignment (nucleotide)

MxDZ2_Kirby_RS0225565 atgaacgtgatttcgttgtttccgccccctggaacgaccatcgacggatggagtgttgtt
MxDZ2_Tam_RS21100     atgaacgtgatttcgttgtttccgccccctggaacgaccatcgacggatggagtgttgtt
MxDZ2_Nan_RS21765     atgaacgtgatttcgttgtttccgccccctggaacgaccatcgacggatggagtgttgtt
                      ************************************************************

MxDZ2_Kirby_RS0225565 cgggagcttggaaacggagggtttgcagtcgtctacctcgtcgagaagcacggtctcaga
MxDZ2_Tam_RS21100     cgggagcttggaaacggagggtttgcagtcgtctacctcgtcgagaagcacggtctcaga
MxDZ2_Nan_RS21765     cgggagcttggaaacggagggtttgcagtcgtctacctcgtcgagaagcacggtctcaga
                      ************************************************************

MxDZ2_Kirby_RS0225565 tgcgcgctcaagttggcgcgccaccgggattcgagtggggacgacaagcagactcacgca
MxDZ2_Tam_RS21100     tgcgcgctcaagttggcgcgccaccgggattcgagtggggacgacaagcagactcacgca
MxDZ2_Nan_RS21765     tgcgcgctcaagttggcgcgccaccgggattcgagtggggacgacaagcagactcacgca
                      ************************************************************

MxDZ2_Kirby_RS0225565 cggacgcttcgggagctttcggccctcctcctcctggaccatccgaacatcgtcaagcac
MxDZ2_Tam_RS21100     cggacgcttcgggagctttcggccctcctcctcctggaccatccgaacatcgtcaagcac
MxDZ2_Nan_RS21765     cggacgcttcgggagctttcggccctcctcctcctggaccatccgaacatcgtcaagcac
                      ************************************************************

MxDZ2_Kirby_RS0225565 cgtgggtatggatactctgagcaggggaatgtctatctcgcgcttgagtacgtagatggg
MxDZ2_Tam_RS21100     cgtgggtatggatactctgagcaggggaatgtctatctcgcgcttgagtacgtagatggg
MxDZ2_Nan_RS21765     cgtgggtatggatactctgagcaggggaatgtctatctcgcgcttgagtacgtagatggg
                      ************************************************************

MxDZ2_Kirby_RS0225565 tggacccttgccgaatgggcggagcgtaaacaccccacggttcaggaggttttgcatgtc
MxDZ2_Tam_RS21100     tggacccttgccgaatgggcggagcgtaaacaccccacggttcaggaggttttgcatgtc
MxDZ2_Nan_RS21765     tggacccttgccgaatgggcggagcgtaaacaccccacggttcaggaggttttgcatgtc
                      ************************************************************

MxDZ2_Kirby_RS0225565 ttcgacaagatttccgccgcgctttcgtacatgcacggccgtggtgtcctgcatcgggat
MxDZ2_Tam_RS21100     ttcgacaagatttccgccgcgctttcgtacatgcacggccgtggtgtcctgcatcgggat
MxDZ2_Nan_RS21765     ttcgacaagatttccgccgcgctttcgtacatgcacggccgtggtgtcctgcatcgggat
                      ************************************************************

MxDZ2_Kirby_RS0225565 ttgaaactgtccaacgttctgattcggaagagcgatggagagccggtcatcatcgacttt
MxDZ2_Tam_RS21100     ttgaaactgtccaacgttctgattcggaagagcgatggagagccggtcatcatcgacttt
MxDZ2_Nan_RS21765     ttgaaactgtccaacgttctgattcggaagagcgatggagagccggtcatcatcgacttt
                      ************************************************************

MxDZ2_Kirby_RS0225565 agctgtgcaagctactcgttggccgaagagctgacggattggggcttgccgccgggaact
MxDZ2_Tam_RS21100     agctgtgcaagctactcgttggccgaagagctgacggattggggcttgccgccgggaact
MxDZ2_Nan_RS21765     agctgtgcaagctactcgttggccgaagagctgacggattggggcttgccgccgggaact
                      ************************************************************

MxDZ2_Kirby_RS0225565 gaccgctttcgtgcgccggaacagttcacatggctccgggagcacaaggccgaacagcga
MxDZ2_Tam_RS21100     gaccgctttcgtgcgccggaacagttcacatggctccgggagcacaaggccgaacagcga
MxDZ2_Nan_RS21765     gaccgctttcgtgcgccggaacagttcacatggctccgggagcacaaggccgaacagcga
                      ************************************************************

MxDZ2_Kirby_RS0225565 gcgaagtacgccttccaagttgcggacgagattttcgccgtcggggcgatgctctatgag
MxDZ2_Tam_RS21100     gcgaagtacgccttccaagttgcggacgagattttcgccgtcggggcgatgctctatgag
MxDZ2_Nan_RS21765     gcgaagtacgccttccaagttgcggacgagattttcgccgtcggggcgatgctctatgag
                      ************************************************************

MxDZ2_Kirby_RS0225565 ttgctgaccgacccccgaccgacggaggttcaagcgcgagttacgctcaacagcaccgtc
MxDZ2_Tam_RS21100     ttgctgaccgacccccgaccgacggaggttcaagcgcgagttacgctcaacagcaccgtc
MxDZ2_Nan_RS21765     ttgctgaccgacccccgaccgacggaggttcaagcgcgagttacgctcaacagcaccgtc
                      ************************************************************

MxDZ2_Kirby_RS0225565 atgaagccgcctcctgcgcgtgcgttgaacgtgcgtgttccggaagcgctgaacgacctc
MxDZ2_Tam_RS21100     atgaagccgcctcctgcgcgtgcgttgaacgtgcgtgttccggaagcgctgaacgacctc
MxDZ2_Nan_RS21765     atgaagccgcctcctgcgcgtgcgttgaacgtgcgtgttccggaagcgctgaacgacctc
                      ************************************************************

MxDZ2_Kirby_RS0225565 gttgattgcatcctgtcgcgtgaaccggcaaggcgccctgtcgacactgaggcgttgcgc
MxDZ2_Tam_RS21100     gttgattgcatcctgtcgcgtgaaccggcaaggcgccctgtcgacactgaggcgttgcgc
MxDZ2_Nan_RS21765     gttgattgcatcctgtcgcgtgaaccggcaaggcgccctgtcgacactgaggcgttgcgc
                      ************************************************************

MxDZ2_Kirby_RS0225565 cgggagctgggcgaactcctggcctattcgagcgcggagtacctatccccggtgcatccg
MxDZ2_Tam_RS21100     cgggagctgggcgaactcctggcctattcgagcgcggagtacctatccccggtgcatccg
MxDZ2_Nan_RS21765     cgggagctgggcgaactcctggcctattcgagcgcggagtacctatccccggtgcatccg
                      ************************************************************

MxDZ2_Kirby_RS0225565 ccgtccgaacagcggccattggagccgccggatcaggtgatgcccgaagttgccaacccg
MxDZ2_Tam_RS21100     ccgtccgaacagcggccattggagccgccggatcaggtgatgcccgaagttgccaacccg
MxDZ2_Nan_RS21765     ccgtccgaacagcggccattggagccgccggatcaggtgatgcccgaagttgccaacccg
                      ************************************************************

MxDZ2_Kirby_RS0225565 cgccttcctgtgtcgccaacgcgttctggtcgggagaggtggggactcgcggcgggcttg
MxDZ2_Tam_RS21100     cgccttcctgtgtcgccaacgcgttctggtcgggagaggtggggactcgcggcgggcttg
MxDZ2_Nan_RS21765     cgccttcctgtgtcgccaacgcgttctggtcgggagaggtggggactcgcggcgggcttg
                      ************************************************************

MxDZ2_Kirby_RS0225565 gctgccctcatcgcgcttaccgtggccgggagcttcttgctcagccgaggggaacccacg
MxDZ2_Tam_RS21100     gctgccctcatcgcgcttaccgtggccgggagcttcttgctcagccgaggggaacccacg
MxDZ2_Nan_RS21765     gctgccctcatcgcgcttaccgtggccgggagcttcttgctcagccgaggggaacccacg
                      ************************************************************

MxDZ2_Kirby_RS0225565 gagtccggggcgcagaccgtcgttggtgtgtcgcgcccgcagatccctccacattccgcg
MxDZ2_Tam_RS21100     gagtccggggcgcagaccgtcgttggtgtgtcgcgcccgcagatccctccacattccgcg
MxDZ2_Nan_RS21765     gagtccggggcgcagaccgtcgttggtgtgtcgcgcccgcagatccctccacattccgcg
                      ************************************************************

MxDZ2_Kirby_RS0225565 ccgctcacgtcgactgcccctgctatgtcaccaccaagCcccccgcccatgacgctgacg
MxDZ2_Tam_RS21100     ccgctcacgtcgactgcccctgctatgtcaccaccaagCcccccgcccatgacgctgacg
MxDZ2_Nan_RS21765     ccgctcacgtcgactgcccctgctatgtcaccaccaag-cccccgcccatgacgctgacg
                      ************************************** *********************

MxDZ2_Kirby_RS0225565 ggccttgcgactgccgttccgaaggaaggttcaaccgtgaagacgcctccgtcacctgag
MxDZ2_Tam_RS21100     ggccttgcgactgccgttccgaaggaaggttcaaccgtgaagacgcctccgtcacctgag
MxDZ2_Nan_RS21765     ggccttgcgactgccgttccgaaggaaggttcaaccgtgaagacgcctccgtcacctgag
                      ************************************************************

MxDZ2_Kirby_RS0225565 tccccactccaaggacgcccgtcacgcgggagaacgaaggccgccgctgctgccgactgc
MxDZ2_Tam_RS21100     tccccactccaaggacgcccgtcacgcgggagaacgaaggccgccgctgctgccgactgc
MxDZ2_Nan_RS21765     tccccactccaaggacgcccgtcacgcgggagaacgaaggccgccgctgctgccgactgc
                      ************************************************************

MxDZ2_Kirby_RS0225565 gcgacgatgactctcgttgcggcactcgcggcaggttgccccagtgcccagattcgacct
MxDZ2_Tam_RS21100     gcgacgatgactctcgttgcggcactcgcggcaggttgccccagtgcccagattcgacct
MxDZ2_Nan_RS21765     gcgacgatgactctcgttgcggcactcgcggcaggttgccccagtgcccagattcgacct
                      ************************************************************


MxDZ2_Kirby_RS0225565 gaagcgttcacttgcccggctggtgcggaggaggtgatgcaggaggatctccgctggaag
MxDZ2_Tam_RS21100     gaagcgttcacttgcccggctggtgcggaggaggtgatgcaggaggatctccgctggaag
MxDZ2_Nan_RS21765     gaagcgttcacttgcccggctggtgcggaggaggtgatgcaggaggatctccgctggaag
                      ************************************************************

MxDZ2_Kirby_RS0225565 gtgaatcagagtttcgcgctcaccttggatgcccgccatgagacggacgcctacgtttgg
MxDZ2_Tam_RS21100     gtgaatcagagtttcgcgctcaccttggatgcccgccatgagacggacgcctacgtttgg
MxDZ2_Nan_RS21765     gtgaatcagagtttcgcgctcaccttggatgcccgccatgagacggacgcctacgtttgg
                      ************************************************************

MxDZ2_Kirby_RS0225565 ttcactgcgggggcggaggtgatgggggtcgttccaaagggcgttccatcagcccaaagg
MxDZ2_Tam_RS21100     ttcactgcgggggcggaggtgatgggggtcgttccaaagggcgttccatcagcccaaagg
MxDZ2_Nan_RS21765     ttcactgcgggggcggaggtgatgggggtcgttccaaagggcgttccatcagcccaaagg
                      ************************************************************

MxDZ2_Kirby_RS0225565 gcggtcgcccctcccggaacgcgcttctacggcaaggcatacttcctttccgatcgaatg
MxDZ2_Tam_RS21100     gcggtcgcccctcccggaacgcgcttctacggcaaggcatacttcctttccgatcgaatg
MxDZ2_Nan_RS21765     gcggtcgcccctcccggaacgcgcttctacggcaaggcatacttcctttccgatcgaatg
                      ************************************************************

MxDZ2_Kirby_RS0225565 ggccgctctgaggggcctgcgctggtcatccgctacgatcgtgtgaagctccctgggcag
MxDZ2_Tam_RS21100     ggccgctctgaggggcctgcgctggtcatccgctacgatcgtgtgaagctccctgggcag
MxDZ2_Nan_RS21765     ggccgctctgaggggcctgcgctggtcatccgctacgatcgtgtgaagctccctgggcag
                      ************************************************************

MxDZ2_Kirby_RS0225565 gacgagcgcccggtttgcttcgtcgtcgagtcgccttccaaggggtacgaagatggcagg
MxDZ2_Tam_RS21100     gacgagcgcccggtttgcttcgtcgtcgagtcgccttccaaggggtacgaagatggcagg
MxDZ2_Nan_RS21765     gacgagcgcccggtttgcttcgtcgtcgagtcgccttccaaggggtacgaagatggcagg
                      ************************************************************

MxDZ2_Kirby_RS0225565 gtgaaggcgtacaacttaggtggcggctacgtcgtagaccgttggccctga
MxDZ2_Tam_RS21100     gtgaaggcgtacaacttaggtggcggctacgtcgtagaccgttggccctga
MxDZ2_Nan_RS21765     gtgaaggcgtacaacttaggtggcggctacgtcgtagaccgttggccctga
                      ***************************************************


-----------------------------------------------------------------------------------


multiple sequence alignment (protein)


MxDZ2_Kirby_RS0225565 MNVISLFPPPGTTIDGWSVVRELGNGGFAVVYLVEKHGLRCALKLARHRDSSGDDKQTHA
MxDZ2_Tam_RS21100     MNVISLFPPPGTTIDGWSVVRELGNGGFAVVYLVEKHGLRCALKLARHRDSSGDDKQTHA
MxDZ2_Nan_RS21765     MNVISLFPPPGTTIDGWSVVRELGNGGFAVVYLVEKHGLRCALKLARHRDSSGDDKQTHA
                      ************************************************************

MxDZ2_Kirby_RS0225565 RTLRELSALLLLDHPNIVKHRGYGYSEQGNVYLALEYVDGWTLAEWAERKHPTVQEVLHV
MxDZ2_Tam_RS21100     RTLRELSALLLLDHPNIVKHRGYGYSEQGNVYLALEYVDGWTLAEWAERKHPTVQEVLHV
MxDZ2_Nan_RS21765     RTLRELSALLLLDHPNIVKHRGYGYSEQGNVYLALEYVDGWTLAEWAERKHPTVQEVLHV
                      ************************************************************

MxDZ2_Kirby_RS0225565 FDKISAALSYMHGRGVLHRDLKLSNVLIRKSDGEPVIIDFSCASYSLAEELTDWGLPPGT
MxDZ2_Tam_RS21100     FDKISAALSYMHGRGVLHRDLKLSNVLIRKSDGEPVIIDFSCASYSLAEELTDWGLPPGT
MxDZ2_Nan_RS21765     FDKISAALSYMHGRGVLHRDLKLSNVLIRKSDGEPVIIDFSCASYSLAEELTDWGLPPGT
                      ************************************************************

MxDZ2_Kirby_RS0225565 DRFRAPEQFTWLREHKAEQRAKYAFQVADEIFAVGAMLYELLTDPRPTEVQARVTLNSTV
MxDZ2_Tam_RS21100     DRFRAPEQFTWLREHKAEQRAKYAFQVADEIFAVGAMLYELLTDPRPTEVQARVTLNSTV
MxDZ2_Nan_RS21765     DRFRAPEQFTWLREHKAEQRAKYAFQVADEIFAVGAMLYELLTDPRPTEVQARVTLNSTV
                      ************************************************************

MxDZ2_Kirby_RS0225565 MKPPPARALNVRVPEALNDLVDCILSREPARRPVDTEALRRELGELLAYSSAEYLSPVHP
MxDZ2_Tam_RS21100     MKPPPARALNVRVPEALNDLVDCILSREPARRPVDTEALRRELGELLAYSSAEYLSPVHP
MxDZ2_Nan_RS21765     MKPPPARALNVRVPEALNDLVDCILSREPARRPVDTEALRRELGELLAYSSAEYLSPVHP
                      ************************************************************

MxDZ2_Kirby_RS0225565 PSEQRPLEPPDQVMPEVANPRLPVSPTRSGRERWGLAAGLAALIALTVAGSFLLSRGEPT
MxDZ2_Tam_RS21100     PSEQRPLEPPDQVMPEVANPRLPVSPTRSGRERWGLAAGLAALIALTVAGSFLLSRGEPT
MxDZ2_Nan_RS21765     PSEQRPLEPPDQVMPEVANPRLPVSPTRSGRERWGLAAGLAALIALTVAGSFLLSRGEPT
                      ************************************************************

MxDZ2_Kirby_RS0225565 ESGAQTVVGVSRPQIPPHSAPLTSTAPAMSPPSPPPMTLTGLA-------------TAVP
MxDZ2_Tam_RS21100     ESGAQTVVGVSRPQIPPHSAPLTSTAPAMSPPSPPPMTLTGLA-------------TAVP
MxDZ2_Nan_RS21765     ESGAQTVVGVSRPQIPPHSAPLTSTAPAMSPPSPRPRRALRLPFRRKVQPRRLRHLSPHS
                      ********************************** *     *.             :. .


MxDZ2_Kirby_RS0225565 KEGSTVKTPPSPESPLQGRPSRGRTKAAAAA-----DCATMTLVAALAAGCPSAQIR-PE
MxDZ2_Tam_RS21100     KEGSTVKTPPSPESPLQGRPSRGRTKAAAAA-----DCATMTLVAALAAGCPSAQIR-PE
MxDZ2_Nan_RS21765     KDARHAGERRPPLLPTARRLSLRHSRQVAPVPRFDLKRSLARLVRRRCRRISAGRIRVSR
                      *:.  .    .*  *   * *  ::: .*..     . :   **   .   .:.:** ..

MxDZ2_Kirby_RS0225565 AFTCPAGAEEVMQEDLRWKVNQSFALTL--------------DARHETDAYVWFTAG---
MxDZ2_Tam_RS21100     AFTCPAGAEEVMQEDLRWKVNQSFALTL--------------DARHETDAYVWFTAG---
MxDZ2_Nan_RS21765     SPWMPAMRRTPTFGSLRGRRWGSFQRAFHQPKGRSPLPERASTARHTSFPIEWAALRGLR
                      :   **  .     .** :   **  ::               *** : .  * :

MxDZ2_Kirby_RS0225565 -AEVMGVVPKGVPSAQRAVA----PPGTRFYGKAYFLSDRMGRSEGPALVIRYDRVKLPG
MxDZ2_Tam_RS21100     -AEVMGVVPKGVPSAQRAVA----PPGTRFYGKAYFLSDRMGRSEGPALVIRYDRVKLPG
MxDZ2_Nan_RS21765     WSSATIVSSLGRTSARFASSSSRLPRGTKMAGR---------RTT---------------
                       :..  * . * .**: * :    * **:: *:         *:

MxDZ2_Kirby_RS0225565 QDERPVCFVVESPSKGYEDGRVKAYNLGGGYVVDRWP
MxDZ2_Tam_RS21100     QDERPVCFVVESPSKGYEDGRVKAYNLGGGYVVDRWP
MxDZ2_Nan_RS21765     --------VAATSTVG--------------------P
                              *. :.: *                    *
```

**Frameshift mutation 5**

| **Tam_location** | **Tam_nucleotide** | **Kirby_nucleotide** | **Nan_nucleotide** | **Nan_location** |
| --- | --- | --- | --- | --- |
| **4389880** | **C** | **C** | **.** | **4389859** |

```
multiple sequence alignment (nucleotide)

MxDZ2_Kirby_RS0218355 atgcggaaggttctgctgttggtgatggtgtccacgttcgctgggtgtgcgaagcgtcag
MxDZ2_Tam_RS23100     atgcggaaggttctgctgttggtgatggtgtccacgttcgctgggtgtgcgaagcgtcag
MxDZ2_Nan_RS19765     atgcggaaggttctgctgttggtgatggtgtccacgttcgctgggtgtgcgaagcgtcag
                      ************************************************************

MxDZ2_Kirby_RS0218355 gagcccgagtccctgctcaaggcccgcgagctgatggcggaagcccagaatcccagcggc
MxDZ2_Tam_RS23100     gagcccgagtccctgctcaaggcccgcgagctgatggcggaagcccagaatcccagcggc
MxDZ2_Nan_RS19765     gagcccgagtccctgctcaaggcccgcgagctgatggcggaagcccagaatcccagcggc
                      ************************************************************

MxDZ2_Kirby_RS0218355 aacctggcactgctctgcgagcccgtggacgcggaggtgtatctagatggcgtctggcag
MxDZ2_Tam_RS23100     aacctggcactgctctgcgagcccgtggacgcggaggtgtatctagatggcgtctggcag
MxDZ2_Nan_RS19765     aacctggcactgctctgcgagcccgtggacgcggaggtgtatctagatggcgtctggcag
                      ************************************************************

MxDZ2_Kirby_RS0218355 gggctctgcagcgacttctcgggttCccccaaggcgctgcgagtgggcagcggcctgcac
MxDZ2_Tam_RS23100     gggctctgcagcgacttctcgggttCccccaaggcgctgcgagtgggcagcggcctgcac
MxDZ2_Nan_RS19765     gggctctgcagcgacttctcgggtt-ccccaaggcgctgcgagtgggcagcggcctgcac
                      ************************* **********************************

MxDZ2_Kirby_RS0218355 gagattgaagtgaagaagcaggggtactggccctatacgacgtacttcgagcccagccgg
MxDZ2_Tam_RS23100     gagattgaagtgaagaagcaggggtactggccctatacgacgtacttcgagcccagccgg
MxDZ2_Nan_RS19765     gagattgaagtgaagaagcaggggtactggccctatacgacgtacttcgagcccagccgg
                      ************************************************************

MxDZ2_Kirby_RS0218355 gcccgcgcgcggctgaccatccagctccgcgcctctgggccgaaggccggtggttcggag
MxDZ2_Tam_RS23100     gcccgcgcgcggctgaccatccagctccgcgcctctgggccgaaggccggtggttcggag
MxDZ2_Nan_RS19765     gcccgcgcgcggctgaccatccagctccgcgcctctgggccgaaggccggtggttcggag
                      ************************************************************

MxDZ2_Kirby_RS0218355 tga
MxDZ2_Tam_RS23100     tga
MxDZ2_Nan_RS19765     tga
                      ***

-----------------------------------------------------------------------------------
multiple sequence alignment (protein)


MxDZ2_Kirby_RS0218355 MRKVLLLVMVSTFAGCAKRQEPESLLKARELMAEAQNPSGNLALLCEPVDAEVYLDGVWQ
MxDZ2_Tam_RS23100     MRKVLLLVMVSTFAGCAKRQEPESLLKARELMAEAQNPSGNLALLCEPVDAEVYLDGVWQ
MxDZ2_Nan_RS19765     MRKVLLLVMVSTFAGCAKRQEPESLLKARELMAEAQNPSGNLALLCEPVDAEVYLDGVWQ
                      ************************************************************

MxDZ2_Kirby_RS0218355 GLCSDFSGSPKALRVGSGLHEIEVKKQGYWPYTTYFEPSRARARLTIQLRASGPKAGGSE
MxDZ2_Tam_RS23100     GLCSDFSGSPKALRVGSGLHEIEVKKQGYWPYTTYFEPSRARARLTIQLRASGPKAGGSE
MxDZ2_Nan_RS19765     GLCSDFSGSPRRCEWAAA--------------CTRLKRSRGTGPIRRTSSPAGPARGPSS
                      **********:  . .:.               * :: **. . :     .:**  * *.

MxDZ2_Kirby_RS0218355 ------------
MxDZ2_Tam_RS23100     ------------
MxDZ2_Nan_RS19765     SAPLGRRPVVRS
```

**Frameshift mutation 6**

| **Tam_location** | **Tam_nucleotide** | **Kirby_nucleotide** | **Nan_nucleotide** | **Nan_location** |
| --- | --- | --- | --- | --- |
| **5376775** | **G (complement)** | **G (complement)** | **.** | **5376752** |

```
multiple sequence alignment (nucleotide)

MxDZ2_Kirby_RS0232470 atggccaagcccaagtccggggccaagaagacgactcccgctggcaaggctggcgcaaag
MxDZ2_Tam_RS26190     atggccaagcccaagtccggggccaagaagacgactcccgctggcaaggctggcgcaaag
MxDZ2_Nan_RS16675     atggccaagcccaagtccggggccaagaagacgactcccgctggcaaggctggcgcaaag
                      ************************************************************

MxDZ2_Kirby_RS0232470 cccgctgcgaagaaggactccgctgcccggctggatctgatcaagaacgcctccaagcgg
MxDZ2_Tam_RS26190     cccgctgcgaagaaggactccgctgcccggctggatctgatcaagaacgcctccaagcgg
MxDZ2_Nan_RS16675     cccgctgcgaagaaggactccgctgcccggctggatctgatcaagaacgcctccaagcgg
                      ************************************************************

MxDZ2_Kirby_RS0232470 gcggcgaaggcggcgacgaaattggcgaaagcggcatcggacgtaacgaagggggccaag
MxDZ2_Tam_RS26190     gcggcgaaggcggcgacgaaattggcgaaagcggcatcggacgtaacgaagggggccaag
MxDZ2_Nan_RS16675     gcggcgaaggcggcgacgaaattggcgaaagcggcatcggacgtaacgaagggggccaag
                      ************************************************************

MxDZ2_Kirby_RS0232470 agcgcggtgaagaaggccgtgcaggagaaggccaccgcgaagaaggccgcgcccgctggg
MxDZ2_Tam_RS26190     agcgcggtgaagaaggccgtgcaggagaaggccaccgcgaagaaggccgcgcccgctggg
MxDZ2_Nan_RS16675     agcgcggtgaagaaggccgtgcaggagaaggccaccgcgaagaaggccgcgcccgctggg
                      ************************************************************

MxDZ2_Kirby_RS0232470 aagacggcctcggcagcgaaggccgCccccccggcagcgaagaccccgaaggcacctccg
MxDZ2_Tam_RS26190     aagacggcctcggcagcgaaggccgCccccccggcagcgaagaccccgaaggcacctccg
MxDZ2_Nan_RS16675     aagacggcctcggcagcgaaggccg-ccccccggcagcgaagaccccgaaggcacctccg
                      ************************* **********************************

MxDZ2_Kirby_RS0232470 gcgaaggcgcctcccgcggcgaagtccgccgcgagcgccaaggcggcgaaggcgagcgcg
MxDZ2_Tam_RS26190     gcgaaggcgcctcccgcggcgaagtccgccgcgagcgccaaggcggcgaaggcgagcgcg
MxDZ2_Nan_RS16675     gcgaaggcgcctcccgcggcgaagtccgccgcgagcgccaaggcggcgaaggcgagcgcg
                      ************************************************************

MxDZ2_Kirby_RS0232470 gtacccgccgcccctccggtggagaagccccgtcctcgcgccaccaagttgccgcctccg
MxDZ2_Tam_RS26190     gtacccgccgcccctccggtggagaagccccgtcctcgcgccaccaagttgccgcctccg
MxDZ2_Nan_RS16675     gtacccgccgcccctccggtggagaagccccgtcctcgcgccaccaagttgccgcctccg
                      ************************************************************

MxDZ2_Kirby_RS0232470 ggcgagccgctcaccaagcgtgaaatggagcagttgctgacggccggcgaaggccgcggc
MxDZ2_Tam_RS26190     ggcgagccgctcaccaagcgtgaaatggagcagttgctgacggccggcgaaggccgcggc
MxDZ2_Nan_RS16675     ggcgagccgctcaccaagcgtgaaatggagcagttgctgacggccggcgaaggccgcggc
                      ************************************************************

MxDZ2_Kirby_RS0232470 gtcacgggcgagggcagcctcaagggccgcctggtggtcaccaatgacatgccgcacctg
MxDZ2_Tam_RS26190     gtcacgggcgagggcagcctcaagggccgcctggtggtcaccaatgacatgccgcacctg
MxDZ2_Nan_RS16675     gtcacgggcgagggcagcctcaagggccgcctggtggtcaccaatgacatgccgcacctg
                      ************************************************************

MxDZ2_Kirby_RS0232470 gtggtcgtcggccgcgacaagcgcgagctgaccttcctcctccagggcccggatcaggaa
MxDZ2_Tam_RS26190     gtggtcgtcggccgcgacaagcgcgagctgaccttcctcctccagggcccggatcaggaa
MxDZ2_Nan_RS16675     gtggtcgtcggccgcgacaagcgcgagctgaccttcctcctccagggcccggatcaggaa
                      ************************************************************

MxDZ2_Kirby_RS0232470 gtcctcccggcctacgtggaccacaaggtctccgtcagcgggctcatccgtaagacgacc
MxDZ2_Tam_RS26190     gtcctcccggcctacgtggaccacaaggtctccgtcagcgggctcatccgtaagacgacc
MxDZ2_Nan_RS16675     gtcctcccggcctacgtggaccacaaggtctccgtcagcgggctcatccgtaagacgacc
                      ************************************************************

MxDZ2_Kirby_RS0232470 aaccacgctggcgtggtggacgttcgcaagtactcggccaagaagccggatgccgaggtg
MxDZ2_Tam_RS26190     aaccacgctggcgtggtggacgttcgcaagtactcggccaagaagccggatgccgaggtg
MxDZ2_Nan_RS16675     aaccacgctggcgtggtggacgttcgcaagtactcggccaagaagccggatgccgaggtg
                      ************************************************************

MxDZ2_Kirby_RS0232470 gtggaggccgccccggcggagacggaggcgcggctgcgctacctgtcgcccggtgaggtc
MxDZ2_Tam_RS26190     gtggaggccgccccggcggagacggaggcgcggctgcgctacctgtcgcccggtgaggtc
MxDZ2_Nan_RS16675     gtggaggccgccccggcggagacggaggcgcggctgcgctacctgtcgcccggtgaggtc
                      ************************************************************

MxDZ2_Kirby_RS0232470 tccatggtgacggcggccggcatgggcgcgggcatcaagggcttcgcgggcgtgcgtggc
MxDZ2_Tam_RS26190     tccatggtgacggcggccggcatgggcgcgggcatcaagggcttcgcgggcgtgcgtggc
MxDZ2_Nan_RS16675     tccatggtgacggcggccggcatgggcgcgggcatcaagggcttcgcgggcgtgcgtggc
                      ************************************************************

MxDZ2_Kirby_RS0232470 aacctggagatgacgggcgaggagttcgtgctcgtcgtgtccaatggcggcacgcgccag
MxDZ2_Tam_RS26190     aacctggagatgacgggcgaggagttcgtgctcgtcgtgtccaatggcggcacgcgccag
MxDZ2_Nan_RS16675     aacctggagatgacgggcgaggagttcgtgctcgtcgtgtccaatggcggcacgcgccag
                      ************************************************************

MxDZ2_Kirby_RS0232470 caggtatcgttcatcatcgagggcaaggccgctggcaaggccctgcgcaagcacgtcggg
MxDZ2_Tam_RS26190     caggtatcgttcatcatcgagggcaaggccgctggcaaggccctgcgcaagcacgtcggg
MxDZ2_Nan_RS16675     caggtatcgttcatcatcgagggcaaggccgctggcaaggccctgcgcaagcacgtcggg
                      ************************************************************

MxDZ2_Kirby_RS0232470 tacacgctccaggtgcagggcgtcgtcgacaagacgtccggctggggcggccgcatcatg
MxDZ2_Tam_RS26190     tacacgctccaggtgcagggcgtcgtcgacaagacgtccggctggggcggccgcatcatg
MxDZ2_Nan_RS16675     tacacgctccaggtgcagggcgtcgtcgacaagacgtccggctggggcggccgcatcatg
                      ************************************************************

MxDZ2_Kirby_RS0232470 gcggagaacgtggagctgcgtccgtccgaagcgcgcgccgtgtctcgcgacgagatggag
MxDZ2_Tam_RS26190     gcggagaacgtggagctgcgtccgtccgaagcgcgcgccgtgtctcgcgacgagatggag
MxDZ2_Nan_RS16675     gcggagaacgtggagctgcgtccgtccgaagcgcgcgccgtgtctcgcgacgagatggag
                      ************************************************************

MxDZ2_Kirby_RS0232470 ctggtgcacatcgagggagaggtccccacgtccgtggacgtgcgcctcaaccacggcctc
MxDZ2_Tam_RS26190     ctggtgcacatcgagggagaggtccccacgtccgtggacgtgcgcctcaaccacggcctc
MxDZ2_Nan_RS16675     ctggtgcacatcgagggagaggtccccacgtccgtggacgtgcgcctcaaccacggcctc
                      ************************************************************

MxDZ2_Kirby_RS0232470 accgtgcgcctgcccgagcaccccggcttcacctgggccatcgagcccacggtggccaag
MxDZ2_Tam_RS26190     accgtgcgcctgcccgagcaccccggcttcacctgggccatcgagcccacggtggccaag
MxDZ2_Nan_RS16675     accgtgcgcctgcccgagcaccccggcttcacctgggccatcgagcccacggtggccaag
                      ************************************************************

MxDZ2_Kirby_RS0232470 cgcgtgggcctgcgcgaagccaacttcgagcccgcgcccgaggacggtcctggtacccgc
MxDZ2_Tam_RS26190     cgcgtgggcctgcgcgaagccaacttcgagcccgcgcccgaggacggtcctggtacccgc
MxDZ2_Nan_RS16675     cgcgtgggcctgcgcgaagccaacttcgagcccgcgcccgaggacggtcctggtacccgc
                      ************************************************************

MxDZ2_Kirby_RS0232470 gagttcttcttcaccccgcgcaacccgggcgccttcgatgtggagttcttcctggccaag
MxDZ2_Tam_RS26190     gagttcttcttcaccccgcgcaacccgggcgccttcgatgtggagttcttcctggccaag
MxDZ2_Nan_RS16675     gagttcttcttcaccccgcgcaacccgggcgccttcgatgtggagttcttcctggccaag
                      ************************************************************

MxDZ2_Kirby_RS0232470 gcgctgtcgcccggcctggtggaccgctccttcaaaatcaacgtcacggtcaagccctga
MxDZ2_Tam_RS26190     gcgctgtcgcccggcctggtggaccgctccttcaaaatcaacgtcacggtcaagccctga
MxDZ2_Nan_RS16675     gcgctgtcgcccggcctggtggaccgctccttcaaaatcaacgtcacggtcaagccctga
                      ************************************************************

-----------------------------------------------------------------------------------
multiple sequence alignment (protein)

MxDZ2_Kirby_RS0232470 MAKPKSGAKKTTPAGKAGAKPAAKKDSAARLDLIKNASKRAAKAATKLAKAASDVTKGAK
MxDZ2_Tam_RS26190     MAKPKSGAKKTTPAGKAGAKPAAKKDSAARLDLIKNASKRAAKAATKLAKAASDVTKGAK
MxDZ2_Nan_RS16675     MAKPKSGAKKTTPAGKAGAKPAAKKDSAARLDLIKNASKRAAKAATKLAKAASDVTKGAK
                      ************************************************************

MxDZ2_Kirby_RS0232470 SAVKKAVQEKATAKKAAPAGKTASAAKAAP--------------PAAKTPKAP-------
MxDZ2_Tam_RS26190     SAVKKAVQEKATAKKAAPAGKTASAAKAAP--------------PAAKTPKAP-------
MxDZ2_Nan_RS16675     SAVKKAVQEKATAKKAAPAGKTASAAKAAPRQRRPRRHLRRRRLPRRSPPRAPRRRRRAR
                      ******************************              *  ..*:**

MxDZ2_Kirby_RS0232470 ---------PAKAPPAAKSAASAKAAKASAVPAAPPVEKPRPRATKLPPPGEPLTKREME
MxDZ2_Tam_RS26190     ---------PAKAPPAAKSAASAKAAKASAVPAAPPVEKPRPRATKLPPPGEPLTKREME
MxDZ2_Nan_RS16675     YPPPLRWRSPVLAPPSCRLRASRSPSVKWSSCRPAKAAASRARAASRAAWWSPMTCRTWW
                               *. ***:.:  ** ..:   :   .. .  .*.**:. ..  .*:* *

MxDZ2_Kirby_RS0232470 QLLTAGEGRGVTGEGSLKGRLVVTNDMPHLVVVGRDKRELTFLLQGPDQEVLPAYVDHKV
MxDZ2_Tam_RS26190     QLLTAGEGRGVTGEGSLKGRLVVTNDMPHLVVVGRDKRELTFLLQGPDQEVLPAYVDHKV
MxDZ2_Nan_RS16675     SSAATS-----ASPSSSRARI-------------RKSSRPTWTTRSP-------------
                      .  ::.     :. .* :.*:             *.. . *:  :.*

MxDZ2_Kirby_RS0232470 SVSGLIRKTTNHA--------------------------GVVDVRKYSA--KKP---DAE
MxDZ2_Tam_RS26190     SVSGLIRKTTNHA--------------------------GVVDVRKYSA--KKP---DAE
MxDZ2_Nan_RS16675     SAGSSVRRPTTLAWWTFASTRPRSRMPRWWRPPRRRRRRGCATCRPVRSPWRRPAWARAS
                      *... :*:.*. *                          * .  *   :  ::*    *.

MxDZ2_Kirby_RS0232470 VVEAAPAETEARLRYLSPGEVSMVTAAGMGAGIKGFAGVRGNLEMTGEEFVLVVSNGGTR
MxDZ2_Tam_RS26190     VVEAAPAETEARLRYLSPGEVSMVTAAGMGAGIKGFAGVRGNLEMTGEEFVLVVSNGGTR
MxDZ2_Nan_RS16675     RASRACVATWRRAR-------SSCSSCPMAA-------------------------RASR
                       .. * . *  * *       *  ::. *.*                          .:*

MxDZ2_Kirby_RS0232470 QQVSFIIEGKAAGKALRKHVGYTLQVQGVVDKTSGWGGRIMAENVELRPSEARAVSRDEM
MxDZ2_Tam_RS26190     QQVSFIIEGKAAGKALRKHVGYTLQVQGVVDKTSGWGGRIMAENVELRPSEARAVSRDEM
MxDZ2_Nan_RS16675     YRSS------SRARPLARPCA----------STSGTRSRCRASSTR-RPAGAAASWRRTW
                       : *      : .:.* :  .          .***  .*  *.... **: * *  *

MxDZ2_Kirby_RS0232470 ELVHIEGEVPTSVDVRLNHGLTVRLP------------EHPGFTWAIEPTVAKRVGLREA
MxDZ2_Tam_RS26190     ELVHIEGEVPTSVDVRLNHGLTVRLP------------EHPGFTWAIEPTVAKRVGLREA
MxDZ2_Nan_RS16675     SCVRPK-----------------RAPCLATRWSWCTSRERSPRPWTCASTTASPCACPST
                      . *: :                 * *            *:.  .*:  .*.*.  .  .:

MxDZ2_Kirby_RS0232470 NFEPAPEDGPGTREFFFTPRNPGAFDVEFFLAKALS---------------------PGL
MxDZ2_Tam_RS26190     NFEPAPEDGPGTREFFFTPRNPGAFDVEFFLAKALS---------------------PGL
MxDZ2_Nan_RS16675     PASPGPS----------SPRWPSAWAC----AKPTSSPRPRTVLVPASSSSPRATRAPSM
                        .*.*.          :** *.*:      **. *                     *.:

MxDZ2_Kirby_RS0232470 VDRSF-------------KINVTVKP
MxDZ2_Tam_RS26190     VDRSF-------------KINVTVKP
MxDZ2_Nan_RS16675     WSSSWPRRCRPAWWTAPSKSTSRSSP
                       . *:             * .   .*
```

**Frameshift mutation 7**

| **Tam_location** | **Tam_nucleotide** | **Kirby_nucleotide** | **Nan_nucleotide** | **Nan_location** |
| --- | --- | --- | --- | --- |
| **5389814** | **G (complement)** | **G (complement)** | **.** | **5389790** |

```
multiple sequence alignment (nucleotide)

MxDZ2_Kirby_RS0232530 atgacggactcgggtaatctgatgcaggccgccgccgaggtggcgcggatagcgggtgac
MxDZ2_Tam_RS26250     atgacggactcgggtaatctgatgcaggccgccgccgaggtggcgcggatagcgggtgac
MxDZ2_Nan_RS16615     atgacggactcgggtaatctgatgcaggccgccgccgaggtggcgcggatagcgggtgac
                      ************************************************************

MxDZ2_Kirby_RS0232530 gcggcgttggggttcttccgcggtggcatcgcggtggacacgaagtcggacggctctccg
MxDZ2_Tam_RS26250     gcggcgttggggttcttccgcggtggcatcgcggtggacacgaagtcggacggctctccg
MxDZ2_Nan_RS16615     gcggcgttggggttcttccgcggtggcatcgcggtggacacgaagtcggacggctctccg
                      ************************************************************

MxDZ2_Kirby_RS0232530 gtgacggtggcggaccgcacggcggagtcccgtgcgcgcgagtggctggaagcgcgcttc
MxDZ2_Tam_RS26250     gtgacggtggcggaccgcacggcggagtcccgtgcgcgcgagtggctggaagcgcgcttc
MxDZ2_Nan_RS16615     gtgacggtggcggaccgcacggcggagtcccgtgcgcgcgagtggctggaagcgcgcttc
                      ************************************************************

MxDZ2_Kirby_RS0232530 ccccaggacggcatcctgggcgaggagttcggcgagacacgtccgggcgcgaagcgccgg
MxDZ2_Tam_RS26250     ccccaggacggcatcctgggcgaggagttcggcgagacacgtccgggcgcgaagcgccgg
MxDZ2_Nan_RS16615     ccccaggacggcatcctgggcgaggagttcggcgagacacgtccgggcgcgaagcgccgg
                      ************************************************************

MxDZ2_Kirby_RS0232530 tggattctggatcccattgacgggacgaagacgttcatccgcggtgtcccactgtggggc
MxDZ2_Tam_RS26250     tggattctggatcccattgacgggacgaagacgttcatccgcggtgtcccactgtggggc
MxDZ2_Nan_RS16615     tggattctggatcccattgacgggacgaagacgttcatccgcggtgtcccactgtggggc
                      ************************************************************

MxDZ2_Kirby_RS0232530 acgttggtggcgctggcggagggcgagcgcatcctcgtgggggccgcgtacttCcccgcg
MxDZ2_Tam_RS26250     acgttggtggcgctggcggagggcgagcgcatcctcgtgggggccgcgtacttCcccgcg
MxDZ2_Nan_RS16615     acgttggtggcgctggcggagggcgagcgcatcctcgtgggggccgcgtactt-cccgcg
                      ***************************************************** ******

MxDZ2_Kirby_RS0232530 gtgagtgagttgctggtggccgcgccggggcggggctgcttctggaacgaccagcgcgcg
MxDZ2_Tam_RS26250     gtgagtgagttgctggtggccgcgccggggcggggctgcttctggaacgaccagcgcgcg
MxDZ2_Nan_RS16615     gtgagtgagttgctggtggccgcgccggggcggggctgcttctggaacgaccagcgcgcg
                      ************************************************************

MxDZ2_Kirby_RS0232530 gcggtgtccacgcaggcggagctgtcccaggccgtggtgctgtccacggatgagcgcttc
MxDZ2_Tam_RS26250     gcggtgtccacgcaggcggagctgtcccaggccgtggtgctgtccacggatgagcgcttc
MxDZ2_Nan_RS16615     gcggtgtccacgcaggcggagctgtcccaggccgtggtgctgtccacggatgagcgcttc
                      ************************************************************

MxDZ2_Kirby_RS0232530 ccggtgtacccggagcgcggggccgcctggcgctcgctcgcgcgggatgcggccgtggac
MxDZ2_Tam_RS26250     ccggtgtacccggagcgcggggccgcctggcgctcgctcgcgcgggatgcggccgtggac
MxDZ2_Nan_RS16615     ccggtgtacccggagcgcggggccgcctggcgctcgctcgcgcgggatgcggccgtggac
                      ************************************************************

MxDZ2_Kirby_RS0232530 cgcacctggggggattgctacggctacctgttggtcgccaccgggcgcgcggaggtcatg
MxDZ2_Tam_RS26250     cgcacctggggggattgctacggctacctgttggtcgccaccgggcgcgcggaggtcatg
MxDZ2_Nan_RS16615     cgcacctggggggattgctacggctacctgttggtcgccaccgggcgcgcggaggtcatg
                      ************************************************************

MxDZ2_Kirby_RS0232530 gtggatgagctgctgtccccctgggatggagcggccttgcagcccatcatcgaggaggcc
MxDZ2_Tam_RS26250     gtggatgagctgctgtccccctgggatggagcggccttgcagcccatcatcgaggaggcc
MxDZ2_Nan_RS16615     gtggatgagctgctgtccccctgggatggagcggccttgcagcccatcatcgaggaggcc
                      ************************************************************

MxDZ2_Kirby_RS0232530 ggcggtgtgttcaccgactggacggggcggcggaccgcgttcggcggaaatggaatcgcc
MxDZ2_Tam_RS26250     ggcggtgtgttcaccgactggacggggcggcggaccgcgttcggcggaaatggaatcgcc
MxDZ2_Nan_RS16615     ggcggtgtgttcaccgactggacggggcggcggaccgcgttcggcggaaatggaatcgcc
                      ************************************************************

MxDZ2_Kirby_RS0232530 accaacgcggccatggcgcgcgtggtgcgggagcggctcggcgccgtggagacacgctga
MxDZ2_Tam_RS26250     accaacgcggccatggcgcgcgtggtgcgggagcggctcggcgccgtggagacacgctga
MxDZ2_Nan_RS16615     accaacgcggccatggcgcgcgtggtgcgggagcggctcggcgccgtggagacacgctga
                      ************************************************************

-----------------------------------------------------------------------------------


multiple sequence alignment (protein)

MxDZ2_Kirby_RS0232530 MTDSGNLMQAAAEVARIAGDAALGFFRGGIAVDTKSDGSPVTVADRTAESRAREWLEARF
MxDZ2_Tam_RS26250     MTDSGNLMQAAAEVARIAGDAALGFFRGGIAVDTKSDGSPVTVADRTAESRAREWLEARF
MxDZ2_Nan_RS16615     MTDSGNLMQAAAEVARIAGDAALGFFRGGIAVDTKSDGSPVTVADRTAESRAREWLEARF
                      ************************************************************

MxDZ2_Kirby_RS0232530 PQDGILGEEFGETRPGAKRRWILDPIDGTKTFIRGVPLWGTLVALAEGERILVGAAYFPA
MxDZ2_Tam_RS26250     PQDGILGEEFGETRPGAKRRWILDPIDGTKTFIRGVPLWGTLVALAEGERILVGAAYFPA
MxDZ2_Nan_RS16615     PQDGILGEEFGETRPGAKRRWILDPIDGTKTFIRGVPLWGTLVALAEGERILVGAAYFPR
                      ***********************************************************

MxDZ2_Kirby_RS0232530 VSELLVAAPGRGCFWNDQRAAVSTQAELSQAVVLSTDERFPVYPERGAAWRSLARDAAVD
MxDZ2_Tam_RS26250     VSELLVAAPGRGCFWNDQRAAVSTQAELSQAVVLSTDERFPVYPERGAAWRSLARDAAVD
MxDZ2_Nan_RS16615     VS----------CWWPRRGGAASGTTSARRCPRRRSCPRPWCCPRMSASRCTRSAGPPGA
                      **          *:*  : .*.*  :.  :.    :  *    *. .*:  : : ...

MxDZ2_Kirby_RS0232530 RTWGDCYGYLLVATGRAEVMVDELLSPWD---GAALQPIIEEAGGVFTDW----------
MxDZ2_Tam_RS26250     RTWGDCYGYLLVATGRAEVMVDELLSPWD---GAALQPIIEEAGGVFTDW----------
MxDZ2_Nan_RS16615     RSRG--------------------MRPWTAPGGIATATCWSPPGARRSWWMSCCPPGMER
                      *: *                    : **    * *  .  . .*.  : *

MxDZ2_Kirby_RS0232530 ----TGRRTAF----GGNGIATNAAM----ARVVRERLGAVET----R
MxDZ2_Tam_RS26250     ----TGRRTAF----GGNGIATNAAM----ARVVRERLGAVET----R
MxDZ2_Nan_RS16615     PCSPSSRRPAVCSPTGRGGGPRSAEMESPPTRPWRAWCGSGSAPWRHA
                          :.**.*.    * .* . .* *    :*  *   *: .:
```

**Frameshift mutation 8**

| **Tam_location** | **Tam_nucleotide** | **Kirby_nucleotide** | **Nan_nucleotide** | **Nan_location** |
| --- | --- | --- | --- | --- |
| **5495633** | **G (complement)** | **G (complement)** | **.** | **5495607** |

```
multiple sequence alignment (nucleotide)

MxDZ2_Kirby_RS0227330 atgaaccgcaaggacggattcatggcggagttgtcgccactccagcaggcgttgctgacc
MxDZ2_Tam_RS26610     atgaaccgcaaggacggattcatggcggagttgtcgccactccagcaggcgttgctgacc
MxDZ2_Nan_RS16250     atgaaccgcaaggacggattcatggcggagttgtcgccactccagcaggcgttgctgacc
                      ************************************************************

MxDZ2_Kirby_RS0227330 atcgagaagctgcagaagaagctggccacggcctcgaatggtgagcgggaaccgattgcc
MxDZ2_Tam_RS26610     atcgagaagctgcagaagaagctggccacggcctcgaatggtgagcgggaaccgattgcc
MxDZ2_Nan_RS16250     atcgagaagctgcagaagaagctggccacggcctcgaatggtgagcgggaaccgattgcc
                      ************************************************************

MxDZ2_Kirby_RS0227330 atcatcggcatggcctgccgctttccggggggcgccaccagccccgcgaagttccgtgac
MxDZ2_Tam_RS26610     atcatcggcatggcctgccgctttccggggggcgccaccagccccgcgaagttccgtgac
MxDZ2_Nan_RS16250     atcatcggcatggcctgccgctttccggggggcgccaccagccccgcgaagttccgtgac
                      ************************************************************

MxDZ2_Kirby_RS0227330 ctgctgtggggggcacgggaggcgctgacggagattcccagcgaccggcaggcgctccag
MxDZ2_Tam_RS26610     ctgctgtggggggcacgggaggcgctgacggagattcccagcgaccggcaggcgctccag
MxDZ2_Nan_RS16250     ctgctgtggggggcacgggaggcgctgacggagattcccagcgaccggcaggcgctccag
                      ************************************************************

MxDZ2_Kirby_RS0227330 gggctctatgacccggacccctccaggcccgggaagctggccatgcgccgcgccggcttc
MxDZ2_Tam_RS26610     gggctctatgacccggacccctccaggcccgggaagctggccatgcgccgcgccggcttc
MxDZ2_Nan_RS16250     gggctctatgacccggacccctccaggcccgggaagctggccatgcgccgcgccggcttc
                      ************************************************************

MxDZ2_Kirby_RS0227330 gtggacgaggtggaccggttcgacgcggagttcttccacatctcccggcgtgaagccgag
MxDZ2_Tam_RS26610     gtggacgaggtggaccggttcgacgcggagttcttccacatctcccggcgtgaagccgag
MxDZ2_Nan_RS16250     gtggacgaggtggaccggttcgacgcggagttcttccacatctcccggcgtgaagccgag
                      ************************************************************

MxDZ2_Kirby_RS0227330 ggcatggacccgcagcagcgcttcttcctggaggtgagctgggaggccctggaggacgct
MxDZ2_Tam_RS26610     ggcatggacccgcagcagcgcttcttcctggaggtgagctgggaggccctggaggacgct
MxDZ2_Nan_RS16250     ggcatggacccgcagcagcgcttcttcctggaggtgagctgggaggccctggaggacgct
                      ************************************************************
MxDZ2_Kirby_RS0227330 ggcattcccccacaccagctccaggggacgcggaccggtgtcttcgcgggcgtccacgcg
MxDZ2_Tam_RS26610     ggcattcccccacaccagctccaggggacgcggaccggtgtcttcgcgggcgtccacgcg
MxDZ2_Nan_RS16250     ggcattcccccacaccagctccaggggacgcggaccggtgtcttcgcgggcgtccacgcg
                      ************************************************************

MxDZ2_Kirby_RS0227330 aaggactacgcgttcgtcacggggggcgggctggagaaggtgagcgcccactactccacg
MxDZ2_Tam_RS26610     aaggactacgcgttcgtcacggggggcgggctggagaaggtgagcgcccactactccacg
MxDZ2_Nan_RS16250     aaggactacgcgttcgtcacggggggcgggctggagaaggtgagcgcccactactccacg
                      ************************************************************

MxDZ2_Kirby_RS0227330 ggtgtggacgccagctatgtggcgggccggctgtcgtacctcctcgggctcgaagggccg
MxDZ2_Tam_RS26610     ggtgtggacgccagctatgtggcgggccggctgtcgtacctcctcgggctcgaagggccg
MxDZ2_Nan_RS16250     ggtgtggacgccagctatgtggcgggccggctgtcgtacctcctcgggctcgaagggccg
                      ************************************************************

MxDZ2_Kirby_RS0227330 agcatggcggtggacaccgcgtgctcgtcctctttgtccgctgtgcacctggcgtgtcag
MxDZ2_Tam_RS26610     agcatggcggtggacaccgcgtgctcgtcctctttgtccgctgtgcacctggcgtgtcag
MxDZ2_Nan_RS16250     agcatggcggtggacaccgcgtgctcgtcctctttgtccgctgtgcacctggcgtgtcag
                      ************************************************************

MxDZ2_Kirby_RS0227330 agcctgcggacagaagagtcgacgctcgccatcgccggaggcgtgaagctcatcctggcg
MxDZ2_Tam_RS26610     agcctgcggacagaagagtcgacgctcgccatcgccggaggcgtgaagctcatcctggcg
MxDZ2_Nan_RS16250     agcctgcggacagaagagtcgacgctcgccatcgccggaggcgtgaagctcatcctggcg
                      ************************************************************

MxDZ2_Kirby_RS0227330 ccgcaactcagcgtgttcctgtccaaggcgggagccctctcccccagcggccattgccgc
MxDZ2_Tam_RS26610     ccgcaactcagcgtgttcctgtccaaggcgggagccctctcccccagcggccattgccgc
MxDZ2_Nan_RS16250     ccgcaactcagcgtgttcctgtccaaggcgggagccctctcccccagcggccattgccgc
                      ************************************************************

MxDZ2_Kirby_RS0227330 acgttcgaccgggacgcggacggcatggtgcagggggagggctgtggcgtggtcgtcctg
MxDZ2_Tam_RS26610     acgttcgaccgggacgcggacggcatggtgcagggggagggctgtggcgtggtcgtcctg
MxDZ2_Nan_RS16250     acgttcgaccgggacgcggacggcatggtgcagggggagggctgtggcgtggtcgtcctg
                      ************************************************************

MxDZ2_Kirby_RS0227330 aagcggctgcgggacgccgttcgggatggtgaccgcatcctcgcgaccttgcgtggcacg
MxDZ2_Tam_RS26610     aagcggctgcgggacgccgttcgggatggtgaccgcatcctcgcgaccttgcgtggcacg
MxDZ2_Nan_RS16250     aagcggctgcgggacgccgttcgggatggtgaccgcatcctcgcgaccttgcgtggcacg
                      ************************************************************


MxDZ2_Kirby_RS0227330 ggcatgaaccacgacggcgccagcggtggtctcacggtgccgaacgtcggggcacaggaa
MxDZ2_Tam_RS26610     ggcatgaaccacgacggcgccagcggtggtctcacggtgccgaacgtcggggcacaggaa
MxDZ2_Nan_RS16250     ggcatgaaccacgacggcgccagcggtggtctcacggtgccgaacgtcggggcacaggaa
                      ************************************************************

MxDZ2_Kirby_RS0227330 gcgctgtaccggcgcgtgttgcagcgcgcgggcatcgagcccgggcaggtggactacctg
MxDZ2_Tam_RS26610     gcgctgtaccggcgcgtgttgcagcgcgcgggcatcgagcccgggcaggtggactacctg
MxDZ2_Nan_RS16250     gcgctgtaccggcgcgtgttgcagcgcgcgggcatcgagcccgggcaggtggactacctg
                      ************************************************************

MxDZ2_Kirby_RS0227330 gaagcgcacgggacgggaacgcggctgggagatcccatcgagctggacggcgtgtcgcgc
MxDZ2_Tam_RS26610     gaagcgcacgggacgggaacgcggctgggagatcccatcgagctggacggcgtgtcgcgc
MxDZ2_Nan_RS16250     gaagcgcacgggacgggaacgcggctgggagatcccatcgagctggacggcgtgtcgcgc
                      ************************************************************

MxDZ2_Kirby_RS0227330 gtctacggcgcagcgcggacttcggagcgcccgctgtggattggctccgtcaagccgaac
MxDZ2_Tam_RS26610     gtctacggcgcagcgcggacttcggagcgcccgctgtggattggctccgtcaagccgaac
MxDZ2_Nan_RS16250     gtctacggcgcagcgcggacttcggagcgcccgctgtggattggctccgtcaagccgaac
                      ************************************************************

MxDZ2_Kirby_RS0227330 atcggtcacacggaggccgcggccggcatcgcgggactcatcaaggccgtcctggtgctc
MxDZ2_Tam_RS26610     atcggtcacacggaggccgcggccggcatcgcgggactcatcaaggccgtcctggtgctc
MxDZ2_Nan_RS16250     atcggtcacacggaggccgcggccggcatcgcgggactcatcaaggccgtcctggtgctc
                      ************************************************************

MxDZ2_Kirby_RS0227330 caggcgggcgaggtgcctccctccatcaacttcgagcacccgaccccggagttcgcgtgg
MxDZ2_Tam_RS26610     caggcgggcgaggtgcctccctccatcaacttcgagcacccgaccccggagttcgcgtgg
MxDZ2_Nan_RS16250     caggcgggcgaggtgcctccctccatcaacttcgagcacccgaccccggagttcgcgtgg
                      ************************************************************

MxDZ2_Kirby_RS0227330 gagggctcagggctcgcggtgcctcgagcgcgcacggccctggcggagaaggacggccct
MxDZ2_Tam_RS26610     gagggctcagggctcgcggtgcctcgagcgcgcacggccctggcggagaaggacggccct
MxDZ2_Nan_RS16250     gagggctcagggctcgcggtgcctcgagcgcgcacggccctggcggagaaggacggccct
                      ************************************************************

MxDZ2_Kirby_RS0227330 catcgcgtcgccgtgagctcgtttgggatgagcggtgtgaacgcgcatgcgctcgtcgag
MxDZ2_Tam_RS26610     catcgcgtcgccgtgagctcgtttgggatgagcggtgtgaacgcgcatgcgctcgtcgag
MxDZ2_Nan_RS16250     catcgcgtcgccgtgagctcgtttgggatgagcggtgtgaacgcgcatgcgctcgtcgag
                      ************************************************************


MxDZ2_Kirby_RS0227330 gcgtacgcggaggccgcccatgccgccgtggacgccgggccgtatgtgttgcccctctcc
MxDZ2_Tam_RS26610     gcgtacgcggaggccgcccatgccgccgtggacgccgggccgtatgtgttgcccctctcc
MxDZ2_Nan_RS16250     gcgtacgcggaggccgcccatgccgccgtggacgccgggccgtatgtgttgcccctctcc
                      ************************************************************

MxDZ2_Kirby_RS0227330 gcgcgctcggagcccgcgctgcgcacactggcggcgtcatggctccagtacgtgtcggag
MxDZ2_Tam_RS26610     gcgcgctcggagcccgcgctgcgcacactggcggcgtcatggctccagtacgtgtcggag
MxDZ2_Nan_RS16250     gcgcgctcggagcccgcgctgcgcacactggcggcgtcatggctccagtacgtgtcggag
                      ************************************************************

MxDZ2_Kirby_RS0227330 gccggtcctggggcgcggctccaggatgcgtgcttcatggcgggcgctggacgctcacac
MxDZ2_Tam_RS26610     gccggtcctggggcgcggctccaggatgcgtgcttcatggcgggcgctggacgctcacac
MxDZ2_Nan_RS16250     gccggtcctggggcgcggctccaggatgcgtgcttcatggcgggcgctggacgctcacac
                      ************************************************************

MxDZ2_Kirby_RS0227330 catgcgcaccgtgcggtgctcgtggcgaggacgcccgaggcgcttcgagcggcccttcac
MxDZ2_Tam_RS26610     catgcgcaccgtgcggtgctcgtggcgaggacgcccgaggcgcttcgagcggcccttcac
MxDZ2_Nan_RS16250     catgcgcaccgtgcggtgctcgtggcgaggacgcccgaggcgcttcgagcggcccttcac
                      ************************************************************

MxDZ2_Kirby_RS0227330 gctgtcacggagggacgcgatgcgccgggtgtcaatcgaggttcaggcgagggcgagccc
MxDZ2_Tam_RS26610     gctgtcacggagggacgcgatgcgccgggtgtcaatcgaggttcaggcgagggcgagccc
MxDZ2_Nan_RS16250     gctgtcacggagggacgcgatgcgccgggtgtcaatcgaggttcaggcgagggcgagccc
                      ************************************************************

MxDZ2_Kirby_RS0227330 gtcttcctcttcggtggcacggagcccgaggccagccggttgatgcgtgagctgctgggc
MxDZ2_Tam_RS26610     gtcttcctcttcggtggcacggagcccgaggccagccggttgatgcgtgagctgctgggc
MxDZ2_Nan_RS16250     gtcttcctcttcggtggcacggagcccgaggccagccggttgatgcgtgagctgctgggc
                      ************************************************************

MxDZ2_Kirby_RS0227330 cgggacgtcttccgcgcccacgcggagaaggtggatgcggtcttccggcaggtggcaggt
MxDZ2_Tam_RS26610     cgggacgtcttccgcgcccacgcggagaaggtggatgcggtcttccggcaggtggcaggt
MxDZ2_Nan_RS16250     cgggacgtcttccgcgcccacgcggagaaggtggatgcggtcttccggcaggtggcaggt
                      ************************************************************

MxDZ2_Kirby_RS0227330 gggtccgtcatcgaacaggtggcgggagcgcaggcgggcgaaggacacccggtgggcact
MxDZ2_Tam_RS26610     gggtccgtcatcgaacaggtggcgggagcgcaggcgggcgaaggacacccggtgggcact
MxDZ2_Nan_RS16250     gggtccgtcatcgaacaggtggcgggagcgcaggcgggcgaaggacacccggtgggcact
                      ************************************************************


MxDZ2_Kirby_RS0227330 ctcctcgtgctccagctcgcattggcggagctctggcgcgcgtgcggtctgacccccgca
MxDZ2_Tam_RS26610     ctcctcgtgctccagctcgcattggcggagctctggcgcgcgtgcggtctgacccccgca
MxDZ2_Nan_RS16250     ctcctcgtgctccagctcgcattggcggagctctggcgcgcgtgcggtctgacccccgca
                      ************************************************************

MxDZ2_Kirby_RS0227330 gctgtttccggcttcggcgtgggagcgctctccgcgggagtggtcgcgggagcactcagc
MxDZ2_Tam_RS26610     gctgtttccggcttcggcgtgggagcgctctccgcgggagtggtcgcgggagcactcagc
MxDZ2_Nan_RS16250     gctgtttccggcttcggcgtgggagcgctctccgcgggagtggtcgcgggagcactcagc
                      ************************************************************

MxDZ2_Kirby_RS0227330 gtggaagacgccgtgcggctcgccttgcagttgacgccagcgcagcccgtgcgtcctcaa
MxDZ2_Tam_RS26610     gtggaagacgccgtgcggctcgccttgcagttgacgccagcgcagcccgtgcgtcctcaa
MxDZ2_Nan_RS16250     gtggaagacgccgtgcggctcgccttgcagttgacgccagcgcagcccgtgcgtcctcaa
                      ************************************************************

MxDZ2_Kirby_RS0227330 cccgtgaagtacccattcttctccaccgtcgatggcgcctgggtcgaagcgggcgccacg
MxDZ2_Tam_RS26610     cccgtgaagtacccattcttctccaccgtcgatggcgcctgggtcgaagcgggcgccacg
MxDZ2_Nan_RS16250     cccgtgaagtacccattcttctccaccgtcgatggcgcctgggtcgaagcgggcgccacg
                      ************************************************************

MxDZ2_Kirby_RS0227330 gtaccgggcgcgtactgggagagacagcgcgccgcggtgatcgacccggcgtcgtccgtg
MxDZ2_Tam_RS26610     gtaccgggcgcgtactgggagagacagcgcgccgcggtgatcgacccggcgtcgtccgtg
MxDZ2_Nan_RS16250     gtaccgggcgcgtactgggagagacagcgcgccgcggtgatcgacccggcgtcgtccgtg
                      ************************************************************

MxDZ2_Kirby_RS0227330 gcgaacctgtgtgcgcggaacgcctcggctttcatccaggtctcttgtgaggcttccgtt
MxDZ2_Tam_RS26610     gcgaacctgtgtgcgcggaacgcctcggctttcatccaggtctcttgtgaggcttccgtt
MxDZ2_Nan_RS16250     gcgaacctgtgtgcgcggaacgcctcggctttcatccaggtctcttgtgaggcttccgtt
                      ************************************************************

MxDZ2_Kirby_RS0227330 ggggcggcgttggaggccactgcacgcagccgtggaagcaaggccgtcatcctgcgtgcc
MxDZ2_Tam_RS26610     ggggcggcgttggaggccactgcacgcagccgtggaagcaaggccgtcatcctgcgtgcc
MxDZ2_Nan_RS16250     ggggcggcgttggaggccactgcacgcagccgtggaagcaaggccgtcatcctgcgtgcc
                      ************************************************************

MxDZ2_Kirby_RS0227330 agcaggcccggagatgatgcctggacgggtgtcctgggcgtcatcgcgggtgggtactcc
MxDZ2_Tam_RS26610     agcaggcccggagatgatgcctggacgggtgtcctgggcgtcatcgcgggtgggtactcc
MxDZ2_Nan_RS16250     agcaggcccggagatgatgcctggacgggtgtcctgggcgtcatcgcgggtgggtactcc
                      ************************************************************


MxDZ2_Kirby_RS0227330 tcgggtggctccattcgtttcgaggggctcttcacgggggcgcgcaagacggtggtgccg
MxDZ2_Tam_RS26610     tcgggtggctccattcgtttcgaggggctcttcacgggggcgcgcaagacggtggtgccg
MxDZ2_Nan_RS16250     tcgggtggctccattcgtttcgaggggctcttcacgggggcgcgcaagacggtggtgccg
                      ************************************************************

MxDZ2_Kirby_RS0227330 acgtacccgtggcagcgtgagcgctactggtgggacggagcggtcgctccagcaccgagc
MxDZ2_Tam_RS26610     acgtacccgtggcagcgtgagcgctactggtgggacggagcggtcgctccagcaccgagc
MxDZ2_Nan_RS16250     acgtacccgtggcagcgtgagcgctactggtgggacggagcggtcgctccagcaccgagc
                      ************************************************************

MxDZ2_Kirby_RS0227330 agcgtgcagcccgtccgggatgcatCcccccgggacctcttctgcgaactcacgtggcat
MxDZ2_Tam_RS26610     agcgtgcagcccgtccgggatgcatCcccccgggacctcttctgcgaactcacgtggcat
MxDZ2_Nan_RS16250     agcgtgcagcccgtccgggatgcat-cccccgggacctcttctgcgaactcacgtggcat
                      ************************* **********************************

MxDZ2_Kirby_RS0227330 gcgcgtacggtggggcaggggcccgtggacgcggagggccgctggctcatcgtgagcacg
MxDZ2_Tam_RS26610     gcgcgtacggtggggcaggggcccgtggacgcggagggccgctggctcatcgtgagcacg
MxDZ2_Nan_RS16250     gcgcgtacggtggggcaggggcccgtggacgcggagggccgctggctcatcgtgagcacg
                      ************************************************************

MxDZ2_Kirby_RS0227330 cagccggctgcctcggagtcgctgcggcaatccctgtccaccgccggaggcacggtgcgc
MxDZ2_Tam_RS26610     cagccggctgcctcggagtcgctgcggcaatccctgtccaccgccggaggcacggtgcgc
MxDZ2_Nan_RS16250     cagccggctgcctcggagtcgctgcggcaatccctgtccaccgccggaggcacggtgcgc
                      ************************************************************

MxDZ2_Kirby_RS0227330 gtggccatcgtgggctccgagcaggcgtcggtgctccgggagtggctgtccgagagcccg
MxDZ2_Tam_RS26610     gtggccatcgtgggctccgagcaggcgtcggtgctccgggagtggctgtccgagagcccg
MxDZ2_Nan_RS16250     gtggccatcgtgggctccgagcaggcgtcggtgctccgggagtggctgtccgagagcccg
                      ************************************************************

MxDZ2_Kirby_RS0227330 gcgccccgaggcgtcatctacctgtcgggttcggagcatcccgaatcactggaagggctg
MxDZ2_Tam_RS26610     gcgccccgaggcgtcatctacctgtcgggttcggagcatcccgaatcactggaagggctg
MxDZ2_Nan_RS16250     gcgccccgaggcgtcatctacctgtcgggttcggagcatcccgaatcactggaagggctg
                      ************************************************************

MxDZ2_Kirby_RS0227330 cacacatcggttcagcgagaggtccacgcggcggcgtcgctcgtgcagaccctggcgcgg
MxDZ2_Tam_RS26610     cacacatcggttcagcgagaggtccacgcggcggcgtcgctcgtgcagaccctggcgcgg
MxDZ2_Nan_RS16250     cacacatcggttcagcgagaggtccacgcggcggcgtcgctcgtgcagaccctggcgcgg
                      ************************************************************


MxDZ2_Kirby_RS0227330 cactcggcgccaacgctgcctcggctgttcctcgtcacccagggctcgcaagccgctgac
MxDZ2_Tam_RS26610     cactcggcgccaacgctgcctcggctgttcctcgtcacccagggctcgcaagccgctgac
MxDZ2_Nan_RS16250     cactcggcgccaacgctgcctcggctgttcctcgtcacccagggctcgcaagccgctgac
                      ************************************************************

MxDZ2_Kirby_RS0227330 ggagcggctggacccacggcgattgctggcgcgccactgtggggcctgggacgggttgtc
MxDZ2_Tam_RS26610     ggagcggctggacccacggcgattgctggcgcgccactgtggggcctgggacgggttgtc
MxDZ2_Nan_RS16250     ggagcggctggacccacggcgattgctggcgcgccactgtggggcctgggacgggttgtc
                      ************************************************************

MxDZ2_Kirby_RS0227330 gcctacgagcacccggagcttgcctgcaagcgcatcgacatcgaacccgcggccttcggt
MxDZ2_Tam_RS26610     gcctacgagcacccggagcttgcctgcaagcgcatcgacatcgaacccgcggccttcggt
MxDZ2_Nan_RS16250     gcctacgagcacccggagcttgcctgcaagcgcatcgacatcgaacccgcggccttcggt
                      ************************************************************

MxDZ2_Kirby_RS0227330 gcgctggtgacggagctggcgcgcaaggtctctgcttccgctgacgacgacgaggtcgtg
MxDZ2_Tam_RS26610     gcgctggtgacggagctggcgcgcaaggtctctgcttccgctgacgacgacgaggtcgtg
MxDZ2_Nan_RS16250     gcgctggtgacggagctggcgcgcaaggtctctgcttccgctgacgacgacgaggtcgtg
                      ************************************************************

MxDZ2_Kirby_RS0227330 ctgcgcggcgagcagcggctggtgccctcgctcacgcagacccaaacgattccggaaggc
MxDZ2_Tam_RS26610     ctgcgcggcgagcagcggctggtgccctcgctcacgcagacccaaacgattccggaaggc
MxDZ2_Nan_RS16250     ctgcgcggcgagcagcggctggtgccctcgctcacgcagacccaaacgattccggaaggc
                      ************************************************************

MxDZ2_Kirby_RS0227330 acgggagcgtttcgtccccggcaagaccgcacgtatctgattacgggcgggctcggtgga
MxDZ2_Tam_RS26610     acgggagcgtttcgtccccggcaagaccgcacgtatctgattacgggcgggctcggtgga
MxDZ2_Nan_RS16250     acgggagcgtttcgtccccggcaagaccgcacgtatctgattacgggcgggctcggtgga
                      ************************************************************

MxDZ2_Kirby_RS0227330 atcggactgcggctcgcctcctggctggtggagcgcggcgcgcggcaccttgccctctgt
MxDZ2_Tam_RS26610     atcggactgcggctcgcctcctggctggtggagcgcggcgcgcggcaccttgccctctgt
MxDZ2_Nan_RS16250     atcggactgcggctcgcctcctggctggtggagcgcggcgcgcggcaccttgccctctgt
                      ************************************************************

MxDZ2_Kirby_RS0227330 ggaagaaaaggtgagaccgacgaggcccgccaggccctggcgccgctgcgtgcggcgggc
MxDZ2_Tam_RS26610     ggaagaaaaggtgagaccgacgaggcccgccaggccctggcgccgctgcgtgcggcgggc
MxDZ2_Nan_RS16250     ggaagaaaaggtgagaccgacgaggcccgccaggccctggcgccgctgcgtgcggcgggc
                      ************************************************************


MxDZ2_Kirby_RS0227330 gcgcgggtggagaccttccgcgtcgatgtgagccgcccggagtcggtggcggagatgctg
MxDZ2_Tam_RS26610     gcgcgggtggagaccttccgcgtcgatgtgagccgcccggagtcggtggcggagatgctg
MxDZ2_Nan_RS16250     gcgcgggtggagaccttccgcgtcgatgtgagccgcccggagtcggtggcggagatgctg
                      ************************************************************

MxDZ2_Kirby_RS0227330 gctgccgtccgccgaggcggagcgccgctggggggcatcttccacagcgcgggtgtgctg
MxDZ2_Tam_RS26610     gctgccgtccgccgaggcggagcgccgctggggggcatcttccacagcgcgggtgtgctg
MxDZ2_Nan_RS16250     gctgccgtccgccgaggcggagcgccgctggggggcatcttccacagcgcgggtgtgctg
                      ************************************************************

MxDZ2_Kirby_RS0227330 gcggacagctcgttgttgcagtgggagccccagggcttcgagacggtcatgggccccaag
MxDZ2_Tam_RS26610     gcggacagctcgttgttgcagtgggagccccagggcttcgagacggtcatgggccccaag
MxDZ2_Nan_RS16250     gcggacagctcgttgttgcagtgggagccccagggcttcgagacggtcatgggccccaag
                      ************************************************************

MxDZ2_Kirby_RS0227330 gtcgacggtgcgtggaacctccatgcgctggccaccgacaccaccctcgagcacttcgtc
MxDZ2_Tam_RS26610     gtcgacggtgcgtggaacctccatgcgctggccaccgacaccaccctcgagcacttcgtc
MxDZ2_Nan_RS16250     gtcgacggtgcgtggaacctccatgcgctggccaccgacaccaccctcgagcacttcgtc
                      ************************************************************

MxDZ2_Kirby_RS0227330 ctgttctcctcgacggcgtcgctgattggctcgccggggcaggccagctatgccgcggcc
MxDZ2_Tam_RS26610     ctgttctcctcgacggcgtcgctgattggctcgccggggcaggccagctatgccgcggcc
MxDZ2_Nan_RS16250     ctgttctcctcgacggcgtcgctgattggctcgccggggcaggccagctatgccgcggcc
                      ************************************************************

MxDZ2_Kirby_RS0227330 aatgcattcctcgatgcgttggcgcacttccgtcgcgcacggggtctgccggccatcagt
MxDZ2_Tam_RS26610     aatgcattcctcgatgcgttggcgcacttccgtcgcgcacggggtctgccggccatcagt
MxDZ2_Nan_RS16250     aatgcattcctcgatgcgttggcgcacttccgtcgcgcacggggtctgccggccatcagt
                      ************************************************************

MxDZ2_Kirby_RS0227330 ctcaactgggggagctggggcgaggtcggcatggccgtcgcggatgctcggcgtggcgac
MxDZ2_Tam_RS26610     ctcaactgggggagctggggcgaggtcggcatggccgtcgcggatgctcggcgtggcgac
MxDZ2_Nan_RS16250     ctcaactgggggagctggggcgaggtcggcatggccgtcgcggatgctcggcgtggcgac
                      ************************************************************

MxDZ2_Kirby_RS0227330 cggctggcggagcggggcatgtcgcccatgtccgtggatgaagccctggcggccatggcg
MxDZ2_Tam_RS26610     cggctggcggagcggggcatgtcgcccatgtccgtggatgaagccctggcggccatggcg
MxDZ2_Nan_RS16250     cggctggcggagcggggcatgtcgcccatgtccgtggatgaagccctggcggccatggcg
                      ************************************************************

MxDZ2_Kirby_RS0227330 ctggtgctcgcggagaaccgcgtcacccggggcatcgcgcgcttcgatgtgacgcgttgg
MxDZ2_Tam_RS26610     ctggtgctcgcggagaaccgcgtcacccggggcatcgcgcgcttcgatgtgacgcgttgg
MxDZ2_Nan_RS16250     ctggtgctcgcggagaaccgcgtcacccggggcatcgcgcgcttcgatgtgacgcgttgg
                      ************************************************************

MxDZ2_Kirby_RS0227330 actgccagccatccttccatcgtggcttcgtcgctgttccggaccgagtccgtcaccgat
MxDZ2_Tam_RS26610     actgccagccatccttccatcgtggcttcgtcgctgttccggaccgagtccgtcaccgat
MxDZ2_Nan_RS16250     actgccagccatccttccatcgtggcttcgtcgctgttccggaccgagtccgtcaccgat
                      ************************************************************

MxDZ2_Kirby_RS0227330 gccgtccctgacgtcatggaggcgccgcgcaagctgcgcgaggtgttgctcgcgttggaa
MxDZ2_Tam_RS26610     gccgtccctgacgtcatggaggcgccgcgcaagctgcgcgaggtgttgctcgcgttggaa
MxDZ2_Nan_RS16250     gccgtccctgacgtcatggaggcgccgcgcaagctgcgcgaggtgttgctcgcgttggaa
                      ************************************************************

MxDZ2_Kirby_RS0227330 ggctccgctcgccgcggcgtcctggaggcgcggctcaaggagatgatcagcgaggttgga
MxDZ2_Tam_RS26610     ggctccgctcgccgcggcgtcctggaggcgcggctcaaggagatgatcagcgaggttgga
MxDZ2_Nan_RS16250     ggctccgctcgccgcggcgtcctggaggcgcggctcaaggagatgatcagcgaggttgga
                      ************************************************************

MxDZ2_Kirby_RS0227330 aagattccgacgacgaagctctcgtccacggactcgttcgacgcgctggggttcgactcc
MxDZ2_Tam_RS26610     aagattccgacgacgaagctctcgtccacggactcgttcgacgcgctggggttcgactcc
MxDZ2_Nan_RS16250     aagattccgacgacgaagctctcgtccacggactcgttcgacgcgctggggttcgactcc
                      ************************************************************

MxDZ2_Kirby_RS0227330 atcatggcgctcgagttgcgggacatggtgctgggcgaactcgacgtgagcatgcctctc
MxDZ2_Tam_RS26610     atcatggcgctcgagttgcgggacatggtgctgggcgaactcgacgtgagcatgcctctc
MxDZ2_Nan_RS16250     atcatggcgctcgagttgcgggacatggtgctgggcgaactcgacgtgagcatgcctctc
                      ************************************************************

MxDZ2_Kirby_RS0227330 aagagtttcgttgatgaaagttccattgagcaggtgacggctgagcttctcggaaagctc
MxDZ2_Tam_RS26610     aagagtttcgttgatgaaagttccattgagcaggtgacggctgagcttctcggaaagctc
MxDZ2_Nan_RS16250     aagagtttcgttgatgaaagttccattgagcaggtgacggctgagcttctcggaaagctc
                      ************************************************************

MxDZ2_Kirby_RS0227330 gctgtggccagtgtgctcggcgctgcctcctcggagcgcgatgattcgaagcgggtcctg
MxDZ2_Tam_RS26610     gctgtggccagtgtgctcggcgctgcctcctcggagcgcgatgattcgaagcgggtcctg
MxDZ2_Nan_RS16250     gctgtggccagtgtgctcggcgctgcctcctcggagcgcgatgattcgaagcgggtcctg
                      ************************************************************

MxDZ2_Kirby_RS0227330 ctatga
MxDZ2_Tam_RS26610     ctatga
MxDZ2_Nan_RS16250     ctatga
                      ******

-----------------------------------------------------------------------------------

multiple sequence alignment (protein)

MxDZ2_Kirby_RS0227330 MNRKDGFMAELSPLQQALLTIEKLQKKLATASNGEREPIAIIGMACRFPGGATSPAKFRD
MxDZ2_Tam_RS26610     MNRKDGFMAELSPLQQALLTIEKLQKKLATASNGEREPIAIIGMACRFPGGATSPAKFRD
MxDZ2_Nan_RS16250     MNRKDGFMAELSPLQQALLTIEKLQKKLATASNGEREPIAIIGMACRFPGGATSPAKFRD
                      ************************************************************

MxDZ2_Kirby_RS0227330 LLWGAREALTEIPSDRQALQGLYDPDPSRPGKLAMRRAGFVDEVDRFDAEFFHISRREAE
MxDZ2_Tam_RS26610     LLWGAREALTEIPSDRQALQGLYDPDPSRPGKLAMRRAGFVDEVDRFDAEFFHISRREAE
MxDZ2_Nan_RS16250     LLWGAREALTEIPSDRQALQGLYDPDPSRPGKLAMRRAGFVDEVDRFDAEFFHISRREAE
                      ************************************************************

MxDZ2_Kirby_RS0227330 GMDPQQRFFLEVSWEALEDAGIPPHQLQGTRTGVFAGVHAKDYAFVTGGGLEKVSAHYST
MxDZ2_Tam_RS26610     GMDPQQRFFLEVSWEALEDAGIPPHQLQGTRTGVFAGVHAKDYAFVTGGGLEKVSAHYST
MxDZ2_Nan_RS16250     GMDPQQRFFLEVSWEALEDAGIPPHQLQGTRTGVFAGVHAKDYAFVTGGGLEKVSAHYST
                      ************************************************************

MxDZ2_Kirby_RS0227330 GVDASYVAGRLSYLLGLEGPSMAVDTACSSSLSAVHLACQSLRTEESTLAIAGGVKLILA
MxDZ2_Tam_RS26610     GVDASYVAGRLSYLLGLEGPSMAVDTACSSSLSAVHLACQSLRTEESTLAIAGGVKLILA
MxDZ2_Nan_RS16250     GVDASYVAGRLSYLLGLEGPSMAVDTACSSSLSAVHLACQSLRTEESTLAIAGGVKLILA
                      ************************************************************

MxDZ2_Kirby_RS0227330 PQLSVFLSKAGALSPSGHCRTFDRDADGMVQGEGCGVVVLKRLRDAVRDGDRILATLRGT
MxDZ2_Tam_RS26610     PQLSVFLSKAGALSPSGHCRTFDRDADGMVQGEGCGVVVLKRLRDAVRDGDRILATLRGT
MxDZ2_Nan_RS16250     PQLSVFLSKAGALSPSGHCRTFDRDADGMVQGEGCGVVVLKRLRDAVRDGDRILATLRGT
                      ************************************************************

MxDZ2_Kirby_RS0227330 GMNHDGASGGLTVPNVGAQEALYRRVLQRAGIEPGQVDYLEAHGTGTRLGDPIELDGVSR
MxDZ2_Tam_RS26610     GMNHDGASGGLTVPNVGAQEALYRRVLQRAGIEPGQVDYLEAHGTGTRLGDPIELDGVSR
MxDZ2_Nan_RS16250     GMNHDGASGGLTVPNVGAQEALYRRVLQRAGIEPGQVDYLEAHGTGTRLGDPIELDGVSR
                      ************************************************************


MxDZ2_Kirby_RS0227330 VYGAARTSERPLWIGSVKPNIGHTEAAAGIAGLIKAVLVLQAGEVPPSINFEHPTPEFAW
MxDZ2_Tam_RS26610     VYGAARTSERPLWIGSVKPNIGHTEAAAGIAGLIKAVLVLQAGEVPPSINFEHPTPEFAW
MxDZ2_Nan_RS16250     VYGAARTSERPLWIGSVKPNIGHTEAAAGIAGLIKAVLVLQAGEVPPSINFEHPTPEFAW
                      ************************************************************

MxDZ2_Kirby_RS0227330 EGSGLAVPRARTALAEKDGPHRVAVSSFGMSGVNAHALVEAYAEAAHAAVDAGPYVLPLS
MxDZ2_Tam_RS26610     EGSGLAVPRARTALAEKDGPHRVAVSSFGMSGVNAHALVEAYAEAAHAAVDAGPYVLPLS
MxDZ2_Nan_RS16250     EGSGLAVPRARTALAEKDGPHRVAVSSFGMSGVNAHALVEAYAEAAHAAVDAGPYVLPLS
                      ************************************************************

MxDZ2_Kirby_RS0227330 ARSEPALRTLAASWLQYVSEAGPGARLQDACFMAGAGRSHHAHRAVLVARTPEALRAALH
MxDZ2_Tam_RS26610     ARSEPALRTLAASWLQYVSEAGPGARLQDACFMAGAGRSHHAHRAVLVARTPEALRAALH
MxDZ2_Nan_RS16250     ARSEPALRTLAASWLQYVSEAGPGARLQDACFMAGAGRSHHAHRAVLVARTPEALRAALH
                      ************************************************************

MxDZ2_Kirby_RS0227330 AVTEGRDAPGVNRGSGEGEPVFLFGGTEPEASRLMRELLGRDVFRAHAEKVDAVFRQVAG
MxDZ2_Tam_RS26610     AVTEGRDAPGVNRGSGEGEPVFLFGGTEPEASRLMRELLGRDVFRAHAEKVDAVFRQVAG
MxDZ2_Nan_RS16250     AVTEGRDAPGVNRGSGEGEPVFLFGGTEPEASRLMRELLGRDVFRAHAEKVDAVFRQVAG
                      ************************************************************

MxDZ2_Kirby_RS0227330 GSVIEQVAGAQAGEGHPVGTLLVLQLALAELWRACGLTPAAVSGFGVGALSAGVVAGALS
MxDZ2_Tam_RS26610     GSVIEQVAGAQAGEGHPVGTLLVLQLALAELWRACGLTPAAVSGFGVGALSAGVVAGALS
MxDZ2_Nan_RS16250     GSVIEQVAGAQAGEGHPVGTLLVLQLALAELWRACGLTPAAVSGFGVGALSAGVVAGALS
                      ************************************************************

MxDZ2_Kirby_RS0227330 VEDAVRLALQLTPAQPVRPQPVKYPFFSTVDGAWVEAGATVPGAYWERQRAAVIDPASSV
MxDZ2_Tam_RS26610     VEDAVRLALQLTPAQPVRPQPVKYPFFSTVDGAWVEAGATVPGAYWERQRAAVIDPASSV
MxDZ2_Nan_RS16250     VEDAVRLALQLTPAQPVRPQPVKYPFFSTVDGAWVEAGATVPGAYWERQRAAVIDPASSV
                      ************************************************************

MxDZ2_Kirby_RS0227330 ANLCARNASAFIQVSCEASVGAALEATARSRGSKAVILRASRPGDDAWTGVLGVIAGGYS
MxDZ2_Tam_RS26610     ANLCARNASAFIQVSCEASVGAALEATARSRGSKAVILRASRPGDDAWTGVLGVIAGGYS
MxDZ2_Nan_RS16250     ANLCARNASAFIQVSCEASVGAALEATARSRGSKAVILRASRPGDDAWTGVLGVIAGGYS
                      ************************************************************

MxDZ2_Kirby_RS0227330 SGGSIRFEGLFTGARKTVVPTYPWQRERYWWDGAVAPAPSSVQPVRDASP----------
MxDZ2_Tam_RS26610     SGGSIRFEGLFTGARKTVVPTYPWQRERYWWDGAVAPAPSSVQPVRDASP----------
MxDZ2_Nan_RS16250     SGGSIRFEGLFTGARKTVVPTYPWQRERYWWDGAVAPAPSSVQPVRDASPGTSSANSRGM
                      **************************************************


MxDZ2_Kirby_RS0227330 RDLFCELTWHARTVGQG---------------PVDAEGRW--------------LIVSTQ
MxDZ2_Tam_RS26610     RDLFCELTWHARTVGQG---------------PVDAEGRW--------------LIVSTQ
MxDZ2_Nan_RS16250     RVRWGRGPWTRRAAGSSARSRLPRSRCGNPCPPPEARCAWPSWAPSRRRCSGSGCPRARR
                      *  : . .*  *:.*..               * :*.  *                 : :

MxDZ2_Kirby_RS0227330 PAASESL-----------------------RQSLSTAGGTVRVAIVGSEQASVLREWLSE
MxDZ2_Tam_RS26610     PAASESL-----------------------RQSLSTAGGTVRVAIVGSEQASVLREWLSE
MxDZ2_Nan_RS16250     PEASSTCRVRSIPNHWKGCTHRFSERSTRRRRSCRPWRGTRRQRCLGCSSSPRARKPLTE
                      * **.:                        *:*  .  ** *   :*...:.  *: *:*

MxDZ2_Kirby_RS0227330 SPAPRGVIYLSGSEHPESLEGLHTSVQREVHAAASL------VQTLARHSAPTLPRLFLV
MxDZ2_Tam_RS26610     SPAPRGVIYLSGSEHPESLEGLHTSVQREVHAAASL------VQTLARHSAPTLPRLFLV
MxDZ2_Nan_RS16250     RLDPRRLL----ARHCGAWDGLSPTSTRSLPASASTSNPRPSVRWRSWRARSLLP--LTT
                         ** ::    :.*  : :** .:  *.: *:**       *:  : :: . **  : .

MxDZ2_Kirby_RS0227330 TQGSQAADGAAGPTA-------------IAGAPLWGLGRVVAYEHPELACKRIDIEPAAF
MxDZ2_Tam_RS26610     TQGSQAADGAAGPTA-------------IAGAPLWGLGRVVAYEHPELACKRIDIEPAAF
MxDZ2_Nan_RS16250     TRSCCAASSGWCPRSRRPKRFRKARERFVPGKT----ARILRAGSVESDCGSPPGWWSAA
                      *:.. **...  * :             :.* .    .*::     *  *       :*

MxDZ2_Kirby_RS0227330 GALVTELARKVSASADDDEVVLRGEQRLVPSLTQTQTIPEGTGAFRPRQDRTYLITGGLG
MxDZ2_Tam_RS26610     GALVTELARKVSASADDDEVVLRGEQRLVPSLTQTQTIPEGTGAFRPRQDRTYLITGGLG
MxDZ2_Nan_RS16250     RGTLPSVEEKVRPTRP-----ARPWRRCVRRARGWR--PSASMAARSRWRRCWLPSAEAE
                       . :..: .** .:        *  :* *      :  *..: * *.*  * :* :.

MxDZ2_Kirby_RS0227330 GIGLRLASWLVERGARHLALCGR--KGETDEARQALAPLRAAGARVETFRVDVSRPESVA
MxDZ2_Tam_RS26610     GIGLRLASWLVERGARHLALCGR--KGETDEARQALAPLRAAGARVETFRVDVSRPESVA
MxDZ2_Nan_RS16250     ------RRWGASSTARVCWRTARCCSGSPRASRRSWAP--RSTVRGTSMRWPPTPPSSTS
                              * ..  **     .*  .*..  :*:: **   : .*  ::*   : *.*.:

MxDZ2_Kirby_RS0227330 EMLAA----VRRGGAPLGGIFHSAGVLADSSLLQWEPQGFETVMGPKVD-------GAWN
MxDZ2_Tam_RS26610     EMLAA----VRRGGAPLGGIFHSAGVLADSSLLQWEPQGFETVMGPKVD-------GAWN
MxDZ2_Nan_RS16250     SCSPRRRRLARRGRPAMPRPMHSS--------MRWRTSVAHGVCRPSVSTGGAGARSAWP
                      .  .     .*** ..:   :**:        ::*...  . *  *.*.       .**

MxDZ2_Kirby_RS0227330 LHALATDTT-----------LEHFVLFSSTASLIGSPGQASYAAANAFLDAL--AHFRRA
MxDZ2_Tam_RS26610     LHALATDTT-----------LEHFVLFSSTASLIGSPGQASYAAANAFLDAL--AHFRRA
MxDZ2_Nan_RS16250     SRMLGVATGWRSGACRPCPWMKPWRPWRWCSRRTASPG-ASRASMRVGLPAILPSWLRRC
                       : *.. *            :: :  :   :   .*** ** *: .. * *:  : :**.


MxDZ2_Kirby_RS0227330 RG-LPAISLNWGSWG------------EVGMAVADARRGDRLAERGMSPMSVDEALAAMA
MxDZ2_Tam_RS26610     RG-LPAISLNWGSWG------------EVGMAVADARRGDRLAERGMSPMSVDEALAAMA
MxDZ2_Nan_RS16250     SGPSPSPMPSLTSWRRRASCARCCSRWKAPLAAASWRRGSRRSAR---------------
                       *  *:   .  **             :. :*.*. ***.* : *

MxDZ2_Kirby_RS0227330 LVLAENRVTRGIARFDVTRW--TASHPSIVASSLFRTESVTDAVPDVMEAPRKLREVLLA
MxDZ2_Tam_RS26610     LVLAENRVTRGIARFDVTRW--TASHPSIVASSLFRTESVTDAVPDVMEAPRKLREVLLA
MxDZ2_Nan_RS16250     --LERFRRRSSRPRTRSTRWGSTPSWRSSCGTWCWANSTACLSRVSLMKVPLSRR-----
                        * . *   . .*   ***  *.*  *  .:  : ..:.  :  .:*:.* . *

MxDZ2_Kirby_RS0227330 LEGSARRGVLEARLKEMISEVGKIPTTKLSSTDSFDALGFDSIMALELRDMVLGELDVSM
MxDZ2_Tam_RS26610     LEGSARRGVLEARLKEMISEVGKIPTTKLSSTDSFDALGFDSIMALELRDMVLGELDVSM
MxDZ2_Nan_RS16250     -------------------------------------LSF--------------------
                                                           *.*

MxDZ2_Kirby_RS0227330 PLKSFVDESSIEQVTAELLGKLAVASVLGAASSERDDSKRVLL
MxDZ2_Tam_RS26610     PLKSFVDESSIEQVTAELLGKLAVASVLGAASSERDDSKRVLL
MxDZ2_Nan_RS16250     ------SESSLWPVCSALPPRSAMI---------RSGS---CY
                            .***:  * : *  : *:          *..*
```

**Premature stop codon followed by second ORF prediction 1**

| **Tam_location** | **Tam_nucleotide** | **Kirby_nucleotide** | **Nan_nucleotide** | **Nan_location** |
| --- | --- | --- | --- | --- |
| **2102329** | **C (complement)** | **C (complement)** | **.** | **2102322** |

```
multiple sequence alignment (nucleotide)

MxDZ2_Kirby_RS0202735 gtggcgagcctcctcaccctcctgggcgcgggagtcgaggtcgcgtactacgcgctcgac
MxDZ2_Tam_RS13120     gtggcgagcctcctcaccctcctgggcgcgggagtcgaggtcgcgtactacgcgctcgac
MxDZ2_Nan_RS29755     gtggcgagcctcctcaccctcctgggcgcgggagtcgaggtcgcgtactacgcgctcgac
                      ************************************************************

MxDZ2_Kirby_RS0202735 gcccgggggctggagccattcgtcgcgttcctctcgttgagctacgagcggaacctgccc
MxDZ2_Tam_RS13120     gcccgggggctggagccattcgtcgcgttcctctcgttgagctacgagcggaacctgccc
MxDZ2_Nan_RS29755     gcccgggggctggagccattcgtcgcgttcctctcgttgagctacgagcggaacctgccc
                      ************************************************************

MxDZ2_Kirby_RS0202735 acctggtacgcgtcggctttgctgctgagctgcgcgctcgtgctcgcgtgtatcggccat
MxDZ2_Tam_RS13120     acctggtacgcgtcggctttgctgctgagctgcgcgctcgtgctcgcgtgtatcggccat
MxDZ2_Nan_RS29755     acctggtacgcgtcggctttgctgctgagctgcgcgctcgtgctcgcgtgtatcggccat
                      ************************************************************

MxDZ2_Kirby_RS0202735 atggcgggacagggccgggagcagggatggcgtcattggcgggcgctggcgggcgcgttc
MxDZ2_Tam_RS13120     atggcgggacagggccgggagcagggatggcgtcattggcgggcgctggcgggcgcgttc
MxDZ2_Nan_RS29755     atggcgggacagggccgggagcagggatggcgtcattggcgggcgctggcgggcgcgttc
                      ************************************************************

MxDZ2_Kirby_RS0202735 gcgtacatctctctcgacgagagcgtgggcctccacgagcacctcggggggcagctctcc
MxDZ2_Tam_RS13120     gcgtacatctctctcgacgagagcgtgggcctccacgagcacctcggggggcagctctcc
MxDZ2_Nan_RS29755     gcgtacatctctctcgacgagagcgtgggcctccacgagcacctcggggggcagctctcc
                      ************************************************************

MxDZ2_Kirby_RS0202735 ctgggtggcgtgctcttcttcacctgggtagtgcccGgggctgtcctcgtgctgctcggc
MxDZ2_Tam_RS13120     ctgggtggcgtgctcttcttcacctgggtagtgcccGgggctgtcctcgtgctgctcggc
MxDZ2_Nan_RS29755     ctgggtggcgtgctcttcttcacctgggtagtgccc-gggctgtcctcgtgctgctcggc
                      ************************************ ***********************

MxDZ2_Kirby_RS0202735 gggcttgccttcctccccttcctcgccgggctccgcccgcgccgacgccgacagttcatc
MxDZ2_Tam_RS13120     gggcttgccttcctccccttcctcgccgggctccgcccgcgccgacgccgacagttcatc
MxDZ2_Nan_RS29755     gggcttgccttcctccccttcctcgccgggctccgcccgcgccgacgccgacagttcatc
                      ************************************************************

MxDZ2_Kirby_RS0202735 ctcgctggggtgctgtacgtgggtggggcgctggtgatggagcttccgctggggtggtgg
MxDZ2_Tam_RS13120     ctcgctggggtgctgtacgtgggtggggcgctggtgatggagcttccgctggggtggtgg
MxDZ2_Nan_RS29755     ctcgctggggtgctgtacgtgggtggggcgctggtga-----------------------
                      *************************************

MxDZ2_Kirby_RS0202735 gcggagcaacacggcaacgacaacctcgtgtacgcactcatcgaccacgtggaagaggcc
MxDZ2_Tam_RS13120     gcggagcaacacggcaacgacaacctcgtgtacgcactcatcgaccacgtggaagaggcc
MxDZ2_Nan_RS29755     ------------------------------------------------------------


MxDZ2_Kirby_RS0202735 ctggaactgattggtgccagcctgttcctcgcggcgctggcagaggaactgggcgagcgg
MxDZ2_Tam_RS13120     ctggaactgattggtgccagcctgttcctcgcggcgctggcagaggaactgggcgagcgg
MxDZ2_Nan_RS29755     ------------------------------------------------------------


MxDZ2_Kirby_RS0202735 gtggtgctcacggtacgaaaggaggagccccatgggcctggcccagccgctccccgctga
MxDZ2_Tam_RS13120     gtggtgctcacggtacgaaaggaggagccccatgggcctggcccagccgctccccgctga
MxDZ2_Nan_RS29755     ------------------------------------------------------------

-----------------------------------------------------------------------------------
multiple sequence alignment (protein)


MxDZ2_Kirby_RS0202735 MASLLTLLGAGVEVAYYALDARGLEPFVAFLSLSYERNLPTWYASALLLSCALVLACIGH
MxDZ2_Tam_RS13120     MASLLTLLGAGVEVAYYALDARGLEPFVAFLSLSYERNLPTWYASALLLSCALVLACIGH
MxDZ2_Nan_RS29755     MASLLTLLGAGVEVAYYALDARGLEPFVAFLSLSYERNLPTWYASALLLSCALVLACIGH
                      ************************************************************

MxDZ2_Kirby_RS0202735 MAGQGREQGWRHWRALAGAFAYISLDESVGLHEHLGGQLSLGGVLFFTWVVPGAVLVLLG
MxDZ2_Tam_RS13120     MAGQGREQGWRHWRALAGAFAYISLDESVGLHEHLGGQLSLGGVLFFTWVVPGAVLVLLG
MxDZ2_Nan_RS29755     MAGQGREQGWRHWRALAGAFAYISLDESVGLHEHLGGQLSLGGVLFFTWVVPGLSSCCSA
                      *****************************************************      .

MxDZ2_Kirby_RS0202735 GLAFLPFLAG-LRPRRRRQFILAGVLYVGGALVMELPLGWWAEQHGNDNLVYALIDHVEE
MxDZ2_Tam_RS13120     GLAFLPFLAG-LRPRRRRQFILAGVLYVGGALVMELPLGWWAEQHGNDNLVYALIDHVEE
MxDZ2_Nan_RS29755     GLPSSPSSPGSARADADSSSSLGCCTWVGR----------W-------------------
                      **.  *  .*  *.    .  *.   :**           *

MxDZ2_Kirby_RS0202735 ALELIGASLFLAALAEELGERVVLTVRKEEPHGPGPAAPR
MxDZ2_Tam_RS13120     ALELIGASLFLAALAEELGERVVLTVRKEEPHGPGPAAPR
MxDZ2_Nan_RS29755     ----------------------------------------
```

**Premature stop codon followed by second ORF prediction 2**

| **Tam_location** | **Tam_nucleotide** | **Kirby_nucleotide** | **Nan_nucleotide** | **Nan_location** |
| --- | --- | --- | --- | --- |
| **2225127** | **G (complement)** | **G (complement)** | **.** | **2225117** |

```
multiple sequence alignment (nucleotide)

MxDZ2_Kirby_RS0235795 atgatggattcgtacgagcagatggacgatatcgacctgcggcccctaacacgcgaaatg
MxDZ2_Tam_RS13780     atgatggattcgtacgagcagatggacgatatcgacctgcggcccctaacacgcgaaatg
MxDZ2_Nan_RS29095     atgatggattcgtacgagcagatggacgatatcgacctgcggcccctaacacgcgaaatg
                      ************************************************************

MxDZ2_Kirby_RS0235795 gaccttgagaatctcggcctgagccaggcccggacggtaacgggcgtgcacgtctatctc
MxDZ2_Tam_RS13780     gaccttgagaatctcggcctgagccaggcccggacggtaacgggcgtgcacgtctatctc
MxDZ2_Nan_RS29095     gaccttgagaatctcggcctgagccaggcccggacggtaacgggcgtgcacgtctatctc
                      ************************************************************

MxDZ2_Kirby_RS0235795 gacgtcaccaactttcaaagcttgctcgaggcgcgacgcgagaataccgccgaactaaag
MxDZ2_Tam_RS13780     gacgtcaccaactttcaaagcttgctcgaggcgcgacgcgagaataccgccgaactaaag
MxDZ2_Nan_RS29095     gacgtcaccaactttcaaagcttgctcgaggcgcgacgcgagaataccgccgaactaaag
                      ************************************************************

MxDZ2_Kirby_RS0235795 aagcttcttcgccacctcaccgtctttcagcggcaggtcgagcgattgcttaggcagcag
MxDZ2_Tam_RS13780     aagcttcttcgccacctcaccgtctttcagcggcaggtcgagcgattgcttaggcagcag
MxDZ2_Nan_RS29095     aagcttcttcgccacctcaccgtctttcagcggcaggtcgagcgattgcttaggcagcag
                      ************************************************************

MxDZ2_Kirby_RS0235795 gacggcgccgccgtccgggtgcactttcagggtgcgcggctccacctcatcgtcttcaag
MxDZ2_Tam_RS13780     gacggcgccgccgtccgggtgcactttcagggtgcgcggctccacctcatcgtcttcaag
MxDZ2_Nan_RS29095     gacggcgccgccgtccgggtgcactttcagggtgcgcggctccacctcatcgtcttcaag
                      ************************************************************

MxDZ2_Kirby_RS0235795 ccgtacgacaccgaggaaggggcaactctcgctcgcttggtgaaggccgtcacgctggtc
MxDZ2_Tam_RS13780     ccgtacgacaccgaggaaggggcaactctcgctcgcttggtgaaggccgtcacgctggtc
MxDZ2_Nan_RS29095     ccgtacgacaccgaggaaggggcaactctcgctcgcttggtgaaggccgtcacgctggtc
                      ************************************************************

MxDZ2_Kirby_RS0235795 gagcaggtgcgccaactggcggacatgattcagttccacacggacatccacttcgagttt
MxDZ2_Tam_RS13780     gagcaggtgcgccaactggcggacatgattcagttccacacggacatccacttcgagttt
MxDZ2_Nan_RS29095     gagcaggtgcgccaactggcggacatgattcagttccacacggacatccacttcgagttt
                      ************************************************************

MxDZ2_Kirby_RS0235795 gaggccggggtagacactggcgaggcaatcgccacgatgaacgcgcgggctagacgtcgc
MxDZ2_Tam_RS13780     gaggccggggtagacactggcgaggcaatcgccacgatgaacgcgcgggctagacgtcgc
MxDZ2_Nan_RS29095     gaggccggggtagacactggcgaggcaatcgccacgatgaacgcgcgggctagacgtcgc
                      ************************************************************

MxDZ2_Kirby_RS0235795 cagttgctgtttgttggccgcccggcgaatgaagccgcaaaggcgctgacgggcaacccc
MxDZ2_Tam_RS13780     cagttgctgtttgttggccgcccggcgaatgaagccgcaaaggcgctgacgggcaacccc
MxDZ2_Nan_RS29095     cagttgctgtttgttggccgcccggcgaatgaagccgcaaaggcgctgacgggcaacccc
                      ************************************************************

MxDZ2_Kirby_RS0235795 agtgCccccggagaagagcggctcgccgccggagcggcccccttcacgaagcaactcccc
MxDZ2_Tam_RS13780     agtgCccccggagaagagcggctcgccgccggagcggcccccttcacgaagcaactcccc
MxDZ2_Nan_RS29095     agtg-ccccggagaagagcggctcgccgccggagcggcccccttcacgaagcaactcccc
                      **** *******************************************************

MxDZ2_Kirby_RS0235795 tccgctgctgctcgagagaaactaccgaccagcttcgaggccgccgtcaaggaggacatt
MxDZ2_Tam_RS13780     tccgctgctgctcgagagaaactaccgaccagcttcgaggccgccgtcaaggaggacatt
MxDZ2_Nan_RS29095     tccgctgctgctcgagagaaactaccgaccagcttcgaggccgccgtcaaggaggacatt
                      ************************************************************

MxDZ2_Kirby_RS0235795 gaggcccatccgttggaaagcctggatatcaccgacgttcgagcacctattgactacggc
MxDZ2_Tam_RS13780     gaggcccatccgttggaaagcctggatatcaccgacgttcgagcacctattgactacggc
MxDZ2_Nan_RS29095     gaggcccatccgttggaaagcctggatatcaccgacgttcgagcacctattgactacggc
                      ************************************************************

MxDZ2_Kirby_RS0235795 tcactgggcaagaaggtagcgcggctagaggaggccataacattcttcggggacctatcc
MxDZ2_Tam_RS13780     tcactgggcaagaaggtagcgcggctagaggaggccataacattcttcggggacctatcc
MxDZ2_Nan_RS29095     tcactgggcaagaaggtag-----------------------------------------
                      *******************

MxDZ2_Kirby_RS0235795 ggcttcacgagctatgtcgccagcctctctaccgaagatgaaaagaaggaggcgctgcga
MxDZ2_Tam_RS13780     ggcttcacgagctatgtcgccagcctctctaccgaagatgaaaagaaggaggcgctgcga
MxDZ2_Nan_RS29095     ------------------------------------------------------------


MxDZ2_Kirby_RS0235795 ctccttcatgtcctgcgctcggagatgcaggccgtcgttgctgactacgacggcgacttc
MxDZ2_Tam_RS13780     ctccttcatgtcctgcgctcggagatgcaggccgtcgttgctgactacgacggcgacttc
MxDZ2_Nan_RS29095     ------------------------------------------------------------


MxDZ2_Kirby_RS0235795 atccagttccagggcgaccgagtccaagccctcctgtacgaggcgcgcacttctgggcgg
MxDZ2_Tam_RS13780     atccagttccagggcgaccgagtccaagccctcctgtacgaggcgcgcacttctgggcgg
MxDZ2_Nan_RS29095     ------------------------------------------------------------


MxDZ2_Kirby_RS0235795 ttcaagagcaaagccgtggagacggccgccgccctatgctccgccgtcgaccttgccaag
MxDZ2_Tam_RS13780     ttcaagagcaaagccgtggagacggccgccgccctatgctccgccgtcgaccttgccaag
MxDZ2_Nan_RS29095     ------------------------------------------------------------


MxDZ2_Kirby_RS0235795 gaactcttcccagcgctgtcgacgctggacgtttcctgcggggcggatgttgggaaggtt
MxDZ2_Tam_RS13780     gaactcttcccagcgctgtcgacgctggacgtttcctgcggggcggatgttgggaaggtt
MxDZ2_Nan_RS29095     ------------------------------------------------------------


MxDZ2_Kirby_RS0235795 cttgttggccgtgtggggttgcgcgatgacaaggaggttactgtcctcggccgctccatc
MxDZ2_Tam_RS13780     cttgttggccgtgtggggttgcgcgatgacaaggaggttactgtcctcggccgctccatc
MxDZ2_Nan_RS29095     ------------------------------------------------------------


MxDZ2_Kirby_RS0235795 ccatgtgcctccgagttacaggacggcgctggtgacggccacaccgccatcagcgtacgt
MxDZ2_Tam_RS13780     ccatgtgcctccgagttacaggacggcgctggtgacggccacaccgccatcagcgtacgt
MxDZ2_Nan_RS29095     ------------------------------------------------------------


MxDZ2_Kirby_RS0235795 ttgtgggagggcatcgagaaatcgcttcagggcttcttctcgctgaagaaggagcgctac
MxDZ2_Tam_RS13780     ttgtgggagggcatcgagaaatcgcttcagggcttcttctcgctgaagaaggagcgctac
MxDZ2_Nan_RS29095     ------------------------------------------------------------


MxDZ2_Kirby_RS0235795 gtggcagacgtcgacctcgagcgcatcgaagcggaagaggcggccaaagcatacgactcg
MxDZ2_Tam_RS13780     gtggcagacgtcgacctcgagcgcatcgaagcggaagaggcggccaaagcatacgactcg
MxDZ2_Nan_RS29095     ------------------------------------------------------------


MxDZ2_Kirby_RS0235795 ggaggaagagtggttgttggtggaacgctctcggacggcctgcgtgtgacgccagcagca
MxDZ2_Tam_RS13780     ggaggaagagtggttgttggtggaacgctctcggacggcctgcgtgtgacgccagcagca
MxDZ2_Nan_RS29095     ------------------------------------------------------------


MxDZ2_Kirby_RS0235795 accggcggcatccgaccggcccgttcctggggagagtag
MxDZ2_Tam_RS13780     accggcggcatccgaccggcccgttcctggggagagtag
MxDZ2_Nan_RS29095     ---------------------------------------

-----------------------------------------------------------------------------------
multiple sequence alignment (protein)


MxDZ2_Kirby_RS0235795 MMDSYEQMDDIDLRPLTREMDLENLGLSQARTVTGVHVYLDVTNFQSLLEARRENTAELK
MxDZ2_Tam_RS13780     MMDSYEQMDDIDLRPLTREMDLENLGLSQARTVTGVHVYLDVTNFQSLLEARRENTAELK
MxDZ2_Nan_RS29095     MMDSYEQMDDIDLRPLTREMDLENLGLSQARTVTGVHVYLDVTNFQSLLEARRENTAELK
                      ************************************************************

MxDZ2_Kirby_RS0235795 KLLRHLTVFQRQVERLLRQQDGAAVRVHFQGARLHLIVFKPYDTEEGATLARLVKAVTLV
MxDZ2_Tam_RS13780     KLLRHLTVFQRQVERLLRQQDGAAVRVHFQGARLHLIVFKPYDTEEGATLARLVKAVTLV
MxDZ2_Nan_RS29095     KLLRHLTVFQRQVERLLRQQDGAAVRVHFQGARLHLIVFKPYDTEEGATLARLVKAVTLV
                      ************************************************************

MxDZ2_Kirby_RS0235795 EQVRQLADMIQFHTDIHFEFEAGVDTGEAIATMNARARRRQLLFVGRPANEAAKALTGNP
MxDZ2_Tam_RS13780     EQVRQLADMIQFHTDIHFEFEAGVDTGEAIATMNARARRRQLLFVGRPANEAAKALTGNP
MxDZ2_Nan_RS29095     EQVRQLADMIQFHTDIHFEFEAGVDTGEAIATMNARARRRQLLFVGRPANEAAKALTGNP
                      ************************************************************

MxDZ2_Kirby_RS0235795 SAPGEERLAAGAAPFTKQLP-------------SAAAREKLPTSFEAAVKEDIEAHPLES
MxDZ2_Tam_RS13780     SAPGEERLAAGAAPFTKQLP-------------SAAAREKLPTSFEAAVKEDIEAHPLES
MxDZ2_Nan_RS29095     SAPEKSGSPPERPPSRSNSPPLLLERNYRPASRPPSRRTLRPIRWKAWISPTFE------
                      *** :.  ..  .*  .: *             ..: *   *  ::* :.  :*

MxDZ2_Kirby_RS0235795 LDITDVRAPIDYGSLGKKVARLEEAITFFGDLSGFTSYVASLSTEDEKKEALRLLHVLRS
MxDZ2_Tam_RS13780     LDITDVRAPIDYGSLGKKVARLEEAITFFGDLSGFTSYVASLSTEDEKKEALRLLHVLRS
MxDZ2_Nan_RS29095     -------------------------------------------------------HLLTT
                                                                             *:* :

MxDZ2_Kirby_RS0235795 EMQAVVADYDGDFIQFQGDRVQALLYEARTSGRFKSKAVETAAALCSAVDLAKELFPALS
MxDZ2_Tam_RS13780     EMQAVVADYDGDFIQFQGDRVQALLYEARTSGRFKSKAVETAAALCSAVDLAKELFPALS
MxDZ2_Nan_RS29095     ------------------------------------------------------------

MxDZ2_Kirby_RS0235795 TLDVSCGADVGKVLVGRVGLRDDKEVTVLGRSIPCASELQDGAGDGHTAISVRLWEGIEK
MxDZ2_Tam_RS13780     TLDVSCGADVGKVLVGRVGLRDDKEVTVLGRSIPCASELQDGAGDGHTAISVRLWEGIEK
MxDZ2_Nan_RS29095     ---------------------------------------------AH-------W-----
                                                                   .*       *

MxDZ2_Kirby_RS0235795 SLQGFFSLKKERYVADVDLERIEAEEAAKAYDSGGRVVVGGTLSDGLRVTPAATGGIRPA
MxDZ2_Tam_RS13780     SLQGFFSLKKERYVADVDLERIEAEEAAKAYDSGGRVVVGGTLSDGLRVTPAATGGIRPA
MxDZ2_Nan_RS29095     -----------------------------------------------------------A
                                                                                 *

MxDZ2_Kirby_RS0235795 RSWGE
MxDZ2_Tam_RS13780     RSWGE
MxDZ2_Nan_RS29095     R---R
                      *   .
```

**Premature stop codon followed by second ORF prediction 3**

| **Tam_location** | **Tam_nucleotide** | **Kirby_nucleotide** | **Nan_nucleotide** | **Nan_location** |
| --- | --- | --- | --- | --- |
| **4101535** | **G** | **G** | **.** | **4101516** |

```
multiple sequence alignment (nucleotide)

MxDZ2_Kirby_RS0229920 atgaagcccttcgagcagcagacctattacgagattctcgaagtccccgtcaccgctccg
MxDZ2_Tam_RS21920     atgaagcccttcgagcagcagacctattacgagattctcgaagtccccgtcaccgctccg
MxDZ2_Nan_RS20945     atgaagcccttcgagcagcagacctattacgagattctcgaagtccccgtcaccgctccg
                      ************************************************************

MxDZ2_Kirby_RS0229920 aaggaggagattcgggcggcgtacgaacggctgacggagctgtatgcaccggactccatc
MxDZ2_Tam_RS21920     aaggaggagattcgggcggcgtacgaacggctgacggagctgtatgcaccggactccatc
MxDZ2_Nan_RS20945     aaggaggagattcgggcggcgtacgaacggctgacggagctgtatgcaccggactccatc
                      ************************************************************

MxDZ2_Kirby_RS0229920 gccgtctatgcgctcgtggatgagggccaggtggatgagctccgcactcggctcgccgag
MxDZ2_Tam_RS21920     gccgtctatgcgctcgtggatgagggccaggtggatgagctccgcactcggctcgccgag
MxDZ2_Nan_RS20945     gccgtctatgcgctcgtggatgagggccaggtggatgagctccgcactcggctcgccgag
                      ************************************************************

MxDZ2_Kirby_RS0229920 gcgctggagattctctccgacgaagagctgcgtgccgagtacgacaaggacctggggctc
MxDZ2_Tam_RS21920     gcgctggagattctctccgacgaagagctgcgtgccgagtacgacaaggacctggggctc
MxDZ2_Nan_RS20945     gcgctggagattctctccgacgaagagctgcgtgccgagtacgacaaggacctggggctc
                      ************************************************************

MxDZ2_Kirby_RS0229920 ccggccctccggttgatggacgcggtgtcgaccgggggcgaggtggcacacgccgccgcg
MxDZ2_Tam_RS21920     ccggccctccggttgatggacgcggtgtcgaccgggggcgaggtggcacacgccgccgcg
MxDZ2_Nan_RS20945     ccggccctccggttgatggacgcggtgtcgaccgggggcgaggtggcacacgccgccgcg
                      ************************************************************

MxDZ2_Kirby_RS0229920 ccggtggagcagcgcgcggagccggagctcgttggggttgccgcttccggcgcggaagat
MxDZ2_Tam_RS21920     ccggtggagcagcgcgcggagccggagctcgttggggttgccgcttccggcgcggaagat
MxDZ2_Nan_RS20945     ccggtggagcagcgcgcggagccggagctcgttggggttgccgcttccggcgcggaagat
                      ************************************************************

MxDZ2_Kirby_RS0229920 caaactggccacctttcgacgcagatccccgaaacggggacgaacgggaggacaggagca
MxDZ2_Tam_RS21920     caaactggccacctttcgacgcagatccccgaaacggggacgaacgggaggacaggagca
MxDZ2_Nan_RS20945     caaactggccacctttcgacgcagatccccgaaacggggacgaacgggaggacaggagca
                      ************************************************************

MxDZ2_Kirby_RS0229920 gtgcgagacgtgccggtggaggactcggagaccgcgatgacggggcgggatggggaaggg
MxDZ2_Tam_RS21920     gtgcgagacgtgccggtggaggactcggagaccgcgatgacggggcgggatggggaaggg
MxDZ2_Nan_RS20945     gtgcgagacgtgccggtggaggactcggagaccgcgatgacggggcgggatggggaaggg
                      ************************************************************

MxDZ2_Kirby_RS0229920 gaggccggggcccaggcgacggagcctcgggtggaggcgatgccggcgtcctcgggacag
MxDZ2_Tam_RS21920     gaggccggggcccaggcgacggagcctcgggtggaggcgatgccggcgtcctcgggacag
MxDZ2_Nan_RS20945     gaggccggggcccaggcgacggagcctcgggtggaggcgatgccggcgtcctcgggacag
                      ************************************************************

MxDZ2_Kirby_RS0229920 gcggacttccgggcgtcgtttttccgtgggttctcgttcgcatacgtgtccagctcgttg
MxDZ2_Tam_RS21920     gcggacttccgggcgtcgtttttccgtgggttctcgttcgcatacgtgtccagctcgttg
MxDZ2_Nan_RS20945     gcggacttccgggcgtcgtttttccgtgggttctcgttcgcatacgtgtccagctcgttg
                      ************************************************************

MxDZ2_Kirby_RS0229920 caggacacgcagcggctggggagcgcggtggatgttcccgtcgcgtcgattcagcctcag
MxDZ2_Tam_RS21920     caggacacgcagcggctggggagcgcggtggatgttcccgtcgcgtcgattcagcctcag
MxDZ2_Nan_RS20945     caggacacgcagcggctggggagcgcggtggatgttcccgtcgcgtcgattcagcctcag
                      ************************************************************

MxDZ2_Kirby_RS0229920 gtttcgggtagtggcacgggggccgcgagcgctgagtcgcagcgcgaggcccccggtgcg
MxDZ2_Tam_RS21920     gtttcgggtagtggcacgggggccgcgagcgctgagtcgcagcgcgaggcccccggtgcg
MxDZ2_Nan_RS20945     gtttcgggtagtggcacgggggccgcgagcgctgagtcgcagcgcgaggcccccggtgcg
                      ************************************************************

MxDZ2_Kirby_RS0229920 gtggctgcggctccggtgagtgagccgtccgcggcgcatgaggccgctgcggtagtgcct
MxDZ2_Tam_RS21920     gtggctgcggctccggtgagtgagccgtccgcggcgcatgaggccgctgcggtagtgcct
MxDZ2_Nan_RS20945     gtggctgcggctccggtgagtgagccgtccgcggcgcatgaggccgctgcggtagtgcct
                      ************************************************************

MxDZ2_Kirby_RS0229920 ggcgtcgtggcggccagtgcaactgcttcggtttccccaggaggccctgcggtaacgggc
MxDZ2_Tam_RS21920     ggcgtcgtggcggccagtgcaactgcttcggtttccccaggaggccctgcggtaacgggc
MxDZ2_Nan_RS20945     ggcgtcgtggcggccagtgcaactgcttcggtttccccaggaggccctgcggtaacgggc
                      ************************************************************

MxDZ2_Kirby_RS0229920 agcgcgccggctccagcgacggctccccaggtcagcgtcccttcttcgggcagcgacgcc
MxDZ2_Tam_RS21920     agcgcgccggctccagcgacggctccccaggtcagcgtcccttcttcgggcagcgacgcc
MxDZ2_Nan_RS20945     agcgcgccggctccagcgacggctccccaggtcagcgtcccttcttcgggcagcgacgcc
                      ************************************************************


MxDZ2_Kirby_RS0229920 tcggtgcctgcctcggccgccgcgccgacgcctggtccggcaggcgctgtcagcgcgctg
MxDZ2_Tam_RS21920     tcggtgcctgcctcggccgccgcgccgacgcctggtccggcaggcgctgtcagcgcgctg
MxDZ2_Nan_RS20945     tcggtgcctgcctcggccgccgcgccgacgcctggtccggcaggcgctgtcagcgcgctg
                      ************************************************************

MxDZ2_Kirby_RS0229920 gcttcggtcagcgatgcgccggcgcctgcgacggctgccgtgccgacgcccgcctcggcc
MxDZ2_Tam_RS21920     gcttcggtcagcgatgcgccggcgcctgcgacggctgccgtgccgacgcccgcctcggcc
MxDZ2_Nan_RS20945     gcttcggtcagcgatgcgccggcgcctgcgacggctgccgtgccgacgcccgcctcggcc
                      ************************************************************

MxDZ2_Kirby_RS0229920 gcccccgtcagctccccggtggcggaaggcagtgccgcggtggctgcctcggccactcag
MxDZ2_Tam_RS21920     gcccccgtcagctccccggtggcggaaggcagtgccgcggtggctgcctcggccactcag
MxDZ2_Nan_RS20945     gcccccgtcagctccccggtggcggaaggcagtgccgcggtggctgcctcggccactcag
                      ************************************************************

MxDZ2_Kirby_RS0229920 gccagcagcccagccgccgcgccggtccccacgcaaggcgctccggtcagcgccgtggcg
MxDZ2_Tam_RS21920     gccagcagcccagccgccgcgccggtccccacgcaaggcgctccggtcagcgccgtggcg
MxDZ2_Nan_RS20945     gccagcagcccagccgccgcgccggtccccacgcaaggcgctccggtcagcgccgtggcg
                      ************************************************************

MxDZ2_Kirby_RS0229920 gctctgcccccttcggcgcctgccaccgccgtgactggcgccccggcatcggaaaccagc
MxDZ2_Tam_RS21920     gctctgcccccttcggcgcctgccaccgccgtgactggcgccccggcatcggaaaccagc
MxDZ2_Nan_RS20945     gctctgcccccttcggcgcctgccaccgccgtgactggcgccccggcatcggaaaccagc
                      ************************************************************

MxDZ2_Kirby_RS0229920 actgcgccccacctgccggttccagcgagtgcgagcccggtaccgccggccccccaggcg
MxDZ2_Tam_RS21920     actgcgccccacctgccggttccagcgagtgcgagcccggtaccgccggccccccaggcg
MxDZ2_Nan_RS20945     actgcgccccacctgccggttccagcgagtgcgagcccggtaccgccggccccccaggcg
                      ************************************************************

MxDZ2_Kirby_RS0229920 cccgaggccgagtcgaccgccatcgtccctgcccgtccgtcgcccacctcggaggctccg
MxDZ2_Tam_RS21920     cccgaggccgagtcgaccgccatcgtccctgcccgtccgtcgcccacctcggaggctccg
MxDZ2_Nan_RS20945     cccgaggccgagtcgaccgccatcgtccctgcccgtccgtcgcccacctcggaggctccg
                      ************************************************************

MxDZ2_Kirby_RS0229920 tccgagccgggccgggctggcgcgcgtcccgggcggcccctgggagacgcgcagcagatc
MxDZ2_Tam_RS21920     tccgagccgggccgggctggcgcgcgtcccgggcggcccctgggagacgcgcagcagatc
MxDZ2_Nan_RS20945     tccgagccgggccgggctggcgcgcgtcccgggcggcccctgggagacgcgcagcagatc
                      ************************************************************


MxDZ2_Kirby_RS0229920 gcccaggactccgccatcgccacggcggaggcggcgctggcgcaggcgtcccaggccgcg
MxDZ2_Tam_RS21920     gcccaggactccgccatcgccacggcggaggcggcgctggcgcaggcgtcccaggccgcg
MxDZ2_Nan_RS20945     gcccaggactccgccatcgccacggcggaggcggcgctggcgcaggcgtcccaggccgcg
                      ************************************************************

MxDZ2_Kirby_RS0229920 tcgcgggcccgtgagccccggcctcgcgtcccggacatcccgtccgacgcagagttcaac
MxDZ2_Tam_RS21920     tcgcgggcccgtgagccccggcctcgcgtcccggacatcccgtccgacgcagagttcaac
MxDZ2_Nan_RS20945     tcgcgggcccgtgagccccggcctcgcgtcccggacatcccgtccgacgcagagttcaac
                      ************************************************************

MxDZ2_Kirby_RS0229920 ggcgagctgctccggcaggtccgagaggcccgggggGtgaccctccagcaggtggccgac
MxDZ2_Tam_RS21920     ggcgagctgctccggcaggtccgagaggcccgggggGtgaccctccagcaggtggccgac
MxDZ2_Nan_RS20945     ggcgagctgctccggcaggtccgagaggcccggggg-tga--------------------
                      ************************************ ***

MxDZ2_Kirby_RS0229920 cgcacccgcatcacgcgcatgcacctggagaacgtcgaggcggaccgctacaacctgctc
MxDZ2_Tam_RS21920     cgcacccgcatcacgcgcatgcacctggagaacgtcgaggcggaccgctacaacctgctc
MxDZ2_Nan_RS20945     ------------------------------------------------------------


MxDZ2_Kirby_RS0229920 ccgccctcggtctacctgcgaggtatcctgatgagcctcgcccgcgagctggggttggat
MxDZ2_Tam_RS21920     ccgccctcggtctacctgcgaggtatcctgatgagcctcgcccgcgagctggggttggat
MxDZ2_Nan_RS20945     ------------------------------------------------------------


MxDZ2_Kirby_RS0229920 ccactccgcgtctccaagagctacctggccctgtcttctgagaagtcaggccggaagtag
MxDZ2_Tam_RS21920     ccactccgcgtctccaagagctacctggccctgtcttctgagaagtcaggccggaagtag
MxDZ2_Nan_RS20945     ------------------------------------------------------------

-----------------------------------------------------------------------------------
multiple sequence alignment (protein)

MxDZ2_Tam_RS21920     MKPFEQQTYYEILEVPVTAPKEEIRAAYERLTELYAPDSIAVYALVDEGQVDELRTRLAE
MxDZ2_Nan_RS20945     MKPFEQQTYYEILEVPVTAPKEEIRAAYERLTELYAPDSIAVYALVDEGQVDELRTRLAE
MxDZ2_Kirby_RS0229920 MKPFEQQTYYEILEVPVTAPKEEIRAAYERLTELYAPDSIAVYALVDEGQVDELRTRLAE
                      ************************************************************

MxDZ2_Tam_RS21920     ALEILSDEELRAEYDKDLGLPALRLMDAVSTGGEVAHAAAPVEQRAEPELVGVAASGAED
MxDZ2_Nan_RS20945     ALEILSDEELRAEYDKDLGLPALRLMDAVSTGGEVAHAAAPVEQRAEPELVGVAASGAED
MxDZ2_Kirby_RS0229920 ALEILSDEELRAEYDKDLGLPALRLMDAVSTGGEVAHAAAPVEQRAEPELVGVAASGAED
                      ************************************************************

MxDZ2_Tam_RS21920     QTGHLSTQIPETGTNGRTGAVRDVPVEDSETAMTGRDGEGEAGAQATEPRVEAMPASSGQ
MxDZ2_Nan_RS20945     QTGHLSTQIPETGTNGRTGAVRDVPVEDSETAMTGRDGEGEAGAQATEPRVEAMPASSGQ
MxDZ2_Kirby_RS0229920 QTGHLSTQIPETGTNGRTGAVRDVPVEDSETAMTGRDGEGEAGAQATEPRVEAMPASSGQ
                      ************************************************************

MxDZ2_Tam_RS21920     ADFRASFFRGFSFAYVSSSLQDTQRLGSAVDVPVASIQPQVSGSGTGAASAESQREAPGA
MxDZ2_Nan_RS20945     ADFRASFFRGFSFAYVSSSLQDTQRLGSAVDVPVASIQPQVSGSGTGAASAESQREAPGA
MxDZ2_Kirby_RS0229920 ADFRASFFRGFSFAYVSSSLQDTQRLGSAVDVPVASIQPQVSGSGTGAASAESQREAPGA
                      ************************************************************

MxDZ2_Tam_RS21920     VAAAPVSEPSAAHEAAAVVPGVVAASATASVSPGGPAVTGSAPAPATAPQVSVPSSGSDA
MxDZ2_Nan_RS20945     VAAAPVSEPSAAHEAAAVVPGVVAASATASVSPGGPAVTGSAPAPATAPQVSVPSSGSDA
MxDZ2_Kirby_RS0229920 VAAAPVSEPSAAHEAAAVVPGVVAASATASVSPGGPAVTGSAPAPATAPQVSVPSSGSDA
                      ************************************************************

MxDZ2_Tam_RS21920     SVPASAAAPTPGPAGAVSALASVSDAPAPATAAVPTPASAAPVSSPVAEGSAAVAASATQ
MxDZ2_Nan_RS20945     SVPASAAAPTPGPAGAVSALASVSDAPAPATAAVPTPASAAPVSSPVAEGSAAVAASATQ
MxDZ2_Kirby_RS0229920 SVPASAAAPTPGPAGAVSALASVSDAPAPATAAVPTPASAAPVSSPVAEGSAAVAASATQ
                      ************************************************************

MxDZ2_Tam_RS21920     ASSPAAAPVPTQGAPVSAVAALPPSAPATAVTGAPASETSTAPHLPVPASASPVPPAPQA
MxDZ2_Nan_RS20945     ASSPAAAPVPTQGAPVSAVAALPPSAPATAVTGAPASETSTAPHLPVPASASPVPPAPQA
MxDZ2_Kirby_RS0229920 ASSPAAAPVPTQGAPVSAVAALPPSAPATAVTGAPASETSTAPHLPVPASASPVPPAPQA
                      ************************************************************

MxDZ2_Tam_RS21920     PEAESTAIVPARPSPTSEAPSEPGRAGARPGRPLGDAQQIAQDSAIATAEAALAQASQAA
MxDZ2_Nan_RS20945     PEAESTAIVPARPSPTSEAPSEPGRAGARPGRPLGDAQQIAQDSAIATAEAALAQASQAA
MxDZ2_Kirby_RS0229920 PEAESTAIVPARPSPTSEAPSEPGRAGARPGRPLGDAQQIAQDSAIATAEAALAQASQAA
                      ************************************************************

MxDZ2_Tam_RS21920     SRAREPRPRVPDIPSDAEFNGELLRQVREARGVTLQQVADRTRITRMHLENVEADRYNLL
MxDZ2_Nan_RS20945     SRAREPRPRVPDIPSDAEFNGELLRQVREARG----------------------------
MxDZ2_Kirby_RS0229920 SRAREPRPRVPDIPSDAEFNGELLRQVREARGVTLQQVADRTRITRMHLENVEADRYNLL
                      ********************************

MxDZ2_Tam_RS21920     PPSVYLRGILMSLARELGLDPLRVSKSYLALSSEKSGRK
MxDZ2_Nan_RS20945     ---------------------------------------
MxDZ2_Kirby_RS0229920 PPSVYLRGILMSLARELGLDPLRVSKSYLALSSEKSGRK
```

**Premature stop codon followed by second ORF prediction 4**

| **Tam_location** | **Tam_nucleotide** | **Kirby_nucleotide** | **Nan_nucleotide** | **Nan_location** |
| --- | --- | --- | --- | --- |
| **5471988** | **G (complement)** | **G (complement)** | **.** | **5471963** |

```
multiple sequence alignment (nucleotide)

MxDZ2_Kirby_RS0227310 atgaatctgggggaactgctcgccgagctgaatcggcggggcctggaagtccgggtggaa
MxDZ2_Tam_RS26590     atgaatctgggggaactgctcgccgagctgaatcggcggggcctggaagtccgggtggaa
MxDZ2_Nan_RS16270     atgaatctgggggaactgctcgccgagctgaatcggcggggcctggaagtccgggtggaa
                      ************************************************************

MxDZ2_Kirby_RS0227310 ggcgacgcgctgaggcttcgcggaccgaaaggcgccgccaccgcagagcttcggcaggca
MxDZ2_Tam_RS26590     ggcgacgcgctgaggcttcgcggaccgaaaggcgccgccaccgcagagcttcggcaggca
MxDZ2_Nan_RS16270     ggcgacgcgctgaggcttcgcggaccgaaaggcgccgccaccgcagagcttcggcaggca
                      ************************************************************

MxDZ2_Kirby_RS0227310 ctggcggaacacaaacccgagttgctcgaactcctgcgtgcccatgaccgggggcaggag
MxDZ2_Tam_RS26590     ctggcggaacacaaacccgagttgctcgaactcctgcgtgcccatgaccgggggcaggag
MxDZ2_Nan_RS16270     ctggcggaacacaaacccgagttgctcgaactcctgcgtgcccatgaccgggggcaggag
                      ************************************************************

MxDZ2_Kirby_RS0227310 gaggcgcccatcgtcccggttcaacgcaccgggcccgcgccgctctcctatgggcagacg
MxDZ2_Tam_RS26590     gaggcgcccatcgtcccggttcaacgcaccgggcccgcgccgctctcctatgggcagacg
MxDZ2_Nan_RS16270     gaggcgcccatcgtcccggttcaacgcaccgggcccgcgccgctctcctatgggcagacg
                      ************************************************************

MxDZ2_Kirby_RS0227310 cggctgtggttcctcgaccgcttggagcctgggagcaccgcctacaacctgatgctggcg
MxDZ2_Tam_RS26590     cggctgtggttcctcgaccgcttggagcctgggagcaccgcctacaacctgatgctggcg
MxDZ2_Nan_RS16270     cggctgtggttcctcgaccgcttggagcctgggagcaccgcctacaacctgatgctggcg
                      ************************************************************

MxDZ2_Kirby_RS0227310 ttgcaggtcgaagggcacctcgacgtagccatcgtgaagcgctgcttcaccgaggtcatc
MxDZ2_Tam_RS26590     ttgcaggtcgaagggcacctcgacgtagccatcgtgaagcgctgcttcaccgaggtcatc
MxDZ2_Nan_RS16270     ttgcaggtcgaagggcacctcgacgtagccatcgtgaagcgctgcttcaccgaggtcatc
                      ************************************************************

MxDZ2_Kirby_RS0227310 cgccgccacgaggtgctgcgcacgcggtatgccgagcacgcgggcgtcccggtccagatt
MxDZ2_Tam_RS26590     cgccgccacgaggtgctgcgcacgcggtatgccgagcacgcgggcgtcccggtccagatt
MxDZ2_Nan_RS16270     cgccgccacgaggtgctgcgcacgcggtatgccgagcacgcgggcgtcccggtccagatt
                      ************************************************************

MxDZ2_Kirby_RS0227310 gtcgacccggagccccggttcgacttccgcgtgctcgacgagcgcgaggtgctggcctcc
MxDZ2_Tam_RS26590     gtcgacccggagccccggttcgacttccgcgtgctcgacgagcgcgaggtgctggcctcc
MxDZ2_Nan_RS16270     gtcgacccggagccccggttcgacttccgcgtgctcgacgagcgcgaggtgctggcctcc
                      ************************************************************

MxDZ2_Kirby_RS0227310 gaacccgcgggcgtggaggccttcctccagcgagagggcaagcggcccttcgacctgtcc
MxDZ2_Tam_RS26590     gaacccgcgggcgtggaggccttcctccagcgagagggcaagcggcccttcgacctgtcc
MxDZ2_Nan_RS16270     gaacccgcgggcgtggaggccttcctccagcgagagggcaagcggcccttcgacctgtcc
                      ************************************************************

MxDZ2_Kirby_RS0227310 acgggccccgtgatgcgtgtgctcatcatcgagcgggggccccaggggcagtacctccag
MxDZ2_Tam_RS26590     acgggccccgtgatgcgtgtgctcatcatcgagcgggggccccaggggcagtacctccag
MxDZ2_Nan_RS16270     acgggccccgtgatgcgtgtgctcatcatcgagcgggggccccaggggcagtacctccag
                      ************************************************************

MxDZ2_Kirby_RS0227310 ctctgcttgcaccacatcgcggcggacgtgtgggcgcaggccatcctcgtccgcgaaatc
MxDZ2_Tam_RS26590     ctctgcttgcaccacatcgcggcggacgtgtgggcgcaggccatcctcgtccgcgaaatc
MxDZ2_Nan_RS16270     ctctgcttgcaccacatcgcggcggacgtgtgggcgcaggccatcctcgtccgcgaaatc
                      ************************************************************

MxDZ2_Kirby_RS0227310 gtgacgttatatggcgccttcgccagcggccaaccttcgccgttgccgcccctgacgctc
MxDZ2_Tam_RS26590     gtgacgttatatggcgccttcgccagcggccaaccttcgccgttgccgcccctgacgctc
MxDZ2_Nan_RS16270     gtgacgttatatggcgccttcgccagcggccaaccttcgccgttgccgcccctgacgctc
                      ************************************************************

MxDZ2_Kirby_RS0227310 cagtactccgacttcgctgtctggcagcgggagtacctccagggcgaggttcgccagaag
MxDZ2_Tam_RS26590     cagtactccgacttcgctgtctggcagcgggagtacctccagggcgaggttcgccagaag
MxDZ2_Nan_RS16270     cagtactccgacttcgctgtctggcagcgggagtacctccagggcgaggttcgccagaag
                      ************************************************************

MxDZ2_Kirby_RS0227310 ctggtggactactggcgcaggaagctggagggggcgcctccgctgctcgagctccccacg
MxDZ2_Tam_RS26590     ctggtggactactggcgcaggaagctggagggggcgcctccgctgctcgagctccccacg
MxDZ2_Nan_RS16270     ctggtggactactggcgcaggaagctggagggggcgcctccgctgctcgagctccccacg
                      ************************************************************

MxDZ2_Kirby_RS0227310 gaccatcctcgccccaaggtgcagtcgaaccgaggcggcgaggtccgcttcgaagtgggc
MxDZ2_Tam_RS26590     gaccatcctcgccccaaggtgcagtcgaaccgaggcggcgaggtccgcttcgaagtgggc
MxDZ2_Nan_RS16270     gaccatcctcgccccaaggtgcagtcgaaccgaggcggcgaggtccgcttcgaagtgggc
                      ************************************************************


MxDZ2_Kirby_RS0227310 ccccagctcaccgccgccctgaaggcacagagccacacggcccgcaccacgcccttcgtg
MxDZ2_Tam_RS26590     ccccagctcaccgccgccctgaaggcacagagccacacggcccgcaccacgcccttcgtg
MxDZ2_Nan_RS16270     ccccagctcaccgccgccctgaaggcacagagccacacggcccgcaccacgcccttcgtg
                      ************************************************************

MxDZ2_Kirby_RS0227310 gggatgctgagcgcgttcttcgtgctcctgcaccgcctcacggggcgtgaggacctggtc
MxDZ2_Tam_RS26590     gggatgctgagcgcgttcttcgtgctcctgcaccgcctcacggggcgtgaggacctggtc
MxDZ2_Nan_RS16270     gggatgctgagcgcgttcttcgtgctcctgcaccgcctcacggggcgtgaggacctggtc
                      ************************************************************

MxDZ2_Kirby_RS0227310 ctgggcgcgaactccatcaaccggacgcgtacggagctggagccgttggtgggcttcttc
MxDZ2_Tam_RS26590     ctgggcgcgaactccatcaaccggacgcgtacggagctggagccgttggtgggcttcttc
MxDZ2_Nan_RS16270     ctgggcgcgaactccatcaaccggacgcgtacggagctggagccgttggtgggcttcttc
                      ************************************************************

MxDZ2_Kirby_RS0227310 gtggacaacctggtgatgcgggtggacctcggtggcggcccggggttcgccacggtgctg
MxDZ2_Tam_RS26590     gtggacaacctggtgatgcgggtggacctcggtggcggcccggggttcgccacggtgctg
MxDZ2_Nan_RS16270     gtggacaacctggtgatgcgggtggacctcggtggcggcccggggttcgccacggtgctg
                      ************************************************************

MxDZ2_Kirby_RS0227310 gagcgcgtgcgtgagacggtgctcgatgccttcgctcaccaggacctgcccttcgacctg
MxDZ2_Tam_RS26590     gagcgcgtgcgtgagacggtgctcgatgccttcgctcaccaggacctgcccttcgacctg
MxDZ2_Nan_RS16270     gagcgcgtgcgtgagacggtgctcgatgccttcgctcaccaggacctgcccttcgacctg
                      ************************************************************

MxDZ2_Kirby_RS0227310 ctcgtggaggagctgaggcccgcgcgcagcctgggcttcaacccgctgttccaggtcgtg
MxDZ2_Tam_RS26590     ctcgtggaggagctgaggcccgcgcgcagcctgggcttcaacccgctgttccaggtcgtg
MxDZ2_Nan_RS16270     ctcgtggaggagctgaggcccgcgcgcagcctgggcttcaacccgctgttccaggtcgtg
                      ************************************************************

MxDZ2_Kirby_RS0227310 ttcgcctgggtccgcgcttccggcgaggcccaggacgagacgggcctccgcatgcggccg
MxDZ2_Tam_RS26590     ttcgcctgggtccgcgcttccggcgaggcccaggacgagacgggcctccgcatgcggccg
MxDZ2_Nan_RS16270     ttcgcctgggtccgcgcttccggcgaggcccaggacgagacgggcctccgcatgcggccg
                      ************************************************************

MxDZ2_Kirby_RS0227310 ctcgagttcgaggcggaatcatcgcgcttcgacctcaacctcttcgtggatgaccacggc
MxDZ2_Tam_RS26590     ctcgagttcgaggcggaatcatcgcgcttcgacctcaacctcttcgtggatgaccacggc
MxDZ2_Nan_RS16270     ctcgagttcgaggcggaatcatcgcgcttcgacctcaacctcttcgtggatgaccacggc
                      ************************************************************


MxDZ2_Kirby_RS0227310 gaccggctgtcggcgcggatggtgttcaaccgcgacctcttcgagcagggcaccatccag
MxDZ2_Tam_RS26590     gaccggctgtcggcgcggatggtgttcaaccgcgacctcttcgagcagggcaccatccag
MxDZ2_Nan_RS16270     gaccggctgtcggcgcggatggtgttcaaccgcgacctcttcgagcagggcaccatccag
                      ************************************************************

MxDZ2_Kirby_RS0227310 cactacgtggactgcttccaggtgctgctcgaagggctcgtggccgagccccagcggccg
MxDZ2_Tam_RS26590     cactacgtggactgcttccaggtgctgctcgaagggctcgtggccgagccccagcggccg
MxDZ2_Nan_RS16270     cactacgtggactgcttccaggtgctgctcgaagggctcgtggccgagccccagcggccg
                      ************************************************************

MxDZ2_Kirby_RS0227310 gtgggcgagctgcccgtgctccccgcggatgtccgcgagcgcgtgctccggcagtggaac
MxDZ2_Tam_RS26590     gtgggcgagctgcccgtgctccccgcggatgtccgcgagcgcgtgctccggcagtggaac
MxDZ2_Nan_RS16270     gtgggcgagctgcccgtgctccccgcggatgtccgcgagcgcgtgctccggcagtggaac
                      ************************************************************

MxDZ2_Kirby_RS0227310 gacacggggacgccggaggcggaaggcgcgtgcctccatgcgctcatcgaggcccaggcg
MxDZ2_Tam_RS26590     gacacggggacgccggaggcggaaggcgcgtgcctccatgcgctcatcgaggcccaggcg
MxDZ2_Nan_RS16270     gacacggggacgccggaggcggaaggcgcgtgcctccatgcgctcatcgaggcccaggcg
                      ************************************************************

MxDZ2_Kirby_RS0227310 ctgcgcacgccggatgcggtggccatcgtcgtggacgactgggagctgacctacggcgag
MxDZ2_Tam_RS26590     ctgcgcacgccggatgcggtggccatcgtcgtggacgactgggagctgacctacggcgag
MxDZ2_Nan_RS16270     ctgcgcacgccggatgcggtggccatcgtcgtggacgactgggagctgacctacggcgag
                      ************************************************************

MxDZ2_Kirby_RS0227310 ctcgaccagctcagcgaccgggtcgcagcgtcactccaggatttggacgtgggaccggag
MxDZ2_Tam_RS26590     ctcgaccagctcagcgaccgggtcgcagcgtcactccaggatttggacgtgggaccggag
MxDZ2_Nan_RS16270     ctcgaccagctcagcgaccgggtcgcagcgtcactccaggatttggacgtgggaccggag
                      ************************************************************

MxDZ2_Kirby_RS0227310 gtggtggtcggcgtctacctggaccgttcggcggagctcatcgtgagcctgctggcggtg
MxDZ2_Tam_RS26590     gtggtggtcggcgtctacctggaccgttcggcggagctcatcgtgagcctgctggcggtg
MxDZ2_Nan_RS16270     gtggtggtcggcgtctacctggaccgttcggcggagctcatcgtgagcctgctggcggtg
                      ************************************************************

MxDZ2_Kirby_RS0227310 atgaaggcagggggcgcgttcctcgcgctcgacgcagacgagcccgtggaccggctccgc
MxDZ2_Tam_RS26590     atgaaggcagggggcgcgttcctcgcgctcgacgcagacgagcccgtggaccggctccgc
MxDZ2_Nan_RS16270     atgaaggcagggggcgcgttcctcgcgctcgacgcagacgagcccgtggaccggctccgc
                      ************************************************************


MxDZ2_Kirby_RS0227310 cacatcgtggcggatgcccggCcccgggtggtgatttcttccgcgaagctgtccgagcgc
MxDZ2_Tam_RS26590     cacatcgtggcggatgcccggCcccgggtggtgatttcttccgcgaagctgtccgagcgc
MxDZ2_Nan_RS16270     cacatcgtggcggatgcccgg-cccgggtggtga--------------------------
                      ********************* ************

MxDZ2_Kirby_RS0227310 ctctggggcatggggggcttcgtcacgctccacgtggacgaaggctaccgggacatgccc
MxDZ2_Tam_RS26590     ctctggggcatggggggcttcgtcacgctccacgtggacgaaggctaccgggacatgccc
MxDZ2_Nan_RS16270     ------------------------------------------------------------


MxDZ2_Kirby_RS0227310 gccgcgccgggccagcagctccgccgcgacgtcctccccgaccacctggcctacatcctc
MxDZ2_Tam_RS26590     gccgcgccgggccagcagctccgccgcgacgtcctccccgaccacctggcctacatcctc
MxDZ2_Nan_RS16270     ------------------------------------------------------------


MxDZ2_Kirby_RS0227310 tacacgtcaggctcgacgggccggcccaagggcacggaaatcacgcaccggagcatcgtc
MxDZ2_Tam_RS26590     tacacgtcaggctcgacgggccggcccaagggcacggaaatcacgcaccggagcatcgtc
MxDZ2_Nan_RS16270     ------------------------------------------------------------


MxDZ2_Kirby_RS0227310 aactacctgcgctggagcgtggacgcctaccggcttcgcgaagggacggggagcccggtg
MxDZ2_Tam_RS26590     aactacctgcgctggagcgtggacgcctaccggcttcgcgaagggacggggagcccggtg
MxDZ2_Nan_RS16270     ------------------------------------------------------------


MxDZ2_Kirby_RS0227310 attggctccgtcagcttcgacggaacgctgacgagcctcttcgcgccgctcctggcgggc
MxDZ2_Tam_RS26590     attggctccgtcagcttcgacggaacgctgacgagcctcttcgcgccgctcctggcgggc
MxDZ2_Nan_RS16270     ------------------------------------------------------------


MxDZ2_Kirby_RS0227310 cgcgcgctgttcctcgtgccacgtggtcacgaaatcgaccagttgacgtcccgggactac
MxDZ2_Tam_RS26590     cgcgcgctgttcctcgtgccacgtggtcacgaaatcgaccagttgacgtcccgggactac
MxDZ2_Nan_RS16270     ------------------------------------------------------------


MxDZ2_Kirby_RS0227310 cccgagcagggcttcagcttcatcaagatgacgccgtcgcacctgcgggccttcaacggc
MxDZ2_Tam_RS26590     cccgagcagggcttcagcttcatcaagatgacgccgtcgcacctgcgggccttcaacggc
MxDZ2_Nan_RS16270     ------------------------------------------------------------


MxDZ2_Kirby_RS0227310 ctggggcgcacgcgcgaggtgctcggccgcacccatgcggtggtgctgggcggggaaggg
MxDZ2_Tam_RS26590     ctggggcgcacgcgcgaggtgctcggccgcacccatgcggtggtgctgggcggggaaggg
MxDZ2_Nan_RS16270     ------------------------------------------------------------


MxDZ2_Kirby_RS0227310 ctgcacggcgtggatctggccccatggcgggagcagggcctccccacgcgcgtcatcaac
MxDZ2_Tam_RS26590     ctgcacggcgtggatctggccccatggcgggagcagggcctccccacgcgcgtcatcaac
MxDZ2_Nan_RS16270     ------------------------------------------------------------


MxDZ2_Kirby_RS0227310 gagtacggccccaccgaggccgcggtggcgtgctgcttcgagacactgctgccggatgga
MxDZ2_Tam_RS26590     gagtacggccccaccgaggccgcggtggcgtgctgcttcgagacactgctgccggatgga
MxDZ2_Nan_RS16270     ------------------------------------------------------------


MxDZ2_Kirby_RS0227310 acacctcccccggagcgggtgcccattggccggcccatctcgcacatgcggttgtacatc
MxDZ2_Tam_RS26590     acacctcccccggagcgggtgcccattggccggcccatctcgcacatgcggttgtacatc
MxDZ2_Nan_RS16270     ------------------------------------------------------------


MxDZ2_Kirby_RS0227310 ctggaccggtacctccagcccgtgcccgttggcgtacccggcgagctgtacatcggcggc
MxDZ2_Tam_RS26590     ctggaccggtacctccagcccgtgcccgttggcgtacccggcgagctgtacatcggcggc
MxDZ2_Nan_RS16270     ------------------------------------------------------------


MxDZ2_Kirby_RS0227310 gtcggactggcccgaggctacctgcgccggccggacctgacggcggagcgcttcgtcccc
MxDZ2_Tam_RS26590     gtcggactggcccgaggctacctgcgccggccggacctgacggcggagcgcttcgtcccc
MxDZ2_Nan_RS16270     ------------------------------------------------------------


MxDZ2_Kirby_RS0227310 aaccccttcgacgggggcctggagggctcgcggctctaccgcacgggggaccacgcccgg
MxDZ2_Tam_RS26590     aaccccttcgacgggggcctggagggctcgcggctctaccgcacgggggaccacgcccgg
MxDZ2_Nan_RS16270     ------------------------------------------------------------


MxDZ2_Kirby_RS0227310 tacctctccgacggccgcatcgaatacctgggccgccaggacgaccagctcaagattcgc
MxDZ2_Tam_RS26590     tacctctccgacggccgcatcgaatacctgggccgccaggacgaccagctcaagattcgc
MxDZ2_Nan_RS16270     ------------------------------------------------------------


MxDZ2_Kirby_RS0227310 ggtcaccgcgtcgagacgggagaggtggaggccgcgctgggccgccatcccgacgtcgtc
MxDZ2_Tam_RS26590     ggtcaccgcgtcgagacgggagaggtggaggccgcgctgggccgccatcccgacgtcgtc
MxDZ2_Nan_RS16270     ------------------------------------------------------------


MxDZ2_Kirby_RS0227310 caggccgcggtgctcctgcagcgcctcccgagcggggcgccgcgcctggtggcctacgtg
MxDZ2_Tam_RS26590     caggccgcggtgctcctgcagcgcctcccgagcggggcgccgcgcctggtggcctacgtg
MxDZ2_Nan_RS16270     ------------------------------------------------------------


MxDZ2_Kirby_RS0227310 cagccccagtcgatggagaacggtgacctgcgggccgagctgcgcaagtccctgcgtgag
MxDZ2_Tam_RS26590     cagccccagtcgatggagaacggtgacctgcgggccgagctgcgcaagtccctgcgtgag
MxDZ2_Nan_RS16270     ------------------------------------------------------------


MxDZ2_Kirby_RS0227310 gtgttgcccgagtacatgatgccggaggtcatcgcggtcctcccggagctgccgctgacg
MxDZ2_Tam_RS26590     gtgttgcccgagtacatgatgccggaggtcatcgcggtcctcccggagctgccgctgacg
MxDZ2_Nan_RS16270     ------------------------------------------------------------


MxDZ2_Kirby_RS0227310 cccagcgggaagattgaccgcaaggcgctcccgcccgtggcgtccgaggcccccgtcgcc
MxDZ2_Tam_RS26590     cccagcgggaagattgaccgcaaggcgctcccgcccgtggcgtccgaggcccccgtcgcc
MxDZ2_Nan_RS16270     ------------------------------------------------------------


MxDZ2_Kirby_RS0227310 agcgcgctggcccgcacggaagcgcggaccgagacggagcggcaactccaggcgctgttc
MxDZ2_Tam_RS26590     agcgcgctggcccgcacggaagcgcggaccgagacggagcggcaactccaggcgctgttc
MxDZ2_Nan_RS16270     ------------------------------------------------------------


MxDZ2_Kirby_RS0227310 ggtgaactcctgggactgagctccgtggcgcccaccgacagcttcttcgaactgggaggc
MxDZ2_Tam_RS26590     ggtgaactcctgggactgagctccgtggcgcccaccgacagcttcttcgaactgggaggc
MxDZ2_Nan_RS16270     ------------------------------------------------------------


MxDZ2_Kirby_RS0227310 cattcgctcctcgccgtcacgctcatctcgcgcatctccgcgaagctggacatcgaggtg
MxDZ2_Tam_RS26590     cattcgctcctcgccgtcacgctcatctcgcgcatctccgcgaagctggacatcgaggtg
MxDZ2_Nan_RS16270     ------------------------------------------------------------


MxDZ2_Kirby_RS0227310 cccctcaacgaggtgttcgaccggccctccgtggagtccctggcgcactggattgacgaa
MxDZ2_Tam_RS26590     cccctcaacgaggtgttcgaccggccctccgtggagtccctggcgcactggattgacgaa
MxDZ2_Nan_RS16270     ------------------------------------------------------------


MxDZ2_Kirby_RS0227310 cactccagcaaggtcaccgcgctgattcgccagcttcccgcctgtgtggtggcgctgaag
MxDZ2_Tam_RS26590     cactccagcaaggtcaccgcgctgattcgccagcttcccgcctgtgtggtggcgctgaag
MxDZ2_Nan_RS16270     ------------------------------------------------------------


MxDZ2_Kirby_RS0227310 cccctgggacacaaaccgccgctgttccttgccccgccgtccgctggcagtccggccgtc
MxDZ2_Tam_RS26590     cccctgggacacaaaccgccgctgttccttgccccgccgtccgctggcagtccggccgtc
MxDZ2_Nan_RS16270     ------------------------------------------------------------


MxDZ2_Kirby_RS0227310 tacgtcaccctggcccgccacctggacgcggagcagcccgtgttcggcttccagatgcct
MxDZ2_Tam_RS26590     tacgtcaccctggcccgccacctggacgcggagcagcccgtgttcggcttccagatgcct
MxDZ2_Nan_RS16270     ------------------------------------------------------------


MxDZ2_Kirby_RS0227310 ggcgtcatggacgaccaactgccacccgagaccatcgaggacaccgccgcgctgtatgtg
MxDZ2_Tam_RS26590     ggcgtcatggacgaccaactgccacccgagaccatcgaggacaccgccgcgctgtatgtg
MxDZ2_Nan_RS16270     ------------------------------------------------------------


MxDZ2_Kirby_RS0227310 gcggccctgcggcaggtccagccgcgcgggccgtaccggatcgcgggctggtcctacgga
MxDZ2_Tam_RS26590     gcggccctgcggcaggtccagccgcgcgggccgtaccggatcgcgggctggtcctacgga
MxDZ2_Nan_RS16270     ------------------------------------------------------------


MxDZ2_Kirby_RS0227310 ggactcgtcgtctgtgaaatggcgcggcagctcgaagcgctcggggaacaggtggcgctg
MxDZ2_Tam_RS26590     ggactcgtcgtctgtgaaatggcgcggcagctcgaagcgctcggggaacaggtggcgctg
MxDZ2_Nan_RS16270     ------------------------------------------------------------


MxDZ2_Kirby_RS0227310 ctcggcttgattgacggcgcggcgttggaccggttgtccgcgcacgacggtaccccgcag
MxDZ2_Tam_RS26590     ctcggcttgattgacggcgcggcgttggaccggttgtccgcgcacgacggtaccccgcag
MxDZ2_Nan_RS16270     ------------------------------------------------------------


MxDZ2_Kirby_RS0227310 gagattcccgcgggctcccagctcatcaaggtgctgggggaagcccagatgccgcgggac
MxDZ2_Tam_RS26590     gagattcccgcgggctcccagctcatcaaggtgctgggggaagcccagatgccgcgggac
MxDZ2_Nan_RS16270     ------------------------------------------------------------


MxDZ2_Kirby_RS0227310 tatgccagcctgagattgataggcgaatggatgggaatcagcctgcccgaggcatgggca
MxDZ2_Tam_RS26590     tatgccagcctgagattgataggcgaatggatgggaatcagcctgcccgaggcatgggca
MxDZ2_Nan_RS16270     ------------------------------------------------------------


MxDZ2_Kirby_RS0227310 gacctgcttcgtcgggatacagatgggcagtattcatacctgtggcgtttcttgcgagac
MxDZ2_Tam_RS26590     gacctgcttcgtcgggatacagatgggcagtattcatacctgtggcgtttcttgcgagac
MxDZ2_Nan_RS16270     ------------------------------------------------------------


MxDZ2_Kirby_RS0227310 agcgcgctgacggcccggaatttcctggcaacacgccgctcggagcaactctatacgttc
MxDZ2_Tam_RS26590     agcgcgctgacggcccggaatttcctggcaacacgccgctcggagcaactctatacgttc
MxDZ2_Nan_RS16270     ------------------------------------------------------------


MxDZ2_Kirby_RS0227310 tcttcgtatggaggctccgccacacttttcaggacgggtccggtgacgattggaggcgat
MxDZ2_Tam_RS26590     tcttcgtatggaggctccgccacacttttcaggacgggtccggtgacgattggaggcgat
MxDZ2_Nan_RS16270     ------------------------------------------------------------


MxDZ2_Kirby_RS0227310 ccgctcgtcgacagtgtgcgcagattcgcgctcgcggggacggaagtgataccggttccg
MxDZ2_Tam_RS26590     ccgctcgtcgacagtgtgcgcagattcgcgctcgcggggacggaagtgataccggttccg
MxDZ2_Nan_RS16270     ------------------------------------------------------------


MxDZ2_Kirby_RS0227310 ggcaatcacatgacgctcatcatggatgaacggaatgtcgccgtgcttgcaattgaaata
MxDZ2_Tam_RS26590     ggcaatcacatgacgctcatcatggatgaacggaatgtcgccgtgcttgcaattgaaata
MxDZ2_Nan_RS16270     ------------------------------------------------------------


MxDZ2_Kirby_RS0227310 cagaaatgtctggatagggctttgacgctcccgccaatcgcagaaatgtctcgggcggaa
MxDZ2_Tam_RS26590     cagaaatgtctggatagggctttgacgctcccgccaatcgcagaaatgtctcgggcggaa
MxDZ2_Nan_RS16270     ------------------------------------------------------------


MxDZ2_Kirby_RS0227310 atgccagcaaaggggctggctgcgcagctcctcatggaggtcatgtaa
MxDZ2_Tam_RS26590     atgccagcaaaggggctggctgcgcagctcctcatggaggtcatgtaa
MxDZ2_Nan_RS16270     ------------------------------------------------

-----------------------------------------------------------------------------------
multiple sequence alignment (protein)

MxDZ2_Kirby_RS0227310 MNLGELLAELNRRGLEVRVEGDALRLRGPKGAATAELRQALAEHKPELLELLRAHDRGQE
MxDZ2_Tam_RS26590     MNLGELLAELNRRGLEVRVEGDALRLRGPKGAATAELRQALAEHKPELLELLRAHDRGQE
MxDZ2_Nan_RS16270     MNLGELLAELNRRGLEVRVEGDALRLRGPKGAATAELRQALAEHKPELLELLRAHDRGQE
                      ************************************************************

MxDZ2_Kirby_RS0227310 EAPIVPVQRTGPAPLSYGQTRLWFLDRLEPGSTAYNLMLALQVEGHLDVAIVKRCFTEVI
MxDZ2_Tam_RS26590     EAPIVPVQRTGPAPLSYGQTRLWFLDRLEPGSTAYNLMLALQVEGHLDVAIVKRCFTEVI
MxDZ2_Nan_RS16270     EAPIVPVQRTGPAPLSYGQTRLWFLDRLEPGSTAYNLMLALQVEGHLDVAIVKRCFTEVI
                      ************************************************************

MxDZ2_Kirby_RS0227310 RRHEVLRTRYAEHAGVPVQIVDPEPRFDFRVLDEREVLASEPAGVEAFLQREGKRPFDLS
MxDZ2_Tam_RS26590     RRHEVLRTRYAEHAGVPVQIVDPEPRFDFRVLDEREVLASEPAGVEAFLQREGKRPFDLS
MxDZ2_Nan_RS16270     RRHEVLRTRYAEHAGVPVQIVDPEPRFDFRVLDEREVLASEPAGVEAFLQREGKRPFDLS
                      ************************************************************

MxDZ2_Kirby_RS0227310 TGPVMRVLIIERGPQGQYLQLCLHHIAADVWAQAILVREIVTLYGAFASGQPSPLPPLTL
MxDZ2_Tam_RS26590     TGPVMRVLIIERGPQGQYLQLCLHHIAADVWAQAILVREIVTLYGAFASGQPSPLPPLTL
MxDZ2_Nan_RS16270     TGPVMRVLIIERGPQGQYLQLCLHHIAADVWAQAILVREIVTLYGAFASGQPSPLPPLTL
                      ************************************************************

MxDZ2_Kirby_RS0227310 QYSDFAVWQREYLQGEVRQKLVDYWRRKLEGAPPLLELPTDHPRPKVQSNRGGEVRFEVG
MxDZ2_Tam_RS26590     QYSDFAVWQREYLQGEVRQKLVDYWRRKLEGAPPLLELPTDHPRPKVQSNRGGEVRFEVG
MxDZ2_Nan_RS16270     QYSDFAVWQREYLQGEVRQKLVDYWRRKLEGAPPLLELPTDHPRPKVQSNRGGEVRFEVG
                      ************************************************************

MxDZ2_Kirby_RS0227310 PQLTAALKAQSHTARTTPFVGMLSAFFVLLHRLTGREDLVLGANSINRTRTELEPLVGFF
MxDZ2_Tam_RS26590     PQLTAALKAQSHTARTTPFVGMLSAFFVLLHRLTGREDLVLGANSINRTRTELEPLVGFF
MxDZ2_Nan_RS16270     PQLTAALKAQSHTARTTPFVGMLSAFFVLLHRLTGREDLVLGANSINRTRTELEPLVGFF
                      ************************************************************

MxDZ2_Kirby_RS0227310 VDNLVMRVDLGGGPGFATVLERVRETVLDAFAHQDLPFDLLVEELRPARSLGFNPLFQVV
MxDZ2_Tam_RS26590     VDNLVMRVDLGGGPGFATVLERVRETVLDAFAHQDLPFDLLVEELRPARSLGFNPLFQVV
MxDZ2_Nan_RS16270     VDNLVMRVDLGGGPGFATVLERVRETVLDAFAHQDLPFDLLVEELRPARSLGFNPLFQVV
                      ************************************************************


MxDZ2_Kirby_RS0227310 FAWVRASGEAQDETGLRMRPLEFEAESSRFDLNLFVDDHGDRLSARMVFNRDLFEQGTIQ
MxDZ2_Tam_RS26590     FAWVRASGEAQDETGLRMRPLEFEAESSRFDLNLFVDDHGDRLSARMVFNRDLFEQGTIQ
MxDZ2_Nan_RS16270     FAWVRASGEAQDETGLRMRPLEFEAESSRFDLNLFVDDHGDRLSARMVFNRDLFEQGTIQ
                      ************************************************************

MxDZ2_Kirby_RS0227310 HYVDCFQVLLEGLVAEPQRPVGELPVLPADVRERVLRQWNDTGTPEAEGACLHALIEAQA
MxDZ2_Tam_RS26590     HYVDCFQVLLEGLVAEPQRPVGELPVLPADVRERVLRQWNDTGTPEAEGACLHALIEAQA
MxDZ2_Nan_RS16270     HYVDCFQVLLEGLVAEPQRPVGELPVLPADVRERVLRQWNDTGTPEAEGACLHALIEAQA
                      ************************************************************

MxDZ2_Kirby_RS0227310 LRTPDAVAIVVDDWELTYGELDQLSDRVAASLQDLDVGPEVVVGVYLDRSAELIVSLLAV
MxDZ2_Tam_RS26590     LRTPDAVAIVVDDWELTYGELDQLSDRVAASLQDLDVGPEVVVGVYLDRSAELIVSLLAV
MxDZ2_Nan_RS16270     LRTPDAVAIVVDDWELTYGELDQLSDRVAASLQDLDVGPEVVVGVYLDRSAELIVSLLAV
                      ************************************************************

MxDZ2_Kirby_RS0227310 MKAGGAFLALDADEPVDRLRHIVADARPRVVISSAKLSERLWGMGGFVTLHVDEGYRDMP
MxDZ2_Tam_RS26590     MKAGGAFLALDADEPVDRLRHIVADARPRVVISSAKLSERLWGMGGFVTLHVDEGYRDMP
MxDZ2_Nan_RS16270     MKAGGAFLALDADEPVDRLRHIVADARP--------------------------------
                      ****************************

MxDZ2_Kirby_RS0227310 AAPGQQLRRDVLPDHLAYILYTSGSTGRPKGTEITHRSIVNYLRWSVDAYRLREGTGSPV
MxDZ2_Tam_RS26590     AAPGQQLRRDVLPDHLAYILYTSGSTGRPKGTEITHRSIVNYLRWSVDAYRLREGTGSPV
MxDZ2_Nan_RS16270     ------------------------------------------------------------


MxDZ2_Kirby_RS0227310 IGSVSFDGTLTSLFAPLLAGRALFLVPRGHEIDQLTSRDYPEQGFSFIKMTPSHLRAFNG
MxDZ2_Tam_RS26590     IGSVSFDGTLTSLFAPLLAGRALFLVPRGHEIDQLTSRDYPEQGFSFIKMTPSHLRAFNG
MxDZ2_Nan_RS16270     ------------------------------------------------------------


MxDZ2_Kirby_RS0227310 LGRTREVLGRTHAVVLGGEGLHGVDLAPWREQGLPTRVINEYGPTEAAVACCFETLLPDG
MxDZ2_Tam_RS26590     LGRTREVLGRTHAVVLGGEGLHGVDLAPWREQGLPTRVINEYGPTEAAVACCFETLLPDG
MxDZ2_Nan_RS16270     ------------------------------------------------------------


MxDZ2_Kirby_RS0227310 TPPPERVPIGRPISHMRLYILDRYLQPVPVGVPGELYIGGVGLARGYLRRPDLTAERFVP
MxDZ2_Tam_RS26590     TPPPERVPIGRPISHMRLYILDRYLQPVPVGVPGELYIGGVGLARGYLRRPDLTAERFVP
MxDZ2_Nan_RS16270     ------------------------------------------------------------
                                                                                MxDZ2_Kirby_RS0227310 NPFDGGLEGSRLYRTGDHARYLSDGRIEYLGRQDDQLKIRGHRVETGEVEAALGRHPDVV
MxDZ2_Tam_RS26590     NPFDGGLEGSRLYRTGDHARYLSDGRIEYLGRQDDQLKIRGHRVETGEVEAALGRHPDVV
MxDZ2_Nan_RS16270     ------------------------------------------------------------


MxDZ2_Kirby_RS0227310 QAAVLLQRLPSGAPRLVAYVQPQSMENGDLRAELRKSLREVLPEYMMPEVIAVLPELPLT
MxDZ2_Tam_RS26590     QAAVLLQRLPSGAPRLVAYVQPQSMENGDLRAELRKSLREVLPEYMMPEVIAVLPELPLT
MxDZ2_Nan_RS16270     ------------------------------------------------------------


MxDZ2_Kirby_RS0227310 PSGKIDRKALPPVASEAPVASALARTEARTETERQLQALFGELLGLSSVAPTDSFFELGG
MxDZ2_Tam_RS26590     PSGKIDRKALPPVASEAPVASALARTEARTETERQLQALFGELLGLSSVAPTDSFFELGG
MxDZ2_Nan_RS16270     ------------------------------------------------------------


MxDZ2_Kirby_RS0227310 HSLLAVTLISRISAKLDIEVPLNEVFDRPSVESLAHWIDEHSSKVTALIRQLPACVVALK
MxDZ2_Tam_RS26590     HSLLAVTLISRISAKLDIEVPLNEVFDRPSVESLAHWIDEHSSKVTALIRQLPACVVALK
MxDZ2_Nan_RS16270     ------------------------------------------------------------


MxDZ2_Kirby_RS0227310 PLGHKPPLFLAPPSAGSPAVYVTLARHLDAEQPVFGFQMPGVMDDQLPPETIEDTAALYV
MxDZ2_Tam_RS26590     PLGHKPPLFLAPPSAGSPAVYVTLARHLDAEQPVFGFQMPGVMDDQLPPETIEDTAALYV
MxDZ2_Nan_RS16270     ------------------------------------------------------------


MxDZ2_Kirby_RS0227310 AALRQVQPRGPYRIAGWSYGGLVVCEMARQLEALGEQVALLGLIDGAALDRLSAHDGTPQ
MxDZ2_Tam_RS26590     AALRQVQPRGPYRIAGWSYGGLVVCEMARQLEALGEQVALLGLIDGAALDRLSAHDGTPQ
MxDZ2_Nan_RS16270     ---------------GW-------------------------------------------
                                     **

MxDZ2_Kirby_RS0227310 EIPAGSQLIKVLGEAQMPRDYASLRLIGEWMGISLPEAWADLLRRDTDGQYSYLWRFLRD
MxDZ2_Tam_RS26590     EIPAGSQLIKVLGEAQMPRDYASLRLIGEWMGISLPEAWADLLRRDTDGQYSYLWRFLRD
MxDZ2_Nan_RS16270     ------------------------------------------------------------


MxDZ2_Kirby_RS0227310 SALTARNFLATRRSEQLYTFSSYGGSATLFRTGPVTIGGDPLVDSVRRFALAGTEVIPVP
MxDZ2_Tam_RS26590     SALTARNFLATRRSEQLYTFSSYGGSATLFRTGPVTIGGDPLVDSVRRFALAGTEVIPVP
MxDZ2_Nan_RS16270     ------------------------------------------------------------


MxDZ2_Kirby_RS0227310 GNHMTLIMDERNVAVLAIEIQKCLDRALTLPPIAEMSRAEMPAKGLAAQLLMEVM
MxDZ2_Tam_RS26590     GNHMTLIMDERNVAVLAIEIQKCLDRALTLPPIAEMSRAEMPAKGLAAQLLMEVM
MxDZ2_Nan_RS16270     -------------------------------------------------------
```

**Premature stop codon followed by second ORF prediction 5**

| **Tam_location** | **Tam_nucleotide** | **Kirby_nucleotide** | **Nan_nucleotide** | **Nan_location** |
| --- | --- | --- | --- | --- |
| **5888709** | **C (complement)** | **C (complement)** | **.** | **5888681** |

```
multiple sequence alignment (nucleotide)

MxDZ2_Kirby_RS0206915 atgaagggctggaccctcttccctcacctcgtggttcgcaccacgggttttcccttcgat
MxDZ2_Tam_RS27885     atgaagggctggaccctcttccctcacctcgtggttcgcaccacgggttttcccttcgat
MxDZ2_Nan_RS14965     atgaagggctggaccctcttccctcacctcgtggttcgcaccacgggttttcccttcgat
                      ************************************************************

MxDZ2_Kirby_RS0206915 tggctggagcggctggggtgcccggaggccgcacgggccgcgcgccaactggcctcggcg
MxDZ2_Tam_RS27885     tggctggagcggctggggtgcccggaggccgcacgggccgcgcgccaactggcctcggcg
MxDZ2_Nan_RS14965     tggctggagcggctggggtgcccggaggccgcacgggccgcgcgccaactggcctcggcg
                      ************************************************************

MxDZ2_Kirby_RS0206915 cggcgggagctggaggcgctgaaagcccagggccctcgggtgaagcggccatcgcgcgcg
MxDZ2_Tam_RS27885     cggcgggagctggaggcgctgaaagcccagggccctcgggtgaagcggccatcgcgcgcg
MxDZ2_Nan_RS14965     cggcgggagctggaggcgctgaaagcccagggccctcgggtgaagcggccatcgcgcgcg
                      ************************************************************

MxDZ2_Kirby_RS0206915 gtgctgtcggccctgaaggccgggcgcccggtggacatcgagggaatggagtctcccgag
MxDZ2_Tam_RS27885     gtgctgtcggccctgaaggccgggcgcccggtggacatcgagggaatggagtctcccgag
MxDZ2_Nan_RS14965     gtgctgtcggccctgaaggccgggcgcccggtggacatcgagggaatggagtctcccgag
                      ************************************************************

MxDZ2_Kirby_RS0206915 ctgttcgctgaatggaacgaccgtgcccgggccgcgcaggaggcggaggccacctttcat
MxDZ2_Tam_RS27885     ctgttcgctgaatggaacgaccgtgcccgggccgcgcaggaggcggaggccacctttcat
MxDZ2_Nan_RS14965     ctgttcgctgaatggaacgaccgtgcccgggccgcgcaggaggcggaggccacctttcat
                      ************************************************************

MxDZ2_Kirby_RS0206915 gcggcgatgaagcaggagtccctggcggtggaggacgcgctgagggccctgcggcaggag
MxDZ2_Tam_RS27885     gcggcgatgaagcaggagtccctggcggtggaggacgcgctgagggccctgcggcaggag
MxDZ2_Nan_RS14965     gcggcgatgaagcaggagtccctggcggtggaggacgcgctgagggccctgcggcaggag
                      ************************************************************

MxDZ2_Kirby_RS0206915 ccgcgcttcctggaggcggtggccagctccagcccccccgtggcgagagacctgctggag
MxDZ2_Tam_RS27885     ccgcgcttcctggaggcggtggccagctccagcccccccgtggcgagagacctgctggag
MxDZ2_Nan_RS14965     ccgcgcttcctggaggcggtggccagctccagcccccccgtggcgagagacctgctggag
                      ************************************************************

MxDZ2_Kirby_RS0206915 gggcgagacggcgcgcggttgcggcggcaggtggccagctacctgcaacgcatgtgcgcg
MxDZ2_Tam_RS27885     gggcgagacggcgcgcggttgcggcggcaggtggccagctacctgcaacgcatgtgcgcg
MxDZ2_Nan_RS14965     gggcgagacggcgcgcggttgcggcggcaggtggccagctacctgcaacgcatgtgcgcg
                      ************************************************************

MxDZ2_Kirby_RS0206915 aagaacgagacgatgggcttcttcggtcccatcaactacggccgcgcggacgcggcggcg
MxDZ2_Tam_RS27885     aagaacgagacgatgggcttcttcggtcccatcaactacggccgcgcggacgcggcggcg
MxDZ2_Nan_RS14965     aagaacgagacgatgggcttcttcggtcccatcaactacggccgcgcggacgcggcggcg
                      ************************************************************

MxDZ2_Kirby_RS0206915 ccgacgggtgtgacgctgcgttggtccgggccggaggtgctcacggggcggaacaccttc
MxDZ2_Tam_RS27885     ccgacgggtgtgacgctgcgttggtccgggccggaggtgctcacggggcggaacaccttc
MxDZ2_Nan_RS14965     ccgacgggtgtgacgctgcgttggtccgggccggaggtgctcacggggcggaacaccttc
                      ************************************************************

MxDZ2_Kirby_RS0206915 gcggcgtcgtggctggtgcagggactggttcgcgccatcgccttcgaccccgaggtcgcc
MxDZ2_Tam_RS27885     gcggcgtcgtggctggtgcagggactggttcgcgccatcgccttcgaccccgaggtcgcc
MxDZ2_Nan_RS14965     gcggcgtcgtggctggtgcagggactggttcgcgccatcgccttcgaccccgaggtcgcc
                      ************************************************************

MxDZ2_Kirby_RS0206915 gcgtggctggtgctgcgtcgcaaggccttcgcggaggtgccatcgcgcaagacgctgccc
MxDZ2_Tam_RS27885     gcgtggctggtgctgcgtcgcaaggccttcgcggaggtgccatcgcgcaagacgctgccc
MxDZ2_Nan_RS14965     gcgtggctggtgctgcgtcgcaaggccttcgcggaggtgccatcgcgcaagacgctgccc
                      ************************************************************

MxDZ2_Kirby_RS0206915 gcgccggagagcgcggaggcgttgctgccccggctggtggaggcggtggatggaacgcgc
MxDZ2_Tam_RS27885     gcgccggagagcgcggaggcgttgctgccccggctggtggaggcggtggatggaacgcgc
MxDZ2_Nan_RS14965     gcgccggagagcgcggaggcgttgctgccccggctggtggaggcggtggatggaacgcgc
                      ************************************************************

MxDZ2_Kirby_RS0206915 acgctcgcGgggctcgcatcatcgctgggcgtggccccgggcatggcgcgcgaggctgcg
MxDZ2_Tam_RS27885     acgctcgcGgggctcgcatcatcgctgggcgtggccccgggcatggcgcgcgaggctgcg
MxDZ2_Nan_RS14965     acgctcgc-gggctcgcatcatcgctgggcgtggccccgggcatggcgcgcgaggctgcg
                      ******** ***************************************************

MxDZ2_Kirby_RS0206915 cggatgggctgcgaaaagggactgctgacgcatcagctcgaggtgcccgcggccacgcat
MxDZ2_Tam_RS27885     cggatgggctgcgaaaagggactgctgacgcatcagctcgaggtgcccgcggccacgcat
MxDZ2_Nan_RS14965     cggatgggctgcgaaaagggactgctga--------------------------------
                      ****************************


MxDZ2_Kirby_RS0206915 cacccggtggacgacctggccgagcgcgttgcgggcctgccgtgttccgcggcccggcgg
MxDZ2_Tam_RS27885     cacccggtggacgacctggccgagcgcgttgcgggcctgccgtgttccgcggcccggcgg
MxDZ2_Nan_RS14965     ------------------------------------------------------------


MxDZ2_Kirby_RS0206915 cacgtggagggcttgagcgcgctgctggcgttgatggccgggtatggcgccgcggatgcc
MxDZ2_Tam_RS27885     cacgtggagggcttgagcgcgctgctggcgttgatggccgggtatggcgccgcggatgcc
MxDZ2_Nan_RS14965     ------------------------------------------------------------


MxDZ2_Kirby_RS0206915 gcggggaagatggcgctccaggccgccttcgcgaagcgcgcgaatgagcagtggggcgtg
MxDZ2_Tam_RS27885     gcggggaagatggcgctccaggccgccttcgcgaagcgcgcgaatgagcagtggggcgtg
MxDZ2_Nan_RS14965     ------------------------------------------------------------


MxDZ2_Kirby_RS0206915 gcgccgccttccgagcgtgggcccgcgtcggagtcgcacaacttctaccaggaccggttg
MxDZ2_Tam_RS27885     gcgccgccttccgagcgtgggcccgcgtcggagtcgcacaacttctaccaggaccggttg
MxDZ2_Nan_RS14965     ------------------------------------------------------------


MxDZ2_Kirby_RS0206915 gcgctccgggaggagtgcggcggtgacctgcggctggaggccggaggtgagcgggcgcgt
MxDZ2_Tam_RS27885     gcgctccgggaggagtgcggcggtgacctgcggctggaggccggaggtgagcgggcgcgt
MxDZ2_Nan_RS14965     ------------------------------------------------------------


MxDZ2_Kirby_RS0206915 gagctggtgacgcgcctggagcccgcgctggcgtggatgggagcggcggcgcggcggacc
MxDZ2_Tam_RS27885     gagctggtgacgcgcctggagcccgcgctggcgtggatgggagcggcggcgcggcggacc
MxDZ2_Nan_RS14965     ------------------------------------------------------------


MxDZ2_Kirby_RS0206915 cgggaggccgctcgcaccaccgtggcggagctggtgggggcgcgcacggtgccgttctgg
MxDZ2_Tam_RS27885     cgggaggccgctcgcaccaccgtggcggagctggtgggggcgcgcacggtgccgttctgg
MxDZ2_Nan_RS14965     ------------------------------------------------------------


MxDZ2_Kirby_RS0206915 aaggtggcggccgcgtactcggaccggccggtgccgttggatggctcggtggcggaggtg
MxDZ2_Tam_RS27885     aaggtggcggccgcgtactcggaccggccggtgccgttggatggctcggtggcggaggtg
MxDZ2_Nan_RS14965     ------------------------------------------------------------


MxDZ2_Kirby_RS0206915 ctggctggcgcagtgggagacgcgtccgcgcgctgcgcggacctgggcttggtgacgccg
MxDZ2_Tam_RS27885     ctggctggcgcagtgggagacgcgtccgcgcgctgcgcggacctgggcttggtgacgccg
MxDZ2_Nan_RS14965     ------------------------------------------------------------


MxDZ2_Kirby_RS0206915 ccgtcgctggacggagacgcgaaggcgctttcgctcgtgacgtccatcgacctgctcgtg
MxDZ2_Tam_RS27885     ccgtcgctggacggagacgcgaaggcgctttcgctcgtgacgtccatcgacctgctcgtg
MxDZ2_Nan_RS14965     ------------------------------------------------------------


MxDZ2_Kirby_RS0206915 ggggcccgggacgtggaggcctggagccgaggtgagtacgagttggtgatgggcgacgtg
MxDZ2_Tam_RS27885     ggggcccgggacgtggaggcctggagccgaggtgagtacgagttggtgatgggcgacgtg
MxDZ2_Nan_RS14965     ------------------------------------------------------------


MxDZ2_Kirby_RS0206915 cacgacacggcgctggtgtggggctgggcgctccagttccacgaggcgcgcggccgggtg
MxDZ2_Tam_RS27885     cacgacacggcgctggtgtggggctgggcgctccagttccacgaggcgcgcggccgggtg
MxDZ2_Nan_RS14965     ------------------------------------------------------------


MxDZ2_Kirby_RS0206915 gagagcgccatggtgcgggcgctcggcgccttgagcaggccggtgcccctggtgacggtg
MxDZ2_Tam_RS27885     gagagcgccatggtgcgggcgctcggcgccttgagcaggccggtgcccctggtgacggtg
MxDZ2_Nan_RS14965     ------------------------------------------------------------


MxDZ2_Kirby_RS0206915 ttggcgtcgcggcgcacgggcctgcttccgtcggagttcccggggcccgtggtggagctg
MxDZ2_Tam_RS27885     ttggcgtcgcggcgcacgggcctgcttccgtcggagttcccggggcccgtggtggagctg
MxDZ2_Nan_RS14965     ------------------------------------------------------------


MxDZ2_Kirby_RS0206915 ggcggtgtcagtgcccgcgcctccgcgtggcggctgccgctggatgacctgttcgtggag
MxDZ2_Tam_RS27885     ggcggtgtcagtgcccgcgcctccgcgtggcggctgccgctggatgacctgttcgtggag
MxDZ2_Nan_RS14965     ------------------------------------------------------------


MxDZ2_Kirby_RS0206915 agtgatggaacgcgggcgcggctggtgtcgaagcggctgggctccgaggtgtgtctctac
MxDZ2_Tam_RS27885     agtgatggaacgcgggcgcggctggtgtcgaagcggctgggctccgaggtgtgtctctac
MxDZ2_Nan_RS14965     ------------------------------------------------------------


MxDZ2_Kirby_RS0206915 aacggtgagctggacagtctggtgcacaccgccttctccctgccgcgcatccgtccgctg
MxDZ2_Tam_RS27885     aacggtgagctggacagtctggtgcacaccgccttctccctgccgcgcatccgtccgctg
MxDZ2_Nan_RS14965     ------------------------------------------------------------


MxDZ2_Kirby_RS0206915 cgcgtgtcgctgggcgagcacacgcctcggttgaccctgggcggtgtcgtggtgcagcgc
MxDZ2_Tam_RS27885     cgcgtgtcgctgggcgagcacacgcctcggttgaccctgggcggtgtcgtggtgcagcgc
MxDZ2_Nan_RS14965     ------------------------------------------------------------


MxDZ2_Kirby_RS0206915 gagcagtggcggctgtcgcaggcggagcgggaggcgctgctcgcgggccgcgatgactcg
MxDZ2_Tam_RS27885     gagcagtggcggctgtcgcaggcggagcgggaggcgctgctcgcgggccgcgatgactcg
MxDZ2_Nan_RS14965     ------------------------------------------------------------


MxDZ2_Kirby_RS0206915 gcgagacttcgcgcggcggtgagcgtgtggagcgagcgcgggatgccggactgcgtcttc
MxDZ2_Tam_RS27885     gcgagacttcgcgcggcggtgagcgtgtggagcgagcgcgggatgccggactgcgtcttc
MxDZ2_Nan_RS14965     ------------------------------------------------------------


MxDZ2_Kirby_RS0206915 gcgaagttcaaggacgagcgaaagccggtgctggtggacgtgcgcagcccgccgctgctc
MxDZ2_Tam_RS27885     gcgaagttcaaggacgagcgaaagccggtgctggtggacgtgcgcagcccgccgctgctc
MxDZ2_Nan_RS14965     ------------------------------------------------------------


MxDZ2_Kirby_RS0206915 cgcgtgttcctcaacctgctggagcagaaggaggaggtcatcctgtcggagatgctgccc
MxDZ2_Tam_RS27885     cgcgtgttcctcaacctgctggagcagaaggaggaggtcatcctgtcggagatgctgccc
MxDZ2_Nan_RS14965     ------------------------------------------------------------


MxDZ2_Kirby_RS0206915 tcgccggaccagctctggctgcgaagtgcctcgggtgggcgccataccgtggagctccgt
MxDZ2_Tam_RS27885     tcgccggaccagctctggctgcgaagtgcctcgggtgggcgccataccgtggagctccgt
MxDZ2_Nan_RS14965     ------------------------------------------------------------


MxDZ2_Kirby_RS0206915 tgcacgctgatgtggggctcggagtccgctgccggagcgcgggaatga
MxDZ2_Tam_RS27885     tgcacgctgatgtggggctcggagtccgctgccggagcgcgggaatga
MxDZ2_Nan_RS14965     ------------------------------------------------


-----------------------------------------------------------------------------------
multiple sequence alignment (protein)


MxDZ2_Kirby_RS0206915 MKGWTLFPHLVVRTTGFPFDWLERLGCPEAARAARQLASARRELEALKAQGPRVKRPSRA
MxDZ2_Tam_RS27885     MKGWTLFPHLVVRTTGFPFDWLERLGCPEAARAARQLASARRELEALKAQGPRVKRPSRA
MxDZ2_Nan_RS14965     MKGWTLFPHLVVRTTGFPFDWLERLGCPEAARAARQLASARRELEALKAQGPRVKRPSRA
                      ************************************************************

MxDZ2_Kirby_RS0206915 VLSALKAGRPVDIEGMESPELFAEWNDRARAAQEAEATFHAAMKQESLAVEDALRALRQE
MxDZ2_Tam_RS27885     VLSALKAGRPVDIEGMESPELFAEWNDRARAAQEAEATFHAAMKQESLAVEDALRALRQE
MxDZ2_Nan_RS14965     VLSALKAGRPVDIEGMESPELFAEWNDRARAAQEAEATFHAAMKQESLAVEDALRALRQE
                      ************************************************************

MxDZ2_Kirby_RS0206915 PRFLEAVASSSPPVARDLLEGRDGARLRRQVASYLQRMCAKNETMGFFGPINYGRADAAA
MxDZ2_Tam_RS27885     PRFLEAVASSSPPVARDLLEGRDGARLRRQVASYLQRMCAKNETMGFFGPINYGRADAAA
MxDZ2_Nan_RS14965     PRFLEAVASSSPPVARDLLEGRDGARLRRQVASYLQRMCAKNETMGFFGPINYGRADAAA
                      ************************************************************

MxDZ2_Kirby_RS0206915 PTGVTLRWSGPEVLTGRNTFAASWLVQGLVRAIAFDPEVAAWLVLRRKAFAEVPSRKTLP
MxDZ2_Tam_RS27885     PTGVTLRWSGPEVLTGRNTFAASWLVQGLVRAIAFDPEVAAWLVLRRKAFAEVPSRKTLP
MxDZ2_Nan_RS14965     PTGVTLRWSGPEVLTGRNTFAASWLVQGLVRAIAFDPEVAAWLVLRRKAFAEVPSRKTLP
                      ************************************************************

MxDZ2_Kirby_RS0206915 APESAEALLPRLVEAVDGTRTLAGLASSLGVAPGMAREAARMGCEKGLLTHQLEVPAATH
MxDZ2_Tam_RS27885     APESAEALLPRLVEAVDGTRTLAGLASSLGVAPGMAREAARMGCEKGLLTHQLEVPAATH
MxDZ2_Nan_RS14965     APESAEALLPRLVEAVDGTRTLAG----------------------------------SH
                      ************************                                  :*

MxDZ2_Kirby_RS0206915 HPVDDLAERVAGLPCSAARRHVEGLSALLALMAGYGAADAAGKMALQAAFAKRANEQWGV
MxDZ2_Tam_RS27885     HPVDDLAERVAGLPCSAARRHVEGLSALLALMAGYGAADAAGKMALQAAFAKRANEQWGV
MxDZ2_Nan_RS14965     H-------------------------------------------------------RW--
                      *                                                       :*

MxDZ2_Kirby_RS0206915 APPSERGPASESHNFYQDRLALREECGGDLRLEAGGERARELVTRLEPALAWMGAAARRT
MxDZ2_Tam_RS27885     APPSERGPASESHNFYQDRLALREECGGDLRLEAGGERARELVTRLEPALAWMGAAARRT
MxDZ2_Nan_RS14965     ------------------------------------------------------------


MxDZ2_Kirby_RS0206915 REAARTTVAELVGARTVPFWKVAAAYSDRPVPLDGSVAEVLAGAVGDASARCADLGLVTP
MxDZ2_Tam_RS27885     REAARTTVAELVGARTVPFWKVAAAYSDRPVPLDGSVAEVLAGAVGDASARCADLGLVTP
MxDZ2_Nan_RS14965     ------------------------------------------------------------


MxDZ2_Kirby_RS0206915 PSLDGDAKALSLVTSIDLLVGARDVEAWSRGEYELVMGDVHDTALVWGWALQFHEARGRV
MxDZ2_Tam_RS27885     PSLDGDAKALSLVTSIDLLVGARDVEAWSRGEYELVMGDVHDTALVWGWALQFHEARGRV
MxDZ2_Nan_RS14965     --------------------------AWPR------------------------------
                                                **.*

MxDZ2_Kirby_RS0206915 ESAMVRALGALSRPVPLVTVLASRRTGLLPSEFPGPVVELGGVSARASAWRLPLDDLFVE
MxDZ2_Tam_RS27885     ESAMVRALGALSRPVPLVTVLASRRTGLLPSEFPGPVVELGGVSARASAWRLPLDDLFVE
MxDZ2_Nan_RS14965     ------------------------------------------------AWR---------
                                                                      ***

MxDZ2_Kirby_RS0206915 SDGTRARLVSKRLGSEVCLYNGELDSLVHTAFSLPRIRPLRVSLGEHTPRLTLGGVVVQR
MxDZ2_Tam_RS27885     SDGTRARLVSKRLGSEVCLYNGELDSLVHTAFSLPRIRPLRVSLGEHTPRLTLGGVVVQR
MxDZ2_Nan_RS14965     ------------------------------------------------------------


MxDZ2_Kirby_RS0206915 EQWRLSQAEREALLAGRDDSARLRAAVSVWSERGMPDCVFAKFKDERKPVLVDVRSPPLL
MxDZ2_Tam_RS27885     EQWRLSQAEREALLAGRDDSARLRAAVSVWSERGMPDCVFAKFKDERKPVLVDVRSPPLL
MxDZ2_Nan_RS14965     --------------------ARLRG----WAAK--RDC----------------------
                                          ****.    *: :   **

MxDZ2_Kirby_RS0206915 RVFLNLLEQKEEVILSEMLPSPDQLWLRSASGGRHTVELRCTLMWGSESAAGARE
MxDZ2_Tam_RS27885     RVFLNLLEQKEEVILSEMLPSPDQLWLRSASGGRHTVELRCTLMWGSESAAGARE
MxDZ2_Nan_RS14965     -------------------------------------------------------
```

**Premature stop codon followed by second ORF prediction 6**

| **Tam_location** | **Tam_nucleotide** | **Kirby_nucleotide** | **Nan_nucleotide** | **Nan_location** |
| --- | --- | --- | --- | --- |
| **7592018** | **C** | **C** | **.** | **7591988** |

```
multiple sequence alignment (nucleotide)

MxDZ2_Kirby_RS0231740 atgcctgagcggctgtacccgcacgaagtcctgctggcggcgttcggcatcgtcctctcc
MxDZ2_Tam_RS34805     atgcctgagcggctgtacccgcacgaagtcctgctggcggcgttcggcatcgtcctctcc
MxDZ2_Nan_RS08040     atgcctgagcggctgtacccgcacgaagtcctgctggcggcgttcggcatcgtcctctcc
                      ************************************************************

MxDZ2_Kirby_RS0231740 acagcgctggtgttcgtcgcgggaatctccgcgaccgtcaccgcgcaagccgttggtggc
MxDZ2_Tam_RS34805     acagcgctggtgttcgtcgcgggaatctccgcgaccgtcaccgcgcaagccgttggtggc
MxDZ2_Nan_RS08040     acagcgctggtgttcgtcgcgggaatctccgcgaccgtcaccgcgcaagccgttggtggc
                      ************************************************************

MxDZ2_Kirby_RS0231740 acggtgctgttcatgggcagcgtggcgctgctggcgcgactggacgcagtcgcccatgtc
MxDZ2_Tam_RS34805     acggtgctgttcatgggcagcgtggcgctgctggcgcgactggacgcagtcgcccatgtc
MxDZ2_Nan_RS08040     acggtgctgttcatgggcagcgtggcgctgctggcgcgactggacgcagtcgcccatgtc
                      ************************************************************

MxDZ2_Kirby_RS0231740 ttccgggcacggctgctgctcgcctacctcgcgacgttcttcttctatgcctccgtgaag
MxDZ2_Tam_RS34805     ttccgggcacggctgctgctcgcctacctcgcgacgttcttcttctatgcctccgtgaag
MxDZ2_Nan_RS08040     ttccgggcacggctgctgctcgcctacctcgcgacgttcttcttctatgcctccgtgaag
                      ************************************************************

MxDZ2_Kirby_RS0231740 caggccgtccccgcgctggggctcgtgacgcgtgatgcgtggctcctgtccgctgacgtg
MxDZ2_Tam_RS34805     caggccgtccccgcgctggggctcgtgacgcgtgatgcgtggctcctgtccgctgacgtg
MxDZ2_Nan_RS08040     caggccgtccccgcgctggggctcgtgacgcgtgatgcgtggctcctgtccgctgacgtg
                      ************************************************************

MxDZ2_Kirby_RS0231740 ttcctgttcggcatcacaccggccgcatggctccagcggtggagcacgccctgggtcaac
MxDZ2_Tam_RS34805     ttcctgttcggcatcacaccggccgcatggctccagcggtggagcacgccctgggtcaac
MxDZ2_Nan_RS08040     ttcctgttcggcatcacaccggccgcatggctccagcggtggagcacgccctgggtcaac
                      ************************************************************

MxDZ2_Kirby_RS0231740 gagctcttcagtgccagctatctagcgttccacggatacctgcacctcgccatggcctgg
MxDZ2_Tam_RS34805     gagctcttcagtgccagctatctagcgttccacggatacctgcacctcgccatggcctgg
MxDZ2_Nan_RS08040     gagctcttcagtgccagctatctagcgttccacggatacctgcacctcgccatggcctgg
                      ************************************************************

MxDZ2_Kirby_RS0231740 gcgctggtggggccgcgtgcgcgagcggagcgcttcttcatcgacgtcttctccgcctat
MxDZ2_Tam_RS34805     gcgctggtggggccgcgtgcgcgagcggagcgcttcttcatcgacgtcttctccgcctat
MxDZ2_Nan_RS08040     gcgctggtggggccgcgtgcgcgagcggagcgcttcttcatcgacgtcttctccgcctat
                      ************************************************************

MxDZ2_Kirby_RS0231740 ctcccgggcatcgccggctactacctgatgcccgccatcggtcccgtggccgcctggccg
MxDZ2_Tam_RS34805     ctcccgggcatcgccggctactacctgatgcccgccatcggtcccgtggccgcctggccg
MxDZ2_Nan_RS08040     ctcccgggcatcgccggctactacctgatgcccgccatcggtcccgtggccgcctggccg
                      ************************************************************

MxDZ2_Kirby_RS0231740 gagctgttcaccgtccccatcgagggtggctggttcacgcaactgaatgccgccgtcgtg
MxDZ2_Tam_RS34805     gagctgttcaccgtccccatcgagggtggctggttcacgcaactgaatgccgccgtcgtg
MxDZ2_Nan_RS08040     gagctgttcaccgtccccatcgagggtggctggttcacgcaactgaatgccgccgtcgtg
                      ************************************************************

MxDZ2_Kirby_RS0231740 gcgagtggttcgtccacctacgacctcttccccagcctgcacacgtacatcacgctggtg
MxDZ2_Tam_RS34805     gcgagtggttcgtccacctacgacctcttccccagcctgcacacgtacatcacgctggtg
MxDZ2_Nan_RS08040     gcgagtggttcgtccacctacgacctcttccccagcctgcacacgtacatcacgctggtg
                      ************************************************************

MxDZ2_Kirby_RS0231740 ctgctcgaacacgaccggcgccagcacccgtggcggttccggctgatggtgcccgtcgcc
MxDZ2_Tam_RS34805     ctgctcgaacacgaccggcgccagcacccgtggcggttccggctgatggtgcccgtcgcc
MxDZ2_Nan_RS08040     ctgctcgaacacgaccggcgccagcacccgtggcggttccggctgatggtgcccgtcgcc
                      ************************************************************

MxDZ2_Kirby_RS0231740 gtggccatcctcatgtccacgctggtgctgcgttatcactatgccgtggacctgctggcg
MxDZ2_Tam_RS34805     gtggccatcctcatgtccacgctggtgctgcgttatcactatgccgtggacctgctggcg
MxDZ2_Nan_RS08040     gtggccatcctcatgtccacgctggtgctgcgttatcactatgccgtggacctgctggcg
                      ************************************************************

MxDZ2_Kirby_RS0231740 ggcgccgtgtggttcatggtgtttcgcgcctgcttCccccggcttcaggcgcgctgggag
MxDZ2_Tam_RS34805     ggcgccgtgtggttcatggtgtttcgcgcctgcttCccccggcttcaggcgcgctgggag
MxDZ2_Nan_RS08040     ggcgccgtgtggttcatggtgtttcgcgcctgctt-ccccggcttcaggcgcgctgggag
                      *********************************** ************************

MxDZ2_Kirby_RS0231740 gcgcggacggcgctatcgaagcacgaagcgccgacggtgggcgtcgggtga---------
MxDZ2_Tam_RS34805     gcgcggacggcgctatcgaagcacgaagcgccgacggtgggcgtcgggtga---------
MxDZ2_Nan_RS08040     gcgcggacggcgctatcgaagcacgaagcgccgacggtgggcgtcgggtgacagtccggt
                      ***************************************************

MxDZ2_Kirby_RS0231740 ------------------------------------------------------------
MxDZ2_Tam_RS34805     ------------------------------------------------------------
MxDZ2_Nan_RS08040     gcgcgccttgaaggaggaatggaccgtggtcatccgctacacgcatgacgtggagaagac


MxDZ2_Kirby_RS0231740 ------------------------------------------------------------
MxDZ2_Tam_RS34805     ------------------------------------------------------------
MxDZ2_Nan_RS08040     gtccgagggctggcgcatccgccgcgtcatgctcgccccatccactaccggggcaactct


MxDZ2_Kirby_RS0231740 -------------------
MxDZ2_Tam_RS34805     -------------------
MxDZ2_Nan_RS08040     gtggggctggagtgcgtga


-----------------------------------------------------------------------------------
multiple sequence alignment (protein)


MxDZ2_Kirby_RS0231740 MPERLYPHEVLLAAFGIVLSTALVFVAGISATVTAQAVGGTVLFMGSVALLARLDAVAHV
MxDZ2_Tam_RS34805     MPERLYPHEVLLAAFGIVLSTALVFVAGISATVTAQAVGGTVLFMGSVALLARLDAVAHV
MxDZ2_Nan_RS08040     MPERLYPHEVLLAAFGIVLSTALVFVAGISATVTAQAVGGTVLFMGSVALLARLDAVAHV
                      ************************************************************

MxDZ2_Kirby_RS0231740 FRARLLLAYLATFFFYASVKQAVPALGLVTRDAWLLSADVFLFGITPAAWLQRWSTPWVN
MxDZ2_Tam_RS34805     FRARLLLAYLATFFFYASVKQAVPALGLVTRDAWLLSADVFLFGITPAAWLQRWSTPWVN
MxDZ2_Nan_RS08040     FRARLLLAYLATFFFYASVKQAVPALGLVTRDAWLLSADVFLFGITPAAWLQRWSTPWVN
                      ************************************************************

MxDZ2_Kirby_RS0231740 ELFSASYLAFHGYLHLAMAWALVGPRARAERFFIDVFSAYLPGIAGYYLMPAIGPVAAWP
MxDZ2_Tam_RS34805     ELFSASYLAFHGYLHLAMAWALVGPRARAERFFIDVFSAYLPGIAGYYLMPAIGPVAAWP
MxDZ2_Nan_RS08040     ELFSASYLAFHGYLHLAMAWALVGPRARAERFFIDVFSAYLPGIAGYYLMPAIGPVAAWP
                      ************************************************************

MxDZ2_Kirby_RS0231740 ELFTVPIEGGWFTQLNAAVVASGSSTYDLFPSLHTYITLVLLEHDRRQHPWRFRLMVPVA
MxDZ2_Tam_RS34805     ELFTVPIEGGWFTQLNAAVVASGSSTYDLFPSLHTYITLVLLEHDRRQHPWRFRLMVPVA
MxDZ2_Nan_RS08040     ELFTVPIEGGWFTQLNAAVVASGSSTYDLFPSLHTYITLVLLEHDRRQHPWRFRLMVPVA
                      ************************************************************


MxDZ2_Kirby_RS0231740 VAILMSTLVLRYHYAVDLLAGAVWFMVFRACFP--RLQARWEARTALSKHEAPTVG----
MxDZ2_Tam_RS34805     VAILMSTLVLRYHYAVDLLAGAVWFMVFRACFP--RLQARWEARTALSKHEAPTVG----
MxDZ2_Nan_RS08040     VAILMSTLVLRYHYAVDLLAGAVWFMVFRACFPGFRRAGRRGRRYRSTKRRRWASGDSPV
                      *********************************  *  .*   *   :*:.  : *

MxDZ2_Kirby_RS0231740 -------------------------------------------VG
MxDZ2_Tam_RS34805     -------------------------------------------VG
MxDZ2_Nan_RS08040     RALKEEWTVVIRYTHDVEKTSEGWRIRRVMLAPSTTGATLWGWSA
                                                                  .
```

**Premature stop codon followed by second ORF prediction 7**

| **Tam_location** | **Tam_nucleotide** | **Kirby_nucleotide** | **Nan_nucleotide** | **Nan_location** |
| --- | --- | --- | --- | --- |
| **8820990** | **C** | **C** | **A** | **8820959** |

```
multiple sequence alignment (nucleotide)

MxDZ2_Tam_RS02175     atgagaaaggtgttgttcctcggggcctttgccctgggcgcctcggcctgtactcccgat
MxDZ2_Kirby_RS0212590 atgagaaaggtgttgttcctcggggcctttgccctgggcgcctcggcctgtactcccgat
MxDZ2_Nan_RS02810     atgagaaaggtgttgttcctcggggcctttgccctgggcgcctcggcctgtactcccgat
                      ************************************************************

MxDZ2_Tam_RS02175     atcgcccaggacccgccgcggagccccgaagactccgtcgtcgcggagttcgatccaacc
MxDZ2_Kirby_RS0212590 atcgcccaggacccgccgcggagccccgaagactccgtcgtcgcggagttcgatccaacc
MxDZ2_Nan_RS02810     atcgcccaggacccgccgcggagccccgaagactccgtcgtcgcggagttcgatccaacc
                      ************************************************************

MxDZ2_Tam_RS02175     gggtctccacccgtcgtcccgtcgcccaacgacctggccattgtcaacggtctggtcaac
MxDZ2_Kirby_RS0212590 gggtctccacccgtcgtcccgtcgcccaacgacctggccattgtcaacggtctggtcaac
MxDZ2_Nan_RS02810     gggtctccacccgtcgtcccgtcgcccaacgacctggccattgtcaacggtctggtcaac
                      ************************************************************

MxDZ2_Tam_RS02175     gcgcccatcaacccgcaggcgccggcggcggagcaggagttcacgcgcgactacgtgaac
MxDZ2_Kirby_RS0212590 gcgcccatcaacccgcaggcgccggcggcggagcaggagttcacgcgcgactacgtgaac
MxDZ2_Nan_RS02810     gcgcccatcaacccgcaggcgccggcggcggagcaggagttcacgcgcgactacgtgaac
                      ************************************************************

MxDZ2_Tam_RS02175     acgctcaacggcttcccgaccacggtcaccgcgaccacccgcatccggggcctggaggcg
MxDZ2_Kirby_RS0212590 acgctcaacggcttcccgaccacggtcaccgcgaccacccgcatccggggcctggaggcg
MxDZ2_Nan_RS02810     acgctcaacggcttcccgaccacggtcaccgcgaccacccgcatccggggcctggaggcg
                      ************************************************************

MxDZ2_Tam_RS02175     ggaacggtcaacacgaacacggtgaagatcatcgacgtgtacgcgggtacgccgctggcg
MxDZ2_Kirby_RS0212590 ggaacggtcaacacgaacacggtgaagatcatcgacgtgtacgcgggtacgccgctggcg
MxDZ2_Nan_RS02810     ggaacggtcaacacgaacacggtgaagatcatcgacgtgtacgcgggtacgccgctggcg
                      ************************************************************

MxDZ2_Tam_RS02175     aagccggcgacgcccagctacatcgggtacaacgaggaaaccaacctcgtcaccatcatc
MxDZ2_Kirby_RS0212590 aagccggcgacgcccagctacatcgggtacaacgaggaaaccaacctcgtcaccatcatc
MxDZ2_Nan_RS02810     aagccggcgacgcccagctacatcgggtacaacgaggaaaccaacctcgtcaccatcatc
                      ************************************************************

MxDZ2_Tam_RS02175     cccccgctccccggaggctggcccaagggtggccgctacgcggtggccatcatcggcggt
MxDZ2_Kirby_RS0212590 cccccgctccccggaggctggcccaagggtggccgctacgcggtggccatcatcggcggt
MxDZ2_Nan_RS02810     cccccgctccccggaggctggcccaagggtggccgctacgcggtggccatcatcggcggt
                      ************************************************************

MxDZ2_Tam_RS02175     gagaacggcgtgaagacgacggccggcaagccggtcatcccgtccgccacgtgggccttc
MxDZ2_Kirby_RS0212590 gagaacggcgtgaagacgacggccggcaagccggtcatcccgtccgccacgtgggccttc
MxDZ2_Nan_RS02810     gagaacggcgtgaagacgacggccggcaagccggtcatcccgtccgccacgtgggccttc
                      ************************************************************

MxDZ2_Tam_RS02175     gccagctccgccgagtcgctggtcacctgcgacgacctggcggcgcccaactgccagacg
MxDZ2_Kirby_RS0212590 gccagctccgccgagtcgctggtcacctgcgacgacctggcggcgcccaactgccagacg
MxDZ2_Nan_RS02810     gccagctccgccgagtcgctggtcacctgcgacgacctggcggcgcccaactgccagacg
                      ************************************************************

MxDZ2_Tam_RS02175     accaccgagctcatcccctccacggtgacggagcccgccgcccgcctggcggaccagacg
MxDZ2_Kirby_RS0212590 accaccgagctcatcccctccacggtgacggagcccgccgcccgcctggcggaccagacg
MxDZ2_Nan_RS02810     accaccgagctcatcccctccacggtgacggagcccgccgcccgcctggcggaccagacg
                      ************************************************************

MxDZ2_Tam_RS02175     gccagcgccctgcggctggagcagctgcgccgcggctacgcgcccatcctgaagctggtg
MxDZ2_Kirby_RS0212590 gccagcgccctgcggctggagcagctgcgccgcggctacgcgcccatcctgaagctggtg
MxDZ2_Nan_RS02810     gccagcgccctgcggctggagcagctgcgccgcggctacgcgcccatcctgaagctggtg
                      ************************************************************

MxDZ2_Tam_RS02175     tccgagcagttcagcgtgaagcgcgaggacatcgtcctcctgtggaccttctccatcatg
MxDZ2_Kirby_RS0212590 tccgagcagttcagcgtgaagcgcgaggacatcgtcctcctgtggaccttctccatcatg
MxDZ2_Nan_RS02810     tccgagcagttcagcgtgaagcgcgaggacatcgtcctcctgtggaccttctccatcatg
                      ************************************************************

MxDZ2_Tam_RS02175     gaccacccggaggccaccttcgatccgggcaacggcgtcgtccccttccccaatgacctg
MxDZ2_Kirby_RS0212590 gaccacccggaggccaccttcgatccgggcaacggcgtcgtccccttccccaatgacctg
MxDZ2_Nan_RS02810     gaccacccggaggccaccttcgatccgggcaacggcgtcgtccccttccccaatgacctg
                      ************************************************************

MxDZ2_Tam_RS02175     ctgcgcaaccccgaaaccgggctgctggccctgccggagcccaccgaggccggccccgcg
MxDZ2_Kirby_RS0212590 ctgcgcaaccccgaaaccgggctgctggccctgccggagcccaccgaggccggccccgcg
MxDZ2_Nan_RS02810     ctgcgcaaccccgaaaccgggctgctggccctgccggagcccaccgaggccggccccgcg
                      ************************************************************


MxDZ2_Tam_RS02175     cagatgctcatcaagggcctcaacaccctggacggctggtccaccaccgcgcccatcgtg
MxDZ2_Kirby_RS0212590 cagatgctcatcaagggcctcaacaccctggacggctggtccaccaccgcgcccatcgtg
MxDZ2_Nan_RS02810     cagatgctcatcaagggcctcaacaccctggacggctggtccaccaccgcgcccatcgtg
                      ************************************************************

MxDZ2_Tam_RS02175     tcggagaacggcgcgaaacgcggccccatcgacacgggttccctgctggacgcgaacacc
MxDZ2_Kirby_RS0212590 tcggagaacggcgcgaaacgcggccccatcgacacgggttccctgctggacgcgaacacc
MxDZ2_Nan_RS02810     tcggagaacggcgcgaaacgcggccccatcgacacgggttccctgctggacgcgaacacc
                      ************************************************************

MxDZ2_Tam_RS02175     gtgcgcatgaagatcggcttcacgccggacaccgaggaggagcagaagaccacgctgcgc
MxDZ2_Kirby_RS0212590 gtgcgcatgaagatcggcttcacgccggacaccgaggaggagcagaagaccacgctgcgc
MxDZ2_Nan_RS02810     gtgcgcatgaagatcggcttcacgccggacaccgaggaggagcagaagaccacgctgcgc
                      ************************************************************

MxDZ2_Tam_RS02175     ttcgtcaagctgaccaaccccacctcgggcaccaacccccaggtgaagctgtgcttcaac
MxDZ2_Kirby_RS0212590 ttcgtcaagctgaccaaccccacctcgggcaccaacccccaggtgaagctgtgcttcaac
MxDZ2_Nan_RS02810     ttcgtcaagctgaccaaccccacctcgggcaccaacccccaggtgaagctgtgcttcaac
                      ************************************************************

MxDZ2_Tam_RS02175     tgcgagcccgtcgtggagggcctgacgccggcgccgaactcgccgcagcagctccagatt
MxDZ2_Kirby_RS0212590 tgcgagcccgtcgtggagggcctgacgccggcgccgaactcgccgcagcagctccagatt
MxDZ2_Nan_RS02810     tgcgagcccgtcgtggagggcctgacgccggcgccgaactcgccgcagcagctccagatt
                      ************************************************************

MxDZ2_Tam_RS02175     gtcccggaggtgccgctggacgaggccacccagtatggcgtggtgatgctgcgcggcatg
MxDZ2_Kirby_RS0212590 gtcccggaggtgccgctggacgaggccacccagtatggcgtggtgatgctgcgcggcatg
MxDZ2_Nan_RS02810     gtcccggaggtgccgctggacgaggccacccagtatggcgtggtgatgctgcgcggcatg
                      ************************************************************

MxDZ2_Tam_RS02175     aaggacacgctgggccgcaccgtggcgcccaccgccgcgcaggcgctgatgcgcatggtc
MxDZ2_Kirby_RS0212590 aaggacacgctgggccgcaccgtggcgcccaccgccgcgcaggcgctgatgcgcatggtc
MxDZ2_Nan_RS02810     aaggacacgctgggccgcaccgtggcgcccaccgccgcgcaggcgctgatgcgcatggtc
                      ************************************************************

MxDZ2_Tam_RS02175     aacccgctggtggacgccaacggcaagagccaggtggcggccgttcctgacacgctcgcc
MxDZ2_Kirby_RS0212590 aacccgctggtggacgccaacggcaagagccaggtggcggccgttcctgacacgctcgcc
MxDZ2_Nan_RS02810     aacccgctggtggacgccaacggcaagagccaggtggcggccgttcctgacacgctcgcc
                      ************************************************************


MxDZ2_Tam_RS02175     cgggacctggagcgggcgcgtgtgggcatgaagccgctgttcgacggcctggaggccgcg
MxDZ2_Kirby_RS0212590 cgggacctggagcgggcgcgtgtgggcatgaagccgctgttcgacggcctggaggccgcg
MxDZ2_Nan_RS02810     cgggacctggagcgggcgcgtgtgggcatgaagccgctgttcgacggcctggaggccgcg
                      ************************************************************

MxDZ2_Tam_RS02175     ggcatcgcgcgcaaggacatcaacctggcgtgggccttcaccacccagagcacccgttcc
MxDZ2_Kirby_RS0212590 ggcatcgcgcgcaaggacatcaacctggcgtgggccttcaccacccagagcacccgttcc
MxDZ2_Nan_RS02810     ggcatcgcgcgcaaggacatcaacctggcgtgggccttcaccacccagagcacccgttcc
                      ************************************************************

MxDZ2_Tam_RS02175     atcctcgagcggctgaacgcggcccccacgcaggtgccggtgccggccgaccccgtctac
MxDZ2_Kirby_RS0212590 atcctcgagcggctgaacgcggcccccacgcaggtgccggtgccggccgaccccgtctac
MxDZ2_Nan_RS02810     atcctcgagcggctgaacgcggcccccacgcaggtgccggtgccggccgaccccgtctac
                      ************************************************************

MxDZ2_Tam_RS02175     ctgctggaccagacggccaccatcaagggcacgatggccggcctgacgctggacaacgac
MxDZ2_Kirby_RS0212590 ctgctggaccagacggccaccatcaagggcacgatggccggcctgacgctggacaacgac
MxDZ2_Nan_RS02810     ctgctggaccagacggccaccatcaagggcacgatggccggcctgacgctggacaacgac
                      ************************************************************

MxDZ2_Tam_RS02175     gcggtgggccgcgtcttcatcggcgcgtaccactcgcccttcttcctggatgacgcgcag
MxDZ2_Kirby_RS0212590 gcggtgggccgcgtcttcatcggcgcgtaccactcgcccttcttcctggatgacgcgcag
MxDZ2_Nan_RS02810     gcggtgggccgcgtcttcatcggcgcgtaccactcgcccttcttcctggatgacgcgcag
                      ************************************************************

MxDZ2_Tam_RS02175     ggcacgctcaaccgcgccacgccgcgcatcgaccgcgtcccgttcatgctcttcacgccc
MxDZ2_Kirby_RS0212590 ggcacgctcaaccgcgccacgccgcgcatcgaccgcgtcccgttcatgctcttcacgccc
MxDZ2_Nan_RS02810     ggcacgctcaaccgcgccacgccgcgcatcgaccgcgtcccgttcatgctcttcacgccc
                      ************************************************************

MxDZ2_Tam_RS02175     gcgggggcggcgcccgcggacggctacccggtggtcatctacggccatgggctgacgggc
MxDZ2_Kirby_RS0212590 gcgggggcggcgcccgcggacggctacccggtggtcatctacggccatgggctgacgggc
MxDZ2_Nan_RS02810     gcgggggcggcgcccgcggacggctacccggtggtcatctacggccatgggctgacgggc
                      ************************************************************

MxDZ2_Tam_RS02175     aaccgcaccaacatcttgacggtggccaacaacttcaacgcggcgggctacgccgtggcg
MxDZ2_Kirby_RS0212590 aaccgcaccaacatcttgacggtggccaacaacttcaacgcggcgggctacgccgtggcg
MxDZ2_Nan_RS02810     aaccgcaccaacatcttgacggtggccaacaacttcaacgcggcgggctacgccgtggcg
                      ************************************************************


MxDZ2_Tam_RS02175     gccatcgacaccgtgttccacggtgagcgcgccagctgcgccggcatcagcgaggacgcc
MxDZ2_Kirby_RS0212590 gccatcgacaccgtgttccacggtgagcgcgccagctgcgccggcatcagcgaggacgcc
MxDZ2_Nan_RS02810     gccatcgacaccgtgttccacggtgagcgcgccagctgcgccggcatcagcgaggacgcc
                      ************************************************************

MxDZ2_Tam_RS02175     ccggttgtcgtcgacgagaaccgcgatggcgtccccgagctcaccatcaccacgccggac
MxDZ2_Kirby_RS0212590 ccggttgtcgtcgacgagaaccgcgatggcgtccccgagctcaccatcaccacgccggac
MxDZ2_Nan_RS02810     ccggttgtcgtcgacgagaaccgcgatggcgtccccgagctcaccatcaccacgccggac
                      ************************************************************

MxDZ2_Tam_RS02175     gcggcctgCgacgacggcgacacctgcgacgtgacggcgggcagccccaccttcggccgc
MxDZ2_Kirby_RS0212590 gcggcctgCgacgacggcgacacctgcgacgtgacggcgggcagccccaccttcggccgc
MxDZ2_Nan_RS02810     gcggcctgA---------------------------------------------------
                      ********

MxDZ2_Tam_RS02175     tgcgtgtcgaacatccagccggcctgcgacgtggggccgaccgcgtcgccgcacggtgac
MxDZ2_Kirby_RS0212590 tgcgtgtcgaacatccagccggcctgcgacgtggggccgaccgcgtcgccgcacggtgac
MxDZ2_Nan_RS02810     ------------------------------------------------------------


MxDZ2_Tam_RS02175     ctgttctgctccagccagggccggggccgctgcgtcgtgaccgagcccgacagcaccacc
MxDZ2_Kirby_RS0212590 ctgttctgctccagccagggccggggccgctgcgtcgtgaccgagcccgacagcaccacc
MxDZ2_Nan_RS02810     ------------------------------------------------------------


MxDZ2_Tam_RS02175     ggcacctgcgagggcggcgacttcgcccgtccgggtcccaatcagccgccgtacgtctcc
MxDZ2_Kirby_RS0212590 ggcacctgcgagggcggcgacttcgcccgtccgggtcccaatcagccgccgtacgtctcc
MxDZ2_Nan_RS02810     ------------------------------------------------------------


MxDZ2_Tam_RS02175     ggcgcgggcttcctgaacctggtcaacctgttcgccacccgcgacaacttccgccaccac
MxDZ2_Kirby_RS0212590 ggcgcgggcttcctgaacctggtcaacctgttcgccacccgcgacaacttccgccaccac
MxDZ2_Nan_RS02810     ------------------------------------------------------------


MxDZ2_Tam_RS02175     gtggtggacttcgcccagctggcgcgcgcgctggacaccgacaccctcaacggccgcctg
MxDZ2_Kirby_RS0212590 gtggtggacttcgcccagctggcgcgcgcgctggacaccgacaccctcaacggccgcctg
MxDZ2_Nan_RS02810     ------------------------------------------------------------


MxDZ2_Tam_RS02175     gctgccgcgggcgccggtacgctgaacaccagccagttcagctacgtgggccagagcctg
MxDZ2_Kirby_RS0212590 gctgccgcgggcgccggtacgctgaacaccagccagttcagctacgtgggccagagcctg
MxDZ2_Nan_RS02810     ------------------------------------------------------------


MxDZ2_Tam_RS02175     ggcggcatgcagggcgtcctgtccgcctccgtgtcccccgtcgtgggccgcacagcactc
MxDZ2_Kirby_RS0212590 ggcggcatgcagggcgtcctgtccgcctccgtgtcccccgtcgtgggccgcacagcactc
MxDZ2_Nan_RS02810     ------------------------------------------------------------


MxDZ2_Tam_RS02175     aacgcggtgggcggtgacctggtggacatcctcctgacggccacggactcgacgttcgtg
MxDZ2_Kirby_RS0212590 aacgcggtgggcggtgacctggtggacatcctcctgacggccacggactcgacgttcgtg
MxDZ2_Nan_RS02810     ------------------------------------------------------------


MxDZ2_Tam_RS02175     cggtaccgctcgggcttcttcgaaaccctggaaaacagcggccgcgccccgggcacgccg
MxDZ2_Kirby_RS0212590 cggtaccgctcgggcttcttcgaaaccctggaaaacagcggccgcgccccgggcacgccg
MxDZ2_Nan_RS02810     ------------------------------------------------------------


MxDZ2_Tam_RS02175     gagttctacgagttcatcaccctggcgcgcaccatcctggacccggcctcgccccgcaac
MxDZ2_Kirby_RS0212590 gagttctacgagttcatcaccctggcgcgcaccatcctggacccggcctcgccccgcaac
MxDZ2_Nan_RS02810     ------------------------------------------------------------


MxDZ2_Tam_RS02175     tacggcaggtacctggagaacgcgccgtccgcgcccgccaaccgcaatgcgttcatccag
MxDZ2_Kirby_RS0212590 tacggcaggtacctggagaacgcgccgtccgcgcccgccaaccgcaatgcgttcatccag
MxDZ2_Nan_RS02810     ------------------------------------------------------------


MxDZ2_Tam_RS02175     tacatcgagggtgacatggtcatccccaacgcggtgaccgagggcctcatcagcgcggcc
MxDZ2_Kirby_RS0212590 tacatcgagggtgacatggtcatccccaacgcggtgaccgagggcctcatcagcgcggcc
MxDZ2_Nan_RS02810     ------------------------------------------------------------


MxDZ2_Tam_RS02175     aaccgcccgctggtcggcgtctccgagccccgcacggtgcagacgtacctggcggaggac
MxDZ2_Kirby_RS0212590 aaccgcccgctggtcggcgtctccgagccccgcacggtgcagacgtacctggcggaggac
MxDZ2_Nan_RS02810     ------------------------------------------------------------


MxDZ2_Tam_RS02175     gggatgggcctgccgacgaacgtgcgtcacgcgttcttcggcatcgtggacccgtcgctg
MxDZ2_Kirby_RS0212590 gggatgggcctgccgacgaacgtgcgtcacgcgttcttcggcatcgtggacccgtcgctg
MxDZ2_Nan_RS02810     ------------------------------------------------------------


MxDZ2_Tam_RS02175     gaccccagctcggcggccagccagttcctccggtccgtgcgcaacaccgcgcagaaccag
MxDZ2_Kirby_RS0212590 gaccccagctcggcggccagccagttcctccggtccgtgcgcaacaccgcgcagaaccag
MxDZ2_Nan_RS02810     ------------------------------------------------------------


MxDZ2_Tam_RS02175     gtcatccagttcctgaacacgggcgtcgcgtcgcccaccccgtaa
MxDZ2_Kirby_RS0212590 gtcatccagttcctgaacacgggcgtcgcgtcgcccaccccgtaa
MxDZ2_Nan_RS02810     ---------------------------------------------


-----------------------------------------------------------------------------------
multiple sequence alignment (protein)


MxDZ2_Tam_RS02175     MRKVLFLGAFALGASACTPDIAQDPPRSPEDSVVAEFDPTGSPPVVPSPNDLAIVNGLVN
MxDZ2_Nan_RS02810     MRKVLFLGAFALGASACTPDIAQDPPRSPEDSVVAEFDPTGSPPVVPSPNDLAIVNGLVN
MxDZ2_Kirby_RS0212590 MRKVLFLGAFALGASACTPDIAQDPPRSPEDSVVAEFDPTGSPPVVPSPNDLAIVNGLVN
                      ************************************************************

MxDZ2_Tam_RS02175     APINPQAPAAEQEFTRDYVNTLNGFPTTVTATTRIRGLEAGTVNTNTVKIIDVYAGTPLA
MxDZ2_Nan_RS02810     APINPQAPAAEQEFTRDYVNTLNGFPTTVTATTRIRGLEAGTVNTNTVKIIDVYAGTPLA
MxDZ2_Kirby_RS0212590 APINPQAPAAEQEFTRDYVNTLNGFPTTVTATTRIRGLEAGTVNTNTVKIIDVYAGTPLA
                      ************************************************************

MxDZ2_Tam_RS02175     KPATPSYIGYNEETNLVTIIPPLPGGWPKGGRYAVAIIGGENGVKTTAGKPVIPSATWAF
MxDZ2_Nan_RS02810     KPATPSYIGYNEETNLVTIIPPLPGGWPKGGRYAVAIIGGENGVKTTAGKPVIPSATWAF
MxDZ2_Kirby_RS0212590 KPATPSYIGYNEETNLVTIIPPLPGGWPKGGRYAVAIIGGENGVKTTAGKPVIPSATWAF
                      ************************************************************

MxDZ2_Tam_RS02175     ASSAESLVTCDDLAAPNCQTTTELIPSTVTEPAARLADQTASALRLEQLRRGYAPILKLV
MxDZ2_Nan_RS02810     ASSAESLVTCDDLAAPNCQTTTELIPSTVTEPAARLADQTASALRLEQLRRGYAPILKLV
MxDZ2_Kirby_RS0212590 ASSAESLVTCDDLAAPNCQTTTELIPSTVTEPAARLADQTASALRLEQLRRGYAPILKLV
                      ************************************************************

MxDZ2_Tam_RS02175     SEQFSVKREDIVLLWTFSIMDHPEATFDPGNGVVPFPNDLLRNPETGLLALPEPTEAGPA
MxDZ2_Nan_RS02810     SEQFSVKREDIVLLWTFSIMDHPEATFDPGNGVVPFPNDLLRNPETGLLALPEPTEAGPA
MxDZ2_Kirby_RS0212590 SEQFSVKREDIVLLWTFSIMDHPEATFDPGNGVVPFPNDLLRNPETGLLALPEPTEAGPA
                      ************************************************************

MxDZ2_Tam_RS02175     QMLIKGLNTLDGWSTTAPIVSENGAKRGPIDTGSLLDANTVRMKIGFTPDTEEEQKTTLR
MxDZ2_Nan_RS02810     QMLIKGLNTLDGWSTTAPIVSENGAKRGPIDTGSLLDANTVRMKIGFTPDTEEEQKTTLR
MxDZ2_Kirby_RS0212590 QMLIKGLNTLDGWSTTAPIVSENGAKRGPIDTGSLLDANTVRMKIGFTPDTEEEQKTTLR
                      ************************************************************

MxDZ2_Tam_RS02175     FVKLTNPTSGTNPQVKLCFNCEPVVEGLTPAPNSPQQLQIVPEVPLDEATQYGVVMLRGM
MxDZ2_Nan_RS02810     FVKLTNPTSGTNPQVKLCFNCEPVVEGLTPAPNSPQQLQIVPEVPLDEATQYGVVMLRGM
MxDZ2_Kirby_RS0212590 FVKLTNPTSGTNPQVKLCFNCEPVVEGLTPAPNSPQQLQIVPEVPLDEATQYGVVMLRGM
                      ************************************************************

MxDZ2_Tam_RS02175     KDTLGRTVAPTAAQALMRMVNPLVDANGKSQVAAVPDTLARDLERARVGMKPLFDGLEAA
MxDZ2_Nan_RS02810     KDTLGRTVAPTAAQALMRMVNPLVDANGKSQVAAVPDTLARDLERARVGMKPLFDGLEAA
MxDZ2_Kirby_RS0212590 KDTLGRTVAPTAAQALMRMVNPLVDANGKSQVAAVPDTLARDLERARVGMKPLFDGLEAA
                      ************************************************************

MxDZ2_Tam_RS02175     GIARKDINLAWAFTTQSTRSILERLNAAPTQVPVPADPVYLLDQTATIKGTMAGLTLDND
MxDZ2_Nan_RS02810     GIARKDINLAWAFTTQSTRSILERLNAAPTQVPVPADPVYLLDQTATIKGTMAGLTLDND
MxDZ2_Kirby_RS0212590 GIARKDINLAWAFTTQSTRSILERLNAAPTQVPVPADPVYLLDQTATIKGTMAGLTLDND
                      ************************************************************

MxDZ2_Tam_RS02175     AVGRVFIGAYHSPFFLDDAQGTLNRATPRIDRVPFMLFTPAGAAPADGYPVVIYGHGLTG
MxDZ2_Nan_RS02810     AVGRVFIGAYHSPFFLDDAQGTLNRATPRIDRVPFMLFTPAGAAPADGYPVVIYGHGLTG
MxDZ2_Kirby_RS0212590 AVGRVFIGAYHSPFFLDDAQGTLNRATPRIDRVPFMLFTPAGAAPADGYPVVIYGHGLTG
                      ************************************************************

MxDZ2_Tam_RS02175     NRTNILTVANNFNAAGYAVAAIDTVFHGERASCAGISEDAPVVVDENRDGVPELTITTPD
MxDZ2_Nan_RS02810     NRTNILTVANNFNAAGYAVAAIDTVFHGERASCAGISEDAPVVVDENRDGVPELTITTPD
MxDZ2_Kirby_RS0212590 NRTNILTVANNFNAAGYAVAAIDTVFHGERASCAGISEDAPVVVDENRDGVPELTITTPD
                      ************************************************************

MxDZ2_Tam_RS02175     AACDDGDTCDVTAGSPTFGRCVSNIQPACDVGPTASPHGDLFCSSQGRGRCVVTEPDSTT
MxDZ2_Nan_RS02810     AA----------------------------------------------------------
MxDZ2_Kirby_RS0212590 AACDDGDTCDVTAGSPTFGRCVSNIQPACDVGPTASPHGDLFCSSQGRGRCVVTEPDSTT
                      **


MxDZ2_Tam_RS02175     GTCEGGDFARPGPNQPPYVSGAGFLNLVNLFATRDNFRHHVVDFAQLARALDTDTLNGRL
MxDZ2_Nan_RS02810     ------------------------------------------------------------
MxDZ2_Kirby_RS0212590 GTCEGGDFARPGPNQPPYVSGAGFLNLVNLFATRDNFRHHVVDFAQLARALDTDTLNGRL


MxDZ2_Tam_RS02175     AAAGAGTLNTSQFSYVGQSLGGMQGVLSASVSPVVGRTALNAVGGDLVDILLTATDSTFV
MxDZ2_Nan_RS02810     ------------------------------------------------------------
MxDZ2_Kirby_RS0212590 AAAGAGTLNTSQFSYVGQSLGGMQGVLSASVSPVVGRTALNAVGGDLVDILLTATDSTFV


MxDZ2_Tam_RS02175     RYRSGFFETLENSGRAPGTPEFYEFITLARTILDPASPRNYGRYLENAPSAPANRNAFIQ
MxDZ2_Nan_RS02810     ------------------------------------------------------------
MxDZ2_Kirby_RS0212590 RYRSGFFETLENSGRAPGTPEFYEFITLARTILDPASPRNYGRYLENAPSAPANRNAFIQ


MxDZ2_Tam_RS02175     YIEGDMVIPNAVTEGLISAANRPLVGVSEPRTVQTYLAEDGMGLPTNVRHAFFGIVDPSL
MxDZ2_Nan_RS02810     ------------------------------------------------------------
MxDZ2_Kirby_RS0212590 YIEGDMVIPNAVTEGLISAANRPLVGVSEPRTVQTYLAEDGMGLPTNVRHAFFGIVDPSL


MxDZ2_Tam_RS02175     DPSSAASQFLRSVRNTAQNQVIQFLNTGVASPTP
MxDZ2_Nan_RS02810     ----------------------------------
MxDZ2_Kirby_RS0212590 DPSSAASQFLRSVRNTAQNQVIQFLNTGVASPTP
```

**Premature stop codon followed by non-coding region 1**

| **Tam_location** | **Tam_nucleotide** | **Kirby_nucleotide** | **Nan_nucleotide** | **Nan_location** |
| --- | --- | --- | --- | --- |
| **1332473** | **T** | **.** | **.** | **1332470** |

```
multiple sequence alignment (nucleotide)

MxDZ2_Kirby_RS0217305 atgccgtcacctccggaagaggtggatccgctcgcggacctgcgcgaaagcctggacctg
MxDZ2_Nan_RS32920     atgccgtcacctccggaagaggtggatccgctcgcggacctgcgcgaaagcctggacctg
MxDZ2_Tam_RS09970     atgccgtcacctccggaagaggtggatccgctcgcggacctgcgcgaaagcctggacctg
                      ************************************************************

MxDZ2_Kirby_RS0217305 gaagacaccggtgcctcggtggcccccgccgcccccgcccccgtggcctccaggcccttg
MxDZ2_Nan_RS32920     gaagacaccggtgcctcggtggcccccgccgcccccgcccccgtggcctccaggcccttg
MxDZ2_Tam_RS09970     gaagacaccggtgcctcggtggcccccgccgcccccgcccccgtggcctccaggcccttg
                      ************************************************************

MxDZ2_Kirby_RS0217305 ccaaagccaccgccctcggctccaccgcccgccgctccgcctccgttgccgccgcgccgt
MxDZ2_Nan_RS32920     ccaaagccaccgccctcggctccaccgcccgccgctccgcctccgttgccgccgcgccgt
MxDZ2_Tam_RS09970     ccaaagccaccgccctcggctccaccgcccgccgctccgcctccgttgccgccgcgccgt
                      ************************************************************

MxDZ2_Kirby_RS0217305 ccggccgcccccctggccgccgtgcccaagggcacgggagcgcccgcgtccccgaaggcg
MxDZ2_Nan_RS32920     ccggccgcccccctggccgccgtgcccaagggcacgggagcgcccgcgtccccgaaggcg
MxDZ2_Tam_RS09970     ccggccgcccccctggccgccgtgcccaagggcacgggagcgcccgcgtccccgaaggcg
                      ************************************************************

MxDZ2_Kirby_RS0217305 cccgcgtccccggctgccgctgctgcccccgccgtccgcgccgcgcgggccccggggaat
MxDZ2_Nan_RS32920     cccgcgtccccggctgccgctgctgcccccgccgtccgcgccgcgcgggccccggggaat
MxDZ2_Tam_RS09970     cccgcgtccccggctgccgctgctgcccccgccgtccgcgccgcgcgggccccggggaat
                      ************************************************************

MxDZ2_Kirby_RS0217305 gatccatttggtgagccgcccgaaatccggatgcccatgggcggctcgcccgaggacaag
MxDZ2_Nan_RS32920     gatccatttggtgagccgcccgaaatccggatgcccatgggcggctcgcccgaggacaag
MxDZ2_Tam_RS09970     gatccatttggtgagccgcccgaaatccggatgcccatgggcggctcgcccgaggacaag
                      ************************************************************

MxDZ2_Kirby_RS0217305 ctggagtacttccgggccgtggtccgccagaagacggagaccctggcgcgggcccggacg
MxDZ2_Nan_RS32920     ctggagtacttccgggccgtggtccgccagaagacggagaccctggcgcgggcccggacg
MxDZ2_Tam_RS09970     ctggagtacttccgggccgtggtccgccagaagacggagaccctggcgcgggcccggacg
                      ************************************************************

MxDZ2_Kirby_RS0217305 ctgtacgcggagcgcgacggcgagctcaacggcgtgaagcagtcgctgacgctggcgcgc
MxDZ2_Nan_RS32920     ctgtacgcggagcgcgacggcgagctcaacggcgtgaagcagtcgctgacgctggcgcgc
MxDZ2_Tam_RS09970     ctgtacgcggagcgcgacggcgagctcaacggcgtgaagcagtcgctgacgctggcgcgc
                      ************************************************************

MxDZ2_Kirby_RS0217305 aaggagctggcggagtcgaaggccagtgtctccgccttccaggaggagctggcccgggcg
MxDZ2_Nan_RS32920     aaggagctggcggagtcgaaggccagtgtctccgccttccaggaggagctggcccgggcg
MxDZ2_Tam_RS09970     aaggagctggcggagtcgaaggccagtgtctccgccttccaggaggagctggcccgggcg
                      ************************************************************

MxDZ2_Kirby_RS0217305 gaggcggagtcccagtcgcggctcgccagcctccaggatgagctctccgcggtggaggcg
MxDZ2_Nan_RS32920     gaggcggagtcccagtcgcggctcgccagcctccaggatgagctctccgcggtggaggcg
MxDZ2_Tam_RS09970     gaggcggagtcccagtcgcggctcgccagcctccaggatgagctctccgcggtggaggcg
                      ************************************************************

MxDZ2_Kirby_RS0217305 gaccgcaaggacctgtctcgcgcgctggcggaggtcgaatccgatgtgccgcgcctgacg
MxDZ2_Nan_RS32920     gaccgcaaggacctgtctcgcgcgctggcggaggtcgaatccgatgtgccgcgcctgacg
MxDZ2_Tam_RS09970     gaccgcaaggacctgtctcgcgcgctggcggaggtcgaatccgatgtgccgcgcctgacg
                      ************************************************************

MxDZ2_Kirby_RS0217305 gcggagctccaggaggagcgcgaggcccgtggcgccgtggccgaggagctgattggcgcg
MxDZ2_Nan_RS32920     gcggagctccaggaggagcgcgaggcccgtggcgccgtggccgaggagctgattggcgcg
MxDZ2_Tam_RS09970     gcggagctccaggaggagcgcgaggcccgtggcgccgtggccgaggagctgattggcgcg
                      ************************************************************

MxDZ2_Kirby_RS0217305 aaggaagcgctgtcgctggcccaggaccgcgtggcggagctggccgcggagaagtccgag
MxDZ2_Nan_RS32920     aaggaagcgctgtcgctggcccaggaccgcgtggcggagctggccgcggagaagtccgag
MxDZ2_Tam_RS09970     aaggaagcgctgtcgctggcccaggaccgcgtggcggagctggccgcggagaagtccgag
                      ************************************************************

MxDZ2_Kirby_RS0217305 gcccagggcgccctggaggccgttcaggagcagtaccagcaggccatcgccgacgtggag
MxDZ2_Nan_RS32920     gcccagggcgccctggaggccgttcaggagcagtaccagcaggccatcgccgacgtggag
MxDZ2_Tam_RS09970     gcccagggcgccctggaggccgttcaggagcagtaccagcaggccatcgccgacgtggag
                      ************************************************************

MxDZ2_Kirby_RS0217305 cggctgacctccgagctggaggccgcctcggcggagaaggactcgctgggcctgcgcacc
MxDZ2_Nan_RS32920     cggctgacctccgagctggaggccgcctcggcggagaaggactcgctgggcctgcgcacc
MxDZ2_Tam_RS09970     cggctgacctccgagctggaggccgcctcggcggagaaggactcgctgggcctgcgcacc
                      ************************************************************


MxDZ2_Kirby_RS0217305 gcgcagttggaggccgcgctggaggaggcccaatctgggctgagcgcgctggagagcgag
MxDZ2_Nan_RS32920     gcgcagttggaggccgcgctggaggaggcccaatctgggctgagcgcgctggagagcgag
MxDZ2_Tam_RS09970     gcgcagttggaggccgcgctggaggaggcccaatctgggctgagcgcgctggagagcgag
                      ************************************************************

MxDZ2_Kirby_RS0217305 agcgactggtcgaagagctcgctggaggaggccca-gggccgcgcgggtacgctggaggc
MxDZ2_Nan_RS32920     agcgactggtcgaagagctcgctggaggaggccca-gggccgcgcgggtacgctggaggc
MxDZ2_Tam_RS09970     agcgactggtcgaagagctcgctggaggaggcccaTgggccgcgcgggtacgctggaggc
                      *********************************** ************************

MxDZ2_Kirby_RS0217305 cgagcgcgacgaggcgcgcaagcaactggcggtggtggaggacgggctgcgcaccctgca
MxDZ2_Nan_RS32920     cgagcgcgacgaggcgcgcaagcaactggcggtggtggaggacgggctgcgcaccctgca
MxDZ2_Tam_RS09970     cgagcgcgacgaggcgcgcaagcaactggcggtggtggaggacgggctgcgcaccctgca
                      ************************************************************

MxDZ2_Kirby_RS0217305 ggagcaggtggcggagctggagcgttcgctggcgctgaaggacgccgaaatcgtgggcct
MxDZ2_Nan_RS32920     ggagcaggtggcggagctggagcgttcgctggcgctgaaggacgccgaaatcgtgggcct
MxDZ2_Tam_RS09970     ggagcaggtggcggagctggagcgttcgctggcgctgaaggacgccgaaatcgtgggcct
                      ************************************************************

MxDZ2_Kirby_RS0217305 gcgcgcggcgctcaccgcccgcaccaccgaggccgcggagctgcccgcgctgcgtcaggc
MxDZ2_Nan_RS32920     gcgcgcggcgctcaccgcccgcaccaccgaggccgcggagctgcccgcgctgcgtcaggc
MxDZ2_Tam_RS09970     gcgcgcggcgctcaccgcccgcaccaccgaggccgcggagctgcccgcgctgcgtcaggc
                      ************************************************************

MxDZ2_Kirby_RS0217305 gctggaggcgcgcaccgcggagctggcccagctcaaggcgaagctggaggcggaggcggc
MxDZ2_Nan_RS32920     gctggaggcgcgcaccgcggagctggcccagctcaaggcgaagctggaggcggaggcggc
MxDZ2_Tam_RS09970     gctggaggcgcgcaccgcggagctggcccagctcaaggcgaagctggaggcggaggcggc
                      ************************************************************

MxDZ2_Kirby_RS0217305 caaggccgaggagcgctcacaggccctggaagagggattggcacaggccagtgagcgggc
MxDZ2_Nan_RS32920     caaggccgaggagcgctcacaggccctggaagagggattggcacaggccagtgagcgggc
MxDZ2_Tam_RS09970     caaggccgaggagcgctcacaggccctggaagagggattggcacaggccagtga------
                      ******************************************************

MxDZ2_Kirby_RS0217305 gcacctggcggagggcgaggccgcggcgctgaaggaggcgctggaagccgccgaggtgga
MxDZ2_Nan_RS32920     gcacctggcggagggcgaggccgcggcgctgaaggaggcgctggaagccgccgaggtgga
MxDZ2_Tam_RS09970     ------------------------------------------------------------


MxDZ2_Kirby_RS0217305 gcaggtgtcgttgcgcgaccgcatggaagcggacgcggcggcgctgggcgaggcggtgca
MxDZ2_Nan_RS32920     gcaggtgtcgttgcgcgaccgcatggaagcggacgcggcggcgctgggcgaggcggtgca
MxDZ2_Tam_RS09970     ------------------------------------------------------------


MxDZ2_Kirby_RS0217305 gcaggccgagacgaaggtgtccgagctgacggcggcggtggaggccgcgcacgcggacca
MxDZ2_Nan_RS32920     gcaggccgagacgaaggtgtccgagctgacggcggcggtggaggccgcgcacgcggacca
MxDZ2_Tam_RS09970     ------------------------------------------------------------


MxDZ2_Kirby_RS0217305 tgcgtcgtggaaggaacagctcgccgcggtggagctgaaggcggccaccaaggacgccga
MxDZ2_Nan_RS32920     tgcgtcgtggaaggaacagctcgccgcggtggagctgaaggcggccaccaaggacgccga
MxDZ2_Tam_RS09970     ------------------------------------------------------------


MxDZ2_Kirby_RS0217305 gcgcgtcggcctggccgcccgggtgtccatgctggaggcggcgtccggccagcgcgaggc
MxDZ2_Nan_RS32920     gcgcgtcggcctggccgcccgggtgtccatgctggaggcggcgtccggccagcgcgaggc
MxDZ2_Tam_RS09970     ------------------------------------------------------------


MxDZ2_Kirby_RS0217305 ggagctggtgcggctgcaagcagaggtggcgaaggcccacgaggcgctggcgctggagcg
MxDZ2_Nan_RS32920     ggagctggtgcggctgcaagcagaggtggcgaaggcccacgaggcgctggcgctggagcg
MxDZ2_Tam_RS09970     ------------------------------------------------------------


MxDZ2_Kirby_RS0217305 tgcgcgccgcgaggcggcggaggtcgagcgcaccgatgcggaggtgaaggcggcccaagc
MxDZ2_Nan_RS32920     tgcgcgccgcgaggcggcggaggtcgagcgcaccgatgcggaggtgaaggcggcccaagc
MxDZ2_Tam_RS09970     ------------------------------------------------------------


MxDZ2_Kirby_RS0217305 cgaggcccaggtggagtcgctgacggtggggcagggcggcgccgaggcgcaggtggcctc
MxDZ2_Nan_RS32920     cgaggcccaggtggagtcgctgacggtggggcagggcggcgccgaggcgcaggtggcctc
MxDZ2_Tam_RS09970     ------------------------------------------------------------


MxDZ2_Kirby_RS0217305 gctcaccgaggagctggaggccgcgcgcgccgaggcggacaaggtggagcgcctccaggg
MxDZ2_Nan_RS32920     gctcaccgaggagctggaggccgcgcgcgccgaggcggacaaggtggagcgcctccaggg
MxDZ2_Tam_RS09970     ------------------------------------------------------------


MxDZ2_Kirby_RS0217305 ccggctgaagatgatggagggggccttggaaggcgcgaaggcccaggccgcggcggccgg
MxDZ2_Nan_RS32920     ccggctgaagatgatggagggggccttggaaggcgcgaaggcccaggccgcggcggccgg
MxDZ2_Tam_RS09970     ------------------------------------------------------------


MxDZ2_Kirby_RS0217305 caagtcggacgcggcgcgcgctgcgaccgaggcgaagctggctcgcgcggaggcgtcgct
MxDZ2_Nan_RS32920     caagtcggacgcggcgcgcgctgcgaccgaggcgaagctggctcgcgcggaggcgtcgct
MxDZ2_Tam_RS09970     ------------------------------------------------------------


MxDZ2_Kirby_RS0217305 gaaggccgaggagcagaagcgggccgacgtggaatcctcgctgcgagcagagcaggaggc
MxDZ2_Nan_RS32920     gaaggccgaggagcagaagcgggccgacgtggaatcctcgctgcgagcagagcaggaggc
MxDZ2_Tam_RS09970     ------------------------------------------------------------


MxDZ2_Kirby_RS0217305 acggcgggcgctcgaggcgaagctggcggagcagccggggactccctcccagagcgcggg
MxDZ2_Nan_RS32920     acggcgggcgctcgaggcgaagctggcggagcagccggggactccctcccagagcgcggg
MxDZ2_Tam_RS09970     ------------------------------------------------------------


MxDZ2_Kirby_RS0217305 tgcgccagcggactgggaggcggagcgcgagcagctcaaggcggacgctgccaacctcaa
MxDZ2_Nan_RS32920     tgcgccagcggactgggaggcggagcgcgagcagctcaaggcggacgctgccaacctcaa
MxDZ2_Tam_RS09970     ------------------------------------------------------------


MxDZ2_Kirby_RS0217305 gcgcaagctggtggcggccgagacagcgctcgagaacgcggcgggctacaaggcgaagat
MxDZ2_Nan_RS32920     gcgcaagctggtggcggccgagacagcgctcgagaacgcggcgggctacaaggcgaagat
MxDZ2_Tam_RS09970     ------------------------------------------------------------


MxDZ2_Kirby_RS0217305 tgcccggctggaggcgcaattgaagggcaagaggtag
MxDZ2_Nan_RS32920     tgcccggctggaggcgcaattgaagggcaagaggtag
MxDZ2_Tam_RS09970     -------------------------------------

-----------------------------------------------------------------------------------
multiple sequence alignment (protein)

MxDZ2_Kirby_RS0217305 MPSPPEEVDPLADLRESLDLEDTGASVAPAAPAPVASRPLPKPPPSAPPPAAPPPLPPRR
MxDZ2_Nan_RS32920     MPSPPEEVDPLADLRESLDLEDTGASVAPAAPAPVASRPLPKPPPSAPPPAAPPPLPPRR
MxDZ2_Tam_RS09970     MPSPPEEVDPLADLRESLDLEDTGASVAPAAPAPVASRPLPKPPPSAPPPAAPPPLPPRR
                      ************************************************************

MxDZ2_Kirby_RS0217305 PAAPLAAVPKGTGAPASPKAPASPAAAAAPAVRAARAPGNDPFGEPPEIRMPMGGSPEDK
MxDZ2_Nan_RS32920     PAAPLAAVPKGTGAPASPKAPASPAAAAAPAVRAARAPGNDPFGEPPEIRMPMGGSPEDK
MxDZ2_Tam_RS09970     PAAPLAAVPKGTGAPASPKAPASPAAAAAPAVRAARAPGNDPFGEPPEIRMPMGGSPEDK
                      ************************************************************

MxDZ2_Kirby_RS0217305 LEYFRAVVRQKTETLARARTLYAERDGELNGVKQSLTLARKELAESKASVSAFQEELARA
MxDZ2_Nan_RS32920     LEYFRAVVRQKTETLARARTLYAERDGELNGVKQSLTLARKELAESKASVSAFQEELARA
MxDZ2_Tam_RS09970     LEYFRAVVRQKTETLARARTLYAERDGELNGVKQSLTLARKELAESKASVSAFQEELARA
                      ************************************************************

MxDZ2_Kirby_RS0217305 EAESQSRLASLQDELSAVEADRKDLSRALAEVESDVPRLTAELQEEREARGAVAEELIGA
MxDZ2_Nan_RS32920     EAESQSRLASLQDELSAVEADRKDLSRALAEVESDVPRLTAELQEEREARGAVAEELIGA
MxDZ2_Tam_RS09970     EAESQSRLASLQDELSAVEADRKDLSRALAEVESDVPRLTAELQEEREARGAVAEELIGA
                      ************************************************************

MxDZ2_Kirby_RS0217305 KEALSLAQDRVAELAAEKSEAQGALEAVQEQYQQAIADVERLTSELEAASAEKDSLGLRT
MxDZ2_Nan_RS32920     KEALSLAQDRVAELAAEKSEAQGALEAVQEQYQQAIADVERLTSELEAASAEKDSLGLRT
MxDZ2_Tam_RS09970     KEALSLAQDRVAELAAEKSEAQGALEAVQEQYQQAIADVERLTSELEAASAEKDSLGLRT
                      ************************************************************

MxDZ2_Kirby_RS0217305 AQLEAALEEAQSGLSALESESDWSKSSLEEAQGRAGTLEAERDEARKQLAVVEDGLRTLQ
MxDZ2_Nan_RS32920     AQLEAALEEAQSGLSALESESDWSKSSLEEAQGRAGTLEAERDEARKQLAVVEDGLRTLQ
MxDZ2_Tam_RS09970     AQLEAALEEAQSGLSALESESDWSKSSLEEAHGPRGYAGG---RARRGAQATGGG-----
                      *******************************:*  *   .   .**:   .. .*

MxDZ2_Kirby_RS0217305 EQVAELERSLALKDAEIVGLRAALTARTTEAAELPA----LRQALEARTAELAQLKAKLE
MxDZ2_Nan_RS32920     EQVAELERSLALKDAEIVGLRAALTARTTEAAELPA----LRQALEARTAELAQLKAKLE
MxDZ2_Tam_RS09970     ------------------GGRAAHPAGAGGGAGAFAGAEGRRNRGPARGAHRPHHRGRGA
                                        * *** .* :  .*   *     *:   ** *. .: :.:

MxDZ2_Kirby_RS0217305 AEAAKAEERSQALEEGLAQASERAHLAEGEAAALKEALEAAEVEQVSLRDRMEADAAALG
MxDZ2_Nan_RS32920     AEAAKAEERSQALEEGLAQASERAHLAEGEAAALKEALEAAEVEQVSLRDRMEADAAALG
MxDZ2_Tam_RS09970     ARAA----------SGAGGAHRGAGPAQGEAGG---------------------------
                      *.**          .* . * . *  *:***..

MxDZ2_Kirby_RS0217305 EAVQQAETKVSELTAAVEAAHADHASWKEQLAAVELKAATKDAERVGLAARVSMLEAASG
MxDZ2_Nan_RS32920     EAVQQAETKVSELTAAVEAAHADHASWKEQLAAVELKAATKDAERVGLAARVSMLEAASG
MxDZ2_Tam_RS09970     ------------------------------------------------------------


MxDZ2_Kirby_RS0217305 QREAELVRLQAEVAKAHEALALERARREAAEVERTDAEVKAAQAEAQVESLTVGQGGAEA
MxDZ2_Nan_RS32920     QREAELVRLQAEVAKAHEALALERARREAAEVERTDAEVKAAQAEAQVESLTVGQGGAEA
MxDZ2_Tam_RS09970     -------------------------------------------------------GGG--
                                                                             **.

MxDZ2_Kirby_RS0217305 QVASLTEELEAARAEADKVERLQGRLKMMEGALEGAKAQAAAAGKSDAARAATEAKLARA
MxDZ2_Nan_RS32920     QVASLTEELEAARAEADKVERLQGRLKMMEGALEGAKAQAAAAGKSDAARAATEAKLARA
MxDZ2_Tam_RS09970     ----------------------QGR-----GALTG-------------------------
                                            ***     *** *

MxDZ2_Kirby_RS0217305 EASLKAEEQKRADVESSLRAEQEARRALEAKLAEQPGTPSQSAGAPADWEAEREQLKADA
MxDZ2_Nan_RS32920     EASLKAEEQKRADVESSLRAEQEARRALEAKLAEQPGTPSQSAGAPADWEAEREQLKADA
MxDZ2_Tam_RS09970     --------------------------------------PGRGIGT---------------
                                                            *.:. *:

MxDZ2_Kirby_RS0217305 ANLKRKLVAAETALENAAGYKAKIARLEAQLKGKR
MxDZ2_Nan_RS32920     ANLKRKLVAAETALENAAGYKAKIARLEAQLKGKR
MxDZ2_Tam_RS09970     ---------------------------------GQ
                                                        :
```

**Premature stop codon followed by non-coding region 2**

| **Tam_location** | **Tam_nucleotide** | **Kirby_nucleotide** | **Nan_nucleotide** | **Nan_location** |
| --- | --- | --- | --- | --- |
| **2292711** | **A (complement)** | **.** | **.** | **2292696** |

```
multiple sequence alignment (nucleotide)

MxDZ2_Kirby_RS39310   gtgcccgtgccccaggggcgcctggtcgcgggcttcgatgtgggtcgcaccagagaccgc
MxDZ2_Nan_RS28750     gtgcccgtgccccaggggcgcctggtcgcgggcttcgatgtgggtcgcaccagagaccgc
MxDZ2_Tam_RS38160     gtgcccgtgccccaggggcgcctggtcgcgggcttcgatgtgggtcgcaccagagaccgc
                      ************************************************************

MxDZ2_Kirby_RS39310   tcggagctggccgtcttcgaggagacgggtgggcgcttcacgtgccgcatgctgcgcagc
MxDZ2_Nan_RS28750     tcggagctggccgtcttcgaggagacgggtgggcgcttcacgtgccgcatgctgcgcagc
MxDZ2_Tam_RS38160     tcggagctggccgtcttcgaggagacgggtgggcgcttcacgtgccgcatgctgcgcagc
                      ************************************************************

MxDZ2_Kirby_RS39310   ttcgagggcgtgcccttcgcggagcaggaggcgcacctccgtcgtctgctgtccgtgctt
MxDZ2_Nan_RS28750     ttcgagggcgtgcccttcgcggagcaggaggcgcacctccgtcgtctgctgtccgtgctt
MxDZ2_Tam_RS38160     ttcgagggcgtgcccttcgcggagcaggaggcgcacctccgtcgtctgctgtccgtgctt
                      ************************************************************

MxDZ2_Kirby_RS39310   cccgtggctcgtctcagcgtcgaccgcagcggcatcggaatgaatctcgcggagaacctc
MxDZ2_Nan_RS28750     cccgtggctcgtctcagcgtcgaccgcagcggcatcggaatgaatctcgcggagaacctc
MxDZ2_Tam_RS38160     cccgtggctcgtctcagcgtcgaccgcagcggcatcggaatgaatctcgcggagaacctc
                      ************************************************************

MxDZ2_Kirby_RS39310   gcccgcgactacccccaggtggtggaggagaacttcaccaacgaggccaaggagcgctgg
MxDZ2_Nan_RS28750     gcccgcgactacccccaggtggtggaggagaacttcaccaacgaggccaaggagcgctgg
MxDZ2_Tam_RS38160     gcccgcgactacccccaggtggtggaggagaacttcaccaacgaggccaaggagcgctgg
                      ************************************************************

MxDZ2_Kirby_RS39310   gccaccgacttcaagattctcctccagcgccgcgatgtcacgctgccgcgccagc-gtga
MxDZ2_Nan_RS28750     gccaccgacttcaagattctcctccagcgccgcgatgtcacgctgccgcgccagc-gtga
MxDZ2_Tam_RS38160     gccaccgacttcaagattctcctccagcgccgcgatgtcacgctgccgcgccagcTgtga
                      ******************************************************* ****

MxDZ2_Kirby_RS39310   gcttgtcgggcagattcactccatcaagcgtcgcgtgctgccctcggggaaggtgtcctt
MxDZ2_Nan_RS28750     gcttgtcgggcagattcactccatcaagcgtcgcgtgctgccctcggggaaggtgtcctt
MxDZ2_Tam_RS38160     ------------------------------------------------------------


MxDZ2_Kirby_RS39310   cgacgccgaacgcacctcccgcggccatgcggacaaattctgggccgtggcccttgcctg
MxDZ2_Nan_RS28750     cgacgccgaacgcacctcccgcggccatgcggacaaattctgggccgtggcccttgcctg
MxDZ2_Tam_RS38160     ------------------------------------------------------------


MxDZ2_Kirby_RS39310   ccagcgggagcgc-----------------------------------------------
MxDZ2_Nan_RS28750     ccagcgggagcgccaggccgacaagccgagggcaggcgaaattggggtgcgtgtgattgg
MxDZ2_Tam_RS38160     ------------------------------------------------------------


MxDZ2_Kirby_RS39310   ----
MxDZ2_Nan_RS28750     ctaa
MxDZ2_Tam_RS38160     ----


-----------------------------------------------------------------------------------
multiple sequence alignment (protein)


MxDZ2_Kirby_RS39310   MPVPQGRLVAGFDVGRTRDRSELAVFEETGGRFTCRMLRSFEGVPFAEQEAHLRRLLSVL
MxDZ2_Nan_RS28750     MPVPQGRLVAGFDVGRTRDRSELAVFEETGGRFTCRMLRSFEGVPFAEQEAHLRRLLSVL
MxDZ2_Tam_RS38160     MPVPQGRLVAGFDVGRTRDRSELAVFEETGGRFTCRMLRSFEGVPFAEQEAHLRRLLSVL
                      ************************************************************

MxDZ2_Kirby_RS39310   PVARLSVDRSGIGMNLAENLARDYPQVVEENFTNEAKERWATDFKILLQRRDVTLPRQRE
MxDZ2_Nan_RS28750     PVARLSVDRSGIGMNLAENLARDYPQVVEENFTNEAKERWATDFKILLQRRDVTLPRQRE
MxDZ2_Tam_RS38160     PVARLSVDRSGIGMNLAENLARDYPQVVEENFTNEAKERWATDFKILLQRRDVTLPRQL-
                      **********************************************************

MxDZ2_Kirby_RS39310   LVGQIHSIKRRVLPSGKVSFDAERTSRGHADKFWAVALACQRER----------------
MxDZ2_Nan_RS28750     LVGQIHSIKRRVLPSGKVSFDAERTSRGHADKFWAVALACQRERQADKPRAGEIGVRVIG
MxDZ2_Tam_RS38160     ------------------------------------------------------------
```

**Premature stop codon followed by non-coding region 3**

| **Tam_location** | **Tam_nucleotide** | **Kirby nucleotide** | **Nan_nucleotide** | **Nan_location** |
| --- | --- | --- | --- | --- |
| **4281332** | **A (complement)** | **C (complement)** | **C (complement)** | **3936043** |

```
multiple sequence alignment (nucleotide)

MxDZ2_Kirby_RS0218815 ttgccgccgctggaacgcgcgttccgcaccgtgctggccgaggaactcgccgggcaactg
MxDZ2_Nan_RS20225     ttgccgccgctggaacgcgcgttccgcaccgtgctggccgaggaactcgccgggcaactg
MxDZ2_Tam_RS22640     ttgccgccgctggaacgcgcgttccgcaccgtgctggccgaggaactcgccgggcaactg
                      ************************************************************

MxDZ2_Kirby_RS0218815 tcgccgctgacggacacgcttctgcgcatcgcggagagcctgtcccccagcggcctccgg
MxDZ2_Nan_RS20225     tcgccgctgacggacacgcttctgcgcatcgcggagagcctgtcccccagcggcctccgg
MxDZ2_Tam_RS22640     tcgccgctgacggacacgcttctgcgcatcgcggagagcctgtcccccagcggcctccgg
                      ************************************************************

MxDZ2_Kirby_RS0218815 cgccagccgcggcgtgagctgccgaacgcggccctggacgccctggcgtcgtcgctcgcg
MxDZ2_Nan_RS20225     cgccagccgcggcgtgagctgccgaacgcggccctggacgccctggcgtcgtcgctcgcg
MxDZ2_Tam_RS22640     cgccagccgcggcgtgagctgccgaacgcggccctggacgccctggcgtcgtcgctcgcg
                      ************************************************************

MxDZ2_Kirby_RS0218815 accctggatacggggcccgacgacgccaccgaggaagaagtcgccccggaagcccgagcc
MxDZ2_Nan_RS20225     accctggatacggggcccgacgacgccaccgaggaagaagtcgccccggaagcccgagcc
MxDZ2_Tam_RS22640     accctggatacggggcccgacgacgccaccgaggaagaagtcgccccggaagcccgagcc
                      ************************************************************

MxDZ2_Kirby_RS0218815 tgcgccgtcatcggctgtaagcggccggtgcgcagcctgggctactgcgcggcccactac
MxDZ2_Nan_RS20225     tgcgccgtcatcggctgtaagcggccggtgcgcagcctgggctactgcgcggcccactac
MxDZ2_Tam_RS22640     tgcgccgtcatcggctgtaagcggccggtgcgcagcctgggctactgcgcggcccactac
                      ************************************************************

MxDZ2_Kirby_RS0218815 cagaagcggcgcctcatggtggcgtccggccgcctgcaccacgcgtggacggaagacgcc
MxDZ2_Nan_RS20225     cagaagcggcgcctcatggtggcgtccggccgcctgcaccacgcgtggacggaagacgcc
MxDZ2_Tam_RS22640     cagaagcggcgcctcatggtggcgtccggccgcctgcaccacgcgtggacggaagacgcc
                      ************************************************************

MxDZ2_Kirby_RS0218815 gcgccgaacagcattccggacgtcatcctgggccggaagcgccgcgagtcttcggaggca
MxDZ2_Nan_RS20225     gcgccgaacagcattccggacgtcatcctgggccggaagcgccgcgagtcttcggaggca
MxDZ2_Tam_RS22640     gcgccgaacagcattccggacgtcatcctgggccggaagcgccgcgagtcttcggaggca
                      ************************************************************

MxDZ2_Kirby_RS0218815 ccggagaccgccgtcaccccggagcccgtgcgtcccagcgaggggcctcgcgtctgggtc
MxDZ2_Nan_RS20225     ccggagaccgccgtcaccccggagcccgtgcgtcccagcgaggggcctcgcgtctgggtc
MxDZ2_Tam_RS22640     ccggagaccgccgtcaccccggagcccgtgcgtcccagcgaggggcctcgcgtctgggtc
                      ************************************************************

MxDZ2_Kirby_RS0218815 cgcaagaagggcgccagccccgtggtcccgacgcccagtggcgacagcgcgccgattctg
MxDZ2_Nan_RS20225     cgcaagaagggcgccagccccgtggtcccgacgcccagtggcgacagcgcgccgattctg
MxDZ2_Tam_RS22640     cgcaagaagggcgccagccccgtggtcccgacgcccagtggcgacagcgcgccgattctg
                      ************************************************************

MxDZ2_Kirby_RS0218815 cagttggcccaggcttcggtgattccgaccgcgcagcgctccgagcgcgagcaggtggca
MxDZ2_Nan_RS20225     cagttggcccaggcttcggtgattccgaccgcgcagcgctccgagcgcgagcaggtggca
MxDZ2_Tam_RS22640     cagttggcccaggcttcggtgattccgaccgcgcagcgctccgagcgcgagcaggtggca
                      ************************************************************

MxDZ2_Kirby_RS0218815 tccatcgtcGagcagtgggccagcgagttccgggccaacaagcatcgctcgtaa
MxDZ2_Nan_RS20225     tccatcgtcGagcagtgggccagcgagttccgggccaacaagcatcgctcgtaa
MxDZ2_Tam_RS22640     tccatcgtcTag------------------------------------------
                      ********* **
-----------------------------------------------------------------------------------
multiple sequence alignment (protein)

MxDZ2_Kirby_RS0218815 MPPLERAFRTVLAEELAGQLSPLTDTLLRIAESLSPSGLRRQPRRELPNAALDALASSLA
MxDZ2_Nan_RS20225     MPPLERAFRTVLAEELAGQLSPLTDTLLRIAESLSPSGLRRQPRRELPNAALDALASSLA
MxDZ2_Tam_RS22640     MPPLERAFRTVLAEELAGQLSPLTDTLLRIAESLSPSGLRRQPRRELPNAALDALASSLA
                      ************************************************************

MxDZ2_Kirby_RS0218815 TLDTGPDDATEEEVAPEARACAVIGCKRPVRSLGYCAAHYQKRRLMVASGRLHHAWTEDA
MxDZ2_Nan_RS20225     TLDTGPDDATEEEVAPEARACAVIGCKRPVRSLGYCAAHYQKRRLMVASGRLHHAWTEDA
MxDZ2_Tam_RS22640     TLDTGPDDATEEEVAPEARACAVIGCKRPVRSLGYCAAHYQKRRLMVASGRLHHAWTEDA
                      ************************************************************

MxDZ2_Kirby_RS0218815 APNSIPDVILGRKRRESSEAPETAVTPEPVRPSEGPRVWVRKKGASPVVPTPSGDSAPIL
MxDZ2_Nan_RS20225     APNSIPDVILGRKRRESSEAPETAVTPEPVRPSEGPRVWVRKKGASPVVPTPSGDSAPIL
MxDZ2_Tam_RS22640     APNSIPDVILGRKRRESSEAPETAVTPEPVRPSEGPRVWVRKKGASPVVPTPSGDSAPIL
                      ************************************************************

MxDZ2_Kirby_RS0218815 QLAQASVIPTAQRSEREQVASIVEQWASEFRANKHRS
MxDZ2_Nan_RS20225     QLAQASVIPTAQRSEREQVASIVEQWASEFRANKHRS
MxDZ2_Tam_RS22640     QLAQASVIPTAQRSEREQVASIV--------------
                      ***********************
```

**The yellow highlighted portion indicates the non-coding region.**

**Non-coding region mutation in MxDZ2_Nan 1**

| **Tam_location** | **Tam_nucleotide** | **Kirby nucleotide** | **Nan_nucleotide** | **Nan_location** |
| --- | --- | --- | --- | --- |
| **1992143** | **G (complement)** | **.** | **.** | **1992137** |

```
multiple sequence alignment (nucleotide)

MxDZ2_Tam_RS12675   atgcgattccccttcgttcccctcggcctgatgctcacggcgctgacgggagcccccggc
MxDZ2_Nan_RS30200   atgcgattccccttcgttcccctcggcctgatgctcacggcgctgacgggagcccccggc
MxDZ2_Kirby_RS38395 atgcgattccccttcgttcccctcggcctgatgctcacggcgctgacgggagcccccggc
                    ************************************************************

MxDZ2_Tam_RS12675   ctggcgctggatgacgctcccccccccCtcgccccctccggaaccacgccggatgcgcag
MxDZ2_Nan_RS30200   ctggcgctggatgacgctccccccccc-tcgccccctccggaaccacgccggatgcgcag
MxDZ2_Kirby_RS38395 ctggcgctggatgacgctccccccccc-tcgccccctccggaaccacgccggatgcgcag
                    *************************** ********************************

MxDZ2_Tam_RS12675   gtccaggcgaagctggacctggcggaccggttcgtccgccgggcggaactcctgcgggac
MxDZ2_Nan_RS30200   gtccaggcgaagctggacctggcggaccggttcgtccgccgggcggaactcctgcgggac
MxDZ2_Kirby_RS38395 gtccaggcgaagctggacctggcggaccggttcgtccgccgggcggaactcctgcgggac
                    ************************************************************

MxDZ2_Tam_RS12675   agcctggttccccccaggtggctgcgtgagggcgaccggttggtcttctggtcccgtgag
MxDZ2_Nan_RS30200   agcctggttccccccaggtggctgcgtgagggcgaccggttggtcttctggtcccgtgag
MxDZ2_Kirby_RS38395 agcctggttccccccaggtggctgcgtgagggcgaccggttggtcttctggtcccgtgag
                    ************************************************************

MxDZ2_Tam_RS12675   ggcaaggacggcgggacgtgggtgctggcgcacgcgaagaccggggagctgaagccactc
MxDZ2_Nan_RS30200   ggcaaggacggcgggacgtgggtgctggcgcacgcgaagaccggggagctgaagccactc
MxDZ2_Kirby_RS38395 ggcaaggacggcgggacgtgggtgctggcgcacgcgaagaccggggagctgaagccactc
                    ************************************************************

MxDZ2_Tam_RS12675   ctttccggtgagcaactgaggcagcagctctcgacgctgctgggcaagcccatcaccgcg
MxDZ2_Nan_RS30200   ctttccggtgagcaactgaggcagcagctctcgacgctgctgggcaagcccatcaccgcg
MxDZ2_Kirby_RS38395 ctttccggtgagcaactgaggcagcagctctcgacgctgctgggcaagcccatcaccgcg
                    ************************************************************


MxDZ2_Tam_RS12675   ccccgcttcttcgacgtcgcgctcgctccggatgagcggggcatcgtgttccgcctggag
MxDZ2_Nan_RS30200   ccccgcttcttcgacgtcgcgctcgctccggatgagcggggcatcgtgttccgcctggag
MxDZ2_Kirby_RS38395 ccccgcttcttcgacgtcgcgctcgctccggatgagcggggcatcgtgttccgcctggag
                    ************************************************************

MxDZ2_Tam_RS12675   gggaagacctttggcctgggcctgtcaggtggccatgtcaccttgctgtcgccggaggac
MxDZ2_Nan_RS30200   gggaagacctttggcctgggcctgtcaggtggccatgtcaccttgctgtcgccggaggac
MxDZ2_Kirby_RS38395 gggaagacctttggcctgggcctgtcaggtggccatgtcaccttgctgtcgccggaggac
                    ************************************************************

MxDZ2_Tam_RS12675   cgggccgcgctgacgctgtcgccccagcacttcctcgcgccgaaaggcggcgcgctcgcc
MxDZ2_Nan_RS30200   cgggccgcgctgacgctgtcgccccagcacttcctcgcgccgaaaggcggcgcgctcgcc
MxDZ2_Kirby_RS38395 cgggccgcgctgacgctgtcgccccagcacttcctcgcgccgaaaggcggcgcgctcgcc
                    ************************************************************

MxDZ2_Tam_RS12675   gtgcaacgccagggcggcttcgcggtgctgaacgcggaaggcgccaccgtggtcgagcgc
MxDZ2_Nan_RS30200   gtgcaacgccagggcggcttcgcggtgctgaacgcggaaggcgccaccgtggtcgagcgc
MxDZ2_Kirby_RS38395 gtgcaacgccagggcggcttcgcggtgctgaacgcggaaggcgccaccgtggtcgagcgc
                    ************************************************************

MxDZ2_Tam_RS12675   acgggggaagcgaacctcgactggcggattccggagcgtccctggtcaccggatggccgc
MxDZ2_Nan_RS30200   acgggggaagcgaacctcgactggcggattccggagcgtccctggtcaccggatggccgc
MxDZ2_Kirby_RS38395 acgggggaagcgaacctcgactggcggattccggagcgtccctggtcaccggatggccgc
                    ************************************************************

MxDZ2_Tam_RS12675   ttcctggtggtgtggcgggacgacctccgcgccgttcaccaggtgcccgtcgtggactac
MxDZ2_Nan_RS30200   ttcctggtggtgtggcgggacgacctccgcgccgttcaccaggtgcccgtcgtggactac
MxDZ2_Kirby_RS38395 ttcctggtggtgtggcgggacgacctccgcgccgttcaccaggtgcccgtcgtggactac
                    ************************************************************

MxDZ2_Tam_RS12675   tcctccgcgctggagaaggtgacgacggtgccgtacacgaagtccgggacgccactgccg
MxDZ2_Nan_RS30200   tcctccgcgctggagaaggtgacgacggtgccgtacacgaagtccgggacgccactgccg
MxDZ2_Kirby_RS38395 tcctccgcgctggagaaggtgacgacggtgccgtacacgaagtccgggacgccactgccg
                    ************************************************************

MxDZ2_Tam_RS12675   cgcgcggagctccatgtcgtggaagtggcgacgggccgcgtgacgcgcgttccgcccgtc
MxDZ2_Nan_RS30200   cgcgcggagctccatgtcgtggaagtggcgacgggccgcgtgacgcgcgttccgcccgtc
MxDZ2_Kirby_RS38395 cgcgcggagctccatgtcgtggaagtggcgacgggccgcgtgacgcgcgttccgcccgtc
                    ************************************************************


MxDZ2_Tam_RS12675   gagggcgagacatatgactggttcgccggctggaatcccgagggcaccgaggcgctgtgt
MxDZ2_Nan_RS30200   gagggcgagacatatgactggttcgccggctggaatcccgagggcaccgaggcgctgtgt
MxDZ2_Kirby_RS38395 gagggcgagacatatgactggttcgccggctggaatcccgagggcaccgaggcgctgtgt
                    ************************************************************

MxDZ2_Tam_RS12675   ctccacctctcgcgtgacgccaagcggctggacctgagcgtcgtggaacccgcgtcgggc
MxDZ2_Nan_RS30200   ctccacctctcgcgtgacgccaagcggctggacctgagcgtcgtggaacccgcgtcgggc
MxDZ2_Kirby_RS38395 ctccacctctcgcgtgacgccaagcggctggacctgagcgtcgtggaacccgcgtcgggc
                    ************************************************************

MxDZ2_Tam_RS12675   cagcgtcggcacgtcctgcgcgaggagcgcccggagaccttcgtcaccggcctggacttc
MxDZ2_Nan_RS30200   cagcgtcggcacgtcctgcgcgaggagcgcccggagaccttcgtcaccggcctggacttc
MxDZ2_Kirby_RS38395 cagcgtcggcacgtcctgcgcgaggagcgcccggagaccttcgtcaccggcctggacttc
                    ************************************************************

MxDZ2_Tam_RS12675   gcggtgggcggctgggcgaagcaggtgacggcgctgccaggagggcgcggctacctctgg
MxDZ2_Nan_RS30200   gcggtgggcggctgggcgaagcaggtgacggcgctgccaggagggcgcggctacctctgg
MxDZ2_Kirby_RS38395 gcggtgggcggctgggcgaagcaggtgacggcgctgccaggagggcgcggctacctctgg
                    ************************************************************

MxDZ2_Tam_RS12675   atgtccgagcgtgatggctggcgccacgtgtatgcgtatgaccgctcggggaagcgcgtg
MxDZ2_Nan_RS30200   atgtccgagcgtgatggctggcgccacgtgtatgcgtatgaccgctcggggaagcgcgtg
MxDZ2_Kirby_RS38395 atgtccgagcgtgatggctggcgccacgtgtatgcgtatgaccgctcggggaagcgcgtg
                    ************************************************************

MxDZ2_Tam_RS12675   aggcagctcacgcgaggcgcctttcccgtgcacgaagtggtgggagtcgcccccacggga
MxDZ2_Nan_RS30200   aggcagctcacgcgaggcgcctttcccgtgcacgaagtggtgggagtcgcccccacggga
MxDZ2_Kirby_RS38395 aggcagctcacgcgaggcgcctttcccgtgcacgaagtggtgggagtcgcccccacggga
                    ************************************************************

MxDZ2_Tam_RS12675   gatgccctctacgtgctggcctccgccgacagcggcgcgccatacgagcacctcttctac
MxDZ2_Nan_RS30200   gatgccctctacgtgctggcctccgccgacagcggcgcgccatacgagcacctcttctac
MxDZ2_Kirby_RS38395 gatgccctctacgtgctggcctccgccgacagcggcgcgccatacgagcacctcttctac
                    ************************************************************

MxDZ2_Tam_RS12675   cggggaagcctgaagggaggcgcgttgaagcggctgtcctctgcctcggggatgcaccgc
MxDZ2_Nan_RS30200   cggggaagcctgaagggaggcgcgttgaagcggctgtcctctgcctcggggatgcaccgc
MxDZ2_Kirby_RS38395 cggggaagcctgaagggaggcgcgttgaagcggctgtcctctgcctcggggatgcaccgc
                    ************************************************************


MxDZ2_Tam_RS12675   atcacgccgtccccctccggccagtactacgtggacacatggtcctcgcggacacagccc
MxDZ2_Nan_RS30200   atcacgccgtccccctccggccagtactacgtggacacatggtcctcgcggacacagccc
MxDZ2_Kirby_RS38395 atcacgccgtccccctccggccagtactacgtggacacatggtcctcgcggacacagccc
                    ************************************************************


MxDZ2_Tam_RS12675   cggttgagggagctggtgtccgtggagggtggcaagcgcgtgcggctcaccacgtcggac
MxDZ2_Nan_RS30200   cggttgagggagctggtgtccgtggagggtggcaagcgcgtgcggctcaccacgtcggac
MxDZ2_Kirby_RS38395 cggttgagggagctggtgtccgtggagggtggcaagcgcgtgcggctcaccacgtcggac
                    ************************************************************

MxDZ2_Tam_RS12675   gcgagtgagttcgagtcgctcggcgacacgcggccggaggcgctgctcgtcaaggcggcc
MxDZ2_Nan_RS30200   gcgagtgagttcgagtcgctcggcgacacgcggccggaggcgctgctcgtcaaggcggcc
MxDZ2_Kirby_RS38395 gcgagtgagttcgagtcgctcggcgacacgcggccggaggcgctgctcgtcaaggcggcc
                    ************************************************************

MxDZ2_Tam_RS12675   gatggcgtcacgcccctgcacggcgtgctctacaagccacgcgacttcgacgcctcgaag
MxDZ2_Nan_RS30200   gatggcgtcacgcccctgcacggcgtgctctacaagccacgcgacttcgacgcctcgaag
MxDZ2_Kirby_RS38395 gatggcgtcacgcccctgcacggcgtgctctacaagccacgcgacttcgacgcctcgaag
                    ************************************************************

MxDZ2_Tam_RS12675   cgctaccccgtgctcgcctccatctacgcgggtccgttcaccaccgttgtcccctggagc
MxDZ2_Nan_RS30200   cgctaccccgtgctcgcctccatctacgcgggtccgttcaccaccgttgtcccctggagc
MxDZ2_Kirby_RS38395 cgctaccccgtgctcgcctccatctacgcgggtccgttcaccaccgttgtcccctggagc
                    ************************************************************

MxDZ2_Tam_RS12675   ttcctggggacttcggattcgctgaccgccagcggcctggcgcaactgggcttcatcgtg
MxDZ2_Nan_RS30200   ttcctggggacttcggattcgctgaccgccagcggcctggcgcaactgggcttcatcgtg
MxDZ2_Kirby_RS38395 ttcctggggacttcggattcgctgaccgccagcggcctggcgcaactgggcttcatcgtg
                    ************************************************************

MxDZ2_Tam_RS12675   gtgttgctggacccccggggtggccccggacggagcaagtccttccaggacgcgaactac
MxDZ2_Nan_RS30200   gtgttgctggacccccggggtggccccggacggagcaagtccttccaggacgcgaactac
MxDZ2_Kirby_RS38395 gtgttgctggacccccggggtggccccggacggagcaagtccttccaggacgcgaactac
                    ************************************************************

MxDZ2_Tam_RS12675   ggccgcgtgggccagacggagattcccgactacgtcgcggggctgaagcaggccgcgtcc
MxDZ2_Nan_RS30200   ggccgcgtgggccagacggagattcccgactacgtcgcggggctgaagcaggccgcgtcc
MxDZ2_Kirby_RS38395 ggccgcgtgggccagacggagattcccgactacgtcgcggggctgaagcaggccgcgtcc
                    ************************************************************


MxDZ2_Tam_RS12675   acgcgcccctggatggacctggagcgggtcggcatccacggcggttcgtggggcggctat
MxDZ2_Nan_RS30200   acgcgcccctggatggacctggagcgggtcggcatccacggcggttcgtggggcggctat
MxDZ2_Kirby_RS38395 acgcgcccctggatggacctggagcgggtcggcatccacggcggttcgtggggcggctat
                    ************************************************************


MxDZ2_Tam_RS12675   ttcacgctgcgcggcatgctgacggcgccggacttcttcaaggcgggctacgccggagcc
MxDZ2_Nan_RS30200   ttcacgctgcgcggcatgctgacggcgccggacttcttcaaggcgggctacgccggagcc
MxDZ2_Kirby_RS38395 ttcacgctgcgcggcatgctgacggcgccggacttcttcaaggcgggctacgccggagcc
                    ************************************************************

MxDZ2_Tam_RS12675   cccggagcgctggaggaagaggccatcatcaacgagccctatctcaacctcccaagcgtc
MxDZ2_Nan_RS30200   cccggagcgctggaggaagaggccatcatcaacgagccctatctcaacctcccaagcgtc
MxDZ2_Kirby_RS38395 cccggagcgctggaggaagaggccatcatcaacgagccctatctcaacctcccaagcgtc
                    ************************************************************

MxDZ2_Tam_RS12675   aatccccaggggtacgcggcgggcgacaacctcgcgattgcagacaggctgaggggccac
MxDZ2_Nan_RS30200   aatccccaggggtacgcggcgggcgacaacctcgcgattgcagacaggctgaggggccac
MxDZ2_Kirby_RS38395 aatccccaggggtacgcggcgggcgacaacctcgcgattgcagacaggctgaggggccac
                    ************************************************************

MxDZ2_Tam_RS12675   ctgaagctgatgcacggcacgagcgacgtgaatgccacgctctccgtgacgatgcggatg
MxDZ2_Nan_RS30200   ctgaagctgatgcacggcacgagcgacgtgaatgccacgctctccgtgacgatgcggatg
MxDZ2_Kirby_RS38395 ctgaagctgatgcacggcacgagcgacgtgaatgccacgctctccgtgacgatgcggatg
                    ************************************************************

MxDZ2_Tam_RS12675   gcggacgcgctcatccgcgcgggcaagcgcttcgaattgctcatcatgcccggccagccg
MxDZ2_Nan_RS30200   gcggacgcgctcatccgcgcgggcaagcgcttcgaattgctcatcatgcccggccagccg
MxDZ2_Kirby_RS38395 gcggacgcgctcatccgcgcgggcaagcgcttcgaattgctcatcatgcccggccagccg
                    ************************************************************

MxDZ2_Tam_RS12675   cactcccctcggggtgccgccagccgctactaccgggacgatgtgggcctgttcttcctc
MxDZ2_Nan_RS30200   cactcccctcggggtgccgccagccgctactaccgggacgatgtgggcctgttcttcctc
MxDZ2_Kirby_RS38395 cactcccctcggggtgccgccagccgctactaccgggacgatgtgggcctgttcttcctc
                    ************************************************************

MxDZ2_Tam_RS12675   cggaccctgggcggtccacggtaa
MxDZ2_Nan_RS30200   cggaccctgggcggtccacggtaa
MxDZ2_Kirby_RS38395 cggaccctgggcggtccacggtaa
                    ************************
```

**Non-coding region mutation in MxDZ2_Nan 2**

| **Tam_location** | **Tam_nucleotide** | **Kirby nucleotide** | **Nan_nucleotide** | **Nan_location** |
| --- | --- | --- | --- | --- |
| **2988905** | **C** | **C** | **.** | **2988888** |

```
multiple sequence alignment (nucleotide)

MxDZ2_Kirby_RS38320 atgaaaaccctccccttgttgaccctggcctcggcgctggccttcaccccggcgctcgcc
MxDZ2_Nan_RS25680   atgaaaaccctccccttgttgaccctggcctcggcgctggccttcaccccggcgctcgcc
MxDZ2_Tam_RS17205   atgaaaaccctccccttgttgaccctggcctcggcgctggccttcaccccggcgctcgcc
                    ************************************************************

MxDZ2_Kirby_RS38320 gcggcgccgaagcctcctccccCcgcgaagtccgaacccacgagcgagaagaagcccatg
MxDZ2_Nan_RS25680   gcggcgccgaagcctcctcccc-cgcgaagtccgaacccacgagcgagaagaagcccatg
MxDZ2_Tam_RS17205   gcggcgccgaagcctcctccccCcgcgaagtccgaacccacgagcgagaagaagcccatg
                    ********************** *************************************

MxDZ2_Kirby_RS38320 tccttccatcacctctccgccaaccggctcgacggcaagccggagacgctgtccggctac
MxDZ2_Nan_RS25680   tccttccatcacctctccgccaaccggctcgacggcaagccggagacgctgtccggctac
MxDZ2_Tam_RS17205   tccttccatcacctctccgccaaccggctcgacggcaagccggagacgctgtccggctac
                    ************************************************************

MxDZ2_Kirby_RS38320 cagggcaaggtcgtcctggtggtgaacaccgcgtcggaatgtggctacacgccgcagtac
MxDZ2_Nan_RS25680   cagggcaaggtcgtcctggtggtgaacaccgcgtcggaatgtggctacacgccgcagtac
MxDZ2_Tam_RS17205   cagggcaaggtcgtcctggtggtgaacaccgcgtcggaatgtggctacacgccgcagtac
                    ************************************************************

MxDZ2_Kirby_RS38320 gcgggcttggagaagctccaccaggaatacaaggacaagggcctggtggtggtgggcttc
MxDZ2_Nan_RS25680   gcgggcttggagaagctccaccaggaatacaaggacaagggcctggtggtggtgggcttc
MxDZ2_Tam_RS17205   gcgggcttggagaagctccaccaggaatacaaggacaagggcctggtggtggtgggcttc
                    ************************************************************

MxDZ2_Kirby_RS38320 ccgtccaacgactttggcggccaggagccgggctcgtcggaggagatcaagaagttctgt
MxDZ2_Nan_RS25680   ccgtccaacgactttggcggccaggagccgggctcgtcggaggagatcaagaagttctgt
MxDZ2_Tam_RS17205   ccgtccaacgactttggcggccaggagccgggctcgtcggaggagatcaagaagttctgt
                    ************************************************************

MxDZ2_Kirby_RS38320 gagctccgctacaaggtcaccttccccatgttcgagaaggtgaagacgaagggggacggc
MxDZ2_Nan_RS25680   gagctccgctacaaggtcaccttccccatgttcgagaaggtgaagacgaagggggacggc
MxDZ2_Tam_RS17205   gagctccgctacaaggtcaccttccccatgttcgagaaggtgaagacgaagggggacggc
                    ************************************************************

MxDZ2_Kirby_RS38320 cagtcgcccgtctacgcgttcctgtcgcagcaccaccccgcgcccaagtggaatttccac
MxDZ2_Nan_RS25680   cagtcgcccgtctacgcgttcctgtcgcagcaccaccccgcgcccaagtggaatttccac
MxDZ2_Tam_RS17205   cagtcgcccgtctacgcgttcctgtcgcagcaccaccccgcgcccaagtggaatttccac
                    ************************************************************

MxDZ2_Kirby_RS38320 aagtacgtggtgggcaaggacggccaggtgaaggccggcttccccagtgcggtgacgccg
MxDZ2_Nan_RS25680   aagtacgtggtgggcaaggacggccaggtgaaggccggcttccccagtgcggtgacgccg
MxDZ2_Tam_RS17205   aagtacgtggtgggcaaggacggccaggtgaaggccggcttccccagtgcggtgacgccg
                    ************************************************************

MxDZ2_Kirby_RS38320 gacagcgcggagctgaaggcggccatcgacagcgcgctcgcgcagccatag
MxDZ2_Nan_RS25680   gacagcgcggagctgaaggcggccatcgacagcgcgctcgcgcagccatag
MxDZ2_Tam_RS17205   gacagcgcggagctgaaggcggccatcgacagcgcgctcgcgcagccatag
                    ***************************************************
```

**Non-coding region mutation in MxDZ2_Nan 3**

| **Tam_location** | **Tam_nucleotide** | **Kirby nucleotide** | **Nan_nucleotide** | **Nan_location** |
| --- | --- | --- | --- | --- |
| **4124897** | **G** | **G** | **.** | **4124877** |

```
multiple sequence alignment (nucleotide)

MxDZ2_Kirby_RS0230030 atgaaggtggttcttcgtttcggtgcgctggcggtgggcgcggtgctcatcactggcggg
MxDZ2_Tam_RS22030     atgaaggtggttcttcgtttcggtgcgctggcggtgggcgcggtgctcatcactggcggg
MxDZ2_Nan_RS20830     atgaaggtggttcttcgtttcggtgcgctggcggtgggcgcggtgctcatcactggcggg
                      ************************************************************

MxDZ2_Kirby_RS0230030 gtgggagaggcggcggaaacgcaggcccgcaagggggGgaagaagcccgctgcggcgtcc
MxDZ2_Tam_RS22030     gtgggagaggcggcggaaacgcaggcccgcaagggggGgaagaagcccgctgcggcgtcc
MxDZ2_Nan_RS20830     gtgggagaggcggcggaaacgcaggcccgcaaggggg-gaagaagcccgctgcggcgtcc
                      ************************************* **********************

MxDZ2_Kirby_RS0230030 gcgtcgaagacctccggtgcgtccagcaaggcgggcgggaaaaagaagtccgcgaaggcc
MxDZ2_Tam_RS22030     gcgtcgaagacctccggtgcgtccagcaaggcgggcgggaaaaagaagtccgcgaaggcc
MxDZ2_Nan_RS20830     gcgtcgaagacctccggtgcgtccagcaaggcgggcgggaaaaagaagtccgcgaaggcc
                      ************************************************************

MxDZ2_Kirby_RS0230030 caggtcgaccgcaaggcagaggagaaggctccgccgccgggggttgcgccggaggacgtg
MxDZ2_Tam_RS22030     caggtcgaccgcaaggcagaggagaaggctccgccgccgggggttgcgccggaggacgtg
MxDZ2_Nan_RS20830     caggtcgaccgcaaggcagaggagaaggctccgccgccgggggttgcgccggaggacgtg
                      ************************************************************

MxDZ2_Kirby_RS0230030 cggcaggggccggcgcgcgttcagcccgcgtcggcgaagttcgcggagctgccccgcatc
MxDZ2_Tam_RS22030     cggcaggggccggcgcgcgttcagcccgcgtcggcgaagttcgcggagctgccccgcatc
MxDZ2_Nan_RS20830     cggcaggggccggcgcgcgttcagcccgcgtcggcgaagttcgcggagctgccccgcatc
                      ************************************************************

MxDZ2_Kirby_RS0230030 ccggacgccaagcgggacgcgctggcggacaagaagcgcgacgaggccattgccgccttc
MxDZ2_Tam_RS22030     ccggacgccaagcgggacgcgctggcggacaagaagcgcgacgaggccattgccgccttc
MxDZ2_Nan_RS20830     ccggacgccaagcgggacgcgctggcggacaagaagcgcgacgaggccattgccgccttc
                      ************************************************************

MxDZ2_Kirby_RS0230030 aagcgcctcatccccaagctgcgggacggcaatccgcagaaggcggagatgctctaccgc
MxDZ2_Tam_RS22030     aagcgcctcatccccaagctgcgggacggcaatccgcagaaggcggagatgctctaccgc
MxDZ2_Nan_RS20830     aagcgcctcatccccaagctgcgggacggcaatccgcagaaggcggagatgctctaccgc
                      ************************************************************

MxDZ2_Kirby_RS0230030 ctgtcggagctctactgggagaagtccaagtacctctaccagttggagatgacgcgcttc
MxDZ2_Tam_RS22030     ctgtcggagctctactgggagaagtccaagtacctctaccagttggagatgacgcgcttc
MxDZ2_Nan_RS20830     ctgtcggagctctactgggagaagtccaagtacctctaccagttggagatgacgcgcttc
                      ************************************************************

MxDZ2_Kirby_RS0230030 ctcgcggcggagaaggaatacgacgcggccgtggcgcgcggcgagaaggtggagccgccc
MxDZ2_Tam_RS22030     ctcgcggcggagaaggaatacgacgcggccgtggcgcgcggcgagaaggtggagccgccc
MxDZ2_Nan_RS20830     ctcgcggcggagaaggaatacgacgcggccgtggcgcgcggcgagaaggtggagccgccc
                      ************************************************************

MxDZ2_Kirby_RS0230030 aagaagaaccacgcggacagcgagcgctaccgcaccgaaacgatgggcatctacgaggac
MxDZ2_Tam_RS22030     aagaagaaccacgcggacagcgagcgctaccgcaccgaaacgatgggcatctacgaggac
MxDZ2_Nan_RS20830     aagaagaaccacgcggacagcgagcgctaccgcaccgaaacgatgggcatctacgaggac
                      ************************************************************

MxDZ2_Kirby_RS0230030 atcctccgcgcgtacccggattatccccagcgcgacgaggtcctcttctccatggggtac
MxDZ2_Tam_RS22030     atcctccgcgcgtacccggattatccccagcgcgacgaggtcctcttctccatggggtac
MxDZ2_Nan_RS20830     atcctccgcgcgtacccggattatccccagcgcgacgaggtcctcttctccatggggtac
                      ************************************************************

MxDZ2_Kirby_RS0230030 aactactacgagctgggacgccgcgaggacgcggtggcccgctacgaggagttgatccgc
MxDZ2_Tam_RS22030     aactactacgagctgggacgccgcgaggacgcggtggcccgctacgaggagttgatccgc
MxDZ2_Nan_RS20830     aactactacgagctgggacgccgcgaggacgcggtggcccgctacgaggagttgatccgc
                      ************************************************************

MxDZ2_Kirby_RS0230030 gacttcccgaagtcgcagttcgtgccggacgcgtacatccagctcggcaaccactacttc
MxDZ2_Tam_RS22030     gacttcccgaagtcgcagttcgtgccggacgcgtacatccagctcggcaaccactacttc
MxDZ2_Nan_RS20830     gacttcccgaagtcgcagttcgtgccggacgcgtacatccagctcggcaaccactacttc
                      ************************************************************

MxDZ2_Kirby_RS0230030 gagaacaacaagctcatccccgccaaggagaactatgagaaggcgcgggactcgggcgtg
MxDZ2_Tam_RS22030     gagaacaacaagctcatccccgccaaggagaactatgagaaggcgcgggactcgggcgtg
MxDZ2_Nan_RS20830     gagaacaacaagctcatccccgccaaggagaactatgagaaggcgcgggactcgggcgtg
                      ************************************************************

MxDZ2_Kirby_RS0230030 ccgaaaatctacggctacgccgtctacaagctgtcctggtgcgactacaacaccggcgac
MxDZ2_Tam_RS22030     ccgaaaatctacggctacgccgtctacaagctgtcctggtgcgactacaacaccggcgac
MxDZ2_Nan_RS20830     ccgaaaatctacggctacgccgtctacaagctgtcctggtgcgactacaacaccggcgac
                      ************************************************************

MxDZ2_Kirby_RS0230030 tacgagctggggctgaagaagctccacgaggtggtggactacgccgcgaagagccctgag
MxDZ2_Tam_RS22030     tacgagctggggctgaagaagctccacgaggtggtggactacgccgcgaagagccctgag
MxDZ2_Nan_RS20830     tacgagctggggctgaagaagctccacgaggtggtggactacgccgcgaagagccctgag
                      ************************************************************

MxDZ2_Kirby_RS0230030 ttgggtgacctgcgcaccgaggcgctcaacgacctgaccgtcttctacgtccagttggac
MxDZ2_Tam_RS22030     ttgggtgacctgcgcaccgaggcgctcaacgacctgaccgtcttctacgtccagttggac
MxDZ2_Nan_RS20830     ttgggtgacctgcgcaccgaggcgctcaacgacctgaccgtcttctacgtccagttggac
                      ************************************************************

MxDZ2_Kirby_RS0230030 cagccgaaggaagccatcgcctacttcaaggagaaggcgccggcgcagcgcgtgggccgc
MxDZ2_Tam_RS22030     cagccgaaggaagccatcgcctacttcaaggagaaggcgccggcgcagcgcgtgggccgc
MxDZ2_Nan_RS20830     cagccgaaggaagccatcgcctacttcaaggagaaggcgccggcgcagcgcgtgggccgc
                      ************************************************************

MxDZ2_Kirby_RS0230030 ctgctggccaagacggccgcgggcctggtggacgcgggccacttcgacagcgccatcctc
MxDZ2_Tam_RS22030     ctgctggccaagacggccgcgggcctggtggacgcgggccacttcgacagcgccatcctc
MxDZ2_Nan_RS20830     ctgctggccaagacggccgcgggcctggtggacgcgggccacttcgacagcgccatcctc
                      ************************************************************

MxDZ2_Kirby_RS0230030 gcgtaccgcacgctcgtggacgacgagcccatgggcgccaacgcgccggagtaccagcag
MxDZ2_Tam_RS22030     gcgtaccgcacgctcgtggacgacgagcccatgggcgccaacgcgccggagtaccagcag
MxDZ2_Nan_RS20830     gcgtaccgcacgctcgtggacgacgagcccatgggcgccaacgcgccggagtaccagcag
                      ************************************************************

MxDZ2_Kirby_RS0230030 gccatcgtccgcgcccacgaggggctccgccagcgccagttggtccgcaaggaaatgaag
MxDZ2_Tam_RS22030     gccatcgtccgcgcccacgaggggctccgccagcgccagttggtccgcaaggaaatgaag
MxDZ2_Nan_RS20830     gccatcgtccgcgcccacgaggggctccgccagcgccagttggtccgcaaggaaatgaag
                      ************************************************************

MxDZ2_Kirby_RS0230030 cggatggtggacctctacagccctggtggcgggtggtggaaggccaacgagggcaagacg
MxDZ2_Tam_RS22030     cggatggtggacctctacagccctggtggcgggtggtggaaggccaacgagggcaagacg
MxDZ2_Nan_RS20830     cggatggtggacctctacagccctggtggcgggtggtggaaggccaacgagggcaagacg
                      ************************************************************

MxDZ2_Kirby_RS0230030 gccgtcctgcgaaacgccttcaacgtcactgaagaggccatgcgcgtcatggtcaccgag
MxDZ2_Tam_RS22030     gccgtcctgcgaaacgccttcaacgtcactgaagaggccatgcgcgtcatggtcaccgag
MxDZ2_Nan_RS20830     gccgtcctgcgaaacgccttcaacgtcactgaagaggccatgcgcgtcatggtcaccgag
                      ************************************************************


MxDZ2_Kirby_RS0230030 taccaccaggaggcgcagaagacgcgccaggtggagacctaccggctggcgcgtgacatc
MxDZ2_Tam_RS22030     taccaccaggaggcgcagaagacgcgccaggtggagacctaccggctggcgcgtgacatc
MxDZ2_Nan_RS20830     taccaccaggaggcgcagaagacgcgccaggtggagacctaccggctggcgcgtgacatc
                      ************************************************************

MxDZ2_Kirby_RS0230030 tacaagcagtacgtggacgcgttcgcctccaacgcgaacccggacttcgtggcggactcc
MxDZ2_Tam_RS22030     tacaagcagtacgtggacgcgttcgcctccaacgcgaacccggacttcgtggcggactcc
MxDZ2_Nan_RS20830     tacaagcagtacgtggacgcgttcgcctccaacgcgaacccggacttcgtggcggactcc
                      ************************************************************

MxDZ2_Kirby_RS0230030 gccttcaacctccgcttcttctacgcggagatcctctgggctttggaggagtgggaagcg
MxDZ2_Tam_RS22030     gccttcaacctccgcttcttctacgcggagatcctctgggctttggaggagtgggaagcg
MxDZ2_Nan_RS20830     gccttcaacctccgcttcttctacgcggagatcctctgggctttggaggagtgggaagcg
                      ************************************************************

MxDZ2_Kirby_RS0230030 gccgcggccgagtacgacgcggtggtggccttcaagattccggaccgcgacaccgcgcgc
MxDZ2_Tam_RS22030     gccgcggccgagtacgacgcggtggtggccttcaagattccggaccgcgacaccgcgcgc
MxDZ2_Nan_RS20830     gccgcggccgagtacgacgcggtggtggccttcaagattccggaccgcgacaccgcgcgc
                      ************************************************************

MxDZ2_Kirby_RS0230030 gaggtctccaacgaggcgtaccgcaagagcgccgggtacaacgccatcctcgcctacgac
MxDZ2_Tam_RS22030     gaggtctccaacgaggcgtaccgcaagagcgccgggtacaacgccatcctcgcctacgac
MxDZ2_Nan_RS20830     gaggtctccaacgaggcgtaccgcaagagcgccgggtacaacgccatcctcgcctacgac
                      ************************************************************

MxDZ2_Kirby_RS0230030 aagctggtgaagatcgagcgaggccagctcgccaagagcgacctgcgcgacggccagaag
MxDZ2_Tam_RS22030     aagctggtgaagatcgagcgaggccagctcgccaagagcgacctgcgcgacggccagaag
MxDZ2_Nan_RS20830     aagctggtgaagatcgagcgaggccagctcgccaagagcgacctgcgcgacggccagaag
                      ************************************************************

MxDZ2_Kirby_RS0230030 gtcgacgagaagaaggacaagggcgacgtcgccaagcagaagatcgtcaagcgcgacgcg
MxDZ2_Tam_RS22030     gtcgacgagaagaaggacaagggcgacgtcgccaagcagaagatcgtcaagcgcgacgcg
MxDZ2_Nan_RS20830     gtcgacgagaagaaggacaagggcgacgtcgccaagcagaagatcgtcaagcgcgacgcg
                      ************************************************************

MxDZ2_Kirby_RS0230030 aaggaccgccaggaagaggcgctcacgaagttcgaggaccggctggtcgccgcgtgtgac
MxDZ2_Tam_RS22030     aaggaccgccaggaagaggcgctcacgaagttcgaggaccggctggtcgccgcgtgtgac
MxDZ2_Nan_RS20830     aaggaccgccaggaagaggcgctcacgaagttcgaggaccggctggtcgccgcgtgtgac
                      ************************************************************


MxDZ2_Kirby_RS0230030 gtctatgtgaagctgtatccgaacacgcaggacgaaatcgacctgcgctaccaggccgcc
MxDZ2_Tam_RS22030     gtctatgtgaagctgtatccgaacacgcaggacgaaatcgacctgcgctaccaggccgcc
MxDZ2_Nan_RS20830     gtctatgtgaagctgtatccgaacacgcaggacgaaatcgacctgcgctaccaggccgcc
                      ************************************************************

MxDZ2_Kirby_RS0230030 gtcatcctctatgaccgcagccacttcgtggacgcggcccggcgcttcggcgaaatcatc
MxDZ2_Tam_RS22030     gtcatcctctatgaccgcagccacttcgtggacgcggcccggcgcttcggcgaaatcatc
MxDZ2_Nan_RS20830     gtcatcctctatgaccgcagccacttcgtggacgcggcccggcgcttcggcgaaatcatc
                      ************************************************************

MxDZ2_Kirby_RS0230030 gagaagttccccgaggagcgccgctcgcgcgacgcggccgacctcaccatgtacgtgctg
MxDZ2_Tam_RS22030     gagaagttccccgaggagcgccgctcgcgcgacgcggccgacctcaccatgtacgtgctg
MxDZ2_Nan_RS20830     gagaagttccccgaggagcgccgctcgcgcgacgcggccgacctcaccatgtacgtgctg
                      ************************************************************

MxDZ2_Kirby_RS0230030 gagagccgcgaggagtggctcgagctgaacacgctgtcgaagaagttcctggagaacaag
MxDZ2_Tam_RS22030     gagagccgcgaggagtggctcgagctgaacacgctgtcgaagaagttcctggagaacaag
MxDZ2_Nan_RS20830     gagagccgcgaggagtggctcgagctgaacacgctgtcgaagaagttcctggagaacaag
                      ************************************************************

MxDZ2_Kirby_RS0230030 aagctggccaagcccggaacggacttcgccgtgcgcgtcagccgcgtcgtcgaaggcagc
MxDZ2_Tam_RS22030     aagctggccaagcccggaacggacttcgccgtgcgcgtcagccgcgtcgtcgaaggcagc
MxDZ2_Nan_RS20830     aagctggccaagcccggaacggacttcgccgtgcgcgtcagccgcgtcgtcgaaggcagc
                      ************************************************************

MxDZ2_Kirby_RS0230030 cagtacaagtgggtggacgaggtcgtctacaagaaggagaagaacccgaagaaggccgcc
MxDZ2_Tam_RS22030     cagtacaagtgggtggacgaggtcgtctacaagaaggagaagaacccgaagaaggccgcc
MxDZ2_Nan_RS20830     cagtacaagtgggtggacgaggtcgtctacaagaaggagaagaacccgaagaaggccgcc
                      ************************************************************

MxDZ2_Kirby_RS0230030 gaggagttcctccgcttcgtgtccgacttccccaagtcagagaacgcggaccgtgcgctc
MxDZ2_Tam_RS22030     gaggagttcctccgcttcgtgtccgacttccccaagtcagagaacgcggaccgtgcgctc
MxDZ2_Nan_RS20830     gaggagttcctccgcttcgtgtccgacttccccaagtcagagaacgcggaccgtgcgctc
                      ************************************************************

MxDZ2_Kirby_RS0230030 acttacgcgatggtcatcgcgcaggaggcgggcgaaatcgacaagggcctggccgcgggt
MxDZ2_Tam_RS22030     acttacgcgatggtcatcgcgcaggaggcgggcgaaatcgacaagggcctggccgcgggt
MxDZ2_Nan_RS20830     acttacgcgatggtcatcgcgcaggaggcgggcgaaatcgacaagggcctggccgcgggt
                      ************************************************************


MxDZ2_Kirby_RS0230030 gagcgcttcctcaaggagtacccgcgcagccccttcgagctgaaggcgcgttactcgctg
MxDZ2_Tam_RS22030     gagcgcttcctcaaggagtacccgcgcagccccttcgagctgaaggcgcgttactcgctg
MxDZ2_Nan_RS20830     gagcgcttcctcaaggagtacccgcgcagccccttcgagctgaaggcgcgttactcgctg
                      ************************************************************

MxDZ2_Kirby_RS0230030 gcgggcctctacgagaaggtcgctgagtaccggaaggccgccgtcatggcggagtccctc
MxDZ2_Tam_RS22030     gcgggcctctacgagaaggtcgctgagtaccggaaggccgccgtcatggcggagtccctc
MxDZ2_Nan_RS20830     gcgggcctctacgagaaggtcgctgagtaccggaaggccgccgtcatggcggagtccctc
                      ************************************************************

MxDZ2_Kirby_RS0230030 gtcgccagctacgacgccgcgatgaaggcggacgatgccaacggcaagcgcaaggcgacc
MxDZ2_Tam_RS22030     gtcgccagctacgacgccgcgatgaaggcggacgatgccaacggcaagcgcaaggcgacc
MxDZ2_Nan_RS20830     gtcgccagctacgacgccgcgatgaaggcggacgatgccaacggcaagcgcaaggcgacc
                      ************************************************************

MxDZ2_Kirby_RS0230030 aaggcggccgccaaggtgagcgtcgcgccgggcgccgaggacgcggagtccaagcgcgag
MxDZ2_Tam_RS22030     aaggcggccgccaaggtgagcgtcgcgccgggcgccgaggacgcggagtccaagcgcgag
MxDZ2_Nan_RS20830     aaggcggccgccaaggtgagcgtcgcgccgggcgccgaggacgcggagtccaagcgcgag
                      ************************************************************

MxDZ2_Kirby_RS0230030 cgggtggccgccgagcgcaaggcgctgctggaagaggccggcggctggatggcggatgcg
MxDZ2_Tam_RS22030     cgggtggccgccgagcgcaaggcgctgctggaagaggccggcggctggatggcggatgcg
MxDZ2_Nan_RS20830     cgggtggccgccgagcgcaaggcgctgctggaagaggccggcggctggatggcggatgcg
                      ************************************************************

MxDZ2_Kirby_RS0230030 cagttcaatgcgggcgtctggtgggaaggcgcgggtgagccgcagaaggcggtggctgcc
MxDZ2_Tam_RS22030     cagttcaatgcgggcgtctggtgggaaggcgcgggtgagccgcagaaggcggtggctgcc
MxDZ2_Nan_RS20830     cagttcaatgcgggcgtctggtgggaaggcgcgggtgagccgcagaaggcggtggctgcc
                      ************************************************************

MxDZ2_Kirby_RS0230030 tacaacacgtacgtctcccgcttcaaggaccgcaaggacgtgccgcaggtggccttcgcg
MxDZ2_Tam_RS22030     tacaacacgtacgtctcccgcttcaaggaccgcaaggacgtgccgcaggtggccttcgcg
MxDZ2_Nan_RS20830     tacaacacgtacgtctcccgcttcaaggaccgcaaggacgtgccgcaggtggccttcgcg
                      ************************************************************

MxDZ2_Kirby_RS0230030 gcggcgctcgcgtgggagaaggagaagaagtggagcgaggcggcccgggcgttcggcgcc
MxDZ2_Tam_RS22030     gcggcgctcgcgtgggagaaggagaagaagtggagcgaggcggcccgggcgttcggcgcc
MxDZ2_Nan_RS20830     gcggcgctcgcgtgggagaaggagaagaagtggagcgaggcggcccgggcgttcggcgcc
                      ************************************************************


MxDZ2_Kirby_RS0230030 ttcgcggagacgtacggccgtgactcgcgctccagctcggcgcaggtgtaccaggcgcgc
MxDZ2_Tam_RS22030     ttcgcggagacgtacggccgtgactcgcgctccagctcggcgcaggtgtaccaggcgcgc
MxDZ2_Nan_RS20830     ttcgcggagacgtacggccgtgactcgcgctccagctcggcgcaggtgtaccaggcgcgc
                      ************************************************************

MxDZ2_Kirby_RS0230030 taccacgagctgctggcgtacgagcacctgaggaacgcacgcgagcaggagcgcgtgcag
MxDZ2_Tam_RS22030     taccacgagctgctggcgtacgagcacctgaggaacgcacgcgagcaggagcgcgtgcag
MxDZ2_Nan_RS20830     taccacgagctgctggcgtacgagcacctgaggaacgcacgcgagcaggagcgcgtgcag
                      ************************************************************

MxDZ2_Kirby_RS0230030 ggcgagctggtgcgggcgtggaaccggctgccggagagtgctcgcaaggacgcggcggtg
MxDZ2_Tam_RS22030     ggcgagctggtgcgggcgtggaaccggctgccggagagtgctcgcaaggacgcggcggtg
MxDZ2_Nan_RS20830     ggcgagctggtgcgggcgtggaaccggctgccggagagtgctcgcaaggacgcggcggtg
                      ************************************************************

MxDZ2_Kirby_RS0230030 ctcaatgcttacggccatgcgcgcttcctgtcgctggagccggcgtggaagcgttacgtg
MxDZ2_Tam_RS22030     ctcaatgcttacggccatgcgcgcttcctgtcgctggagccggcgtggaagcgttacgtg
MxDZ2_Nan_RS20830     ctcaatgcttacggccatgcgcgcttcctgtcgctggagccggcgtggaagcgttacgtg
                      ************************************************************

MxDZ2_Kirby_RS0230030 ggcatccgcttctcgcgggtgagcaccatccgccgggacctggcggcgaagcagaaggag
MxDZ2_Tam_RS22030     ggcatccgcttctcgcgggtgagcaccatccgccgggacctggcggcgaagcagaaggag
MxDZ2_Nan_RS20830     ggcatccgcttctcgcgggtgagcaccatccgccgggacctggcggcgaagcagaaggag
                      ************************************************************

MxDZ2_Kirby_RS0230030 attcagcggctggagaaggagtacctcgccgtcctgtccaccggctccggtgattggggc
MxDZ2_Tam_RS22030     attcagcggctggagaaggagtacctcgccgtcctgtccaccggctccggtgattggggc
MxDZ2_Nan_RS20830     attcagcggctggagaaggagtacctcgccgtcctgtccaccggctccggtgattggggc
                      ************************************************************

MxDZ2_Kirby_RS0230030 atcgcggcgctcacgcgcatcggcctggcctatgccgacttcgcgcgcaacatcatggac
MxDZ2_Tam_RS22030     atcgcggcgctcacgcgcatcggcctggcctatgccgacttcgcgcgcaacatcatggac
MxDZ2_Nan_RS20830     atcgcggcgctcacgcgcatcggcctggcctatgccgacttcgcgcgcaacatcatggac
                      ************************************************************

MxDZ2_Kirby_RS0230030 tcgccggacccgtccgggctcgatgaggagcagctcgccatgtaccgcagcgagctggag
MxDZ2_Tam_RS22030     tcgccggacccgtccgggctcgatgaggagcagctcgccatgtaccgcagcgagctggag
MxDZ2_Nan_RS20830     tcgccggacccgtccgggctcgatgaggagcagctcgccatgtaccgcagcgagctggag
                      ************************************************************


MxDZ2_Kirby_RS0230030 aacctggcgttgccgctggaggacaaagccgccgaggccctggagaaggccctggagaag
MxDZ2_Tam_RS22030     aacctggcgttgccgctggaggacaaagccgccgaggccctggagaaggccctggagaag
MxDZ2_Nan_RS20830     aacctggcgttgccgctggaggacaaagccgccgaggccctggagaaggccctggagaag
                      ************************************************************

MxDZ2_Kirby_RS0230030 gcctacgagctgggcgtctacagcccgtggaccctggccgcgcaggaccaggtgaaccgc
MxDZ2_Tam_RS22030     gcctacgagctgggcgtctacagcccgtggaccctggccgcgcaggaccaggtgaaccgc
MxDZ2_Nan_RS20830     gcctacgagctgggcgtctacagcccgtggaccctggccgcgcaggaccaggtgaaccgc
                      ************************************************************

MxDZ2_Kirby_RS0230030 ctgcgtccgggggcctacgcacaggtgcggcaggtggactaccgcggcagcgacacactc
MxDZ2_Tam_RS22030     ctgcgtccgggggcctacgcacaggtgcggcaggtggactaccgcggcagcgacacactc
MxDZ2_Nan_RS20830     ctgcgtccgggggcctacgcacaggtgcggcaggtggactaccgcggcagcgacacactc
                      ************************************************************

MxDZ2_Kirby_RS0230030 gtccgctcggacctggtgcgcgtgctggaaggcgccaccgcgacgacgccggccccggcg
MxDZ2_Tam_RS22030     gtccgctcggacctggtgcgcgtgctggaaggcgccaccgcgacgacgccggccccggcg
MxDZ2_Nan_RS20830     gtccgctcggacctggtgcgcgtgctggaaggcgccaccgcgacgacgccggccccggcg
                      ************************************************************

MxDZ2_Kirby_RS0230030 gactcctcgaagccctcggatgacgaggcgcaggcacccacggcggcgcgcggggaggtg
MxDZ2_Tam_RS22030     gactcctcgaagccctcggatgacgaggcgcaggcacccacggcggcgcgcggggaggtg
MxDZ2_Nan_RS20830     gactcctcgaagccctcggatgacgaggcgcaggcacccacggcggcgcgcggggaggtg
                      ************************************************************

MxDZ2_Kirby_RS0230030 ctgcgatga
MxDZ2_Tam_RS22030     ctgcgatga
MxDZ2_Nan_RS20830     ctgcgatga
                      *********
```

**Non-coding region mutation in MxDZ2_Nan 4**

| **Tam_location** | **Tam_nucleotide** | **Kirby nucleotide** | **Nan_nucleotide** | **Nan_location** |
| --- | --- | --- | --- | --- |
| **5287207** | **C (complement)** | **C (complement)** | **.** | **5287185** |

```
multiple sequence alignment (nucleotide)

MxDZ2_Kirby_RS0232095 gtgtcatcattgagaagctggagctatctccccatcaccgccctcgctctcgcgctggtc
MxDZ2_Nan_RS17045     gtgtcatcattgagaagctggagctatctccccatcaccgccctcgctctcgcgctggtc
MxDZ2_Tam_RS25820     gtgtcatcattgagaagctggagctatctccccatcaccgccctcgctctcgcgctggtc
                      ************************************************************

MxDZ2_Kirby_RS0232095 cttccggagcccgcgaaggcgcaggctgacggaacctatccatccccgGgggctgcgcag
MxDZ2_Nan_RS17045     cttccggagcccgcgaaggcgcaggctgacggaacctatccatccccg-gggctgcgcag
MxDZ2_Tam_RS25820     cttccggagcccgcgaaggcgcaggctgacggaacctatccatccccgGgggctgcgcag
                      ************************************************ ***********

MxDZ2_Kirby_RS0232095 ccgccgtctgcccccgcgggttatgcggcccctccgcagcccacgtatcgccagacggcg
MxDZ2_Nan_RS17045     ccgccgtctgcccccgcgggttatgcggcccctccgcagcccacgtatcgccagacggcg
MxDZ2_Tam_RS25820     ccgccgtctgcccccgcgggttatgcggcccctccgcagcccacgtatcgccagacggcg
                      ************************************************************

MxDZ2_Kirby_RS0232095 ccgcagcagccggcggatccgtacgcgcagccgccgcagacgcagcagcccgcggacccg
MxDZ2_Nan_RS17045     ccgcagcagccggcggatccgtacgcgcagccgccgcagacgcagcagcccgcggacccg
MxDZ2_Tam_RS25820     ccgcagcagccggcggatccgtacgcgcagccgccgcagacgcagcagcccgcggacccg
                      ************************************************************

MxDZ2_Kirby_RS0232095 tacgcgcagccgccggcgtccgcgcagcccgcggacccgtatgcgcagccgccgcagacg
MxDZ2_Nan_RS17045     tacgcgcagccgccggcgtccgcgcagcccgcggacccgtatgcgcagccgccgcagacg
MxDZ2_Tam_RS25820     tacgcgcagccgccggcgtccgcgcagcccgcggacccgtatgcgcagccgccgcagacg
                      ************************************************************

MxDZ2_Kirby_RS0232095 cagctgccggtggacccgtatgcgcagccgcagacgcagcagccgtcagcggatccgtat
MxDZ2_Nan_RS17045     cagctgccggtggacccgtatgcgcagccgcagacgcagcagccgtcagcggatccgtat
MxDZ2_Tam_RS25820     cagctgccggtggacccgtatgcgcagccgcagacgcagcagccgtcagcggatccgtat
                      ************************************************************

MxDZ2_Kirby_RS0232095 gcgcaaccgccgtcgtccacgcagccggtggacccgtacgcgcagccgccgcagacgcag
MxDZ2_Nan_RS17045     gcgcaaccgccgtcgtccacgcagccggtggacccgtacgcgcagccgccgcagacgcag
MxDZ2_Tam_RS25820     gcgcaaccgccgtcgtccacgcagccggtggacccgtacgcgcagccgccgcagacgcag
                      ************************************************************

MxDZ2_Kirby_RS0232095 cagccggtggacccatatgcgcagccgcagacgccgcagggctatcccccggcgcagaat
MxDZ2_Nan_RS17045     cagccggtggacccatatgcgcagccgcagacgccgcagggctatcccccggcgcagaat
MxDZ2_Tam_RS25820     cagccggtggacccatatgcgcagccgcagacgccgcagggctatcccccggcgcagaat
                      ************************************************************

MxDZ2_Kirby_RS0232095 gcctatcccccgtcggatgcgcgggacccggctcgctatccgggcgattccgcgacggcg
MxDZ2_Nan_RS17045     gcctatcccccgtcggatgcgcgggacccggctcgctatccgggcgattccgcgacggcg
MxDZ2_Tam_RS25820     gcctatcccccgtcggatgcgcgggacccggctcgctatccgggcgattccgcgacggcg
                      ************************************************************

MxDZ2_Kirby_RS0232095 agcgacgcgcccacgctgcctccggatccggactcgcttcggacgaagccgcagtcgccg
MxDZ2_Nan_RS17045     agcgacgcgcccacgctgcctccggatccggactcgcttcggacgaagccgcagtcgccg
MxDZ2_Tam_RS25820     agcgacgcgcccacgctgcctccggatccggactcgcttcggacgaagccgcagtcgccg
                      ************************************************************

MxDZ2_Kirby_RS0232095 ctgacgccggacgaccgggccatggcgaagcggggcgtgcctcccatcgaggacactggg
MxDZ2_Nan_RS17045     ctgacgccggacgaccgggccatggcgaagcggggcgtgcctcccatcgaggacactggg
MxDZ2_Tam_RS25820     ctgacgccggacgaccgggccatggcgaagcggggcgtgcctcccatcgaggacactggg
                      ************************************************************

MxDZ2_Kirby_RS0232095 gtgatgctggacggcaggccgcgccagggtccgttcctctccggtccgggcagcttcacc
MxDZ2_Nan_RS17045     gtgatgctggacggcaggccgcgccagggtccgttcctctccggtccgggcagcttcacc
MxDZ2_Tam_RS25820     gtgatgctggacggcaggccgcgccagggtccgttcctctccggtccgggcagcttcacc
                      ************************************************************

MxDZ2_Kirby_RS0232095 ttcatcgtgcaccacacgacgctgggcgcgctgggtggcttcttcacgcaggcgttcgcc
MxDZ2_Nan_RS17045     ttcatcgtgcaccacacgacgctgggcgcgctgggtggcttcttcacgcaggcgttcgcc
MxDZ2_Tam_RS25820     ttcatcgtgcaccacacgacgctgggcgcgctgggtggcttcttcacgcaggcgttcgcc
                      ************************************************************

MxDZ2_Kirby_RS0232095 agcgacttcaacttcgacaagagctcgcgagaggcaatgctcgcgggcacgctcatcggc
MxDZ2_Nan_RS17045     agcgacttcaacttcgacaagagctcgcgagaggcaatgctcgcgggcacgctcatcggc
MxDZ2_Tam_RS25820     agcgacttcaacttcgacaagagctcgcgagaggcaatgctcgcgggcacgctcatcggc
                      ************************************************************

MxDZ2_Kirby_RS0232095 gcgggcctgggcttcggtgcctcgtcgtggtggcagttcaacaactgggtggaccgcccc
MxDZ2_Nan_RS17045     gcgggcctgggcttcggtgcctcgtcgtggtggcagttcaacaactgggtggaccgcccc
MxDZ2_Tam_RS25820     gcgggcctgggcttcggtgcctcgtcgtggtggcagttcaacaactgggtggaccgcccc
                      ************************************************************


MxDZ2_Kirby_RS0232095 atggccaccttcagcatcgccaactcggtggtgggcggcatgttcaccgcgggcctgatg
MxDZ2_Nan_RS17045     atggccaccttcagcatcgccaactcggtggtgggcggcatgttcaccgcgggcctgatg
MxDZ2_Tam_RS25820     atggccaccttcagcatcgccaactcggtggtgggcggcatgttcaccgcgggcctgatg
                      ************************************************************

MxDZ2_Kirby_RS0232095 gacctcttcacgcaggacgcgggcgtcctgacgtggtccgccttcctgggcgcggagctg
MxDZ2_Nan_RS17045     gacctcttcacgcaggacgcgggcgtcctgacgtggtccgccttcctgggcgcggagctg
MxDZ2_Tam_RS25820     gacctcttcacgcaggacgcgggcgtcctgacgtggtccgccttcctgggcgcggagctg
                      ************************************************************

MxDZ2_Kirby_RS0232095 ggcgcatggctcacggcgggtatcggcggtggccagcttccgctcaacgacggcttgctg
MxDZ2_Nan_RS17045     ggcgcatggctcacggcgggtatcggcggtggccagcttccgctcaacgacggcttgctg
MxDZ2_Tam_RS25820     ggcgcatggctcacggcgggtatcggcggtggccagcttccgctcaacgacggcttgctg
                      ************************************************************

MxDZ2_Kirby_RS0232095 atttcgtccggtgcgggctggggcgcggcctactcggcgctgctgttggccatcatccac
MxDZ2_Nan_RS17045     atttcgtccggtgcgggctggggcgcggcctactcggcgctgctgttggccatcatccac
MxDZ2_Tam_RS25820     atttcgtccggtgcgggctggggcgcggcctactcggcgctgctgttggccatcatccac
                      ************************************************************

MxDZ2_Kirby_RS0232095 ttctccgggacgcccgtgtccggcaagacgtggcgggacacgctgctgctggctccgggc
MxDZ2_Nan_RS17045     ttctccgggacgcccgtgtccggcaagacgtggcgggacacgctgctgctggctccgggc
MxDZ2_Tam_RS25820     ttctccgggacgcccgtgtccggcaagacgtggcgggacacgctgctgctggctccgggc
                      ************************************************************

MxDZ2_Kirby_RS0232095 atcggcgcgggcgcgctggcgctggccaccatgcgttacaacccgacgccgtcccaaatc
MxDZ2_Nan_RS17045     atcggcgcgggcgcgctggcgctggccaccatgcgttacaacccgacgccgtcccaaatc
MxDZ2_Tam_RS25820     atcggcgcgggcgcgctggcgctggccaccatgcgttacaacccgacgccgtcccaaatc
                      ************************************************************

MxDZ2_Kirby_RS0232095 gtccgagccaacatcttcggcggtggcgtgggtgcggtggtgctgctcctctccggcgtg
MxDZ2_Nan_RS17045     gtccgagccaacatcttcggcggtggcgtgggtgcggtggtgctgctcctctccggcgtg
MxDZ2_Tam_RS25820     gtccgagccaacatcttcggcggtggcgtgggtgcggtggtgctgctcctctccggcgtg
                      ************************************************************

MxDZ2_Kirby_RS0232095 gtgctgggtggtttcgatcagtccacgccgtacgtcctgtcgttcctcggttcggcgggg
MxDZ2_Nan_RS17045     gtgctgggtggtttcgatcagtccacgccgtacgtcctgtcgttcctcggttcggcgggg
MxDZ2_Tam_RS25820     gtgctgggtggtttcgatcagtccacgccgtacgtcctgtcgttcctcggttcggcgggg
                      ************************************************************


MxDZ2_Kirby_RS0232095 gccatcaccacggtgagcctgctgtgggaggagagcgtggaccgctccgccctgaagatg
MxDZ2_Nan_RS17045     gccatcaccacggtgagcctgctgtgggaggagagcgtggaccgctccgccctgaagatg
MxDZ2_Tam_RS25820     gccatcaccacggtgagcctgctgtgggaggagagcgtggaccgctccgccctgaagatg
                      ************************************************************

MxDZ2_Kirby_RS0232095 gctggccgctcagagaaggaccggccgtacacgaacgtttggtggtag
MxDZ2_Nan_RS17045     gctggccgctcagagaaggaccggccgtacacgaacgtttggtggtag
MxDZ2_Tam_RS25820     gctggccgctcagagaaggaccggccgtacacgaacgtttggtggtag
                      ************************************************
```

**Non-coding region mutation in MxDZ2_Nan 5**

| **Tam_location** | **Tam_nucleotide** | **Kirby nucleotide** | **Nan_nucleotide** | **Nan_location** |
| --- | --- | --- | --- | --- |
| **7087101** | **G (complement)** | **G (complement)** | **.** | **7087072** |

```
multiple sequence alignment (nucleotide)

MxDZ2_Kirby_RS0206555 atggatgcattggacgtgttgaaccaggagcaccgccacatccagcgcgtgctggtggtc
MxDZ2_Nan_RS10175     atggatgcattggacgtgttgaaccaggagcaccgccacatccagcgcgtgctggtggtc
MxDZ2_Tam_RS32670     atggatgcattggacgtgttgaaccaggagcaccgccacatccagcgcgtgctggtggtc
                      ************************************************************

MxDZ2_Kirby_RS0206555 ctggatcgggcggtggcgaagggccgcgatggggagttcgtctcggcctcgctcttcctg
MxDZ2_Nan_RS10175     ctggatcgggcggtggcgaagggccgcgatggggagttcgtctcggcctcgctcttcctg
MxDZ2_Tam_RS32670     ctggatcgggcggtggcgaagggccgcgatggggagttcgtctcggcctcgctcttcctg
                      ************************************************************

MxDZ2_Kirby_RS0206555 cgcgccgcgaacttcttcctcaccttcgtggacggcagccaccacgcgaaggagatggtg
MxDZ2_Nan_RS10175     cgcgccgcgaacttcttcctcaccttcgtggacggcagccaccacgcgaaggagatggtg
MxDZ2_Tam_RS32670     cgcgccgcgaacttcttcctcaccttcgtggacggcagccaccacgcgaaggagatggtg
                      ************************************************************

MxDZ2_Kirby_RS0206555 ctcttccagacgatggtggcgcaccggctgccgctggcgcccgggctgctggcccaggtg
MxDZ2_Nan_RS10175     ctcttccagacgatggtggcgcaccggctgccgctggcgcccgggctgctggcccaggtg
MxDZ2_Tam_RS32670     ctcttccagacgatggtggcgcaccggctgccgctggcgcccgggctgctggcccaggtg
                      ************************************************************

MxDZ2_Kirby_RS0206555 tccggcgagcacggcaccggcagcgagcacgcggtggccatgctctgcgcggcggaggcg
MxDZ2_Nan_RS10175     tccggcgagcacggcaccggcagcgagcacgcggtggccatgctctgcgcggcggaggcg
MxDZ2_Tam_RS32670     tccggcgagcacggcaccggcagcgagcacgcggtggccatgctctgcgcggcggaggcg
                      ************************************************************

MxDZ2_Kirby_RS0206555 atgctgcgtggcggggagacggaccccgcgcgcatgctcgacgcggcggatgactggttg
MxDZ2_Nan_RS10175     atgctgcgtggcggggagacggaccccgcgcgcatgctcgacgcggcggatgactggttg
MxDZ2_Tam_RS32670     atgctgcgtggcggggagacggaccccgcgcgcatgctcgacgcggcggatgactggttg
                      ************************************************************

MxDZ2_Kirby_RS0206555 cgcctgtaccgggggcacaccgcggtggaggaggcgcaggtgttccccatggcgcggcgg
MxDZ2_Nan_RS10175     cgcctgtaccgggggcacaccgcggtggaggaggcgcaggtgttccccatggcgcggcgg
MxDZ2_Tam_RS32670     cgcctgtaccgggggcacaccgcggtggaggaggcgcaggtgttccccatggcgcggcgg
                      ************************************************************

MxDZ2_Kirby_RS0206555 ctgctgcccgcgggaatcctggaccggatgcggacgcgcttcgcccgaatcgaggcgtcg
MxDZ2_Nan_RS10175     ctgctgcccgcgggaatcctggaccggatgcggacgcgcttcgcccgaatcgaggcgtcg
MxDZ2_Tam_RS32670     ctgctgcccgcgggaatcctggaccggatgcggacgcgcttcgcccgaatcgaggcgtcg
                      ************************************************************

MxDZ2_Kirby_RS0206555 catggctcgctggcggatgccgcggatgccatggagcgcgccttcacgcCccccccggcg
MxDZ2_Nan_RS10175     catggctcgctggcggatgccgcggatgccatggagcgcgccttcacgc-ccccccggcg
MxDZ2_Tam_RS32670     catggctcgctggcggatgccgcggatgccatggagcgcgccttcacgcCccccccggcg
                      ************************************************* **********

MxDZ2_Kirby_RS0206555 ggacgtctcttcccgcgcggaccggtgtgctcgttctag
MxDZ2_Nan_RS10175     ggacgtctcttcccgcgcggaccggtgtgctcgttctag
MxDZ2_Tam_RS32670     ggacgtctcttcccgcgcggaccggtgtgctcgttctag
                      ***************************************
```

**Non-coding region mutation in MxDZ2_Nan 6**

| **Tam_location** | **Tam_nucleotide** | **Kirby nucleotide** | **Nan_nucleotide** | **Nan_location** |
| --- | --- | --- | --- | --- |
| **8584270** | **C (complement)** | **C (complement)** | **.** | **8584239** |

```
multiple sequence alignment (nucleotide)

MxDZ2_Kirby_RS0209420 atgaagccagagcaagacgccttgttccctcccgcgccagtctccgtggccacctccgGg
MxDZ2_Nan_RS03855     atgaagccagagcaagacgccttgttccctcccgcgccagtctccgtggccacctccg-g
MxDZ2_Tam_RS01140     atgaagccagagcaagacgccttgttccctcccgcgccagtctccgtggccacctccgGg
                      ********************************************************** *

MxDZ2_Kirby_RS0209420 ggggagccccgattcggtacctaccagggtgagttgcccgaagtggacctgccgcgcctg
MxDZ2_Nan_RS03855     ggggagccccgattcggtacctaccagggtgagttgcccgaagtggacctgccgcgcctg
MxDZ2_Tam_RS01140     ggggagccccgattcggtacctaccagggtgagttgcccgaagtggacctgccgcgcctg
                      ************************************************************

MxDZ2_Kirby_RS0209420 caagggcagtgggccccgggccgcaccacgcgtctgctgaagcgcaagcgttggcactac
MxDZ2_Nan_RS03855     caagggcagtgggccccgggccgcaccacgcgtctgctgaagcgcaagcgttggcactac
MxDZ2_Tam_RS01140     caagggcagtgggccccgggccgcaccacgcgtctgctgaagcgcaagcgttggcactac
                      ************************************************************

MxDZ2_Kirby_RS0209420 accttcgcggccacgcccgaggtcgccgcgctcttcgcggtggtggacctgggctacacg
MxDZ2_Nan_RS03855     accttcgcggccacgcccgaggtcgccgcgctcttcgcggtggtggacctgggctacacg
MxDZ2_Tam_RS01140     accttcgcggccacgcccgaggtcgccgcgctcttcgcggtggtggacctgggctacacg
                      ************************************************************

MxDZ2_Kirby_RS0209420 tcgagcgccttcgcggtggccctggacttgcgcgagcgcaagccgctgtgtgacgtgagc
MxDZ2_Nan_RS03855     tcgagcgccttcgcggtggccctggacttgcgcgagcgcaagccgctgtgtgacgtgagc
MxDZ2_Tam_RS01140     tcgagcgccttcgcggtggccctggacttgcgcgagcgcaagccgctgtgtgacgtgagc
                      ************************************************************

MxDZ2_Kirby_RS0209420 ttcctgggcacgcccggccccatggtggtgctgggcgacaagcccggtcagggcctcaag
MxDZ2_Nan_RS03855     ttcctgggcacgcccggccccatggtggtgctgggcgacaagcccggtcagggcctcaag
MxDZ2_Tam_RS01140     ttcctgggcacgcccggccccatggtggtgctgggcgacaagcccggtcagggcctcaag
                      ************************************************************

MxDZ2_Kirby_RS0209420 gcctccttccgcacgctgggcggacggctgtcggtgcagcgcggcgaggacgacgagcgc
MxDZ2_Nan_RS03855     gcctccttccgcacgctgggcggacggctgtcggtgcagcgcggcgaggacgacgagcgc
MxDZ2_Tam_RS01140     gcctccttccgcacgctgggcggacggctgtcggtgcagcgcggcgaggacgacgagcgc
                      ************************************************************

MxDZ2_Kirby_RS0209420 taccgggtggtggtggacgtcagccggctgcgcaccggaagccttcagtccttccagtgg
MxDZ2_Nan_RS03855     taccgggtggtggtggacgtcagccggctgcgcaccggaagccttcagtccttccagtgg
MxDZ2_Tam_RS01140     taccgggtggtggtggacgtcagccggctgcgcaccggaagccttcagtccttccagtgg
                      ************************************************************

MxDZ2_Kirby_RS0209420 gagggcgacatgatggtggccggcggcccgcccgcgctcaccgtggtggcgccggtggag
MxDZ2_Nan_RS03855     gagggcgacatgatggtggccggcggcccgcccgcgctcaccgtggtggcgccggtggag
MxDZ2_Tam_RS01140     gagggcgacatgatggtggccggcggcccgcccgcgctcaccgtggtggcgccggtggag
                      ************************************************************

MxDZ2_Kirby_RS0209420 ggtgacgggctggtcaacgtcaccatgaagagcaatggcctgctcaccttcggcagcctg
MxDZ2_Nan_RS03855     ggtgacgggctggtcaacgtcaccatgaagagcaatggcctgctcaccttcggcagcctg
MxDZ2_Tam_RS01140     ggtgacgggctggtcaacgtcaccatgaagagcaatggcctgctcaccttcggcagcctg
                      ************************************************************

MxDZ2_Kirby_RS0209420 gaggcgggcgggaagcgcttccggttggacggaggcgtgggcggcatggactacacgcag
MxDZ2_Nan_RS03855     gaggcgggcgggaagcgcttccggttggacggaggcgtgggcggcatggactacacgcag
MxDZ2_Tam_RS01140     gaggcgggcgggaagcgcttccggttggacggaggcgtgggcggcatggactacacgcag
                      ************************************************************

MxDZ2_Kirby_RS0209420 ggctacctggcgcggcacaccgcgtggcgctgggccttcgcgtcggggcggctggcggac
MxDZ2_Nan_RS03855     ggctacctggcgcggcacaccgcgtggcgctgggccttcgcgtcggggcggctggcggac
MxDZ2_Tam_RS01140     ggctacctggcgcggcacaccgcgtggcgctgggccttcgcgtcggggcggctggcggac
                      ************************************************************

MxDZ2_Kirby_RS0209420 ggcacgccggtgggcatcaacctcgtggagggcttcaacgagggcaacgcggacgccaac
MxDZ2_Nan_RS03855     ggcacgccggtgggcatcaacctcgtggagggcttcaacgagggcaacgcggacgccaac
MxDZ2_Tam_RS01140     ggcacgccggtgggcatcaacctcgtggagggcttcaacgagggcaacgcggacgccaac
                      ************************************************************

MxDZ2_Kirby_RS0209420 gagaacgcgctgtggctgggggacaagctgtacccgctggcgcgcgcccgcttcgaatac
MxDZ2_Nan_RS03855     gagaacgcgctgtggctgggggacaagctgtacccgctggcgcgcgcccgcttcgaatac
MxDZ2_Tam_RS01140     gagaacgcgctgtggctgggggacaagctgtacccgctggcgcgcgcccgcttcgaatac
                      ************************************************************

MxDZ2_Kirby_RS0209420 gacgcgaaggacctgatggcgccgtggcggctggcgacggtggatggcgcggtggacctg
MxDZ2_Nan_RS03855     gacgcgaaggacctgatggcgccgtggcggctggcgacggtggatggcgcggtggacctg
MxDZ2_Tam_RS01140     gacgcgaaggacctgatggcgccgtggcggctggcgacggtggatggcgcggtggacctg
                      ************************************************************


MxDZ2_Kirby_RS0209420 cgcttccagccccttcatgtgcaccgcgaggaccggaacctgcgcctcgtcgtgagccac
MxDZ2_Nan_RS03855     cgcttccagccccttcatgtgcaccgcgaggaccggaacctgcgcctcgtcgtgagccac
MxDZ2_Tam_RS01140     cgcttccagccccttcatgtgcaccgcgaggaccggaacctgcgcctcgtcgtgagccac
                      ************************************************************

MxDZ2_Kirby_RS0209420 ttcgcgcagccggtgggcttcttcaacggcacggtgcgcgtgggaagccgcacgctggag
MxDZ2_Nan_RS03855     ttcgcgcagccggtgggcttcttcaacggcacggtgcgcgtgggaagccgcacgctggag
MxDZ2_Tam_RS01140     ttcgcgcagccggtgggcttcttcaacggcacggtgcgcgtgggaagccgcacgctggag
                      ************************************************************

MxDZ2_Kirby_RS0209420 ttgtccaacgtccccggcgtcaccgaggaccaggacatgctctggtga
MxDZ2_Nan_RS03855     ttgtccaacgtccccggcgtcaccgaggaccaggacatgctctggtga
MxDZ2_Tam_RS01140     ttgtccaacgtccccggcgtcaccgaggaccaggacatgctctggtga
                      ************************************************
```

**Non-coding region mutation in MxDZ2_Tam 1**

| **Tam_location** | **Tam_nucleotide** | **Kirby nucleotide** | **Nan_nucleotide** | **Nan_location** |
| --- | --- | --- | --- | --- |
| **2292537** | **A** | **Contig boundary** |  | **2292526** |
| **2292543** | **T** | **Contig boundary** |  | **2292531** |
| **2292577** | **A** | **.** | **.** | **2292577** |
| **2292694** | **T** | **.** | **.** | **2292680** |

```
multiple sequence alignment (nucleotide)

MxDZ2_Tam_RS38160   gtgcccgtgccccaggggcgcctggtcgcgggcttcgatgtgggtcgcaccagagaccgc
MxDZ2_Nan_RS28750   gtgcccgtgccccaggggcgcctggtcgcgggcttcgatgtgggtcgcaccagagaccgc
MxDZ2_Kirby_RS39310 gtgcccgtgccccaggggcgcctggtcgcgggcttcgatgtgggtcgcaccagagaccgc
                    ************************************************************

MxDZ2_Tam_RS38160   tcggagctggccgtcttcgaggagacgggtgggcgcttcacgtgccgcatgctgcgcagc
MxDZ2_Nan_RS28750   tcggagctggccgtcttcgaggagacgggtgggcgcttcacgtgccgcatgctgcgcagc
MxDZ2_Kirby_RS39310 tcggagctggccgtcttcgaggagacgggtgggcgcttcacgtgccgcatgctgcgcagc
                    ************************************************************

MxDZ2_Tam_RS38160   ttcgagggcgtgcccttcgcggagcaggaggcgcacctccgtcgtctgctgtccgtgctt
MxDZ2_Nan_RS28750   ttcgagggcgtgcccttcgcggagcaggaggcgcacctccgtcgtctgctgtccgtgctt
MxDZ2_Kirby_RS39310 ttcgagggcgtgcccttcgcggagcaggaggcgcacctccgtcgtctgctgtccgtgctt
                    ************************************************************

MxDZ2_Tam_RS38160   cccgtggctcgtctcagcgtcgaccgcagcggcatcggaatgaatctcgcggagaacctc
MxDZ2_Nan_RS28750   cccgtggctcgtctcagcgtcgaccgcagcggcatcggaatgaatctcgcggagaacctc
MxDZ2_Kirby_RS39310 cccgtggctcgtctcagcgtcgaccgcagcggcatcggaatgaatctcgcggagaacctc
                    ************************************************************

MxDZ2_Tam_RS38160   gcccgcgactacccccaggtggtggaggagaacttcaccaacgaggccaaggagcgctgg
MxDZ2_Nan_RS28750   gcccgcgactacccccaggtggtggaggagaacttcaccaacgaggccaaggagcgctgg
MxDZ2_Kirby_RS39310 gcccgcgactacccccaggtggtggaggagaacttcaccaacgaggccaaggagcgctgg
                    ************************************************************

MxDZ2_Tam_RS38160   gccaccgacttcaagattctcctccagcgccgcgatgtcacgctgccgcgccagctgtga
MxDZ2_Nan_RS28750   gccaccgacttcaagattctcctccagcgccgcgatgtcacgctgccgcgccagc-gtga
MxDZ2_Kirby_RS39310 gccaccgacttcaagattctcctccagcgccgcgatgtcacgctgccgcgccagc-gtga
                    ******************************************************* ****


MxDZ2_Tam_RS38160   gcttgtcgggcaAgattcactccatcaagcgtcgcgtgctgccctcggggaaggtgtcctt
MxDZ2_Nan_RS28750   gcttgtcgggca-gattcactccatcaagcgtcgcgtgctgccctcggggaaggtgtcctt
MxDZ2_Kirby_RS39310 gcttgtcgggca-gattcactccatcaagcgtcgcgtgctgccctcggggaaggtgtcctt
                    ************ ************************************************

MxDZ2_Tam_RS38160   cgacgccgaacgcacctcccgcggccatgcggacaaattctgggccgtggcccttgcctg
MxDZ2_Nan_RS28750   cgacgccgaacgcacctcccgcggccatgcggacaaattctgggccgtggcccttgcctg
MxDZ2_Kirby_RS39310 cgacgccgaacgcacctcccgcggccatgcggacaaattctgggccgtggcccttgcctg
                    ************************************************************

MxDZ2_Tam_RS38160   ccagcgggTagcgccaggccgacaagccgagggcaggcgaaaAttgggTgtgcgtgtgattgg
MxDZ2_Nan_RS28750   ccagcggg-agcgccaggccgacaagccgagggcaggcgaaa-ttggg-gtgcgtgtgattgg
MxDZ2_Kirby_RS39310 ccagcggg-agcgc-------------------------------------------------
                    ******** *****

MxDZ2_Tam_RS38160   ctaa
MxDZ2_Nan_RS28750   ctaa
MxDZ2_Kirby_RS39310 ----
```

**The non-coding region lies between the two mentioned locus tags.**

**Non-coding region mutation 1**

| **Tam_location** | **Tam_nucleotide** | **Kirby nucleotide** | **Nan_nucleotide** | **Nan_location** |
| --- | --- | --- | --- | --- |
| **277103** | **C** | **C** | **.** | **277102** |

```
multiple sequence alignment (nucleotide)

Tam__RS05505_RS05510          cggcccgcagcgtatccgccgcatcgtcaccgcgctcgttcacctgagtctggaaataCc
Kirby__RS0208105_RS0208110    cggcccgcagcgtatccgccgcatcgtcaccgcgctcgttcacctgagtctggaaataCc
Nan__RS37365_RS37370          cggcccgcagcgtatccgccgcatcgtcaccgcgctcgttcacctgagtctggaaata-c
                              ********************************************************** *

Tam__RS05505_RS05510          cccggtatcagtcctgtcgagggttcgcacacccggtgaggtctcctgcgattcatcaag
Kirby__RS0208105_RS0208110    cccggtatcagtcctgtcgagggttcgcacacccggtgaggtctcctgcgattcatcaag
Nan__RS37365_RS37370          cccggtatcagtcctgtcgagggttcgcacacccggtgaggtctcctgcgattcatcaag
                              ************************************************************

Tam__RS05505_RS05510          tcctgtatttccagatagatatgaagcgcggtgaaccgcttctgcgcacgttgaaaacat
Kirby__RS0208105_RS0208110    tcctgtatttccagatagatatgaagcgcggtgaaccgcttctgcgcacgttgaaaacat
Nan__RS37365_RS37370          tcctgtatttccagatagatatgaagcgcggtgaaccgcttctgcgcacgttgaaaacat
                              ************************************************************

Tam__RS05505_RS05510          gaaccaaccctagagggcgggacagcgtgcaggactcgacggaaccgtaggaaagtcgcc
Kirby__RS0208105_RS0208110    gaaccaaccctagagggcgggacagcgtgcaggactcgacggaaccgtaggaaagtcgcc
Nan__RS37365_RS37370          gaaccaaccctagagggcgggacagcgtgcaggactcgacggaaccgtaggaaagtcgcc
                              ************************************************************

Tam__RS05505_RS05510          gcacaacgtgtcaggtcctcctgcgacccaggtgactgcatgggtccggaaggcatacgg
Kirby__RS0208105_RS0208110    gcacaacgtgtcaggtcctcctgcgacccaggtgactgcatgggtccggaaggcatacgg
Nan__RS37365_RS37370          gcacaacgtgtcaggtcctcctgcgacccaggtgactgcatgggtccggaaggcatacgg
                              ************************************************************

Tam__RS05505_RS05510          aggcctctcactcgacgttttccgtcgagaaaaggcatcggccgcgcgtcgtggatggac
Kirby__RS0208105_RS0208110    aggcctctcactcgacgttttccgtcgagaaaaggcatcggccgcgcgtcgtggatggac
Nan__RS37365_RS37370          aggcctctcactcgacgttttccgtcgagaaaaggcatcggccgcgcgtcgtggatggac
                              ************************************************************


Tam__RS05505_RS05510          tcgcgaagtccaccgcgtgcacagcaaccctatgggtgaacaattggaggctgacccggc
Kirby__RS0208105_RS0208110    tcgcgaagtccaccgcgtgcacagcaaccctatgggtgaacaattggaggctgacccggc
Nan__RS37365_RS37370          tcgcgaagtccaccgcgtgcacagcaaccctatgggtgaacaattggaggctgacccggc
                              ************************************************************

Tam__RS05505_RS05510          aagttctgtctgtctgcccccatacgtcctatgccagcggccaaacgttccgggtcggta
Kirby__RS0208105_RS0208110    aagttctgtctgtctgcccccatacgtcctatgccagcggccaaacgttccgggtcggta
Nan__RS37365_RS37370          aagttctgtctgtctgcccccatacgtcctatgccagcggccaaacgttccgggtcggta
                              ************************************************************

Tam__RS05505_RS05510          aggc
Kirby__RS0208105_RS0208110    aggc
Nan__RS37365_RS37370          aggc
                              ****
```

**Non-coding region mutation 2**

| **Tam_location** | **Tam_nucleotide** | **Kirby nucleotide** | **Nan_nucleotide** | **Nan_location** |
| --- | --- | --- | --- | --- |
| **1755694** | **C** | **C** | **.** | **1755690** |

```
multiple sequence alignment (nucleotide)

Tam__RS11755_RS11760          cagaggcgcctccgagtagcccgagcgtcagcaggactggctggaggggctggcggaggc
Kirby__RS0204160_RS0204170    cagaggcgcctccgagtagcccgagcgtcagcaggactggctggaggggctggcggaggc
Nan__RS31120_RS31125          cagaggcgcctccgagtagcccgagcgtcagcaggactggctggaggggctggcggaggc
                              ************************************************************

Tam__RS11755_RS11760          cttccggcacttcggcggcgtgccgtgcgtcgtgctcggcgacgatgcccgcgcgcacgt
Kirby__RS0204160_RS0204170    cttccggcacttcggcggcgtgccgtgcgtcgtgctcggcgacgatgcccgcgcgcacgt
Nan__RS31120_RS31125          cttccggcacttcggcggcgtgccgtgcgtcgtgctcggcgacgatgcccgcgcgcacgt
                              ************************************************************

Tam__RS11755_RS11760          cgagggtcgcaccggcttcctgcttgagctgaaggagtagggcgcatggccgagaggtga
Kirby__RS0204160_RS0204170    cgagggtcgcaccggcttcctgcttgagctgaaggagtagggcgcatggccgagaggtga
Nan__RS31120_RS31125          cgagggtcgcaccggcttcctgcttgagctgaaggagtagggcgcatggccgagaggtga
                              ************************************************************

Tam__RS11755_RS11760          gagcacgcctggcctcgccacgcgtgggggcaggcgtccgcgctttccgggtccgcgacg
Kirby__RS0204160_RS0204170    gagcacgcctggcctcgccacgcgtgggggcaggcgtccgcgctttccgggtccgcgacg
Nan__RS31120_RS31125          gagcacgcctggcctcgccacgcgtgggggcaggcgtccgcgctttccgggtccgcgacg
                              ************************************************************

Tam__RS11755_RS11760          ggccattggagtcgccatggcagccctctccgctctctccactgccagcagatgccagac
Kirby__RS0204160_RS0204170    ggccattggagtcgccatggcagccctctccgctctctccactgccagcagatgccagac
Nan__RS31120_RS31125          ggccattggagtcgccatggcagccctctccgctctctccactgccagcagatgccagac
                              ************************************************************

Tam__RS11755_RS11760          cccgggtattgccgcaagCccccccggtcgtgctccaccctcgcccgtttcaatccagga
Kirby__RS0204160_RS0204170    cccgggtattgccgcaagCccccccggtcgtgctccaccctcgcccgtttcaatccagga
Nan__RS31120_RS31125          cccgggtattgccgcaag-ccccccggtcgtgctccaccctcgcccgtttcaatccagga
                              ****************** *****************************************

Tam__RS11755_RS11760          gcgggagggccgagaa
Kirby__RS0204160_RS0204170    gcgggagggccgagaa
Nan__RS31120_RS31125          gcgggagggccgagaa
                              ****************
```

**Non-coding region mutation 3**

| **Tam_location** | **Tam_nucleotide** | **Kirby nucleotide** | **Nan_nucleotide** | **Nan_location** |
| --- | --- | --- | --- | --- |
| **2104550** | **G** | **G** | **.** | **2104542** |
| **2104551** | **C** | **C** | **.** | **2104542** |

```
multiple sequence alignment (nucleotide)

Tam__RS13130_RS38120          gcgggacgtcgggagtccctcgttcccggggggcatctccactggactcctgctcgcctt
Kirby__RS41580_RS0202725      gcgggacgtcgggagtccctcgttcccggggggcatctccactggactcctgctcgcctt
Nan__RS29740_RS29745          gcgggacgtcgggagtccctcgttcccggggggcatctccactggactcctgctcgcctt
                              ************************************************************

Tam__RS13130_RS38120          gacgcactgcccactctgaaatcggtgtacccgtgaagcagcgagcgacatttccgtcaa
Kirby__RS41580_RS0202725      gacgcactgcccactctgaaatcggtgtacccgtgaagcagcgagcgacatttccgtcaa
Nan__RS29740_RS29745          gacgcactgcccactctgaaatcggtgtacccgtgaagcagcgagcgacatttccgtcaa
                              ************************************************************

Tam__RS13130_RS38120          cgttgcgcggcgtgtttgtttctgaacggcggagtgagtacaggggctgggcggacacgg
Kirby__RS41580_RS0202725      cgttgcgcggcgtgtttgtttctgaacggcggagtgagtacaggggctgggcggacacgg
Nan__RS29740_RS29745          cgttgcgcggcgtgtttgtttctgaacggcggagtgagtacaggggctgggcggacacgg
                              ************************************************************

Tam__RS13130_RS38120          aggcgctccctggtgtagaccgagcctgaacgagaggaggggccgttgggattcgcttcg
Kirby__RS41580_RS0202725      aggcgctccctggtgtagaccgagcctgaacgagaggaggggccgttgggattcgcttcg
Nan__RS29740_RS29745          aggcgctccctggtgtagaccgagcctgaacgagaggaggggccgttgggattcgcttcg
                              ************************************************************

Tam__RS13130_RS38120          atgaccagcggctccctgtgactatgccgacgccgccacgcgcatccgtattgctcctgg
Kirby__RS41580_RS0202725      atgaccagcggctccctgtgactatgccgacgccgccacgcgcatccgtattgctcctgg
Nan__RS29740_RS29745          atgaccagcggctccctgtgactatgccgacgccgccacgcgcatccgtattgctcctgg
                              ************************************************************

Tam__RS13130_RS38120          aggactacccagacctccgggcgtagtggccttctttgcgcaaaccgatggatttcgacg
Kirby__RS41580_RS0202725      aggactacccagacctccgggcgtagtggccttctttgcgcaaaccgatggatttcgacg
Nan__RS29740_RS29745          aggactacccagacctccgggcgtagtggccttctttgcgcaaaccgatggatttcgacg
                              ************************************************************

Tam__RS13130_RS38120          aactcctcgaactgatcgctcagcactgcccgcgagcgagcacataggcaggtttcgttg
Kirby__RS41580_RS0202725      aactcctcgaactgatcgctcagcactgcccgcgagcgagcacataggcaggtttcgttg
Nan__RS29740_RS29745          aactcctcgaactgatcgctcagcactgcccgcgagcgagcacataggcaggtttcgttg
                              ************************************************************

Tam__RS13130_RS38120          tgcgatgggatgacgcgtcttctctcgtgggaatgagtttgcccgttgtactgctgctgc
Kirby__RS41580_RS0202725      tgcgatgggatgacgcgtcttctctcgtgggaatgagtttgcccgttgtactgctgctgc
Nan__RS29740_RS29745          tgcgatgggatgacgcgtcttctctcgtgggaatgagtttgcccgttgtactgctgctgc
                              ************************************************************

Tam__RS13130_RS38120          agaatgtcgcgcttgcccccccGCgagtggctgctggagtggcggagttgctgacgcccc
Kirby__RS41580_RS0202725      agaatgtcgcgcttgcccccccGCgagtggctgctggagtggcggagttgctgacgcccc
Nan__RS29740_RS29745          agaatgtcgcgcttgccccccc--gagtggctgctggagtggcggagttgctgacgcccc
                              **********************  ************************************

Tam__RS13130_RS38120          gaggc
Kirby__RS41580_RS0202725      gaggc
Nan__RS29740_RS29745          gaggc
                              *****
```

**Non-coding region mutation 4**

| **Tam_location** | **Tam_nucleotide** | **Kirby nucleotide** | **Nan_nucleotide** | **Nan_location** |
| --- | --- | --- | --- | --- |
| **5878114** | **C** | **C** | **.** | **5878087** |

```
multiple sequence alignment (nucleotide)

Tam__RS27845_RS27850          gggctctgctccttcccccggtatgacgggcccgtcctccagcttgttccaaccgcgcgc
Kirby__RS0206870_RS0206880    gggctctgctccttcccccggtatgacgggcccgtcctccagcttgttccaaccgcgcgc
Nan__RS15005_RS15010          gggctctgctccttcccccggtatgacgggcccgtcctccagcttgttccaaccgcgcgc
                              ************************************************************

Tam__RS27845_RS27850          cgtattcctctcacgacggttcattgccccacattCccccccgccgggcgttcaccgatg
Kirby__RS0206870_RS0206880    cgtattcctctcacgacggttcattgccccacattCccccccgccgggcgttcaccgatg
Nan__RS15005_RS15010          cgtattcctctcacgacggttcattgccccacatt-ccccccgccgggcgttcaccgatg
                              *********************************** ************************

Tam__RS27845_RS27850          gaagccttcacacgtggagcgtgacggatcggacgcttcacctgtcagaccccctctcta
Kirby__RS0206870_RS0206880    gaagccttcacacgtggagcgtgacggatcggacgcttcacctgtcagaccccctctcta
Nan__RS15005_RS15010          gaagccttcacacgtggagcgtgacggatcggacgcttcacctgtcagaccccctctcta
                              ************************************************************

Tam__RS27845_RS27850          ccttcatcggcatgcgactggaggtcgtccatgttcgacagttgtggaggatgtgagcgc
Kirby__RS0206870_RS0206880    ccttcatcggcatgcgactggaggtcgtccatgttcgacagttgtggaggatgtgagcgc
Nan__RS15005_RS15010          ccttcatcggcatgcgactggaggtcgtccatgttcgacagttgtggaggatgtgagcgc
                              ************************************************************

Tam__RS27845_RS27850          cgcggaggcgccggtgaagtgttcgactggtgcgcccactgcgagctgtccctgtgcccc
Kirby__RS0206870_RS0206880    cgcggaggcgccggtgaagtgttcgactggtgcgcccactgcgagctgtccctgtgcccc
Nan__RS15005_RS15010          cgcggaggcgccggtgaagtgttcgactggtgcgcccactgcgagctgtccctgtgcccc
                              ************************************************************

Tam__RS27845_RS27850          cgctgcatggagcgcggctgctgcgacaatgagcccgccgattccggccgctcggcgccc
Kirby__RS0206870_RS0206880    cgctgcatggagcgcggctgctgcgacaatgagcccgccgattccggccgctcggcgccc
Nan__RS15005_RS15010          cgctgcatggagcgcggctgctgcgacaatgagcccgccgattccggccgctcggcgccc
                              ************************************************************

Tam__RS27845_RS27850          cggtgcctgccagagcctcccgaggacgacgaggccacggcacttcctgagcacttcggc
Kirby__RS0206870_RS0206880    cggtgcctgccagagcctcccgaggacgacgaggccacggcacttcctgagcacttcggc
Nan__RS15005_RS15010          cggtgcctgccagagcctcccgaggacgacgaggccacggcacttcctgagcacttcggc
                              ************************************************************

Tam__RS27845_RS27850          ggccggtgctgcacccgcgcccgcgccgtggcctgctcctgcgccttccactgggtttgc
Kirby__RS0206870_RS0206880    ggccggtgctgcacccgcgcccgcgccgtggcctgctcctgcgccttccactgggtttgc
Nan__RS15005_RS15010          ggccggtgctgcacccgcgcccgcgccgtggcctgctcctgcgccttccactgggtttgc
                              ************************************************************

Tam__RS27845_RS27850          gctcaccacggcgatcagcacatcggcacgcatgactgacgttcccgttctccgctggga
Kirby__RS0206870_RS0206880    gctcaccacggcgatcagcacatcggcacgcatgactgacgttcccgttctccgctggga
Nan__RS15005_RS15010          gctcaccacggcgatcagcacatcggcacgcatgactgacgttcccgttctccgctggga
                              ************************************************************

Tam__RS27845_RS27850          acggatctccaggtgctgggctcgacacatccggctccgcccagaaagtgacgtttcacc
Kirby__RS0206870_RS0206880    acggatctccaggtgctgggctcgacacatccggctccgcccagaaagtgacgtttcacc
Nan__RS15005_RS15010          acggatctccaggtgctgggctcgacacatccggctccgcccagaaagtgacgtttcacc
                              ************************************************************

Tam__RS27845_RS27850          tgatattttccgggcaggctagattgctcagttcctgagagggtgtgaagatgcgcgtga
Kirby__RS0206870_RS0206880    tgatattttccgggcaggctagattgctcagttcctgagagggtgtgaagatgcgcgtga
Nan__RS15005_RS15010          tgatattttccgggcaggctagattgctcagttcctgagagggtgtgaagatgcgcgtga
                              ************************************************************

Tam__RS27845_RS27850          gacagatgtcgtgtctcacccccaccttgcgcccctccgaggagcctccc
Kirby__RS0206870_RS0206880    gacagatgtcgtgtctcacccccaccttgcgcccctccgaggagcctccc
Nan__RS15005_RS15010          gacagatgtcgtgtctcacccccaccttgcgcccctccgaggagcctccc
                              **************************************************
```

**Non-coding region mutation 5**

| **Tam_location** | **Tam_nucleotide** | **Kirby nucleotide** | **Nan_nucleotide** | **Nan_location** |
| --- | --- | --- | --- | --- |
| **8902777** | **G** | **G** | **.** | **8902745** |

```
multiple sequence alignment (nucleotide)

Tam__RS02590_RS02595          gtgacactcagggacgcgcggccgtggcaggggggcgcgtcccgcggccgtgcggtgaac
Kirby__RS0212165_RS0212170    gtgacactcagggacgcgcggccgtggcaggggggcgcgtcccgcggccgtgcggtgaac
Nan__RS02395_RS02400          gtgacactcagggacgcgcggccgtggcaggggggcgcgtcccgcggccgtgcggtgaac
                              ************************************************************

Tam__RS02590_RS02595          ccgagtcaagcgcgccagcccaccagtgacggcttccgtgcgctccaatgcccccGgggg
Kirby__RS0212165_RS0212170    ccgagtcaagcgcgccagcccaccagtgacggcttccgtgcgctccaatgcccccGgggg
Nan__RS02395_RS02400          ccgagtcaagcgcgccagcccaccagtgacggcttccgtgcgctccaatgccccc-gggg
                              ******************************************************* ****

Tam__RS02590_RS02595          tcgttccgcgtcgaaaggccccgacagaataatgaggcgcccacgcacatggtgaggacg
Kirby__RS0212165_RS0212170    tcgttccgcgtcgaaaggccccgacagaataatgaggcgcccacgcacatggtgaggacg
Nan__RS02395_RS02400          tcgttccgcgtcgaaaggccccgacagaataatgaggcgcccacgcacatggtgaggacg
                              ************************************************************
```
