## Supplementary figures and images for "Same strain, two genomes: Creating a consensus circular genome of *Myxococcus xanthus* DZ2 revealed the diversity of a large polyploid prophage region within the close-related myxobacterial strains"

### Supplementary Figure1

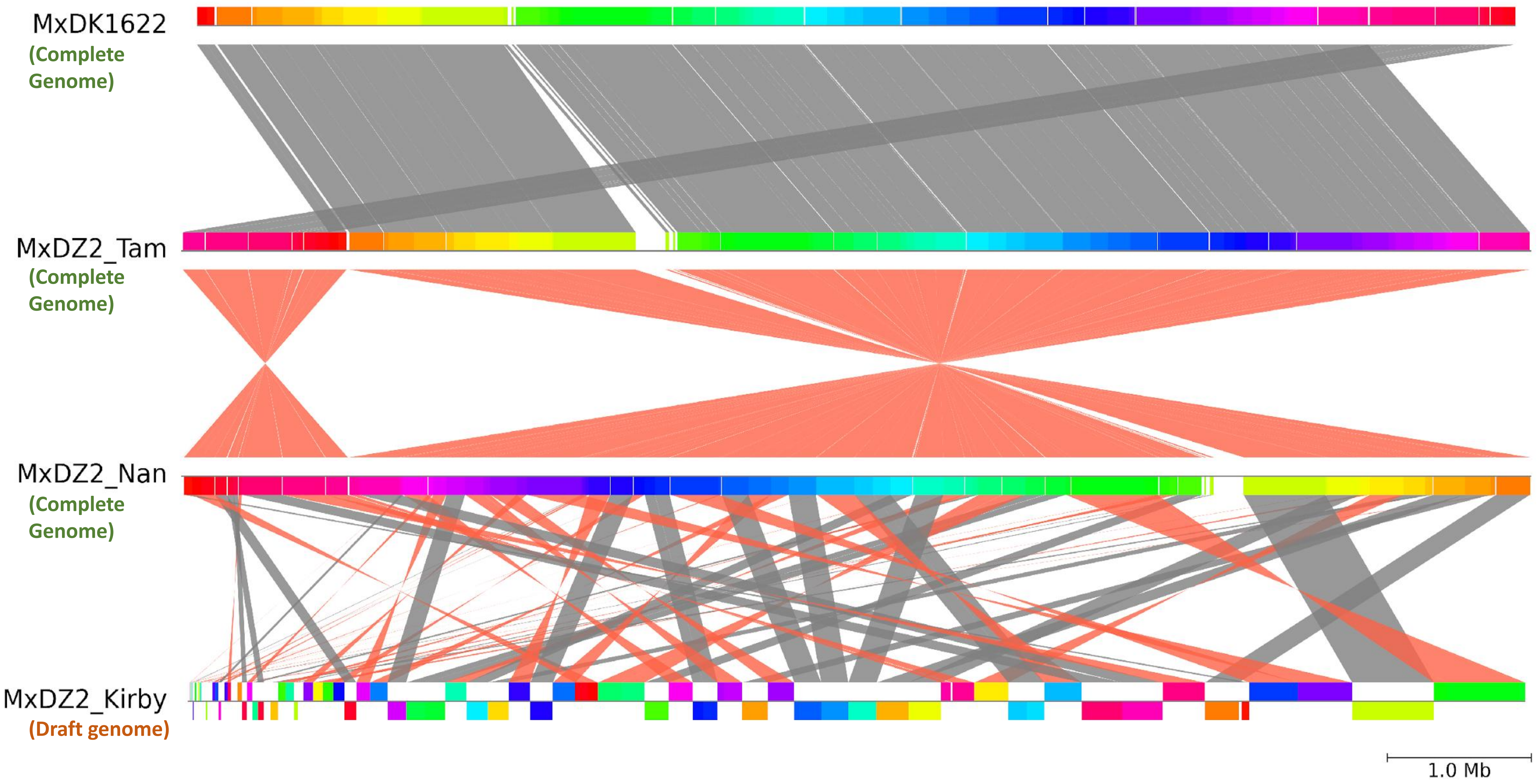

### Supplementary Figure 2

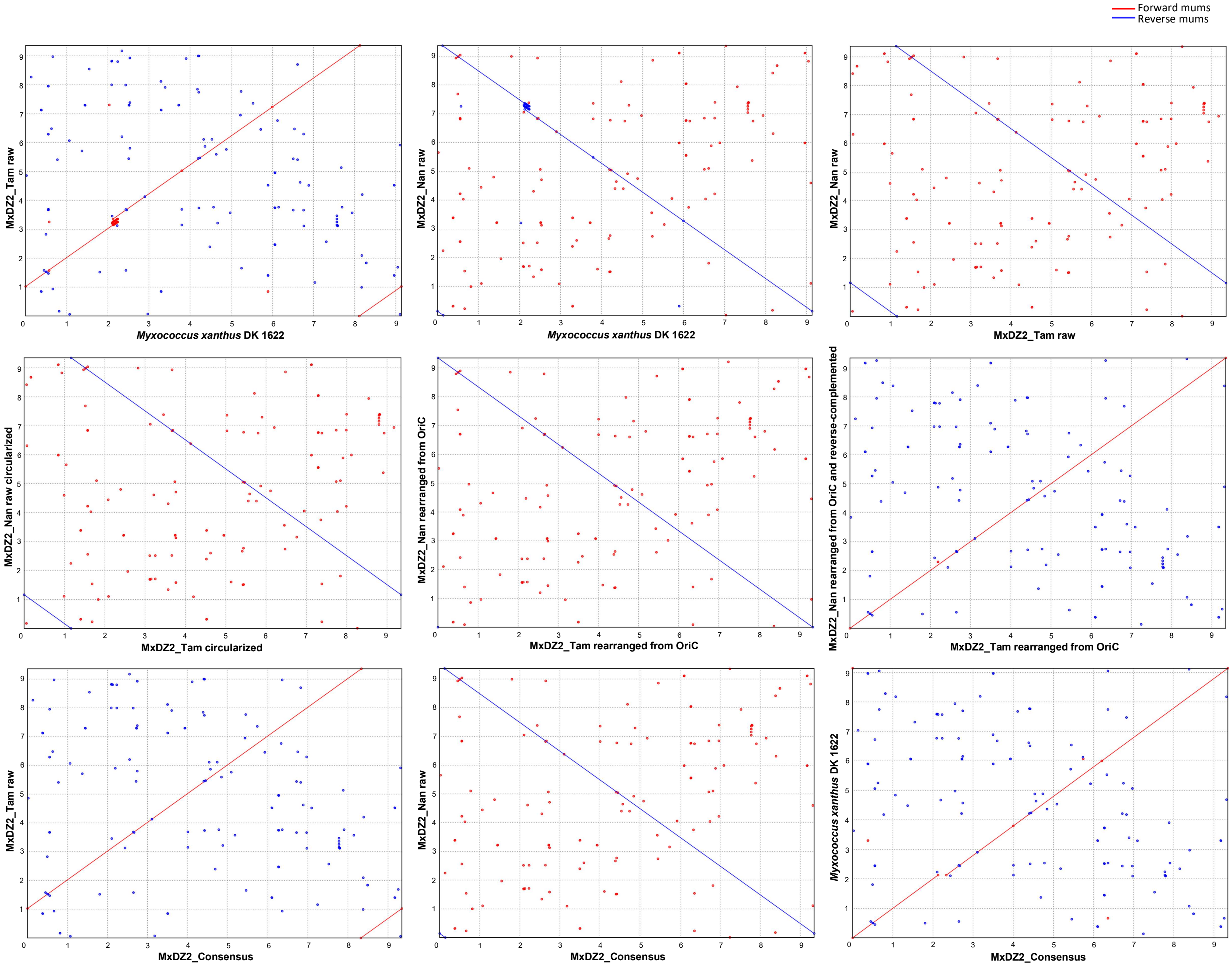

### Supplementary Figure 3

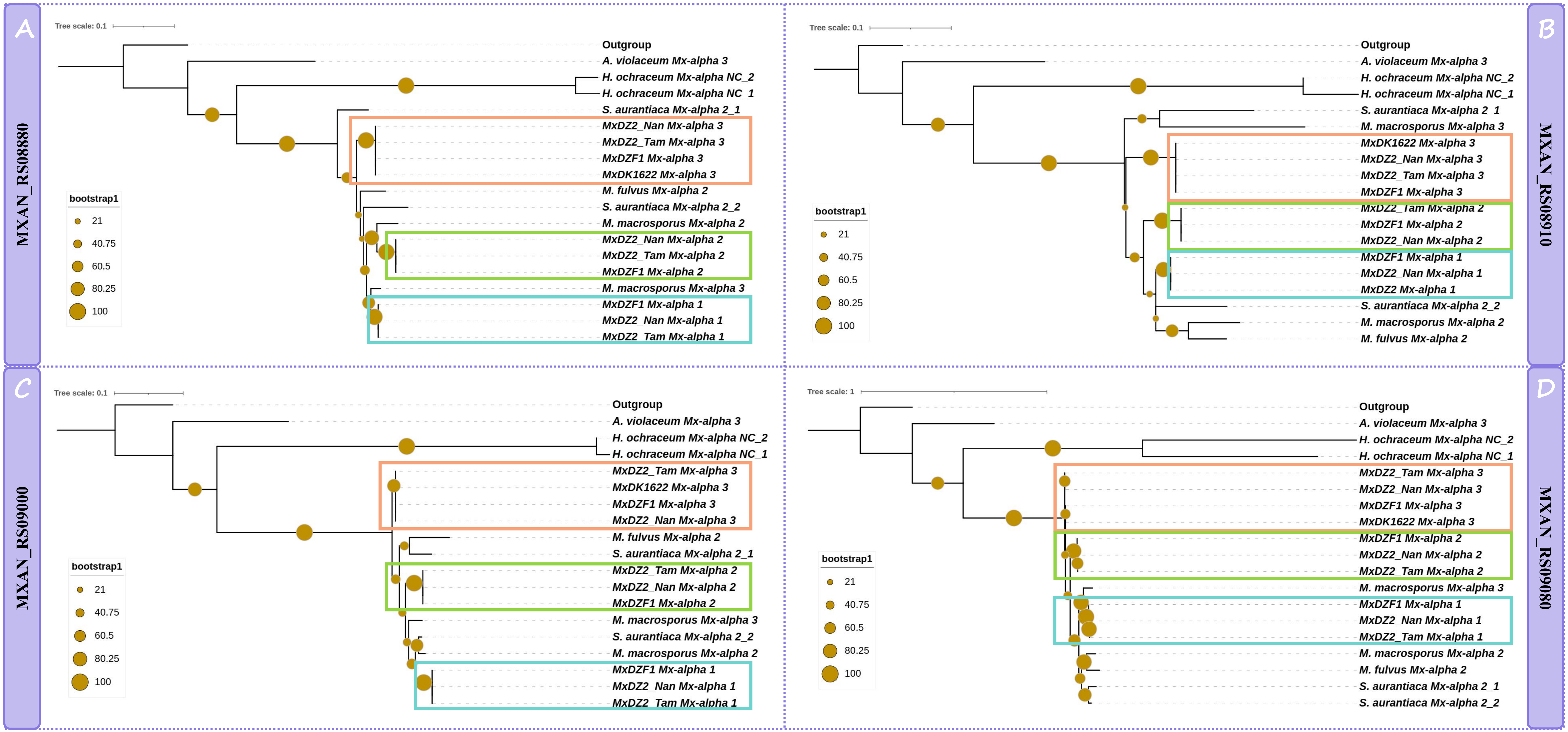
